# Analysis of Two-State Folding Using Parabolic Approximation III: Non-Arrhenius Kinetics of FBP28 WW Part-I

**DOI:** 10.1101/038331

**Authors:** Robert S. Sade

## Abstract

A model which treats the denatured and the native conformers as being confined to harmonic Gibbs energy wells has been used to analyse the non-Arrhenius behaviour of spontaneously-folding fixed two-state systems. The results demonstrate that when pressure and solvent are constant: (*i*) a two-state system is physically defined only for a finite temperature range; (*ii*) irrespective of the primary sequence, the 3-dimensional structure of the native conformer, the residual structure in the denatured state, and the magnitude of the folding and unfolding rate constants, the equilibrium stability of a two-state system is a maximum when its denatured conformers bury the least amount of solvent accessible surface area (SASA) to reach the activated state; (*iii*) the Gibbs barriers to folding and unfolding are not always due to the incomplete compensation of the activation enthalpies and entropies; (*iv*) the difference in heat capacity between the reaction-states is due to both the size of the solvent-shell and the non-covalent interactions; (*v*) the position of the transition state ensemble along the reaction coordinate (RC) depends on the choice of the RC; and (*vi*) the atomic structure of the transiently populated reaction-states cannot be inferred from perturbation-induced changes in their energetics.

## Introduction

It was shown elsewhere, henceforth referred to as Papers I and II, that the equilibrium and kinetic behaviour of spontaneously-folding fixed two-state systems can be analysed by a treatment that is analogous to that given by Marcus for electron transfer.^1-3^ In this framework termed the parabolic approximation, the Gibbs energy functions of the denatured state ensemble (DSE) and the native state ensemble (NSE) are represented by parabolas whose curvature is given by their temperature-invariant force constants, α and ω, respectively. The temperature-invariant mean length of the reaction coordinate (RC) is given by *m*_D-N_ and is identical to the separation between the vertices of the DSE and the NSE-parabolas along the abscissa. Similarly, the position of the transition state ensemble (TSE) relative to the DSE and the NSE are given by *m*_TS-D(*T*)_ and *m*_TS-N(*T*)_, respectively, and are identical to the separation between the *curve-crossing* and the vertices of the DSE and the NSE-parabolas, respectively. The Gibbs energy of unfolding at equilibrium, Δ*G*_D-N(*T*)_, is identical to the separation between the vertices of the DSE and the NSE-parabolas along the ordinate. Similarly, the Gibbs activation energy for folding (Δ*G*_TS-D(*T*)_) and unfolding (Δ*G*_TS-N(*T*)_) are identical to the separation between the *curve-crossing* and the vertices of the DSE and the NSE-parabolas along the ordinate, respectively.

The purpose of this article is to use the framework described in Papers I and II to analyse the non-Arrhenius behaviour of the 37-residue FBP28 WW domain, at an unprecedented range and resolution.^4^

## Equations

The expressions for the position of the TSE relative to the vertices of the DSE and the NSE Gibbs parabolas are given by

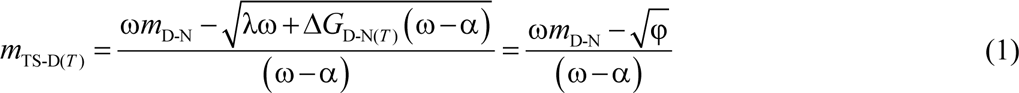

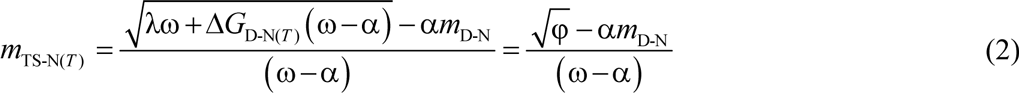

where the discriminant φ = λω + Δ*G*_D-N(*T*)_ (ω − α), and λ = α(*m*_D-N_)^2^ is the *Marcus reorganization energy* for two-state protein folding. The expressions for the activation energies for folding and unfolding are given by

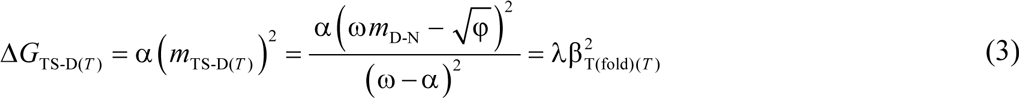

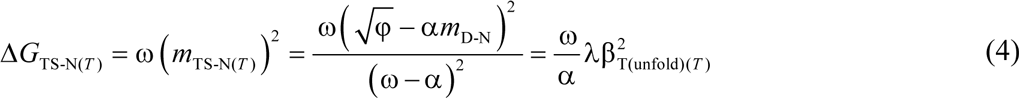

where the parameters β_T(fold)(*T*)_ (= *m*_TS-D(*T*)_/*m*_D-N_) and β_T(unfold)(*T*)_ (= *m*_TS-N(*T*)_/*m*_D-N_) are according to Tanford’s framework.^5^ The expressions for the rate constants for folding (*k*_*f*(*T*)_) and unfolding (*k*_*u*(*T*)_), and Δ*G*_D-N(*T*)_ are given by

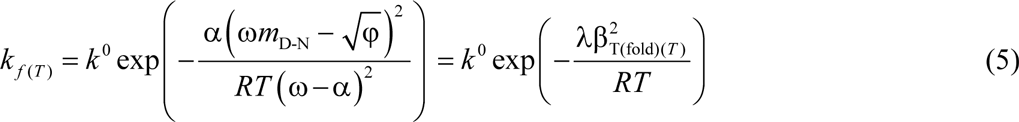

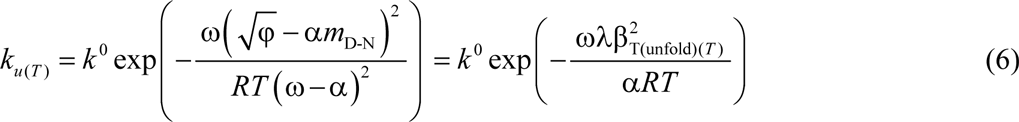

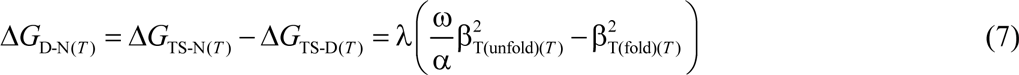

where, *k*^0^ is the temperature-invariant prefactor with units identical to those of the rate constants (s^−1^), *R* is the gas constant, *T* is the absolute temperature. If the temperature-dependence of Δ*G*_D-N(*T*)_ and the values of α, ω, and *m*_D-N_ are known for any two-state system at constant pressure and solvent conditions (see **Methods**), the temperature-dependence of the *curve-crossing* relative to the ground states may be readily ascertained. The temperature-dependence of *curve-crossing* is central to this analysis since all other parameters can be readily derived by manipulating the same using standard kinetic and thermodynamic relationships.

The activation entropies for folding (Δ*S*_TS-D(*T*)_) and unfolding (Δ*S*_TS-N(*T*)_) are given by the first derivatives of Δ*G*_TS-D(*T*)_ and Δ*G*_TS-N(*T*)_ functions with respect to temperature

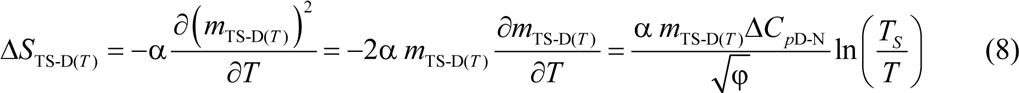

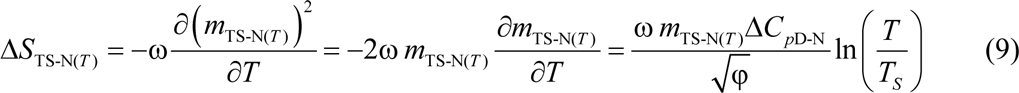

where *T_S_* is the temperature at which the entropy of unfolding at equilibrium is zero (Δ*S*_D-N(*T*)_ = 0) and Δ*C*_*p*D-N_ is the temperature-invariant difference in heat capacity between the DSE and the NSE.^6^ The activation enthalpies for folding (Δ*H*_TS-D(*T*)_) and unfolding (Δ*H*_TS-N(*T*)_) may be readily obtained by recasting the Gibbs equation: Δ*H*_(*T*)_ = Δ*G*_(*T*)_ + *T*Δ*S*_(*T*)_, or from the temperature-dependence of *k*_*f*(*T*)_ and *k*_*u*(*T*)_ to give

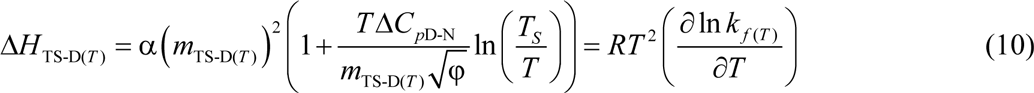

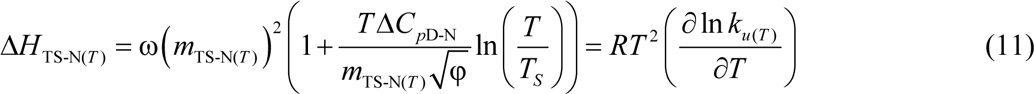

The difference in heat capacity between the DSE and the TSE (i.e., for the partial unfolding reaction [*TS*] ⇌ *D*) is given by

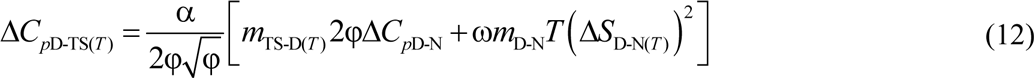

Similarly, the difference in heat capacity between the TSE and the NSE (for the partial unfolding reaction *N* ⇌ [*TS*]) is given by

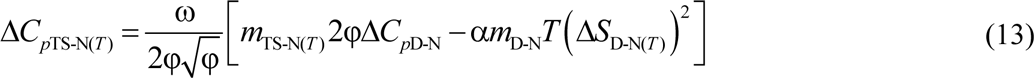

The reader may refer to Papers I and II for the derivations.

## Results and Discussion

As mentioned earlier and discussed in sufficient detail in Papers I and II, the analysis we are going to perform has an explicit requirement for a minimal experimental dataset which are: (*i*) an experimental chevron obtained at constant temperature, pressure and solvent conditions (except for the denaturant); (*ii*) an equilibrium thermal denaturation curve obtained under constant pressure, and in solvent conditions identical to those in which the chevron was acquired but without the denaturant, using either calorimetry or spectroscopy; and (*iii*) the calorimetrically determined Δ*C*_*p*D-N_ value (i.e., the slope of the linear regression of a plot of model-independent Δ*H*_D-N(*T_m_*)_ *vs T_m_*, where Δ*H*_D-N(*T_m_*)_ is the enthalpy of unfolding at the midpoint of thermal denaturation, *T_m_*; see Fig. 4 in Privalov, 1989).^7^ Fitting the chevron to a modified chevron-equation using non-linear regression yields the values of *m*_D-N_, the force constants α and ω, and the prefactor *k*^0^ (*k*^0^ is assumed to be temperature-invariant; see **Methods** in Paper I). Fitting a spectroscopic sigmoidal equilibrium thermal denaturation curve using standard two-state approximation (van’t Hoff analysis using temperature-invariant Δ*C*_*p*D-N_) yields van’t Hoff Δ*H*_D-N(*T_m_*)_ and *T_m_* and enables the temperature-dependence of Δ*H*_D-N(*T*)_, Δ*S*_D-N(*T*)_ and Δ*G*_D-N(*T*)_ functions to be ascertained across a wide temperature regime (Eqs. (A1)-(A3), **Figure 1** and **Figure 1−figure supplement 1**).^6^ Once the values of *m*_D-N_, the force constants, the prefactor, and the temperature dependence of Δ*G*_D-N(*T*)_ are known, the rest of the analysis is fairly straightforward. The values of all the reference temperatures that appear in this article are given in **Table 1**.

**Figure 1.**
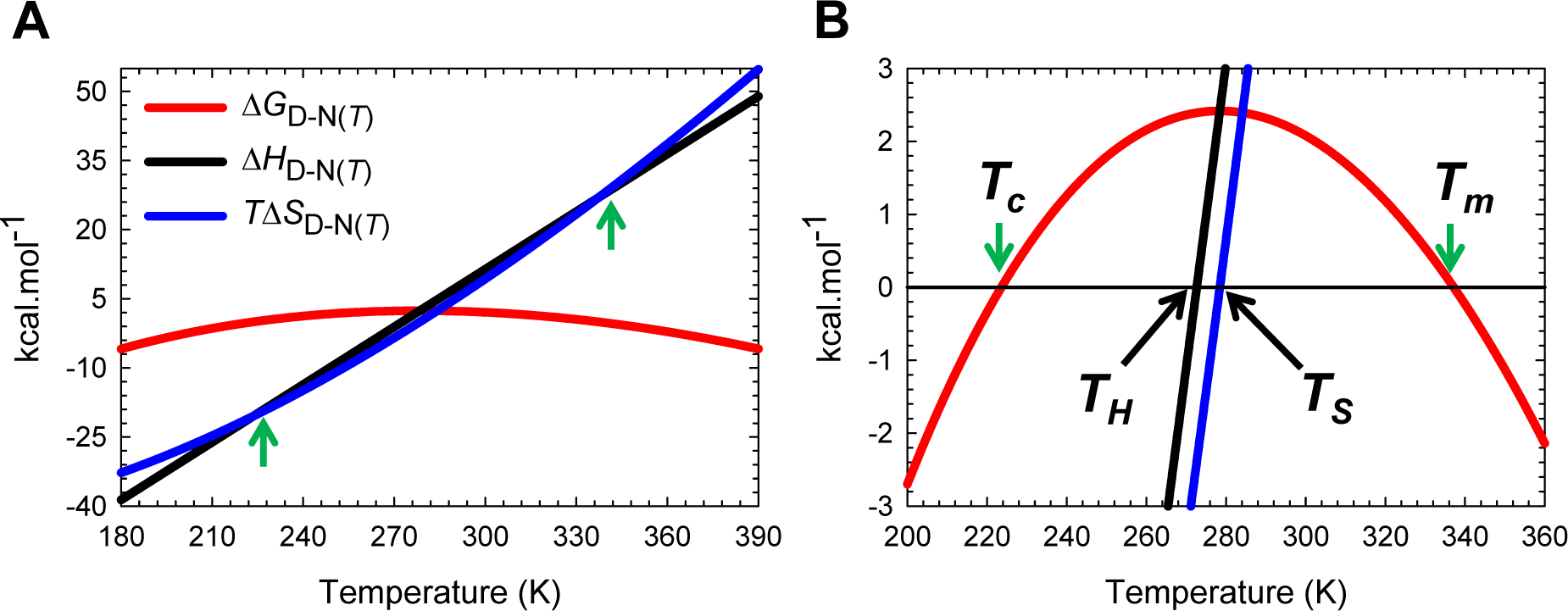
Stability curve for the unfolding reaction *N* ⇌ *D*. **(A)** Temperature-dependence of Δ*H*_D-N(*T*)_, Δ*S*_D-N(*T*)_ and Δ*G*_D-N(*T*)_ according to Eqs. (A1), (A2) and (A3), respectively. The green pointers identify the cold (*T_c_*) and heat (*T_m_*) denaturation temperatures. The slopes of the red and black curves are given by ∂Δ*G*_D-N(*T*)_/∂*T*=-Δ*S*_D-N(*T*)_ and ∂Δ*H*_D-N(*T*)_/∂*T*=Δ*C*_*p*D-N_, respectively. **(B)** An appropriately scaled version of the plot on the left. *T_H_* is the temperature at which Δ*H*_D-N(*T*)_ = 0, and *T_S_* is the temperature at which Δ*S*_D-N(*T*)_ = 0. The values of the reference temperatures are given in **Table 1**.

**Table 1.** Reference temperatures

| Temperature | Value | Remark |
| --- | --- | --- |
| $T_\alpha$ | 182 K | A two-state system is physically undefined for $T < T_\alpha$ |
| $T_{S(\alpha)}$ | 184.4 K | $m_{\text{TS-N}(T)} = 0, \Delta H_{\text{TS-N}(T)} = \Delta S_{\text{TS-N}(T)} = \Delta G_{\text{TS-N}(T)} = 0, k_{u(T)} = k^0$ |
| $T_{C_p\text{TS-N}(\alpha)}$ | 201 K | $\Delta C_{p\text{TS-N}(T)} = 0$ |
| $T_c$ | 223.6 K | Midpoint of cold denaturation, $\Delta G_{\text{D-N}(T)} = 0, k_{f(T)} = k_{u(T)}$ |
| $T_{H(\text{TS-N})}$ | 264.3 K | $\Delta H_{\text{TS-N}(T)} = 0, k_{u(T)}$ is a minimum |
| $T_H$ | 272.9 K | $\Delta H_{\text{TS-D}(T)} = \Delta H_{\text{TS-N}(T)}, \Delta H_{\text{D-N}(T)} = 0, \Delta H_{\text{TS-D}(T)} > 0, \Delta H_{\text{TS-N}(T)} > 0,$ |
| $T_S$ | 278.8 K | $\Delta S_{\text{TS-D}(T)} = \Delta S_{\text{TS-N}(T)} = \Delta S_{\text{D-N}(T)} = 0, \Delta G_{\text{D-N}(T)}$ is a maximum |
| $T_{H(\text{TS-D})}$ | 311.4 K | $\Delta H_{\text{TS-D}(T)} = 0, k_{f(T)}$ is a maximum |
| $T_m$ | 337.2 K | Midpoint of heat denaturation, $\Delta G_{\text{D-N}(T)} = 0, k_{f(T)} = k_{u(T)}$ |
| $T_{C_p\text{TS-N}(\omega)}$ | 361.7 K | $\Delta C_{p\text{TS-N}(T)} = 0$ |
| $T_{S(\omega)}$ | 384.5 K | $m_{\text{TS-N}(T)} = 0, \Delta H_{\text{TS-N}(T)} = \Delta S_{\text{TS-N}(T)} = \Delta G_{\text{TS-N}(T)} = 0, k_{u(T)} = k^0$ |
| $T_\omega$ | 388 K | A two-state system is physically undefined for $T > T_\omega$ |

### Temperature-dependence of *m*_TS-D(*T*)_ and *m*_TS-N(*T*)_

Substituting the expression for the temperature-dependence of *G*_D-N(*T*)_ (Eq. (A3), **Figure 1**) in Eqs. (1) and (2) enables the temperature-dependence of the *curve-crossing* relative to the DSE and the NSE to be ascertained (**Figure 2**; substituted expressions not shown). Because by postulate the force constants, Δ*C*_*p*D-N_, and *m*_D-N_ are temperature-invariant for any given primary sequence that folds in a two-state manner at constant pressure and solvent conditions, we get from inspection of Eqs. (1) and (2) that the discriminant φ, and 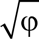 must be a maximum when Δ*G*_D-N(*T*)_ is a maximum. Because Δ*G*_D-N(*T*)_ is a maximum at *T_S_* (the temperature at which the entropy of unfolding at equilibrium, Δ*S*_D-N(*T*)_, is zero),^6^ a corollary is that φ and 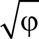 must be a maximum at *T_S_*; and any deviation in the temperature from *T_S_* will only lead to their decrease. Consequently, *m*_TS-D(*T*)_ and β_T(fold)(*T*)_ (= *m*_TS-D(*T*)_/*m*_D-N_) are always a minimum, and *m*_TS-N(*T*)_ and β_T(unfold)(*T*)_ (= *m*_TS-N(*T*)_/*m*_D-N_) are always a maximum at *T_S_*. This gives rise to two further corollaries: Any deviation in the temperature from *T_S_* can only lead to: (*i*) an increase in *m*_TS-D(*T*)_ and β_T(fold)(*T*)_; and (*ii*) a decrease in *m*_TS-N(*T*)_ and β_T(unfold)(*T*)_ (**Figure 2** and **Figure 2−figure supplement 1**). In other words, when *T* = *T_S_*, the TSE is the least native-like in terms of the SASA (solvent accessible surface area), and any deviation in temperature causes the TSE to become more native-like. A further consequence of *m*_TS-D(*T*)_ being a minimum at *T_S_* is that if for a two-state-folding primary sequence there exists a chevron with a well-defined linear folding arm at *T_S_*, then *m*_TS-D(*T*)_ > 0 and β_T(fold)(*T*)_ > 0 for all temperatures (**Figure 2A** and **Figure 2−figure supplement 1A**). Since the *curve-crossing* is physically undefined for φ < 0 owing to there being no real roots, the maximum theoretically possible value of *m*_TS-D(*T*)_ will occur when φ = 0 and is given by: 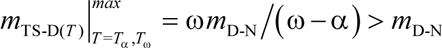 where *T*_α_ and *T*_ω_ are the temperature limits such that for *T* < *T*_α_ and *T* > *T*_ω_, a two-state system is not physically defined (see Paper II). Because *m*_D-N_ = *m*_TS-D(*T*)_ + *m*_TS-N(*T*)_ for a two-state system, and *m*_D-N_ is temperature-invariant by postulate, the theoretical minimum of *m*_TS-N(*T*)_ is given by: 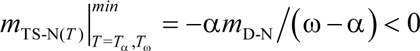. Now, since *m*_TS-N(*T*)_ is a maximum and positive at *T_S_* but its minimum is negative, a consequence is that *m*_TS-N(*T*)_ = β_T(unfold)(*T*)_ = 0 at two unique temperatures, one in the ultralow (*T*_*S*(α)_) and the other in the high (*T*_*S*(ω)_) temperature regime, and negative for *T*_α_ ≤ *T* < *T*_*S*(α)_ and *T*_*S*(ω)_ < *T* ≤ *T*_ω_ (**Figure 2B** and **Figure 2−figure supplement 1B**). Obviously, *m*_TS-D(*T*)_ = *m*_D-N_ and β_T(fold)(*T*)_ is unity at *T*_*S*(α)_ and *T*_*S*(ω)_. To summarize, unlike *m*_TS-D(*T*)_ and β_T(fold)(*T*)_ which are positive for all temperatures and a minimum at *T*_*S*_, *m*_TS-N(*T*)_ and β_T(unfold)(*T*)_ are a maximum at *T*_*S*_, zero at *T*_*S*(α)_ and *T*_*S*(ω)_, and negative for *T*_α_ ≤ *T* < *T*_*S*(α)_ and *T*_*S*(ω)_ < *T* ≤ *T*_ω_.

**Figure 2.**
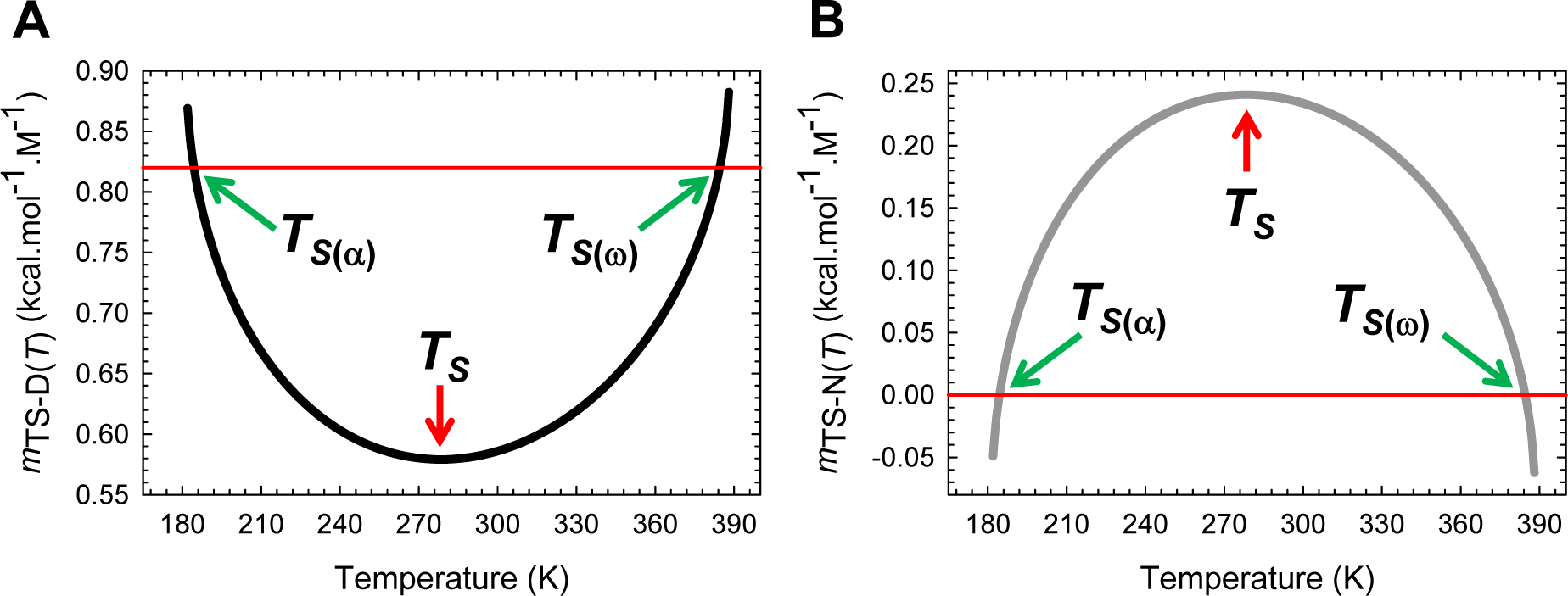
Temperature-dependence of *m*_TS-D(*T*)_ and *m*_TS-N(*T*)_. **(A)** *m*_TS-D(*T*)_ is a minimum at *T_S_*, is identical to *m*_D-N_ at *T*_*S*(*α*)_ and *T*_*S*(*ω*)_, and is greater than *m*_D-N_ for *T_α_* ≤ *T* < *T*_*S*(*α*)_ and *T*_*S*(*ω*)_ < *T* ≤ *T_ω_*. The slope of this curve is given by 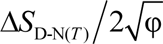 **(B)** *m*_TS-N(*T*)_ is a maximum at *T_S_*, zero at *T*_*S*(*α*)_ and *T*_*S*(*ω*)_, and negative for *T_α_* ≤ *T* < *T*_*S*(*α*)_ and *T*_*S*(*ω*)_ < *T* ≤ *T_ω_*. The slope of this curve is given by 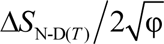. While the slopes of these curves are related to the activation entropies, the second derivatives of these functions with respect to temperature are related to the heat capacities of activation as shown in Paper-II. The values of the reference temperatures are given in **Table 1**.

The predicted Leffler-Hammond shift, which must be valid for any two-state system, is in agreement with the experimental data on the temperature-dependent behaviour of other two-state systems (Table 1 in Dimitriadis et al., 2004; Table 1 in Taskent et al., 2008; Fig. 5C in Otzen and Oliveberg, 2004),^8-12^ with the rate at which the *curve-crossing* shifts with stability (relative to the vertex of the DSE-parabola) being given by 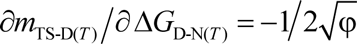. Importantly, just as the Leffler-Hammond movement is rationalized in physical organic chemistry using Marcus theory,^13-15^ we can similarly rationalize these effects in protein folding using parabolic approximation (**Figures 3**, **4**, and **Figure 4−figure supplement 1**). When *T* = *T_S_*, Δ*G*_TS-D(*T*)_ is a minimum, and Δ*G*_D-N(*T*)_ and Δ*G*_TS-N(*T*)_ are both a maximum; and any increase or decrease in the temperature relative to *T_S_* leads to a decrease in Δ*G*_TS-N(*T*)_, and an increase in Δ*G*_TS-D(*T*)_, consequently, leading to a decrease in Δ*G*_D-N(*T*)_ (**Figures 1**, **3B** and **5**). Naturally at *T_c_* and *T_m_*, Δ*G*_TS-D(*T*)_ = Δ*G*_TS-N(*T*)_, *k*_*f*(*T*)_ = *k*_*u*(*T*)_, and Δ*G*_D-N(*T*)_= 0 (**Figure 3C**). The reason why *m*_TS-D(*T*)_ = *m*_D-N_, and *m*_TS-N(*T*)_ = 0 at *T*_*S*(α)_ and *T*_*S*(ω)_ is apparent from **Figures 4A**, **4C** and **Figure 4−figure supplement 1A**: The right arm of the DSE-parabola intersects the vertex of the NSE-parabola leading to Δ*G*_TS-D(*T*)_ = α(*m*_TS-D(*T*)_)^2^ = α(*m*_D-N_)^2^ = λ, Δ*G*_TS-N(*T*)_ = ω(*m*_TS-N(*T*)_)^2^ = 0, and Δ*G*_D-N(*T*)_ = − λ. Importantly, in contrast to unfolding which can become barrierless at *T*_*S*(α)_ and *T*_*S*(ω)_, folding is barrier-limited at all temperatures, with the absolute minimum of Δ*G*_TS-D(*T*)_ occurring at *T_S_*; and any deviation in the temperature from *T_S_* will only lead to an increase in Δ*G*_TS-D(*T*)_ (**Figure 5A**). Thus, a corollary is that if folding is barrier-limited at *T_S_* (i.e., the chevron has a well-defined linear folding arm with a finite slope at *T_S_*), then a protein that folds *via* two-state mechanism can never spontaneously (i.e., unaided by ligands, co-solvents etc.) switch to a downhill mechanism (Type 0 scenario according to the Energy Landscape Theory; see Fig. 6 in Onuchic et al., 1997), no matter what the temperature, and irrespective of how fast or slow it folds. Although unfolding is barrierless at *T*_*S*(α)_ and *T*_*S*(ω)_, it is once again barrier-limited for *T*_α_ ≤ *T* < *T*_*S*(α)_ and *T*_*S*(ω)_ < *T* ≤ *T*_ω_, with the *curve-crossing* occurring to the right of the vertex of the NSE-parabola (**Figures 4A**, **4B**, **Figure 4−figure supplement 1B** and **5B**), such that *m*_TS-D(*T*)_ > *m*_D-N_, *m*_TS-N(*T*)_ < 0, β_T(fold)(*T*)_ > 1 and β_T(unfold)(*T*)_ < 0 (**Figure 2** and **Figure 2−figure supplement 1**).

**Figure 3.**
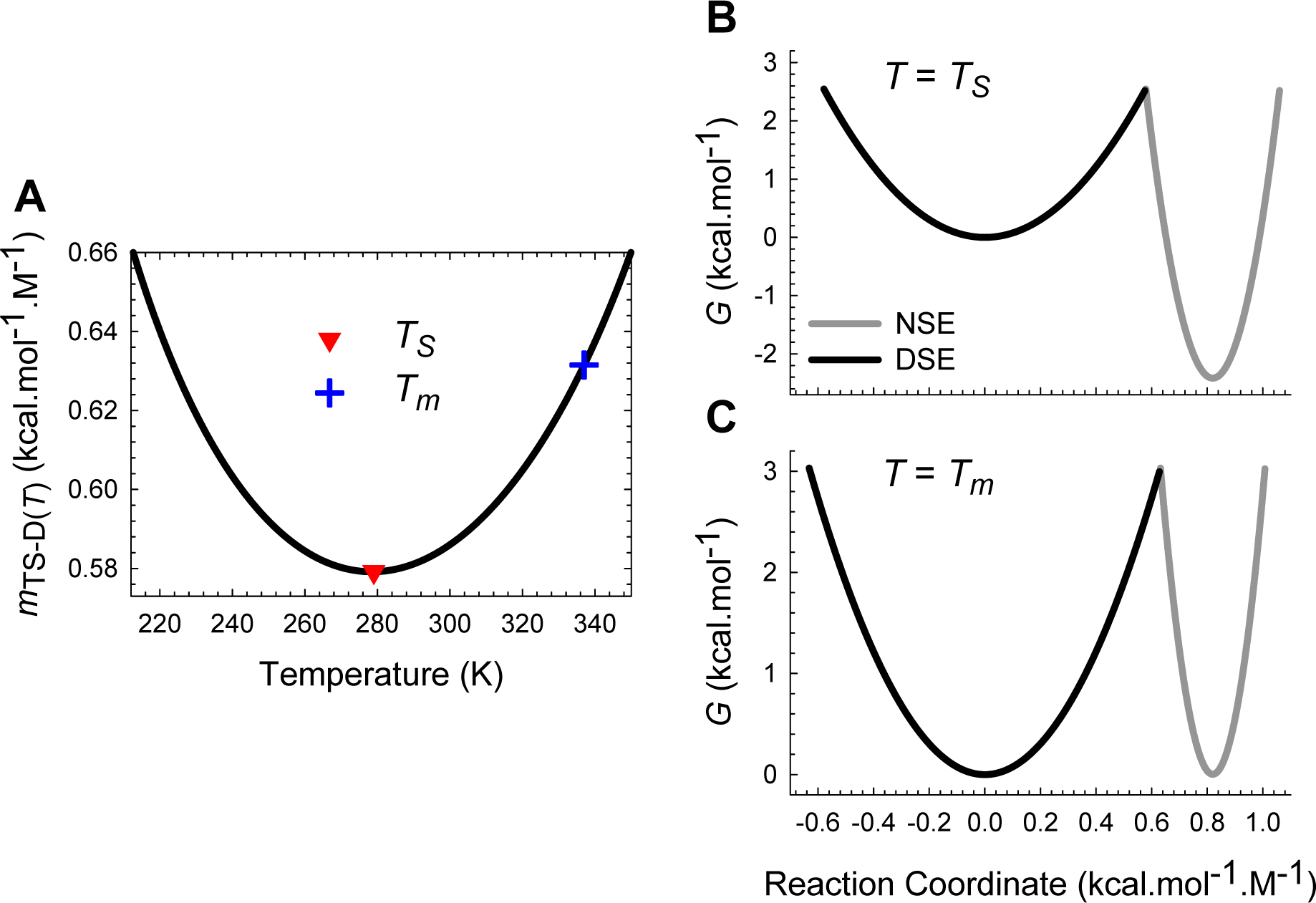
Marcus *curve-crossings* at *T_S_* and *T_m_*. **(A) Figure 2A** reproduced for comparison. **(B)** *Curve-crossing* at *T_S_* where Δ*G*_D-N(*T*)_ is a maximum and purely enthalpic (**Figure 1**). The relevant parameters are as follows: Δ*G*_TS-D(*T*)_ = 2.547 kcal.mol^-1^, Δ*G*_TS-N(*T*)_ = 4.964 kcal.mol^-1^, Δ*G*_D-N(*T*)_ = 2.417 kcal.mol^-1^, *k*_*f*(*T*)_ = 22009 s^-1^, *k*_*u*(*T*)_ = 280.8 s^-1^, *m*_TS-D(*T*)_ = 0.5792 kcal.mol^-1^.M^-1^ and *m*_TS-N(*T*)_ = 0.2408 kcal.mol^-1^.M^-1^. **(C)** *Curve-crossing* at *T_m_* and *T_c_* where Δ*G*_TS-D(*T*)_ = Δ*G*_TS-N(*T*)_ = 3.032 kcal.mol^-1^, Δ*G*_D-N(*T*)_ = 0, *k*_*f*(*T*)_ = *k*_*u*(*T*)_ = 23618 s^-1^, *m*_TS-D(*T*)_ = 0.6319 kcal.mol^-1^.M^-1^ and *m*_TS-N(*T*)_ = 0.1881 kcal.mol^-1^.M^-1^.The DSE and the NSE-parabolas are given by *G*_DSE(*r*,*T*)_=*αr*^2^ and *G*_NSE(*r*,*T*)_ = ω(*m*_D-N_ − *r*)^2^ − Δ*G*_D-N(*T*)_, respectively, α = 7.594 M^2^.mol.kcal^-1^, ω = 85.595 M^2^.mol.kcal^-1^, *m*_D-N_ (0.82 kcal.mol^-1^.M^-1^) is the separation between the vertices of the DSE and the NSE-parabolas along the abscissa, and *r* is any point on the abscissa. The abscissae are identical for plots B and C. The values of the reference temperatures are given in **Table 1**.

**Figure 5.**
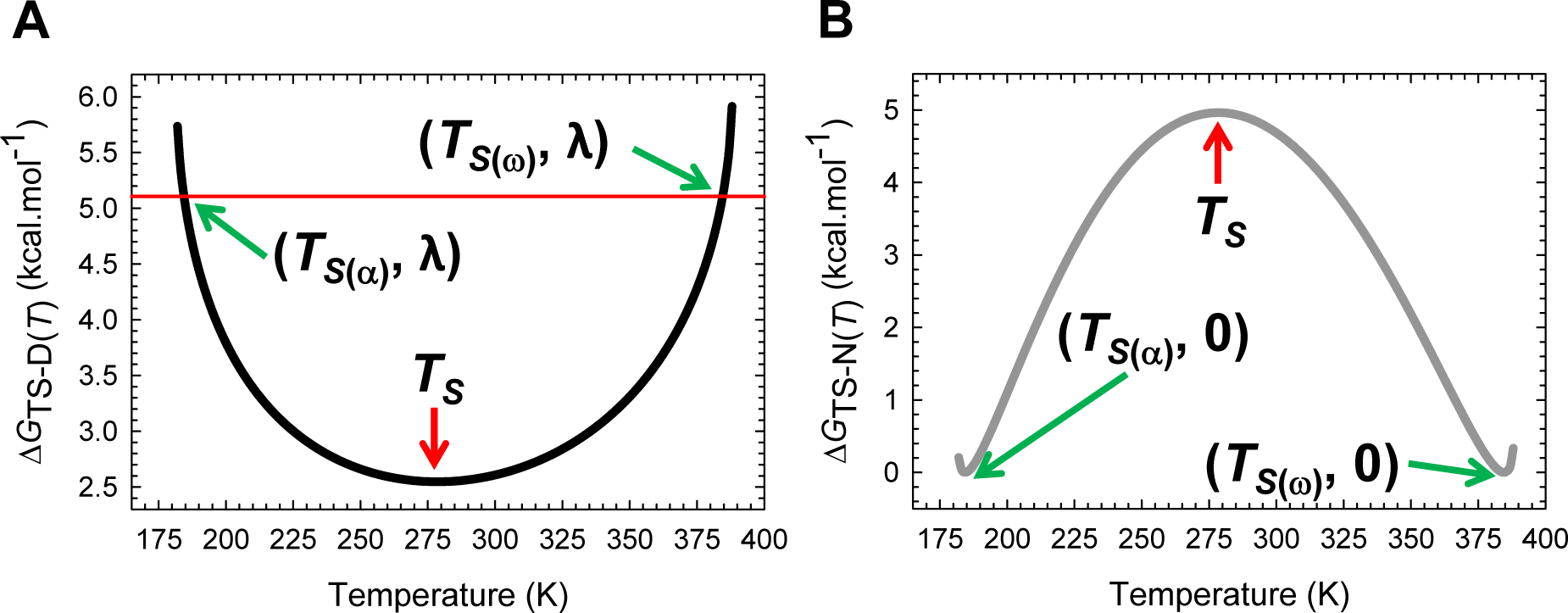
Temperature-dependence of the Gibbs activation energies for folding and unfolding. **(A)** Δ*G*_TS-D(*T*)_ is a minimum at *T_S_*, identical to λ = α(*m*_D-N_)^2^ = 5.106 kcal.mol^-1^ at *T*_*S*(*α*)_ and *T*_*S*(*ω*)_, and greater than λ for *T_α_* ≤ *T* < *T*_*S*(*α*)_ and *T*_*S*(*ω*)_ < *T* ≤ *T_ω_*. Note that ∂Δ*G*_TS-D(*T*)_/∂*T* = −Δ*S*_TS-D(*T*)_ = 0 at *T_S_*. **(B)** In contrast to Δ*G*_TS-D(*T*)_ which has only one extremum, Δ*G*_TS-N(*T*)_ is a maximum at *T_S_* and a minimum (zero) at *T*_*S*(*α*)_ and *T*_*S*(*ω*)_; consequently, ∂Δ*G*_TS-N(*T*)_/∂*T* = −∂*S*_TS-N(*T*)_ = 0 at *T*_*S*(*α*)_, *T_S_* and *T*_*S*(*ω*)_. Although unfolding is barrierless at *T*_*S*(*α*)_ and *T*_*S*(*ω*)_, it is once again barrier-limited for *T_α_* ≤ *T* < *T*_*S*(*α*)_ and *T*_*S*(*ω*)_ < *T* ≤ *T_ω_*; however, unlike the *conventional barrier-limited* unfolding which is characteristic for *T*_*S*(*α*)_ < *T* < *T*_*S*(*ω*)_, these two regimes fall under the *Marcus-inverted-region* and can be rationalized from **Figures 2**, **4**, and their figure supplements. The values of the reference temperatures are given in **Table 1**.

**Figure 6.**
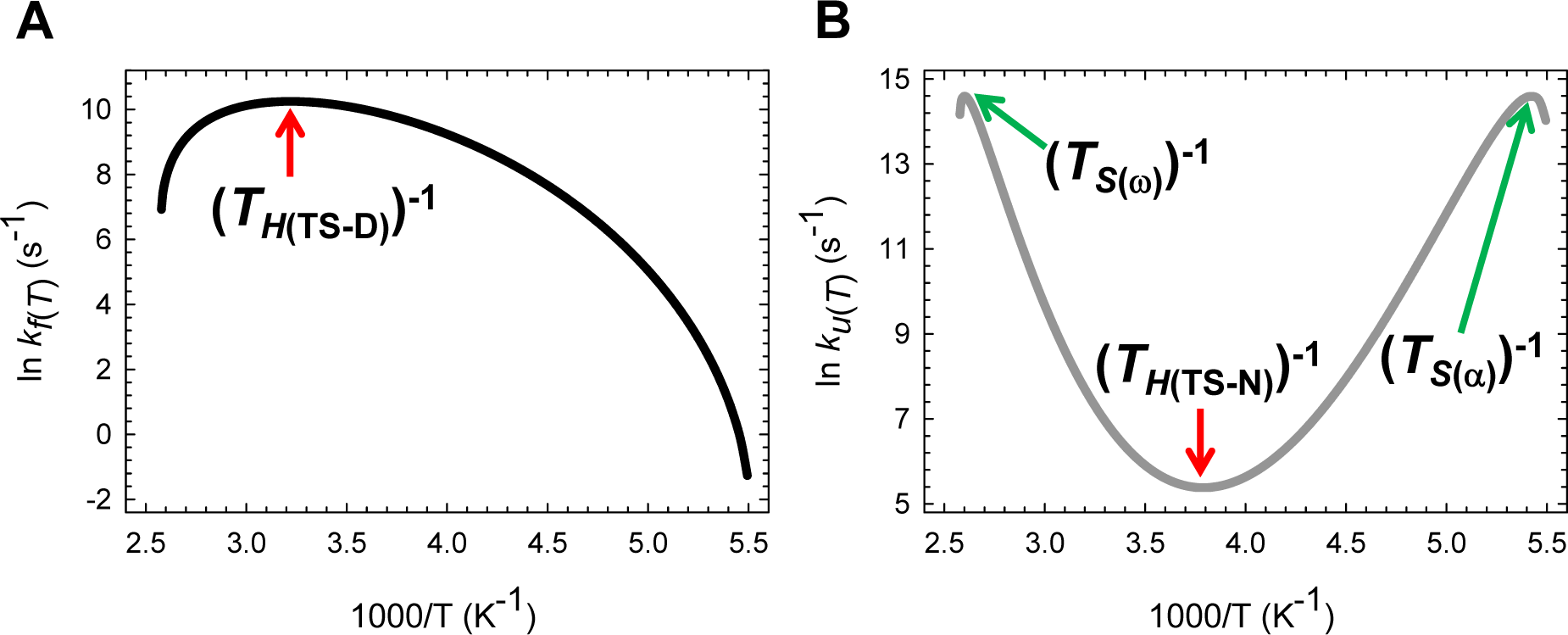
Arrhenius plots for the temperature-dependence of the rate constants. **(A)** *k*_*f*(*T*)_ is a maximum and Δ*H*TS-D(*T*)= 0 at *T*_*H*(TS-D)_. The slope of this curve is given by −Δ*H*_TS-D(*T*)_/*R*. **(B)** Unlike *k*_*f*(*T*)_ which has only one extremum, *k*_*u*(*T*)_ is a minimum at *T*_*H*(TS-N)_ and a maximum at *T*_*S*(*α*)_ and *T*_*S*(*ω*)_. Consequently, Δ*H*_TS-N(*T*)_ = 0 at *T*_*S*(*α*)_, *T*_*H*(TS-N)_ and *T*_*S*(*ω*)_. The slope of this curve is given by −Δ*H*_TS-N(*T*)_/*R*. When *T* = *T*_*S*(*α*)_ or *T*_*S*(*ω*)_, we have a unique scenario: *m*_TS-N(*T*)_ = Δ*G*_TS-N(*T*)_ = Δ*H*_TS-N(*T*)_ = 0 ⇒ Δ*S*TS-N(*T*) = 0, and *k*_*u*(*T*)_ = *k*^0^. Although unfolding is barrier-limited for *T_α_* ≤ *T* < *T*_*S*(*α*)_ and *T*_*S*(*ω*)_ < *T* ≤ *T_ω_*, leading to *k*_*u*(*T*)_ < *k*^0^, these ultra-low and high temperature regimes fall under the *Marcus-inverted-regime* as compared to the *conventional barrier-limited* unfolding which is characteristic for *T*_*S*(*α*)_ < *T* < *T*_*S*(*ω*)_ (the *curve-crossing* occurs in-between the vertices of the DSE and the NSE Gibbs basins) and can be rationalized comprehensively when considered in conjunction with **Figures 2**, **4**, and **5** (see also their figure supplements if any). The maxima of *k*_*f*(*T*)_ and *k*_*u*(*T*)_, as well as the *inverted-region* can be better appreciated on a linear scale as shown in the figure supplement. The values of the reference temperatures are given in **Table 1**.

**Figure 7.**
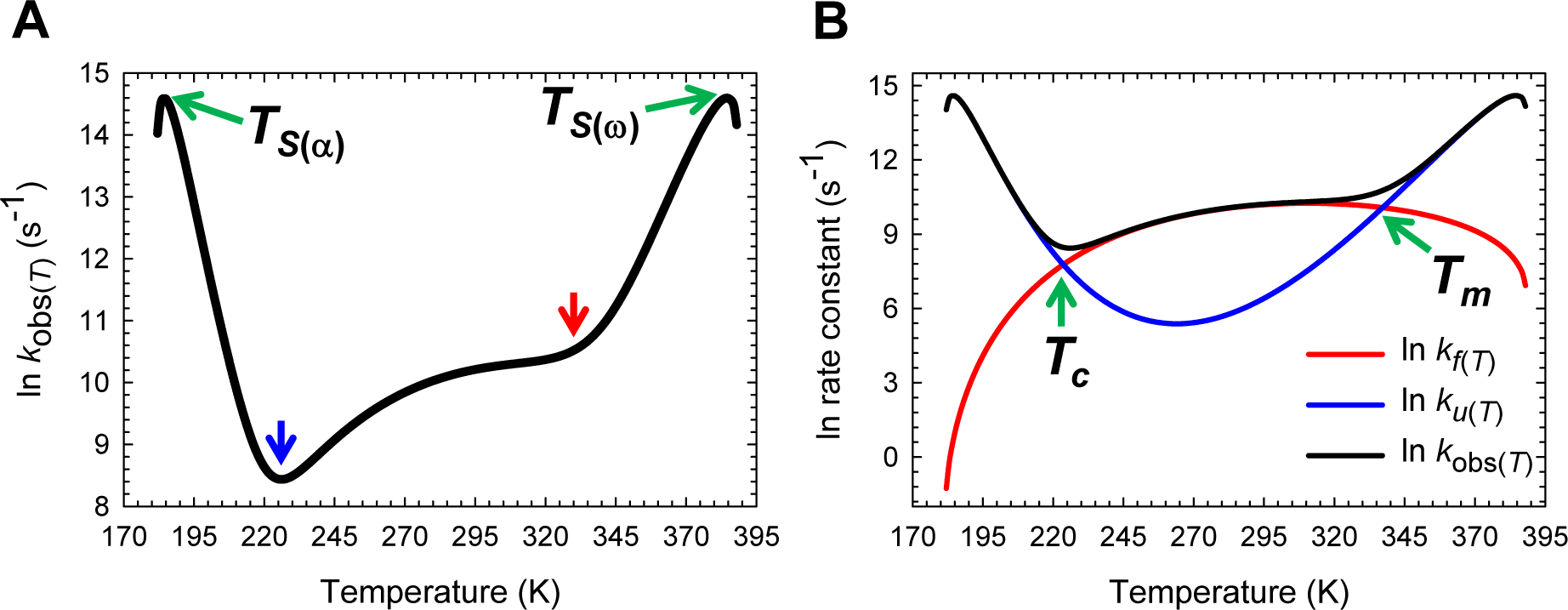
Temperature-dependence of the observed rate constant. **(A)** *k*_obs(*T*)_ is a maximum at *T*_*S*(*α*)_ and *T*_*S*(*ω*)_, and a minimum around *T_c_* (blue pointer). The red pointer indicates *T_m_*. The steep increase in *k*_obs(*T*)_ at very low and high temperatures is due to Δ*G*_TS-N(*T*)_ approaching zero as described in previous figures. **(B)** An overlay of *k*_*f*(*T*)_, *k*_*u*(*T*)_ and *k*_obs(*T*)_ to illuminate how the features of *k*_obs(*T*)_ arise from the sum of *k*_*f*(*T*)_ and *k*_*u*(*T*)_. The slopes of the red and blue curves are given by ∆*H*_TS-D(*T*)_/*RT*^2^ and ∆*H*_TS-N(*T*)_/*RT*^2^, respectively.

To summarize, for any two-state folder, unfolding is *conventional barrier-limited* for *T*_*S*(α)_ < *T* < *T*_*S*(ω)_ and the position of the TSE or the *curve-crossing* occurs in between the vertices of the DSE and the NSE parabolas. As the temperature deviates from *T_S_*, the SASA of the TSE becomes progressively native-like, with a concomitant increase and a decrease in Δ*G*_TS-D(*T*)_ and Δ*G*_TS-N(*T*)_, respectively. When *T* = *T*_*S*(α)_ and *T*_*S*(ω)_, the *curve-crossing* occurs precisely at the vertex of the NSE-parabola, the SASA of the TSE is identical to that of the NSE, and unfolding is barrierless; and for *T*_α_ ≤ *T* < *T*_*S*α_ and *T*_*S*(ω)_ < *T* ≤ *T*_ω_, unfolding is once again barrier-limited but falls under the *Marcus-inverted-regime* with the *curve-crossing* occurring on the right-arm of the NSE-parabola, leading to the SASA of the NSE being greater than that of the TSE (i.e., the TSE is more compact than the NSE). Importantly, for *T* < *T*_α_ and *T* > *T*_ω_, the TSE cannot be physically defined owing to 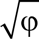 being mathematically undefined for φ < 0. A consequence is that *k_f(T)_* and *k_u(T)_* become physically undefined, leading to Δ*G*_D-N(*T*)_ = *RT* ln (*k_f(T)_*/*k_u(T)_*) being physically undefined, such that all of the conformers will be confined to a single Gibbs energy well, which is the DSE, and the protein will cease to function.^16^ Thus, from the view point of the physics of phase transitions, *T*_α_ ≤ *T* ≤ *T*_ω_ denotes the *coexistence temperature-range* where the DSE and the NSE, which are in a dynamic equilibrium, will coexist as two distinct phases; and for *T* < *T*_α_ and *T* > *T*_ω_ there will be a single phase, which is the DSE, with *T*_α_ and *T*_ω_ being the limiting temperatures for coexistence, or phase boundary temperatures from the view point of the DSE.^17-23^ This is roughly analogous to the operating temperature range of a logic circuit such as a microprocessor; and just as this range is a function of its constituent material, the physically definable temperature range of a two-state system is a function of the primary sequence when pressure and solvent are constant, and importantly, can be modulated by a variety of *cis-*acting and *trans*-acting factors (see Paper-I). The limit of equilibrium stability below which a two-state system becomes physically undefined is given by: Δ*G*_D-N(*T*)_|_*T* = *T*_α_, *T*_ω__ = −λω/(ω − α). Consequently, the physically meaningful range of equilibrium stability for a two-state system is given by: Δ*G*_D-N(*T_S_*)_ + [λω/(ω − α)], where Δ*G*_D-N(*T_S_*)_ is the stability at *T_S_* and is apparent from inspection of **Figure 5−figure supplement 1**. This is akin to the stability range over which Marcus theory is physically realistic (see Kresge, 1973, page 494).^24^

Because by postulate *m*_D-N_, *m*_TS-D(*T*)_ and *m*_TS-N(*T*)_ are true proxies for ΔSASA_D-N_, ΔSASA_D-TS(*T*)_ and ΔSASA_TS-N(*T*)_, respectively (see Paper I), we have three fundamentally important corollaries that must hold for all two-state systems at constant pressure and solvent conditions: (*i*) the Gibbs barrier to folding is the least when the denatured conformers bury the least amount of SASA to reach the TSE (**Figure 5−figure supplement 2A**); (*ii*) the Gibbs barrier to unfolding is the greatest when the native conformers expose the greatest amount of SASA to reach the TSE (**Figure 5−figure supplement 2B**); and (*iii*) equilibrium stability is the greatest when the conformers in the DSE are displaced the least from the mean of their ensemble along the SASA-RC to reach the TSE (the *principle of least displacement*; **Figure 5−figure supplement 1**).

### Temperature-dependence of the folding, unfolding, and the observed rate constants

Inspection of **Figures 6A** and **Figure 6−figure supplement 1A** demonstrates that Eq. (5) makes a remarkable prediction that *k_f(T)_* has a non-linear dependence on temperature. Starting from the lowest temperature (*T*_α_) at which a two-state system is physically defined, *k_f(T)_* initially increases with an increase in the temperature and reaches a maximal value at *T* = *T*_*H*(TS-D)_ where ∂ ln *k_f(T)_*/∂*T* = Δ*H*_TS-D(*T*)_/*RT*^2^ = 0; and any further increase in temperature beyond this point will cause a decrease in *k_f(T)_* until the temperature *T*_ω_ is reached, such that for *T* > *T*_ω_, *k_f(T)_* is undefined. Inspection of **Figures 6B** and **Figure 6−figure supplement 1B** demonstrates that the temperature-dependence of *k_u(T)_* is far more complex: Starting from *T*_α_, *k_u(T)_* increases with temperature for the regime *T*_α_ ≤ *T* < *T*_*S*(α)_ (the low-temperature *Marcus-inverted-regime*), reaches a maximum when *T* = *T*_*S*(α)_ (*k_u(T)_* = *k*^0^; the first extremum of *k_u(T)_*), and decreases with further rise in temperature for the regime *T*_*S*(α)_ < *T* < *T*_*H*(TS-N)_ such that when *T* = *T*_*H*(TS-N)_, *k_u(T)_* is a minimum (the second extremum of *k_u(T)_*). And for *T*_*H*(TS-N)_ < *T* < *T*_*S*(ω)_, an increase in temperature will lead to an increase in *k_u(T)_*, eventually leading to its saturation at *T* = *T*_*S*(ω)_ (*k_u(T)_* = *k*^0^; the third extremum of *k_u(T)_*), and decreases with further rise in temperature for *T*_*S*(ω)_ < *T* ≤ *T*_ω_ (the high-temperature *Marcus-inverted-regime*). Thus, in contrast to *k_f(T)_* which has only one extremum, *k_u(T)_* is characterised by three extrema where ∂ ln *k_u(T)_*/∂*T* = Δ*H*_TS-N(*T*)_/*R*^2^ = 0, and may be rationalized from the temperature-dependence of *m*_TS-D(*T*)_ and *m*_TS-N(*T*)_, the Gibbs barrier heights for folding and unfolding, and the intersection of the DSE and the NSE Gibbs parabolas (**Figures 2**-**5** and their figure supplements). We will show in subsequent publications that the inverted behaviour at very low and high temperatures is not common to all fixed two-state systems and depends on the mean and variance of the Gaussian distribution of the SASA of the conformers in the DSE and the NSE.

Since the ultimate test of any hypothesis is experiment, the most important question now is how well do the calculated rate constants compare with experiment? Although Nguyen et al. have investigated the non-Arrhenius behaviour of the FBP28 WW, they find that the behaviour of its wild type is erratic, with its folding being three-state for *T* < *T_m_* and two-state for *T* > *T_m_* (Fig. 3A in Nguyen et al., 2003). Consequently, non-Arrhenius data for the wild type FBP28 WW are lacking. Incidentally, this atypical behaviour is probably artefactual since the protein aggregates and forms fibrils under the experimental conditions in which the measurements were made (see Figs. 2, 3 and 6 in Ferguson et al., 2003).^25,26^ Nevertheless, data for ΔNΔC Y11R W30F, a variant of FBP28 WW are available between ∽ 298 and ∽357 K (Fig. 4A in Nguyen et al., 2003). Now since the relaxation time constants for the fast phase of wild type FBP28 WW (∽ 30 μs at 39.5 °C and < 15 μs at 65 °C, page 3950, Fig. 3A, Nguyen et al., 2003) are very similar to those of ΔNΔC Y11R W30F (∽ 28 μs at 40 °C and 11 μs at 65 °C, page 3952), a reasonable approximation is that the temperature-dependence of *k_f(T)_* and *k_u(T)_* of the wild type and the mutant must be similar. Consequently, the temperature-dependence of the rate constants for the wild type FBP28 WW calculated using parabolic approximation must be very similar to the data for ΔNΔC Y11R W30F reported by Nguyen et al. The remarkable agreement between the said datasets is readily apparent from a comparison of Fig. 4A of Nguyen et al., and **Figure 6−figure supplement 2**, and serves an important test of the hypothesis.

Since the temperature-dependence of *k_f(T)_* and *k_u(T)_* across a wide temperature range is known, the variation in the observed rate constant (*k*_obs(*T*)_) with temperature may be readily ascertained using (see **Appendix**)

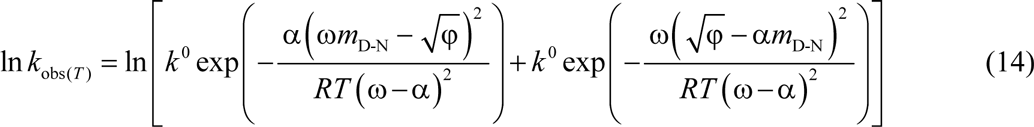

Inspection of **Figure 7** demonstrates that ln(*k*_obs(*T*)_) *vs* temperature is a smooth ‘W-shaped’ curve, with *k*_obs(*T*)_ being dominated by *k_f(T)_* around *T*_*H*(TS-N)_, and by *k_u(T)_* for *T* < *T_c_* and *T* > *T_m_*, which is precisely why the kinks in ln(*k*_obs(*T*)_) occur around these temperatures. It is easy to see that at *T_c_* or *T_m_*, *k_f(T)_* = *k_u(T)_* ⇒ *k*_obs(*T*)_ = 2*k_f(T)_* = 2*k_u(T)_,* Δ*G*_D-N(*T*)_ = *RT* ln (*k_f(T)_*/*k_u(T)_*)) = 0 or Δ*G*_TS-D(*T*)_ = Δ*G*_TS-N(*T*)_ (**Figures 3C** and **Figure 7−figure supplement 1**). In other words, for a two-state system, *T_c_* and *T_m_* determined at equilibrium must be identical to the temperatures at which *k_f(T)_* and *k_u(T)_* intersect. This is a consequence of the *principle of microscopic reversibility*, i.e., the equilibrium and kinetic stabilities must be identical for a two-state system at all temperatures.^27^ It is precisely for this reason that the value of the prefactor in the Arrhenius expressions for the rate constants must be identical for both the folding and the unfolding reactions at all temperatures (Eqs. (5) and (6)). The steep increase in *k*_obs(*T*)_ for *T* < *T_c_* and *T* > *T_m_* is due to the Δ*G*_TS-N(*T*)_ approaching zero as described earlier. The argument that the shapes of the curves must be conserved across two-state systems applies not only to the temperature-dependence of *m*_TS-D(*T*)_, *m*_TS-N(*T*)_, Δ*G*_TS-D(*T*)_ and Δ*G*_TS-N(*T*)_ described so far, but to the rest of the state functions that will be described in this article (see Paper-I).

An important conclusion that we may draw from these data is the following: Because we have assumed a temperature-invariant prefactor and yet find that the kinetics are non-Arrhenius, it essentially implies that one does not need to invoke a *super-Arrhenius temperature-dependence of the configurational diffusion constant* to explain the non-Arrhenius behaviour of proteins.^28-32^ Instead, as long as the enthalpies and the entropies of unfolding/folding at equilibrium display a large variation with temperature, and equilibrium stability is a non-linear function of temperature, both *k_f(T)_* and *k_u(T)_* will have a non-linear dependence on temperature. This leads to two corollaries: (*i*) since the large variation in equilibrium enthalpies and entropies of unfolding, including the pronounced curvature in Δ*G*_D-N(*T*)_ of proteins with temperature is due to the large and positive Δ*C*_*p*D-N_, “*non-Arrhenius kinetics can be particularly acute for reactions that are accompanied by large changes in the heat capacity*”; and (*ii*) because the change in heat capacity upon unfolding is, to a first approximation, proportional to the change in SASA that accompanies it, and since the change in SASA upon unfolding/folding increases with chain-length,^33,34^ “*non-Arrhenius kinetics, in general, can be particularly pronounced for large proteins, as compared to very small proteins and peptides*.”

### Temperature-dependence of activation enthalpies

Inspection of **Figure 8** demonstrates that for the partial folding reaction *D* ⇌ [*TS*]: (*i*) Δ*H*_TS-D(*T*)_ > 0 for *T*_α_ ≤ *T* < *T*_*H*(TS-D)_; (*ii*) Δ*H*_TS-D(*T*)_ < 0 for *T*_*H*(TS-D)_ < *T* ≤ *T*_ω_ and (*iii*) Δ*H*_TS-D(*T*)_ = 0 for *T* = *T*_*H*(TS-D)_. Thus, the activation of the denatured conformers to the TSE is enthalpically: (*i*) unfavourable for *T*_α_ ≤ *T* < *T*_*H*(TS-D)_; (*ii*) favourable for *T*_*H*(TS-D)_ < *T* ≤ *T*_ω_; and (*iii*) neutral when *T* = *T*_*H*(TS-D)_. Consequently, at *T*_*H*(TS-D)_, Δ*G*_TS-D(*T*)_ is purely due to the difference in entropy between the DSE and the TSE (Δ*G*_TS-D(*T*)_ = −*T*Δ*S*_TS-D(*T*)_) with *k_f(T)_* being given by

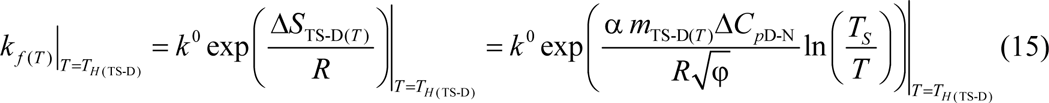

**Figure 8.**
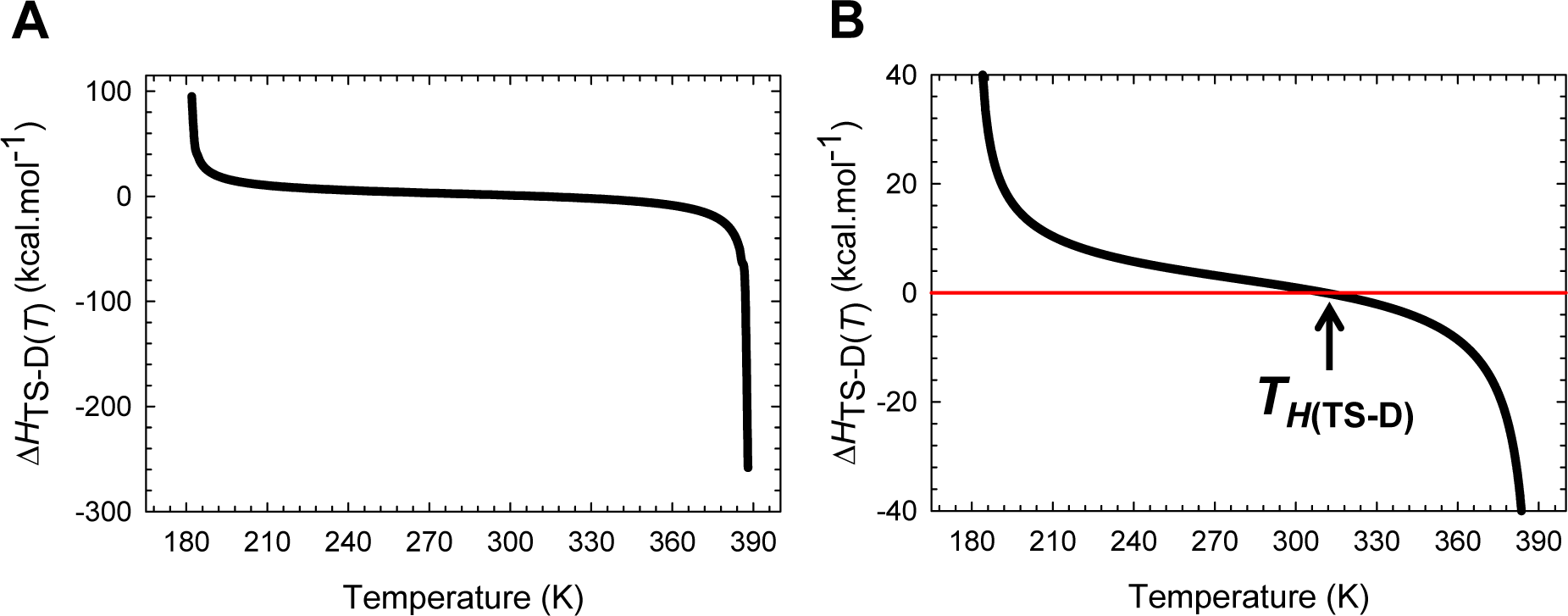
Temperature-dependence of the activation enthalpy for folding. **(A)** The variation in Δ*H*_TS-D(*T*)_ function with temperature. The slope of this curve varies with temperature, equals Δ*C*_*p*TS-D(*T*)_, and is algebraically negative. **(B)** An appropriately scaled version of the plot on the left to illuminate the three important scenarios: (*i*) Δ*H*_TS-D(*T*)_ > 0 for *T_α_* ≤ *T* < *T*_*H*(TS-D)_; (*ii*) Δ*H*_TS-D(*T*)_ < 0 for *T*_*H*(TS-D)_ < *T* ≤ *T_ω_*; and (*iii*) Δ*H*_TS-D(*T*)_ = 0 when *T* = *T*_*H*(TS-D)_. Note that *k*_*f*(*T*)_ is a maximum at *T*_*H*(TS-D)_.

Because *k_f(T)_* is a maximum at *T*_*H*(TS-D)_ (∂ ln *k_f(T)_*/∂*T* = 0), a corollary is that *“for a two-state folder at constant pressure and solvent conditions, if the prefactor is temperature-invariant, then k_f(T)_ will be a maximum when the Gibbs barrier to folding is purely entropic.”* This statement is valid only if the prefactor is temperature-invariant. Now since Δ*G*_TS-D(*T*)_ > 0 for all temperatures (**Figure 5A** and **Table 1**), it is imperative that Δ*S*_TS-D(*T*)_ < 0 at *T*_*H*(TS-D)_ (see activation entropy for folding).

Unlike the Δ*H*_TS-D(*T*)_ function which changes its algebraic sign only once across the entire temperature range over which a two-state system is physically defined, the behaviour of Δ*H*_TS-N(*T*)_ function is far more complex (**Figure 9**): (*i*) Δ*H*_TS-N(*T*)_ > 0 for *T*_α_ ≤ *T* < *T*_*S*(α)_ and *T*_*H*(TS-N)_ < *T* < *T*_*S*(ω)_; (*ii*) Δ*H*_TS-N(*T*)_ < 0 for *T*_*S*(α)_ < *T* < *T*_*H*(TS-N)_ and *T*_*S*(ω)_ < *T* ≤ *T*_ω_; and (*iii*) Δ*H*_TS-N(*T*)_ = 0 at *T*_*S*(α)_, *T*_*H*(TS-N)_, and *T*_*S*(ω)_. Consequently, we may state that the activation of native conformers to the TSE is enthalpically: (*i*) unfavourable for *T*_α_ ≤ *T* < *T*_*S*(α)_ and *T*_*H*(TS-N)_ < *T* < *T*_*S*(ω)_; (*ii*) favourable for *T*_*S*(α)_ < *T* < *T*_*H*(TS-N)_ and *T*_*S*(ω)_ < *T* ≤ *T*_ω_; and (*iii*) neutral at *T*_*S*(α)_, *T*_*H*(TS-N)_, and *T*_*S*(ω)_. If we reverse the reaction-direction, the algebraic signs invert leading to a change in the interpretation. Thus, for the partial folding reaction [*TS*] ⇌ *N*, the flux of the conformers from the TSE to the NSE is enthalpically: (*i*) favourable for *T*_α_ ≤ *T* < *T*_*S*(α)_ and *T*_*H*(TS-N)_ < *T* < *T*_*S*(ω)_ (Δ*H*_N-TS(*T*)_ < 0); (*ii*) unfavourable for *T*_*S*(α)_ < *T* < *T*_*H*(TS-N)_ and *T*_*S*(ω)_ < *T* ≤ *T*_ω_ (Δ*H*_N-TS(*T*)_ > 0); and (*iii*) neither favourable nor unfavourable at *T*_*S*(α)_, *T*_*H*(TS-N)_, and *T*_*S*(ω)_ (**Figure 9−figure supplement 1A**). Note that the term “flux” implies “diffusion of the conformers from one reaction state to the other on the Gibbs energy surface,” and as such is an “operational definition.”

**Figure 9.**
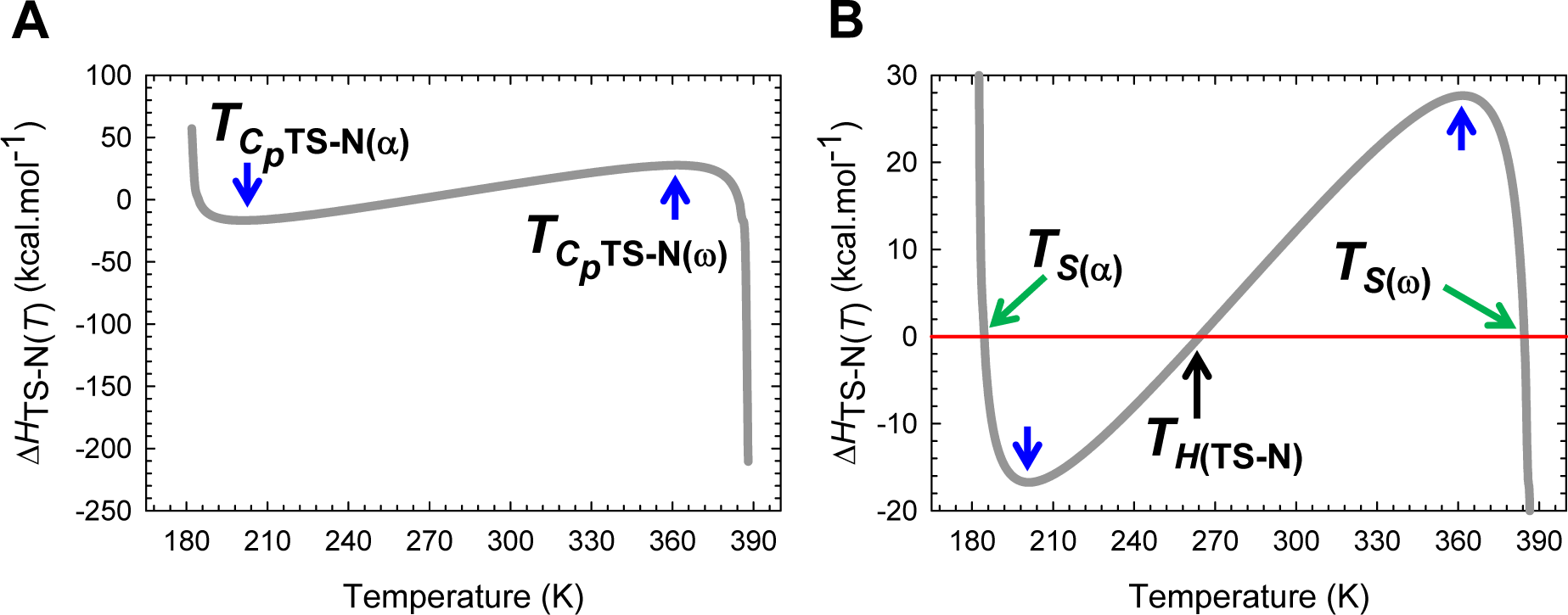
Temperature-dependence of the activation enthalpy for unfolding. **(A)** The variation in Δ*H*TS-N(*T*) function with temperature. The slope of this curve equals Δ*C*_*p*TS-N(*T*)_ and is zero at *T*_*C_p_*TS-N(α)_ and *T*_*C_p_*TS-N(ω)_. **(B)** An appropriately scaled version of the figure on the left to illuminate the various temperature-regimes and their implications: (*i*) Δ*H*_TS-N(*T*)_ > 0 for *T*_α_ ≤ *T* < *T*_*S*(*α*)_ and *T*_*H*(TS-N)_ < *T* < *T*_*S*(*ω*)_; (*ii*) Δ*H*_TS-N(*T*)_ < 0 for *T*_*S*(*α*)_ < *T* < *T*_*H*(TS-N)_ and *T*_*S*(*ω*)_ < *T* ≤ *T_ω_*; and (*iii*) Δ*H*_TS-N(*T*)_ = 0 at *T*_*S*(*α*)_, *T*_*H*(TS-N)_, and *T*_*S*(*ω*)_. Note that at *T*_*S*(*α*)_ and *T*_*S*(*ω*)_, we have the unique scenario: *m*_TS-N(*T*)_ = Δ*G*_TS-N(*T*)_ = Δ*S*_TS-N(*T*)_ = Δ*H*_TS-N(*T*)_ = 0, and *k*_*u*(*T*)_ = *k*^0^. The values of the reference temperatures are given in **Table 1**.

Importantly, although ∂ ln *k_u(T)_*/∂*T* = 0 ⇒ Δ*H*_TS-N(*T*)_ = 0 at *T*_*S*(α)_, *T*_*H*(TS-N)_, and *T*_*S*(ω)_, the behaviour of the system at *T*_*S*(α)_ and *T*_*S*(ω)_ is distinctly different from that at *T*_*H*(TS-N)_: While *m*_TS-N(*T*)_ = Δ*G*_TS-N(*T*)_ = Δ*H*_TS-N(*T*)_ = Δ*S*_TS-N(*T*)_ = 0, *m*_TS-D(*T*)_ = *m*_D-N_, Δ*G*_TS-D(*T*)_ = Δ*G*_N-D(*T*)_ = λ, and *k_u(T)_* = *k*^0^ at *T*_*S*(α)_ and *T*_*S*(ω)_ (note that if both Δ*G*_TS-N(*T*)_ and Δ*H*_TS-N(*T*)_ are zero, then Δ*S*_TS-N(*T*)_ must also be zero, see activation entropies), *k_u(T)_* is a minimum (*k_u(T)_* ≪ *k*^0^) with the Gibbs barrier to unfolding being purely entropic (Δ*G*_TS-N(*T*)_ = −*T*Δ*S*_TS-N(*T*)_) at *T*_*H*(TS-N)_. Consequently, we may write

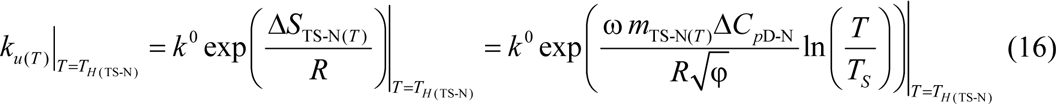

Thus, a corollary is that *“for two-state system at constant pressure and solvent conditions, if the prefactor is temperature-invariant, then k_u(T)_ will be a minimum when the Gibbs barrier to unfolding is purely entropic.”* Since Δ*G*_TS-N(*T*)_ > 0 at *T*_*H*(TS-N)_ (**Figure 5B** and **Table 1**), it is imperative that Δ*S*_TS-N(*T*)_ be negative at *T*_*H*(TS-N)_ (see activation entropy for unfolding).

The criteria for two-state folding from the viewpoint of enthalpy are the following: (*i*) the condition that Δ*H*_D-N(*T*)_ = Δ*H*_TS-N(*T*)_ − Δ*H*_TS-D(*T*)_ must be satisfied at all temperatures; (*ii*) the intersection of Δ*H*_TS-D(*T*)_ and Δ*H*_TS-N(*T*)_ functions calculated directly from the temperature-dependence of the experimentally determined *k_f(T)_* and *k_u(T)_*, respectively, must be identical to the independently estimated *T_H_* from equilibrium thermal denaturation experiments; and (*iii*) the condition that *T*_*H*(TS-N)_ < *T_H_* < *T_S_* < *T*_*H*(TS-D)_ must be satisfied. A corollary of the last statement is that both Δ*H*_TS-D(*T*)_ and Δ*H*_TS-N(*T*)_ functions must be positive at the point of intersection. These aspects are readily apparent from **Figure 9−figure supplement 1B** and **Figure 9−figure supplement 2**.

### Temperature-dependence of activation entropies

Inspection of **Figure 10** shows that for the partial folding reaction *D* ⇌ [*TS*], Δ*S*_TS-D(*T*)_ which is positive at low temperature, decreases in magnitude with an increase in temperature and becomes zero at *T_S_*, where the SASA of the TSE is the least native-like, Δ*G*_TS-D(*T*)_ is a minimum (∂Δ*G*_TS-D(*T*)_/∂*T* = −Δ*S*_TS-D(*T*)_ = 0) and Δ*G*_D-N(*T*)_ is a maximum (∂Δ*G*_D-N(*T*)_/∂*T* = −Δ*S*_D-N(*T*)_ = 0; **Figures 1, 2, 5A, Figure 10−figure supplements 1** and **2**); and any further increase in temperature beyond this point causes Δ*S*_TS-D(*T*)_ to become negative. Thus, the activation of denatured conformers to the TSE is entropically: (*i*) favourable for *T*_α_ ≤ *T* < *T_S_*; (*ii*) unfavourable for *T_S_* < *T* ≤ *T*_ω_; and (*iii*) neutral when *T* = *T_S_*. At *T_S_* the Gibbs barrier to folding is purely due to the difference in enthalpy between the DSE and the TSE with *k_f(T)_* being given by

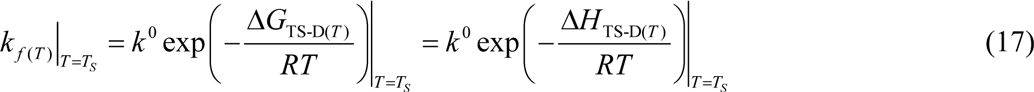

**Figure 10.**
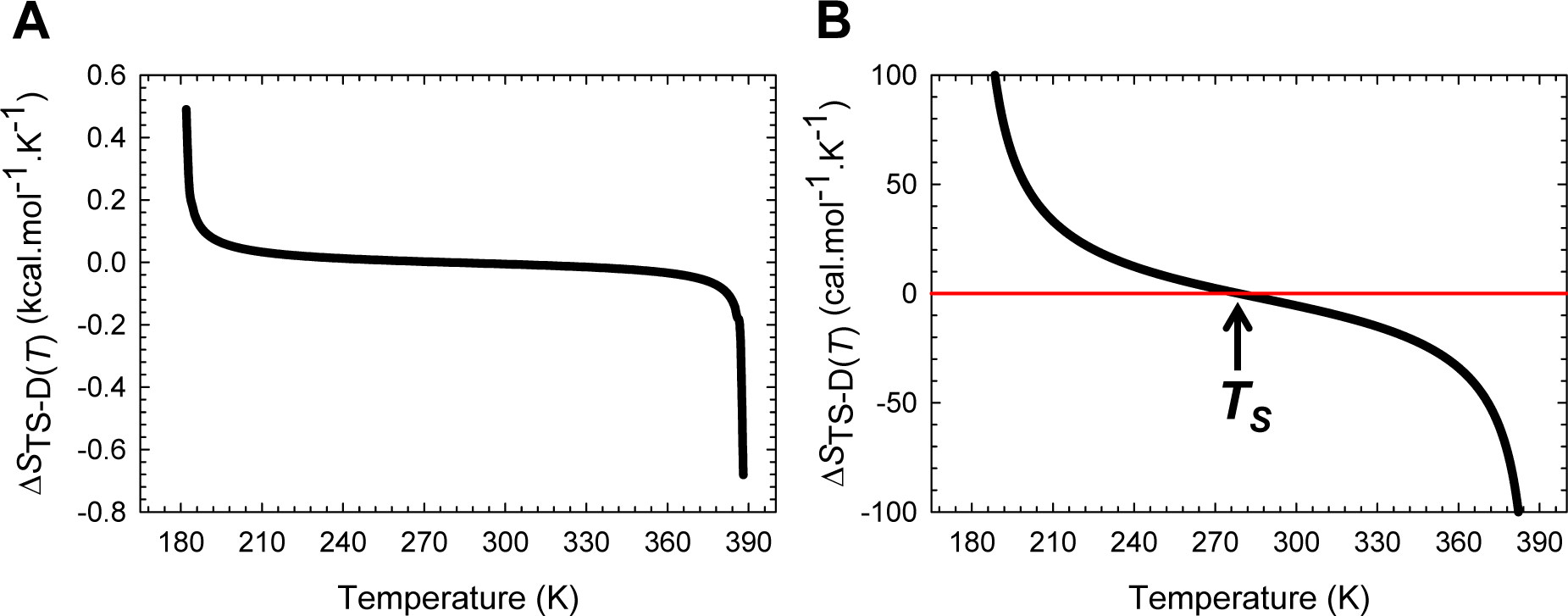
Temperature-dependence of the activation entropy for folding. **(A)** The variation in Δ*S*_TS-D(*T*)_ function with temperature. The slope of this curve varies with temperature and equals Δ*C*_*p*TS-D(*T*)_/*T*. **(B)** An appropriately scaled version of the figure on the left to illuminate the three temperature regimes and their implications: (*i*) Δ*S*_TS-D(*T*)_ > 0 for *T*_*α*_ ≤ *T* < *T_S_*; (*ii*) Δ*S*_TS-D(*T*)_ < 0 for *T_S_* < *T* ≤ *T_ω_*; and (*iii*) Δ*S*_TS-D(*T*)_ = 0 when *T* = *T_S_*.

Inspection of **Figure 11** demonstrates that the behaviour of the Δ*S*_TS-N(*T*)_ function is far more complex than the Δ*S*_TS-D(*T*)_ function: (*i*) Δ*S*_TS-N(*T*)_ > 0 for *T*_α_ ≤ *T* < *T*_*S*(α)_ and *T_S_* < *T* < *T*_*S*(ω)_; (*ii*) Δ*S*_TS-N(*T*)_ < 0 for *T*_*S*(α)_ < *T* < *T_S_* and *T*_*S*(ω)_ < *T* ≤ *T*_ω_; and (*iii*) Δ*S*_TS-N(*T*)_ = 0 at *T*_*S*(α)_, *T_S_*, and *T*_*S*(ω)_. Consequently, we may state that the activation of native conformers to the TSE is entropically: (*i*) favourable for *T*_α_ ≤ *T* < *T*_*S*(α)_ and *T_S_* < *T* < *T*_*S*(ω)_; (*ii*) unfavourable for *T*_*S*(α)_ < *T* < *T_S_* and *T*_*S*(ω)_ < *T* ≤ *T*_ω_; and (*iii*) neutral at *T*_*S*(α)_, *T_S_*, and *T*_*S*(ω)_. If we reverse the reaction-direction (**Figure 11−figure supplement 1A**), the algebraic signs invert leading to a change in the interpretation. Consequently, we may state that for the partial folding reaction [*TS*] ⇌ *N*, the flux of the conformers from the TSE to the NSE is entropically: (*i*) unfavourable for *T*_α_ ≤ *T* < *T*_*S*(α)_ and *T_S_* < *T* < *T*_*S*(ω)_ (Δ*S*_N-TS(*T*)_ < 0); (*ii*) favourable for *T*_*S*(α)_ < *T* < *T_S_* and *T*_*S*(ω)_ < *T* ≤ *T*_ω_ (Δ*S*_N-TS(*T*)_ > 0); and (*iii*) neutral at *T*_*S*(α)_, *T_S_*, and *T*_*S*(ω)_.

**Figure 11.**
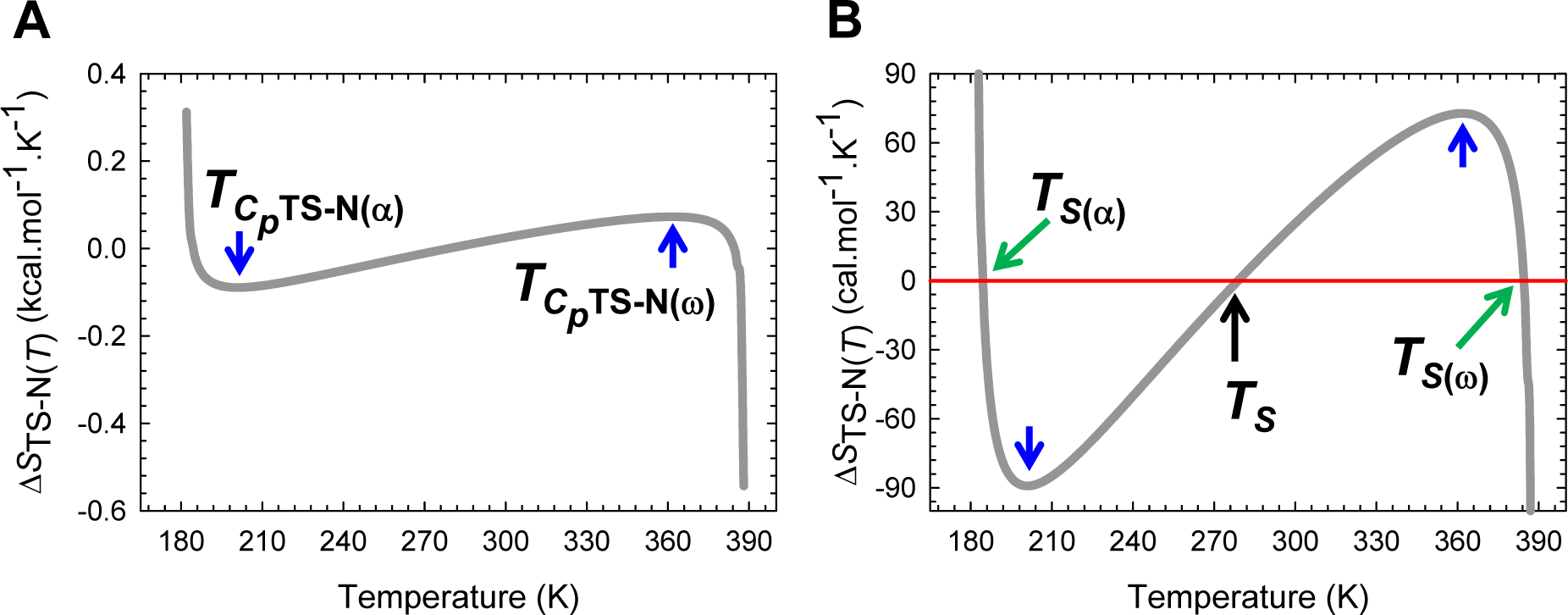
Temperature-dependence of the activation entropy for unfolding. **(A)** The variation in Δ*S*_TS-N(*T*)_ function with temperature. The slope of this curve, given by Δ*C*_*p*TS-N(*T*)_/*T*, varies with temperature, and is zero at *T*_*C_p_*TS-N(α)_ and *T*_*C_p_*TS-N(ω)_. **(B)** An appropriately scaled version of the figure on the left to illuminate the temperature regimes and their implications: (*i*) Δ*S*_TS-N(*T*)_ > 0 for *T_α_* ≤ *T* < *T*_*S*(*α*)_ and *T_S_* < *T* < *T*_*S*(*ω*)_; (*ii*) Δ*S*_TS-N(*T*)_ < 0 for *T*_*S*(*α*)_ < *T* < *T_S_* and *T*_*S*(ω)_ < *T* ≤ *T_ω_*; and (*iii*) Δ*S*_TS-N(*T*)_ = 0 at *T*_*S*(*α*)_, *T_S_*, and *T*_*S*(*ω*)_. Note that at *T*_*S*(*α*)_ and *T*_*S*(*ω*)_, we have the unique scenario: *m*_TS-N(*T*)_ = Δ*G*_TS-N(*T*)_ = Δ*S*_TS-N(*T*)_ = Δ*H*_TS-N(*T*)_ = 0, and *k*_*u*(*T*)_ = *k*^0^. The values of the reference temperatures are given in **Table 1**.

At *T* = *T_S_*, the Gibbs barrier to unfolding is purely due to the difference in enthalpy between the TSE and the NSE (Δ*G*_TS-N(*T*)_ = Δ*H*_TS-N(*T*)_) with *k_u(T)_* being given by

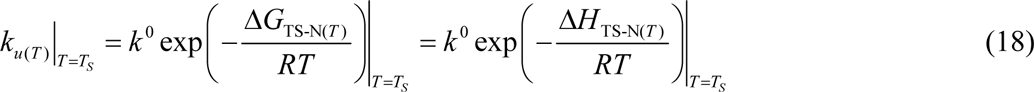

Although Δ*S*_TS-N(*T*)_ = 0 ⇒ *S*_TS(*T*)_ = *S*_N(*T*)_ at *T*_*S*(α)_, *T_S_*, and *T*_*S*(ω)_, the underlying thermodynamics is fundamentally different at *T_S_* as compared to *T*_*S*(α)_ and *T*_*S*(ω)_. While both Δ*G*_TS-N(*T*)_ and *m*_TS-N(*T*)_ are positive and a maximum, and Δ*G*_TS-N(*T*)_ is purely enthalpic at *T_S_* (Δ*G*_TS-N(*T*)_ = Δ*H*_TS-N(*T*)_), at *T*_*S*(α)_ and *T*_*S*(ω)_ we have *m*_TS-N(*T*)_ = 0 ⇒ Δ*G*_TS-N(*T*)_ = ω(*m*_TS-N(*T*)_^2^ = 0 ⇒ Δ*H*_TS-N(*T*)_ = 0, and Δ*G*_N-D(*T*)_ = Δ*G*_TS-D(*T*)_ = λ; and because Δ*G*_TS-N(*T*)_ = 0 at *T*_*S*(α)_ and *T*_*S*(ω)_, the rate constant for unfolding will reach an absolute maximum for that particular solvent and pressure at these two temperatures. To summarize, while at *T_S_* we have *G*_TS(*T*)_ ≫ *G*_N(*T*)_, *S*_D(*T*)_ = *S*_TS(*T*)_ = *S*_N(*T*)_, and *k*_*u*(*T*)_ ≪ *k*^0^, when *T* = *T*_*S*(α)_ and *T*_*S*(ω)_, we have *G*_TS(*T*)_ = *G*_N(*T*)_, *H*_TS(*T*)_ = *H*_N(*T*)_, *S*_TS(*T*)_ = *S*_N(*T*)_, and *k*_*u*(*T*)_ = *k*^0^ (**Figure 11−figure supplements 2** and **3**). Thus, a fundamentally important conclusion that we may draw from these relationships is that “*if two reaction-states on the folding pathway of a two-state system have identical SASA and Gibbs energy under identical environmental conditions, then their absolute enthalpies and entropies must be identical*.” This must hold irrespective of whether or not the two reaction-states have identical, similar or dissimilar structures. We will revisit this scenario when we discuss the heat capacities of activation and the inapplicability of the Hammond postulate to protein folding reactions.

The criteria for two-state folding from the viewpoint of entropy are the following: (*i*) the condition that Δ*S*_D-N(*T*)_ = Δ*S*_TS-N(*T*)_ — Δ*S*_TS-D(*T*)_ must be satisfied at all temperatures; (*ii*) the intersection of Δ*S*_TS-D(*T*)_ and Δ*S*_TS-N(*T*)_ functions calculated directly from the slopes of the temperature-dependent shift in the *curve-crossing* relative to the DSE and the NSE, respectively, must be identical to the independently estimated *T_S_* from equilibrium thermal denaturation experiments (**Figure 11−figure supplements 1B**, **4** and **5**); and (*iii*) both Δ*S*_TS-D(*T*)_ and Δ*S*_TS-N(*T*)_ functions must independently be equal to zero at *T_S_*.

### Temperature-dependence of the Gibbs activation energies

Although the general features of the temperature-dependence of Δ*G*_TS-D(*T*)_ and Δ*G*_TS-N(*T*)_ were described earlier (**Figure 5** and its figure supplements), it is instructive to discuss the same in terms of their constituent enthalpies and entropies.

The determinants of Δ*G*_TS-D(*T*)_ in terms of its activation enthalpy and entropy may be readily deduced by partitioning the entire temperature range over which the two-state system is physically defined (*T*_α_ ≤ *T* ≤ *T*_ω_) into three distinct regimes using four unique reference temperatures: *T*_α_, *T_S_*, *T*_*H*(TS-D)_, and *T*_ω_ (**Figure 12** and **Figure 12−figure supplement 1**). (1) For *T*_α_ ≤ *T* < *T_S_*, the activation of conformers from the DSE to the TSE is entropically favoured (*T*Δ*S*_TS-D(*T*)_ > 0) but is more than offset by the endothermic activation enthalpy (Δ*H*_TS-D(*T*)_ > 0), leading to incomplete compensation and a positive Δ*G*_TS-D(*T*)_ (Δ*H*_TS-D(*T*)_ – *T*Δ*S*_TS-D(*T*)_. When *T* = *T*_*S*_, Δ*G*_TS-D(*T*)_ is a minimum (its lone extremum), and is purely due to the endothermic enthalpy of activation (Δ*G*_TS-D(*T*)_ = Δ*H*_TS-D(*T*)_ > 0. (2) For *T_S_* < *T* < *T*_*H*(TS-D)_, the activation of denatured conformers to the TSE is enthalpically and entropically disfavoured (Δ*H*_TS-D(*T*)_ > 0 and *T*Δ*S*_TS-D(*T*)_< 0) leading to a positive Δ*G*_TS-D(*T*)_. (3) In contrast, for *T*_*H*(TS-D)_ < *T* ≤ *T*_ω_, the favourable exothermic activation enthalpy (Δ*H*_TS-D(*T*)_ < 0) is more than offset by the unfavourable entropy of activation (*T*Δ*S*_TS-D(*T*)_ < 0), leading once again to a positive Δ*G*_TS-D(*T*)_. When *T* = *T*_*H*(TS-D)_, Δ*G*_TS-D(*T*)_ is purely due to the negative change in the activation entropy or the *negentropy* of activation (Δ*G*_TS-D(*T*)_ = –*T*Δ*S*_TS-D(*T*)_ > 0), Δ*G*_TS-D(*T*)_/*T* is a minimum, and *k*_*f*(*T*)_ is a maximum (their lone extrema; see Massieu-Planck functions below). An important conclusion that we may draw from these analyses is the following: While it is true that for the temperature regimes *T*_α_ ≤ *T* < *T_S_* and *T*_*H*(TS-D)_ < *T* ≤ *T*_ω_, Δ*G*_TS-D(*T*)_ is due to the incomplete compensation of the opposing activation enthalpy and entropy, this is clearly not the case for *T_S_* < *T* < *T*_*H*(TS-D)_ where both these two state functions are unfavourable and complement each other to generate a positive Gibbs activation barrier.

**Figure 12.**
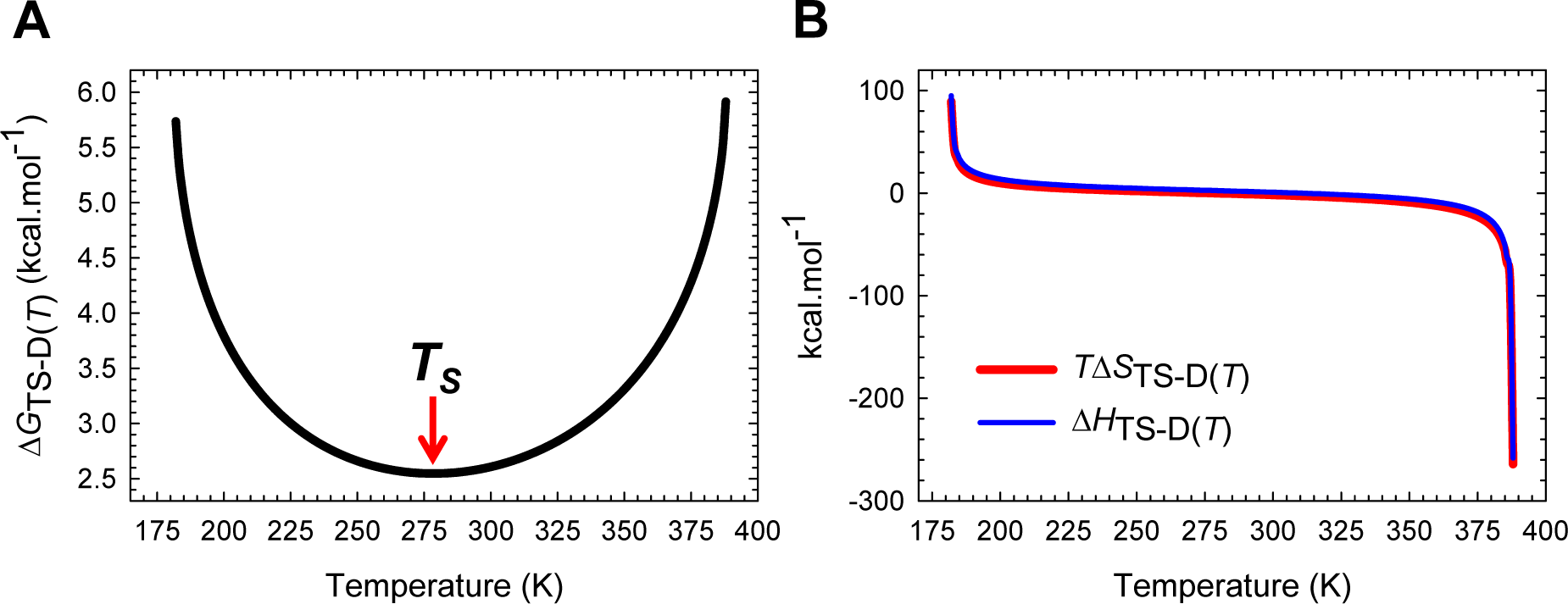
Entropy-enthalpy compensation for the partial folding reaction *D* ⇌ [*TS*]. Despite large changes in Δ*H*_TS-D(*T*)_ (∽ 400 kcal.mol^-1^) Δ*G*_TS-D(*T*)_ which is a minimum at *T_S_*, varies only by ∽3.4 kcal.mol^-1^ due to compensating changes in Δ*S*_TS-D(*T*)_. See the appropriately scaled figure supplement for description. The physical basis for entropy-enthalpy compensation is addressed in the accompanying article.

Similarly, the determinants of Δ*G*_TS-N(*T*)_ in terms of its activation enthalpy and entropy may be readily divined by partitioning the entire temperature range into five distinct regimes using six unique reference temperatures: *T*_α_, *T*_*S*(α)_, *T*_*H*(TS-N)_, *T_S_*, *T*_*S*(ω)_, and *T*_ω_ (**Figure 13** and **Figure 13−figure supplement 1**). (1) For *T*_α_ ≤ *T* < *T*_*S*(α)_, which is the ultralow temperature *Marcus-inverted-regime* for unfolding, the activation of the native conformers to the TSE is entropically favoured (*T*Δ*S*_TS-N(*T*)_ > 0) but is more than offset by the unfavourable enthalpy of activation (Δ*H*_TS-N(*T*)_ > 0) leading to incomplete compensation and a positive Δ*G*_TS-N(*T*)_ (Δ*H*_TS-N(*T*)_ – Δ*T*Δ*S*_TS-N(*T*)_ > 0). When *T* = *T*_*S*(α)_, Δ*S*_TS-N(*T*)_ = Δ*H*_TS-N(*T*)_ = 0 ⇒ Δ*G*_TS-N(*T*)_ = 0. The first extrema of Δ*G*_TS-N(*T*)_ and Δ*G*_TS-N(*T*)_/*T* (which are a minimum), and the first extremum of *k*_*u*(*T*)_ (which is a maximum, *k*_*u*(*T*)_ = *k*^0^) occur at *T*_*S*(α)_. (2) For *T*_*S*(α)_ < *T* < *T_H_*_(TS-N)_, the activation of the native conformers to the TSE is enthalpically favourable (Δ*H*_TS-N(*T*)_ < 0) but is more than offset by the unfavourable negentropy of activation (*T*Δ*S*_TS-N(*T*)_ < 0) leading to Δ*G*_TS-N(*T*)_ > 0. When *T* = *T*_*H*(TS-N)_, Δ*H*_TS-N(*T*)_ = 0 for the second time, and the Gibbs barrier to unfolding is purely due to the negentropy of activation (Δ*G*_TS-N(*T*)_ = –*T*Δ*S*_TS-N(*T*)_ > 0. The second extrema of Δ*G*_TS-N(*T*)_/*T* (which is a maximum) and *k*_*u*(*T*)_ (which is a minimum) occur at *T*_*H*(TS-N)_. (3) For *T*_*H*(TS-N)_ < *T* < *T_S_*, the activation of the native conformers to the TSE is entropically and enthalpically unfavourable (Δ*H*_TS-N(*T*)_ > 0 and *T*Δ*S*_TS-N(*T*)_ < 0) leading to Δ*G*_TS-N(*T*)_ > 0. When *T* = *T_S_*, Δ*S*TS-N(*T*) = 0 for the second time, and the Gibbs barrier to unfolding is purely due to the endothermic enthalpy of activation (Δ*G*_TS-N(*T*)_ = Δ*H*_TS-N(*T*)_ > 0). The second extremum of Δ*G*_TS-N(*T*)_ (which is a maximum) occurs at *T_S_*. (4) For *T_S_* < *T* < *T*_*S*(ω)_, the activation of the native conformers to the TSE is entropically favourable (*T*Δ*S*_TS-N(*T*)_ > 0) but is more than offset by the endothermic enthalpy of activation (Δ*H*_TS-N(*T*)_ > 0) leading to incomplete compensation and a positive Δ*G*_TS-N(*T*)_. When *T* = *T*_*S*(ω)_, Δ*S*_TS-N(*T*)_ = Δ*H*_TS-N(*T*)_ = 0 for the third and the final time, and Δ*G*_TS-N(*T*)_ = 0 for the second and final time. The third extrema of Δ*G*_TS-N(*T*)_ and Δ*G*_TS-N(*T*)_/*T* (which are a minimum), and the third extremum of *k*_*u*(*T*)_ (which is a maximum, *k*_*u*(*T*)_ = *k*^0^) occur at *T*_*S*(ω)_. (5) For *T*_*S*(ω)_< *T* ≤ *T*_ω_, which is the high-temperature *Marcus-inverted-regime* for unfolding, the activation of the native conformers to the TSE is enthalpically favourable (Δ*H*_TS-N(*T*)_ < 0) but is more than offset by the unfavourable negentropy of activation (*T*Δ*S*_TS-N(*T*)_ < 0), leading to Δ*G*_TS-N(*T*)_ > 0. Once again we note that although the Gibbs barrier to unfolding is due to the incomplete compensation of the opposing enthalpies and entropies of activation for the temperature regimes *T*_α_ ≤ *T* < *T*_*S*(α)_, *T*_*S*(α)_ < *T* < *T*_*H*(TS-N)_, *T_S_* < *T* < *T*_*S*(ω)_, and *T*_*S*(ω)_< *T* ≤ *T*_ω_, both the enthalpy and the entropy of activation are unfavourable and collude to generate the Gibbs barrier to unfolding for the temperature regime *T*_*H*(TS-N)_ < *T* < *T_S_*. Thus, a fundamentally important conclusion that we may draw from this analysis is that “*the Gibbs barriers to folding and unfolding are not always due to the incomplete compensation of the opposing enthalpy and entropy*.”

**Figure 13.**
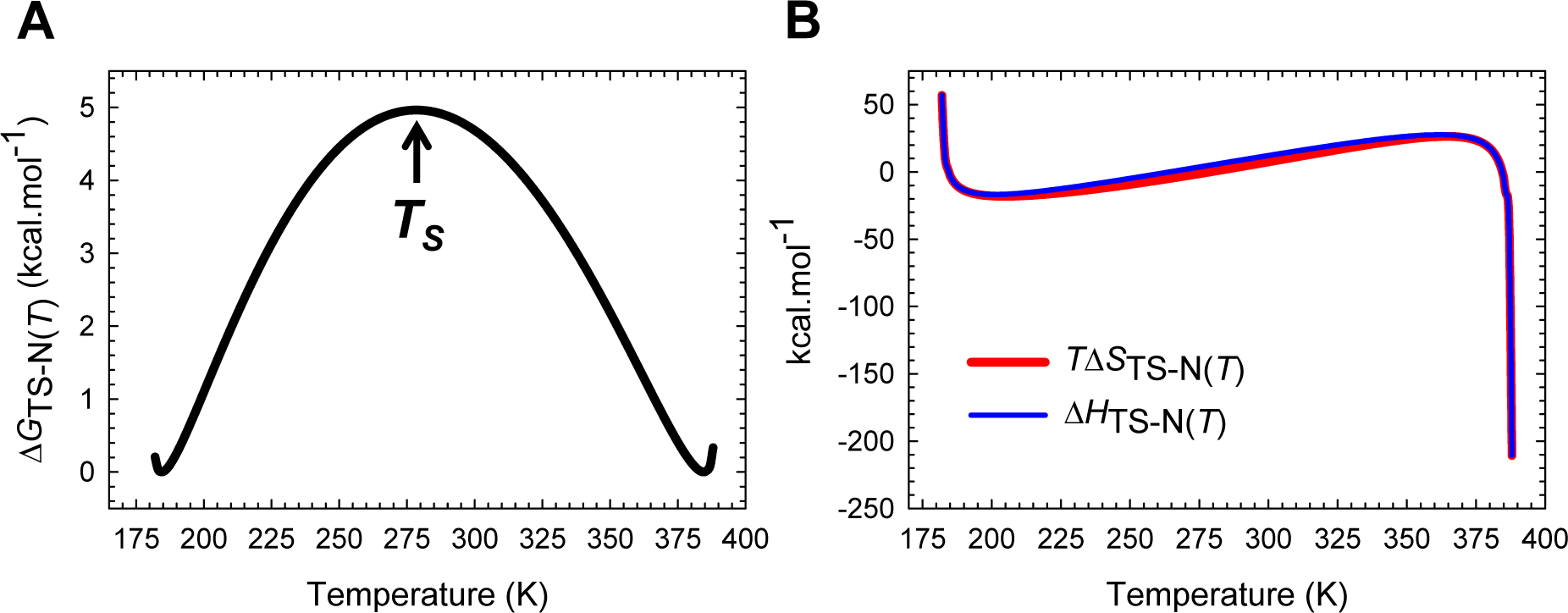
Entropy-enthalpy compensation for the partial unfolding reaction *N* ⇌ [*TS*]. Despite large changes in Δ*H*_TS-N(*T*),_ Δ*G*_TS-N(*T*)_ which is a maximum at *T_S_*, varies only by ∽5 kcal.mol^-1^ due to compensating changes in Δ*S*_TS-N(*T*)_. See the appropriately scaled figure supplement for description.

In a *protein folding* scenario where the activated conformers diffuse on the Gibbs energy surface to reach the NSE, the algebraic signs of the state functions invert leading to a change in the interpretation (**Figure 13−figure supplements 2** and **3**). Thus, for the partial folding reaction[*TS*] ⇌: (1) For *T*_α_ ≤ *T* < *T*_*S*(α)_, the flux of the conformers from the TSE to the NSE is entropically disfavoured (*T*Δ*S*_TS-N(*T*)_ > 0 ⇒ *T*Δ*S*_N-TS(*T*)_ < 0) but is more than compensated by the favourable change in enthalpy (Δ*H*_TS-N(*T*)_ > 0 ⇒ Δ*H*_N-TS(*T*)_ < 0), leading to Δ*G*_N-TS(*T*)_ < 0. (2) For *T*_*S*(α)_ < *T* < *T*_*H*(TS-N)_, the flux of the conformers from the TSE to the NSE is enthalpically unfavourable (Δ*H*_TS-N(*T*)_ < 0 ⇒ Δ*H*_N-TS(*T*)_ > 0) but is more than compensated by the favourable change in entropy (*T*Δ*S*_TS-N(*T*)_ < 0 ⇒ *T*Δ*S*_N-TS(*T*)_ > 0) leading to Δ*G*_N-TS(*T*)_ < 0. When *T* = *T*_*H*(TS-N)_, the flux is driven purely by the positive change in entropy (Δ*G*_N-TS(*T*)_ = –*T*Δ*S*_N-TS(*T*)_ > 0). (3) For *T*_*H*(TS-N)_ < *T* < *T_S_*, the flux of the conformers from the TSE to the NSE is entropically and enthalpically favourable (Δ*H*_N-TS(*T*)_ < 0 and *T*Δ*S*_N-TS(*T*)_ > 0) leading to Δ*G*_N-TS(*T*)_ < 0. When *T* = *T_S_*, the flux is driven purely by the exothermic change in enthalpy (Δ*G*_N-TS(*T*)_ = Δ*H*_N-TS(*T*)_ < 0). (4) For *T_S_* < *T* < *T*_*S*(ω)_, the flux of the conformers from the TSE to the NSE is entropically unfavourable (*T*Δ*S*_TS-N(*T*)_ > 0 ⇒ *T*Δ*S*_N-TS(*T*)_ < 0) but is more than compensated by the exothermic change in enthalpy (Δ*H*_TS-N(*T*)_ > 0 ⇒ Δ*H*_N-TS(*T*)_ < 0) leading to Δ*G*_N-TS(*T*)_ < 0. (5) For *T*_*S*(ω)_< *T* ≤ *T*_ω_, the flux of the conformers from the TSE to the NSE is enthalpically unfavourable (Δ*H*_TS-N(*T*)_ < 0 ⇒ Δ*H*_N__TS(*T*)_ > 0) but is more than compensated by the favourable change in entropy (*T*Δ*S*_TS-N(*T*)_ < 0 ⇒ *T*Δ*S*_N-TS(*T*)_ > 0), leading to Δ*G*_N-TS(*T*)_ < 0.

Thus, the criteria for two-state folding from the viewpoint of Gibbs energy are the following: (*i*) the condition that Δ*G*_D-N(*T*)_ = Δ*G*_TS-N(*T*)_ – Δ*G*_TS-D(*T*)_ must be satisfied at all temperatures; (*ii*) the cold and heat denaturation temperatures estimated from equilibrium thermal denaturation must be identical to independently determined temperatures at which *k*_*f*(*T*)_ and *k*_*u*(*T*)_ are identical, i.e., the temperatures at which Δ*G*_TS-D(*T*)_ and Δ*G*_TS-N(*T*)_ functions intersect must be identical to the temperatures at which Δ*H*_D-N(*T*)_ – *T*Δ*S*_D-N(*T*)_= Δ*G*_D-N(*T*)_ = 0. The basis for these relationships, as mentioned earlier, is the *principle of microscopic reversibility*;^27^ (*iii*) Δ*G*_TS-D(*T*)_ and Δ*G*_TS-N(*T*)_ must be a minimum and a maximum, respectively, at *T_S_*; and (*iv*) the condition that *T*_*H*(TS-N)_ < *T_H_* < *T_S_* < *T*_*H*(TS-D)_ must be satisfied. A far more detailed explanation in terms of chain and desolvation entropies and enthalpies is given in the accompanying article.

### Massieu-Planck functions

The Massieu-Planck function, Δ*G*/*T*, or its equivalent –*R*ln*K* (*K* is the equilibrium constant) predates the Gibbs energy function by a few years and is especially useful when analysing temperature-dependent changes in protein behaviour (see Schellman, 1997, on the use of Massieu-Planck functions to analyse protein folding, and why the use of Δ*G versus T* curves can sometimes lead to ambiguous conclusions).^6,35^ Comparison of **Figure 6−figure supplement 1A** and **Figure 14A** demonstrates that although Δ*G*_TS-D(*T*)_ is a minimum at *T_S_* (**Figure 5A**), *k*_*f*(*T*)_ will be a maximum not at *T_S_* but instead at *T*_*H*(TS-D)_ where the Massieu-Planck activation potential for folding (Δ*G*_TS-D(*T*)_/*T* ≡ –*R* ln *K*_TS-D(*T*)_) is a minimum, and is readily apparent if we recast the Arrhenius expression for *k*_*f*(*T*)_ in terms of the equilibrium constant for the partial folding reaction *D* ⇌ [*TS*].

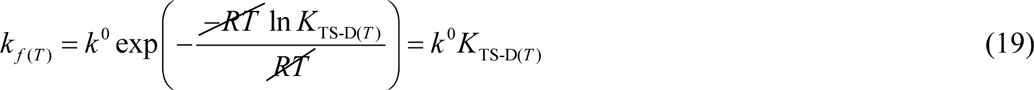

**Figure 14.**
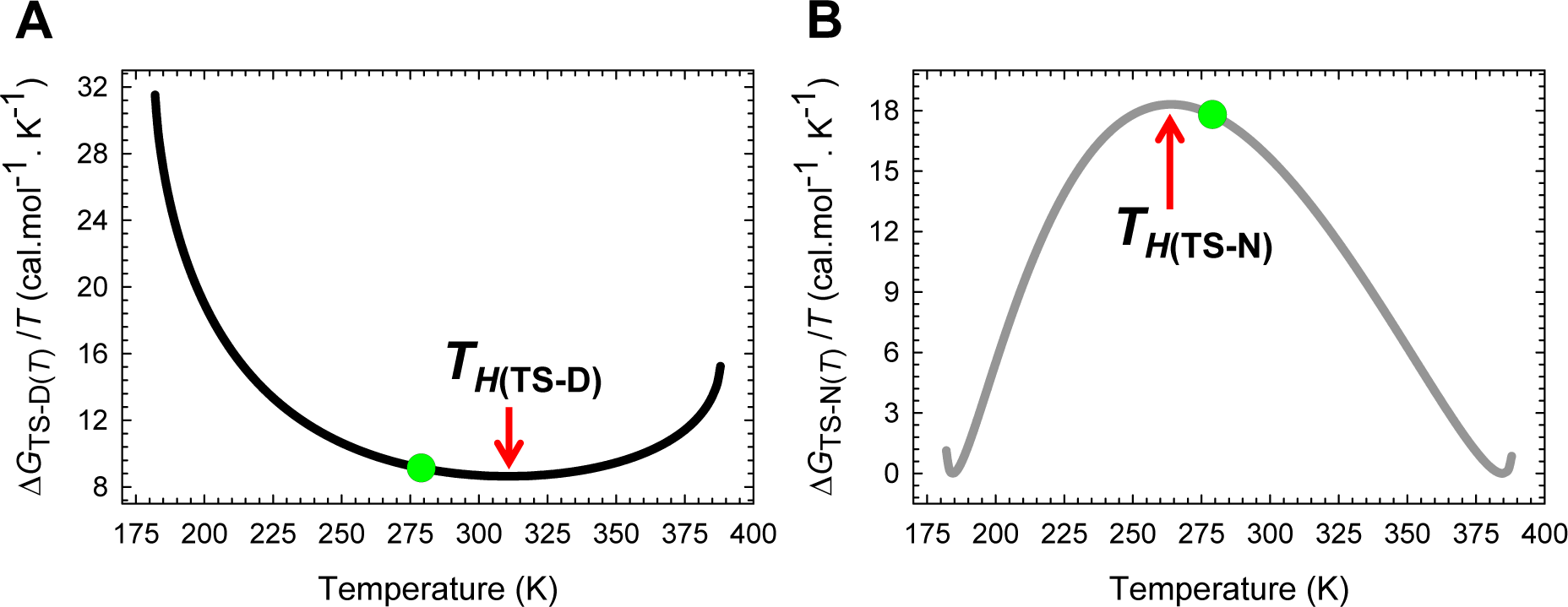
Temperature-dependence of the Massieu-Planck activation potentials. **(A)** The Massieu-Planck activation potential for folding is a minimum at *T*_*H*(TS-D)_. The slope of this curve is given by −Δ*H*_TS-D(*T*)_/*T*^2^. **(B)** The Massieu-Planck activation potential for unfolding is a maximum at *T*_*H*(TS-N)_ and a minimum (zero) at *T*_*S*(α)_ and *T*_*S*(ω)_. The slope of this curve is given by −Δ*H*_TS-N(*T*)_/*T*^2^. The temperature *T_S_* at which Δ*G*_TS-D(*T*)_ and Δ*G*_TS-N(*T*)_ are a minimum and a maximum, respectively, are indicated by green circles.

Eq. (19) shows that the rate determining *K*_TS-D(*T*)_ ([*TS*]/[*D*]) or the population of activated conformers relative to those that nestle at the bottom of the denatured Gibbs energy well is a maximum not at *T_S_* but at *T*_*H(TS-D)*_ (**Figure 14−figure supplement 1A**). Similarly, comparison of **Figure 6−figure supplement 1B** and **Figure 14B** shows that although Δ*G*_TS-N(*T*)_ is a maximum at *T_S_* (**Figure 5B**), the minimum in *k*_*u*(*T*)_ will occur not at *T_S_* but instead at *T*_*H*(TS-N)_ where the Massieu-Planck activation potential for unfolding (Δ*G*_TS-N(*T*)_/*T* ≡ –*R* ln *K*_TS-D(*T*)_) is a maximum (Eq. (20)).

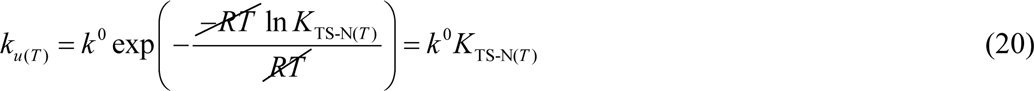

Thus, for the partial unfolding reaction *N* ⇌ [*TS*], the rate determining *K*_TS-N(*T*)_ ([*TS*]/[*N*]) or the population of activated conformers relative to those at the bottom of the native Gibbs basin is a minimum not at *T_S_* but at *T*_*H*(TS-N)_ (**Figure 14−figure supplement 1B**). Similarly, we see that although the Δ*G*_N-D(*T*)_ is a minimum or the most negative at *T*_*S*_ (**Figure 1−figure supplement 1**), *K*_N-D(*T*)_ ([*N*]/[*D*]) is a maximum not at *T_S_* but at *T_H_* where Δ*H*_N-D(*T*)_= 0 and *k*_*f*(*T*)_/*k*_*u*(*T*)_ is a maximum (**Figure 14−figure supplement 2A**).^6^ Because the ratio of the solubilities of any two reaction-states is identical to the equilibrium constant, we may state that for any two-state folder at constant pressure and solvent conditions: (*i*) the solubility of the TSE as compared to the DSE is the greatest when the Gibbs barrier to folding is purely entropic, and this occurs precisely at *T*_*H*(TS-D)_ (**Figure 14−figure supplement 3A**); (*ii*) the solubility of the TSE as compared to the NSE is the least when the Gibbs barrier to unfolding is purely entropic and occurs precisely at *T*_*H*(TS-N)_ (**Figure 14−figure supplement 3B**); (*iii*) the solubilities of the TSE and the NSE are identical at *T*_*S*(α)_ and *T*_*S*(ω)_where Δ*S*_TS-N(*T*)_ = Δ*H*_TS-N(*T*)_ = Δ*G*_TS-N(*T*)_ = 0, and *k*_*u*(*T*)_ = *k*^0^ (**Figure 14−figure supplement 3B**); and (*iv*) the solubility of the NSE as compared to the DSE is the greatest when the net flux of the conformers from the DSE to the NSE is driven purely by the difference in entropy between these two reaction-states and occurs precisely at *T_H_* (**Figure 14−figure supplement 2B**). The notion that “certain aspects of the temperature-dependent protein behaviour are greatly simplified when the Massieu-Planck functions are used in preference to the Gibbs energy” is readily apparent from inspection of **Figure 14−figure supplements 4** and **5**: While the natural logarithms of *k*_*f*(*T*)_ and *k*_*u*(*T*)_ have a complex dependence on their respective Gibbs barriers, a simple linear relationship exists between the rate constants and their respective Massieu-Planck functions.

### Temperature-dependence of Δ*C*_*p*D-TS(*T*)_ and Δ*C*_*p*TS-N(*T*)_

In order to provide a rational explanation for the temperature-dependence of the Δ*C*_*p*D-TS(*T*)_ and Δ*C*_*p*TS-N(*T*)_ functions, it is instructive to first discuss the inter-relationships between ΔSASA_D-N_, *m*_D-N_, and Δ*C*_*p*D-N_. According to the “*liquid-liquid transfer*” model (LLTM) the greater heat capacity of the DSE as compared to the NSE (i.e., Δ*C*_*p*D-N_ > 0 and substantial) is predominantly due to anomalously high heat capacity and low entropy of water that surrounds the exposed non-polar residues in the DSE (referred to as “microscopic icebergs” or “clathrates”; see references in Baldwin, 2014).^36^ Because the size of the solvation shell depends on the SASA of the non-polar solute, it naturally follows that the change in the heat capacity must be proportional to the change in the non-polar SASA that accompanies a reaction. Consequently, protein unfolding reactions which are accompanied by large changes in non-polar SASA lead to large and positive changes in the heat capacity.^33,37,38^ Because the denaturant *m* values are also directly proportional to the change in SASA that accompanies protein unfolding reactions, the expectation is that *m*_D-N_ and Δ*C*_*p*D-N_ values must also be proportional to each other: The greater the *m*_D-N_ value, the greater is the Δ*C*_*p*D-N_ value and *vice versa* (Figs. 2, 3 and 5 in Myers et al., 1995). However, since the residual structure in the DSEs of proteins under folding conditions is both sequence and solvent-dependent (i.e., the SASAs of the DSEs two proteins of identical chain lengths but dissimilar primary sequences need not necessarily be the same even under identical solvent conditions),^39,40^ and because we do not yet have reliable theoretical or experimental methods to accurately quantify the SASA of the DSEs of proteins under folding conditions (i.e., the values are model-dependent),^41-43^ the data scatter in plots that show correlation between the experimentally determined *m*_D-N_ or Δ*C*_*p*D-N_ values (which reflect the true ΔSASA_D-N_) and the calculated values of ΔSASA_D-N_ can be significant (Fig. 2 in Myers et al., 1995, and Fig. 3 in Robertson and Murphy, 1997). Now, since the solvation shell around the DSEs of large proteins is relatively greater than that of small proteins even when the residual structure in the DSEs under folding conditions is taken into consideration, large proteins on average expose relatively greater amount of non-polar SASA upon unfolding than do small proteins; consequently, both *m*_D-N_ and Δ*C*_*p*D-N_ values also correlate linearly with chain-length, albeit with considerable scatter since chain length, owing to the residual structure in the DSEs, is unlikely to be a true descriptor of the SASA of the DSEs of proteins under folding conditions (note that the scatter can also be due to certain proteins having anomalously high or low number of non-polar residues). The point we are trying to make is the following: Because the native structures of proteins are relatively insensitive to small variations in pH and co-solvents,^44^ and since the number of ways in which foldable polypeptides can be packed into their native structures is relatively limited (as inferred from the limited number of protein folds, see SCOP: www.mrc-lmb.cam.ac.uk and CATH: www.cathdb.info databases), one might find a reasonably good correlation between chain lengths and the SASAs of the NSEs of proteins of differing primary sequences under varying solvents (Fig. 1 in Miller et al., 1987).^45,46^ However, since the SASAs of the DSEs under folding conditions, owing to residual structure are variable, until and unless we find a way to accurately simulate the DSEs of proteins, and if and only if these theoretical methods are sensitive to point mutations, changes in pH, co-solvents, temperature and pressure, it is almost impossible to arrive at a universal equation that will describe how the ΔSASA_D-N_ under folding conditions will vary with chain length, and by logical extension, how *m*_D-N_ and Δ*C*_*p*D-N_ will vary with SASA or chain length. Nevertheless, if we consider a single two-state-folding primary sequence under constant pressure and solvent conditions and vary the temperature, and if the properties of the solvent are temperature-invariant (for example, no change in the pH due to the temperature-dependence of the p*K_a_* of the constituent buffer), then the manner in which the Δ*C*_*p*D-TS(*T*)_ and Δ*C*_*p*TS-N(*T*)_ functions vary with temperature must be consistent with the temperature-dependence of *m*_TS-D(*T*)_ and *m*_TS-N(*T*)_, respectively, and by logical extension, with ΔSASA_D-TS(*T*)_ and ΔSASA_TS-N(*T*)_, respectively.

Inspection of **Figures 15** and **Figure 15−figure supplements 1**, **2** and **3** demonstrate that: (*i*) both Δ*C*_*p*D-TS(*T*)_ and Δ*C*_*p*TS-N(*T*)_ vary with temperature; and (*ii*) their gross features stem primarily from the second derivatives of the temperature-dependence of the *curve-crossing* with respect to the DSE and the NSE. The prediction that the change in heat capacities for the partial unfolding reactions, *N* ⇌ [*TS*] and [*TS*] ⇌ *D*, must vary with temperature is due to Eqs. (12) and (13). Although this may not be readily apparent from a casual inspection of the equations, even a cursory examination of **Figures 8** and **9** shows that it is simply not possible for Δ*C*_*p*D-TS(*T*)_ and Δ*C*_*p*TS-N(*T*)_ functions to be temperature-invariant since the slopes of the Δ*H*TS-D(*T*) and the Δ*H*_TS-N(*T*)_ functions are continuously changing with temperature. If we recall that the force constants are temperature-invariant, it becomes readily apparent that the second terms in the brackets on the right-hand-side (RHS) of Eqs. (12) and (13) i.e., ω*T*(Δ*S*_D-N(*T*)_ and α*T*(Δ*S*_D-N(*T*)_)^2^, respectively, will be parabolas with a minimum (zero) at *T_S_*. This is due to Δ*S*_D-N(*T*)_ being negative for *T* < *T_S_*, positive for *T* > *T_S_*, and zero for *T* = *T_S_*. Furthermore, since φ, 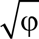 and *m*_TS-N(*T*)_ are a maximum, and *m*_TS-D(*T*)_ a minimum at *T_S_*, the expectation is that Δ*C*_*p*D-TS(*T*)_ must be a minimum (or Δ*C*_*p*TS-D(*TS*)_ is the least negative), and Δ*C*_*p*TS-N(*T*)_ must be a maximum at *T_S_*. Thus, for *T* = *T_S_*, Eqs. (12) and (13) become

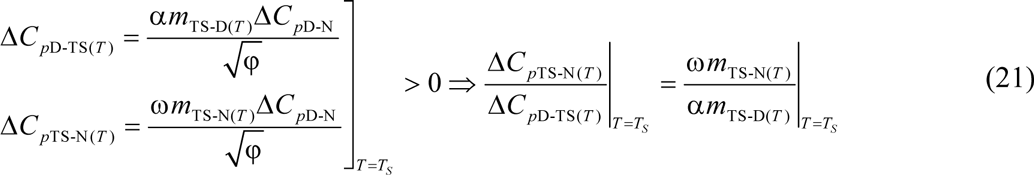

The prediction that the extrema of Δ*C*_*p*D-TS(*T*)_ and Δ*C*_*p*TS-N(*T*)_ functions must occur at *T_S_* is readily apparent from **Figure 15** and **Figure 15−figure supplement 1B**. Importantly, consistent with the relationship between *m*_D-N_ and Δ*C*_*p*D-N_ values, comparison of these two figures with **Figure 2** and **Figure 2−figure supplement 1** demonstrates that just as *m*_TS-D(*T*)_ and *m*_TS-N(*T*)_ are a minimum and a maximum at *T_S_*, respectively, so too are Δ*C*_*p*D-TS(*T*)_and Δ*C*_*p*TS-N(*T*)_ functions. This leads to two obvious corollaries: (*i*) the difference in heat capacity between the DSE and the TSE is a minimum when the difference in SASA between the DSE and the TSE is a minimum; and (*ii*) the difference in heat capacity between the TSE and the NSE is a maximum when the difference in SASA between the TSE and the NSE is a maximum. Because Δ*S*_TS-D(*T*)_ = Δ*S*_TS-N(*T*)_ = 0, Δ*G*_TS-D(*T*)_ is a minimum, and both Δ*G*_TS-N(*T*)_ and Δ*G*_D-N(*T*)_ are a maximum, at *T_S_* (**Figures 1, 5** and **Figure 11−figure supplement 1B**), a fundamentally important conclusion is that the *Gibbs barriers to folding and unfolding are a minimum and a maximum, respectively, and equilibrium stability is a maximum, and are all purely enthalpic when* Δ*C*_*p*D-TS(*T*)_ *and* Δ*C*_*p*TS-N(*T*)_ *are a minimum and a maximum, respectively*.

**Figure 15.**
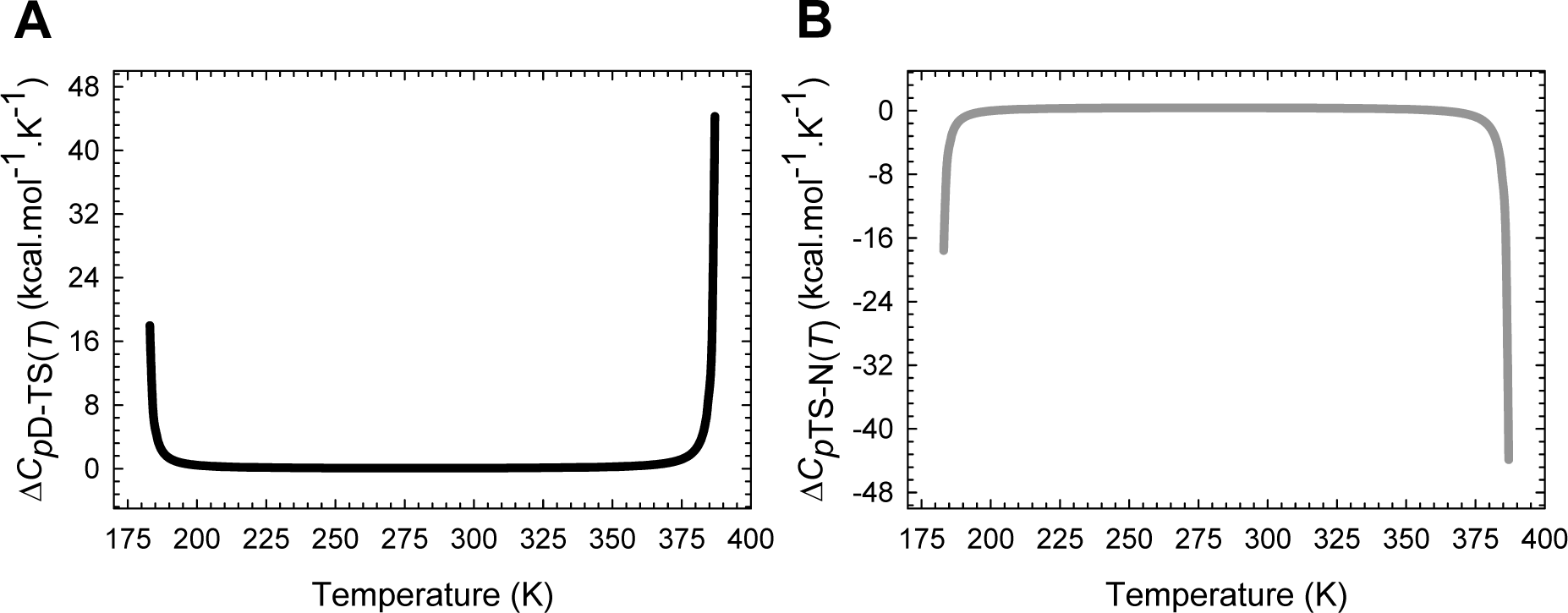
Temperature-dependence of Δ*C*_*p*D-TS(*T*)_ and Δ*C*_*p*TS-N(*T*)_ functions. **(A)** Δ*C*_*p*D-TS(*T*)_ is positive throughout the temperature range and is a minimum at *T_S_* (or Δ*C*_*p*TS-D(*T*)_ is a maximum or the least negative at *T_S_*). **(B)** Δ*C*_*p*TS-N(*T*)_ is a maximum at *T_S_*, positive for *T*_*C_p_*TS-N(α)_ < *T* < *T*_*C_p_*TS-N(ω)_, negative for *T*_α_ ≤ *T* < *T*_*C_p_*TS-N(α)_ and *T*_*C_p_*TS-N(ω)_ < *T* ≤ *T*_ω_, and as described earlier, zero at *T*_*C_p_*TS-N(α)_ and *T*_*C_p_*TS-N(ω)_. These aspects can be better appreciated from the appropriately scaled views shown in the figure supplement. Note that the algebraic sum of Δ*C*_*p*D-TS(*T*)_ and Δ*C*_*p*TS-N(*T*)_ must equal Δ*C*_*p*D-N_ throughout the temperature-regime.

Inspection of **Figure 15** and **Figure 15−figure supplement 1** demonstrates that unlike Δ*C*_*p*D-TS_(*T*) which is positive across the entire temperature range, Δ*C*_*p*TS-N(*T*)_ which is a maximum and positive at *T_S_*, decreases with any deviation in temperature from *T_S_*, and is zero at *T*_C_p_TS-N(α)_ and *T*_*C*_*p*_TS-N(ω)_; consequently, Δ*C*_*p*TS-N(*T*)_ < 0 for *T*_α_ ≤ *T* < *T*_*C*_*p*_TS-N(α)_ and *T*_*C*_p_TS-N(ω)_ < *T* ≤ *T*_ω_. The reason for this behaviour is apparent from inspection of **Figures 9** and **11**: The slope of the Δ*H*_TS-N(*T*)_ and Δ*S*_TS-N(*T*)_ functions becomes zero at *T*_*C*_*p*_TS-N(α)_ and *T*_*C*_p_TS-N(ω)_; and any further decrease or increase in temperature, respectively, causes the slope to invert. This can be mathematically shown as follows: Since *m*_TS-N(*T*)_ = 0 at *T*_*S*(α)_ and *T*_*S*(ω)_, we have 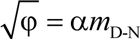 and φ = (α*m*_D-N_)^2^ at *T*_*S*(α)_ and *T*_*S*(ω)_. Substituting these relationships in Eq. (13) leads to

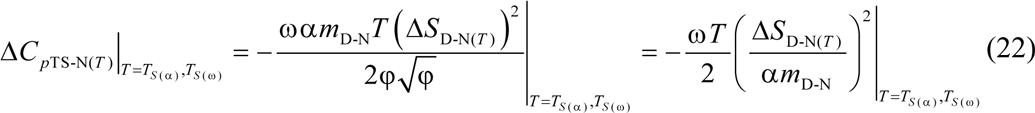

Further, since Δ*C*_*p*D-N_ = Δ*C*_*p*D-TS(*T*)_ + Δ*C*_*p*TS-N(*T*)_ for a two-state system, we have

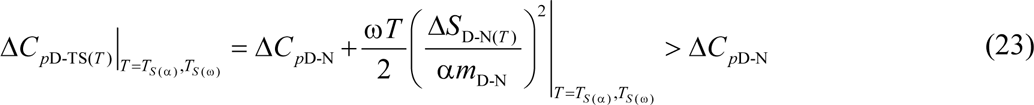

Because Δ*C*_*p*TS-N(*T*)_ < 0 at *T*_*S*(α)_ and *T*_*S*(ω)_, and the lone extremum of Δ*C*_*p*TS-N(*T*)_ (which is algebraically positive and a maximum) occurs at *T_S_*, it implies that there will be two unique temperatures at which Δ*C*_*p*TS-N(*T*)_ = 0, one in the low temperature (*T*_*C*_*p*_TS-N(α)_) such that *T*_*S*(α)_ < *T*_*C*_*p*_TS-N(α)_ < *T_S_*, and the other in the high temperature regime (*T*_*C*_*p*_TS-N(ω)_) such that *T_S_* < *T*_*C*_*p*_TS-N(ω)_ < *T*_*S*(ω)_. Thus, at the these two unique temperatures *T*_*C*_*p*_TS-N(α)_ and *T*_*C*_*p*_TS-N(ω)_, we have Δ*C*_*p*D-TS(*T*)_ = Δ*C*_*p*D-N_ ⇒ β_H(fold)(*T*)_ = 1 and *β*H(unfold)(*T*) = 0; and for the temperature regimes *T*_α_ ≤ *T* < *T*_*C*_*p*_TS-N(α)_ and *T*_*C*_*p*_TS-N(ω)_ < *T* ≤ *T*_ω_, we have Δ*C*_*p*D-TS(*T*)_ > Δ*C*_*p*D-N_ ⇒ β_H(fold)(*T*)_ > 1, and Δ*C*_*p*TS-N(*T*)_ < 0 ⇒ β_H(unfold)(*T*)_ < 0 (see heat capacity RC below for the definition of β_H(fold)(*T*)_ and β_H(unfold)(*T*)_).

Although the prediction that Δ*C*_*p*TS-N(*T*)_ must approach zero at very low and high temperatures may not be readily verified by experiment for the low-temperature regime owing to technical difficulty in making a measurement, the prediction for the high-temperature regime is strongly supported by the data on CI2 from the Fersht lab: Despite the temperature-range not being substantial (320 to 340 K), and the data points that define the Δ*H*_TS-N(*T*)_ function being sparse (7 in total), it is apparent even from a cursory inspection that it is clearly non-linear with temperature (Fig. 5B in Tan et al., 1996).^47^ Although Fersht and co-workers have fitted the data to a linear function and reached the natural conclusion that the heat capacity of activation for unfolding is temperature-invariant, they nevertheless explicitly mention that if the non-linearity of Δ*H*_TS-N(*T*)_ were given due consideration, and the data are fit to an empirical-quadratic instead of a linear function, Δ*C*_*p*TS-N(*T*)_ indeed becomes temperature-dependent and is predicted to approach zero at ∽ 360 K (see text in page 382 in Tan et al., 1996).^47^ Now, since Δ*C*_*p*TS-N(*T*)_ > 0 and a maximum, and Δ*C*_*p*D-TS(*T*)_ is a minimum and positive at *T_S_*, and decrease and increase, respectively, with any deviation in temperature from *T_S_*, and since Δ*C*_*p*TS-N(*T*)_ becomes zero at *T*_*C*_*p*_TS-N(α)_ and *T*_*C*_*p*_TS-N(ω)_, the obvious mathematical consequence is that Δ*C*_*p*D-TS(*T*)_ and Δ*C*_*p*TS-N(*T*)_ functions must intersect at two unique temperatures. Because at the points of intersection we have the relationship: Δ*C*_*p*D-TS(*T*)_ = Δ*C*_*p*TS-N(*T*)_ = Δ*C*_*p*D-N_/2, a consequence is that Δ*C*_*p*TS-N(*T*)_ must be positive at the said temperatures, with the low-temperature intersection occurring between *T*_*C*_*p*_TS-N(α)_ and *T_S_*, and the high-temperature intersection between *T_S_* and *T*_*C*_*p*_TS-N(ω)_. This is readily apparent from inspection of **Figure 15−figure supplement 1B:** Both Δ*C*_*p*D-TS(*T*)_ and Δ*C*_*p*TS-N(*T*)_ are identical at 214.1 K and 345.9 K. An equivalent interpretation is that at these temperatures, the absolute heat capacity of the TSE is exactly half the algebraic sum of the absolute heat capacities of the DSE and the NSE. As we shall show in subsequent publications, the intersection of various state functions is a source of interesting relationships that may be used as constraints in simulations (see also **Figure 9−figure supplement 2**).

### The position of the TSE along the heat capacity RC

Inspection and comparison of **Figure 2−figure supplement 1** and **Figure 15−figure supplement 1B** demonstrates that although the manner in which the Δ*C_p_*D-TS(*T*) and Δ*C*_*p*TS-N(*T*)_ functions vary with temperature is consistent with the relationship between *m*_D-N_ and Δ*C*_*p*D-N_ values, there is nevertheless an intriguing anomaly that is at odds with the LLTM for heat capacity. If we consider the partial folding reaction *D* ⇌ [TS], it is readily apparent from these figures that although the denatured conformer diffuses > ∽ 70% along the normalized SASA-RC to reach the TSE for 240 K < *T* < 320 K, Δ*C*_*p*D-TS(*T*)_ ≪ Δ*C*_*p*TS-N(*T*)_ throughout this regime. Conversely, if we consider the total unfolding reaction *N* ⇌ *D*, a large fraction of Δ*C*_*p*D-N_ is accounted for not by the second-half of the unfolding reaction ([*TS*]) ⇌ *D* but by the first-half ( *N* ⇌ [*TS*]), despite the native conformer diffusing less than ∽30% along the SASA-RC to reach the TSE. To put things into perspective, we will need to normalize the heat capacities of activation. Adopting Leffler’s framework for the relative sensitivities of the activation and equilibrium enthalpies in response to a perturbation in temperature,^48^ we may write

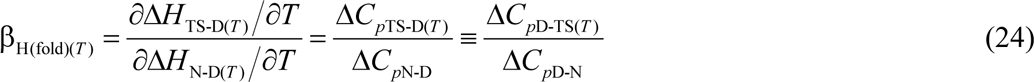

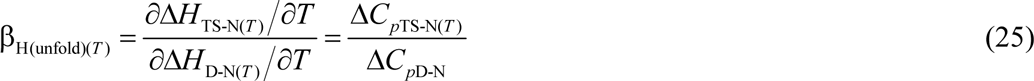

where β_H(fold)(*T*)_ = β_S(fold)(*T*)_ and β_H(unfold)(*T*)_ = β_S(unfold)(*T*)_ (see Paper-II) are classically interpreted to be a measure of the position of the TSE along the heat capacity RC.^49^ Naturally, for a two-state system the algebraic sum of β_H(fold)(*T*)_ and β_H(unfold)(*T*)_ is unity. Recasting Eqs. (24) and (25) in terms of (12) and (13) gives

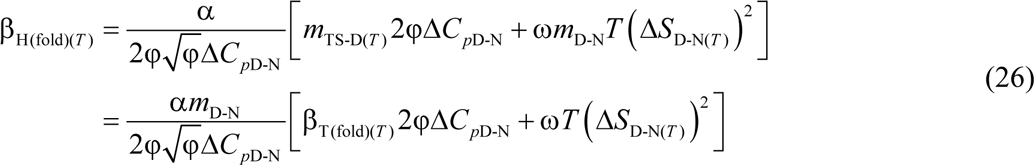

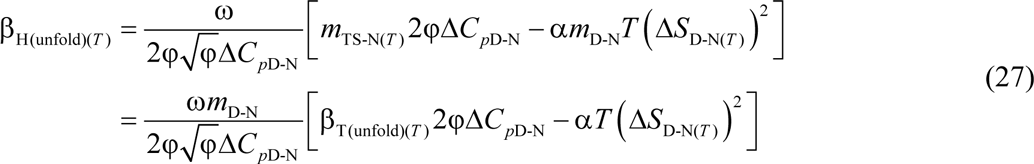

When *T* = *T_S_*, Δ*S*_D-N(*T*)_ = 0 and Eqs. (26) and (27) reduce to

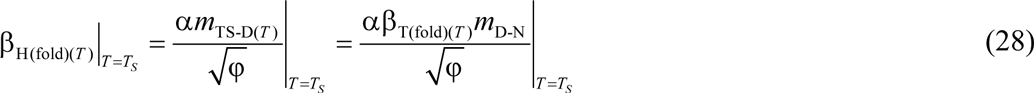

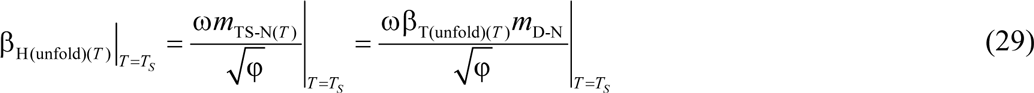

As explained earlier, because Δ*C*_*p*D-N_ is temperature-invariant by postulate, and Δ*C*_*p*D-TS(*T*)_ is a minimum, and Δ*C*_*p*TS-N(*T*)_ is a maximum at *T*_*S*_, β_H(fold)(*T*)_ and β_H(unfold)(*T*)_ are a minimum and a maximum, respectively, at *T*_*S*_. How do β_H(fold)(*T*)_ and β_H(unfold)(*T*)_ compare with their counterparts, β_T(fold)(*T*)_ and β_T(unfold)(*T*)_? This is important because a statistically significant correlation exists between *m*_D-N_ and Δ*C*_*p*D-N_, and both these two parameters independently correlate with ΔSASA_D-N_. Recasting Eqs. (28) and (29) gives

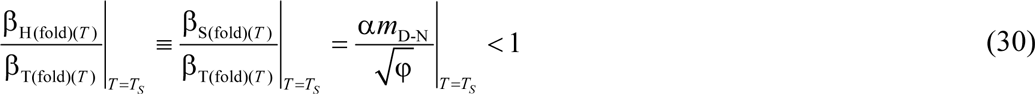

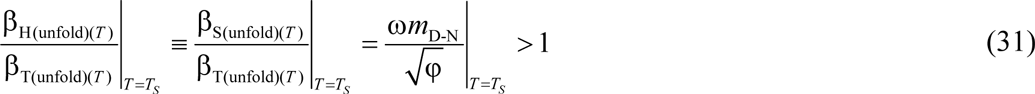

Since *m*_TS-N(*T*)_ > 0 and a maximum, and *m*_TS-D(*T*)_ > 0 and a minimum, respectively, at *T_S_*, it is readily apparent from inspection of Eqs. (1) and (2) that 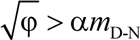 and 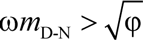 at *T_S_*. Consequently, we have: β_T(fold)(*T*)_|_*T*=*T_S_*_ > β_H(fold)(*T*)_|_*T*=*T_S_*_ and β_T(unfold)(*T*)_|_*T*=*T_S_*_ < β_H(unfold)(*T*)_|_*T*=*T_S_*_.

In agreement with the predictions of Eqs. (30) and (31), inspection of **Figure 16** demonstrates that although the denatured conformer advances by > ∽ 70% along the SASA-RC to reach the TSE when *T* = *T_S_*, it accounts for < ∽20% of the total change in Δ*C*_*p*D-N_ (i.e., β_T(fold)(*T*)_|_*T*=*T_S_*_ > β_H(fold)(*T*)_|_*T*=*T_S_*_), with the rest of the change (> ∽ 80%) in heat capacity coming from a mere ∽ 30% diffusion of the activated conformer along the SASA-RC to reach the bottom of the native Gibbs basin (i.e., β_T(unfold)(*T*)_|_*T*=*T_S_*_ < β_H(unfold)(*T*)_|_*T*=*T_S_*_). The theoretical prediction that β_T(fold)(*T*)_ > β_H(fold)(*T*)_ across a substantial temperature range is supported by the finding by Gloss and Matthews (1998) that the position of the TSE relative to the DSE along the heat capacity RC is consistently lower than the same along the SASA-RC (see also page 178 in Bilsel and Matthews, 2000, and references therein).^50,51^

**Figure 16.**
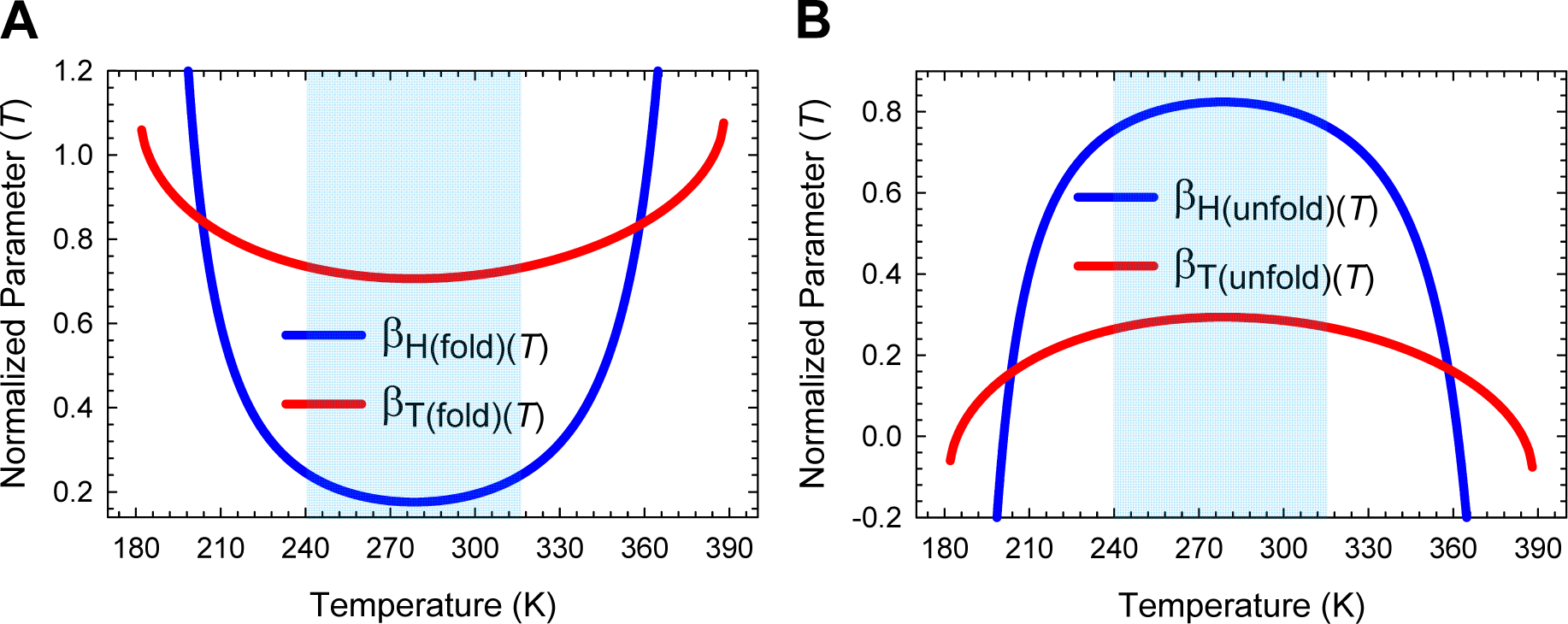
Comparison of the position of the TSE along the heat capacity RC and the SASA-RC. **(A)** Although the temperature-dependences of β_H(fold)(*T*)_ is consistent with β_T(fold)(*T*)_, and both are a minimum at *T_S_*, their magnitudes are not even remotely similar across a large temperature regime (240 K < *T* < 320 K, shaded area); and when *T* = *T_S_*, β_T(fold)(*T*)_ is four fold greater than β_H(fold)(*T*)_ (0.7063/0.1759 = 4.0). Note that the position of the TSE relative to the DSE along the heat capacity and SASA-RCs are identical at the points of intersection (203.6 K and 358.3 K). **(B)** Although the temperature-dependence of β_H(unfold)(*T*)_ is consistent with β_T(unfold)(*T*)_, and both are a maximum at *T_S_*, β_H(unfold)(*T*)_ > β_T(unfold)(*T*)_ for 240 K < *T* < 320 K; and when *T* = *T_S_*, β_H(unfold)(*T*)_ is ∽2.8 fold greater than β_T(unfold)(*T*)_ (0.8241/0.2937 = 2.81). The position of the TSE relative to the NSE along the heat capacity and SASA-RCs are identical at the points of intersection (203.6 K and 358.3 K). See **Figure 16−figure supplement 1** for unscaled plots of β_H(fold)(*T*)_ and β_H(unfold)(*T*)_.

Now, if we accept the long held premise that the greater heat capacity of the DSE as compared to the NSE is purely or predominantly due to structured water around the exposed non-polar residues in the DSE, then the only way we can explain why Δ*C*_*p*D-TS(*T*)_ ≪ Δ*C*_*p*TS-N(*T*)_ despite β_T(fold)(*T*)_ > ∽70% for the partial folding reaction *D* ⇌ [*TS*] is that the non-polar SASA of both the DSE and the TSE are very similar at *T_S_*. Because it is physically near-impossible for the denatured conformer to advance by > ∽ 70% along the SASA-RC to reach the TSE, and yet keep the non-polar SASA fairly constant such that Δ*C*_*p*D-TS(*T*)_ is just about 20% of Δ*C*_*p*D-N_, the natural conclusion is that “*the large and positive difference in heat capacity between the DSE and the NSE cannot be only due to the clathrates of water molecules around exposed non-polar residues in the DSE*.”^38,52-54^ This brings us to two studies on the heat capacities of proteins, one by Sturtevant almost four decades ago, and the other by Lazaridis and Karplus.^55,56^ While Sturtevant identified six possible sources of heat capacity which are: (*i*) the hydrophobic effect; (*ii*) electrostatic charges; (*iii*) hydrogen bonds; (*iv*) conformational entropy; (*v*) intramolecular vibrations; and (*vi*) changes in equilibria, and concluded that the most important of these are the *hydrophobic*, *conformational* and *vibrational* effects, Lazaridis and Karplus concluded from their molecular dynamics simulations on truncated CI2 that the heat capacity can have a significantly large and a positive contribution from intra-protein non-covalent interactions. What these two studies essentially imply is that when the pressure and solvent properties are defined and temperature-invariant, the ability of the conformers in a protein reaction-state to absorb thermal energy and yet resist an increase in temperature is dependent on: (*i*) its molecular structure; and (*ii*) the size and the character of its molecular surface (i.e., the relative proportion of polar and non-polar SASA). While the first variable determines the capacity of the reaction-state to absorb thermal energy and distribute it across its various internal modes of motion (the vibrational, rotational, and to some extent, the translational entropy from elements such as the N and C-terminal regions, loops etc. that can flap around in the solvent), the second variable determines not only the size and thickness of the solvent shell but also how tightly or loosely the solvent molecules are bound to the protein surface and to themselves (i.e., the dynamics of water in the solvation shell as compared to bulk water; see Fig. 1 in Frauenfelder et al., 2009), and by extension, the amount of excess thermal energy needed to disrupt the solvent shell as the reaction-states interconvert due to thermal noise.^36,52,57-61^ Further discussion on the determinants of heat capacity is beyond the scope of this article and will be addressed elsewhere.

### On the inapplicability of the Hammond postulate to protein folding

Although it is difficult to provide a detailed physical explanation for the temperature-dependence of the heat capacities of activation without deconvoluting the activation enthalpies and entropies into their constituent *chain* and *desolvation* enthalpies and entropies (shown in the accompanying article), it is instructive to give one extreme example to emphasize why both the solvent shell and the non-covalent interactions make a significant contribution to heat capacity (note that as long as the difference in the number of covalent bonds between the reaction-states is zero, to a first approximation, their contribution to the *difference in heat capacity* between the reaction-states can be ignored; see Lecture II in Finkelstein and Ptitsyn, 2002, and references therein).^38,56,62,63^

It was shown earlier that when *T* = *T*_*S*(α)_ and *T*_*S*(ω)_, we have *m*_TS-N(*T*)_ = 0 ⇒ ΔSASA_TS-N(*T*)_ = 0, leading to a unique set of relationships: *G*_TS(*T*)_ = *G*_N(*T*)_, *H*_TS(*T*)_ = *H*_N(*T*)_, *S*_TS(*T*)_ = *S*_N(*T*)_, and *k*_*u*(*T*)_ = *k*^0^ (**Figures 2B, Figure 2−figure supplement 1B, 4C, 5B, 6B, 9,** and **11**). However, we note from Eq. (22) that Δ*C*_*p*TS-N(*T*)_ < 0 at these two temperatures and is ∽ −6.2 kcal.mol^-1^.K^-1^ for FBP28 WW (**Figure 15B**). Since the molar concentration of the TSE is identical to that of the NSE at *T*_*S*(α)_ and *T*_*S*(ω)_, what this physically means is that if we were to take a mole of NSE and a mole of TSE and heat them at constant pressure under identical solvent conditions, we will find that the NSE, relative to the TSE, will absorb thermal energy equivalent to ∽6.2 calories before both the TSE and the NSE will independently register a 10^−3^ K rise in temperature. Because at these two temperatures the SASA, the Gibbs energy, the enthalpy, and the entropy of the TSE and the NSE are identical, this large difference in heat capacity which is ∽15-fold greater than Δ*C*_*p*D-N_ (6.2/0.417 = 14.8) must stem from a complex combination of: (*i*) a difference in the number and kinds of non-covalent interactions;^64^ (*ii*) the precise 3D-arrangement of the non-covalent interactions (i.e., the network of interactions) leading to a difference in their fundamental frequencies;^55,56^ and (*iii*) the character of the surface exposed to the solvent (i.e., polar *vs* non-polar SASA) between the said reaction-states.^65-67^ Thus, a fundamentally important conclusion that we may draw from this behaviour is that “*two reaction-states on a protein folding pathway need not necessarily have the same structure even if their interconversion proceeds with concomitant zero net-change in SASA, enthalpy, entropy, and Gibbs energy*.” A corollary is that the reaction-states on a protein folding pathway are distinct entities with respect to both their internal structure and the character of their molecular surface. What this implies is that the Hammond postulate which states that “*if two states, as for example, a transition state and an unstable intermediate, occur consecutively during a reaction process and have nearly the same energy content, their interconversion will involve only a small reorganization of the molecular structures*,”^68^ although may be applicable to reactions of small molecules, is inapplicable to protein folding. The inapplicability stems primarily from the profound differences between non-covalent protein folding reactions and covalent reactions of small molecules. In the simplest reactions of small molecules, except for the one or two bonds that are being reconfigured, the rest of the reactant-structure, to a first approximation, usually remains fairly intact as the reaction proceeds (this need not necessarily hold for all simple chemical reactions and probably not for complex reactions). Consequently, if we were to use the bond-length of the bond that is being reconfigured as the RC, and find that the difference in Gibbs energy between any two reaction-states that occur consecutively along the RC are very similar, a reasonable assumption/expectation would be that their structures must be very similar.^69-77^ However, such an assumption cannot be valid for protein folding since an incredibly large number of chain and solvent configurations can lead to conformers having exactly the same Gibbs energy. Consequently, it is difficult to imagine how one can infer the structure of the transiently populated protein reaction-states, including the TSEs, to a near-atomic resolution purely from energetics (see Φ-value analysis later).^78-80^

### The position of the TSE along the entropic RC

The Leffler parameters for the relative sensitivities of the activation and equilibrium Gibbs energies in response to a perturbation in temperature are given by the ratios of the derivatives of the activation and equilibrium Gibbs energies with respect to temperature.^13-15,81^ Thus, for the partial folding reaction *D* ⇌ [*TS*], we have

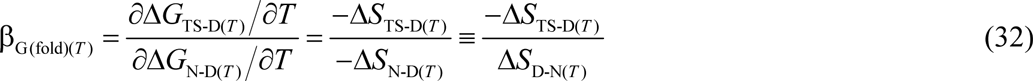

where β_G(fold)(*T*)_ is classically interpreted to be a measure of the position of the TSE relative to the DSE along the entropic RC.^49^ Recasting Eq. (32) in terms of (8) and (A4) and rearranging gives

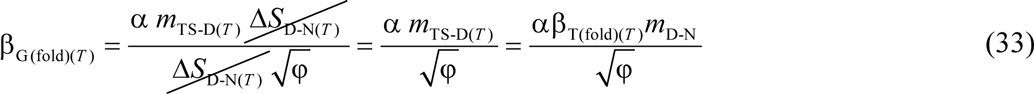

Similarly for the partial unfolding reaction *N* ⇌ [*TS*] we have

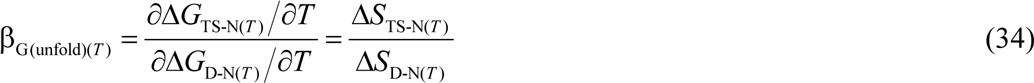

Where β_G(unfold)(*T*)_ is a measure of the position of the TSE relative to the NSE along the entropic RC. Substituting Eqs. (9) and (A6) in (34) gives

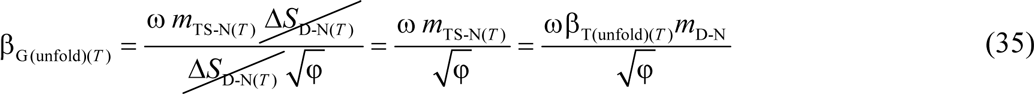

Inspection of Eqs. (32) and (34) shows that β_G(fold)(*T*)_ + β_G(unfold)(*T*)_ = 1 for any given reaction-direction. Now, since Δ*S*_D-N(*T*)_ = Δ*S*_TS-D(*T*)_ = Δ*S*_TS-N(*T*)_ = 0 at *T_S_*, β_G(fold)(*T*)_ and β_G(unfold)(*T*)_ will be undefined for *T* = *T_S_*. However, these are removable discontinuities as is apparent from Eqs. (33) and (35); consequently, curves simulated using the latter set of equations will have a hole at *T_S_*. If we ignore the hole at *T_S_* to enable a physical description and their comparison to other RCs, the extremum of β_G(fold)(*T*)_ (which is positive and a minimum) and the extremum of β_G(unfold)(*T*)_ (which is positive and a maximum) will occur at *T_S_* (**Figure 17** and **Figure 17−figure supplement 1**) and is a consequence of *m*_TS-D(*T*)_ being a minimum, and both *m*_TS-N(*T*)_ and φ being a maximum, respectively, at *T_S_*. This can also be demonstrated by differentiating Eqs. (32) and (34) with respect to temperature (not shown). Comparison of Eqs. (28) and (33), and Eqs. (29) and (35) demonstrate that when *T* = *T_S_*, we have β_H(fold)(*T*)_ = β_G(fold)(*T*)_ and β_H(unfold)(*T*)_ = β_G(unfold)(*T*)_, i.e., the position of the TSE along the heat capacity and entropic RCs are identical at *T_S_*, and non-identical for *T* ≠ *T_S_* (**Figure 17**). Further, since *m*_TS-N(*T*)_ = β_T(unfold)(*T*)_ = 0 at *T*_*S*(α)_ and *T*_*S*(ω)_ (**Figure 2B** and **Figure 2−figure supplement 1B**), β_G(unfold)(*T*)_ ≡ β_T(unfold)(*T*)_ = 0 and β_G(fold)(*T*)_ ≡ β_T(fold)(*T*)_ = 1, and not identical for *T* ≠ *T*_*S*(α)_ and *T*_*S*(ω)_; and for *T*_α_ ≤ *T* < *T*_*S*(α)_ and *T*_*S*(ω)_ < *T* ≤ *T*_ω_ (the ultralow and high temperature *Marcus-inverted-regimes*, respectively), β_G(fold)(*T*)_ and β_T(fold)(*T*)_ are greater than unity, and β_G(unfold)(*T*)_ and β_T(unfold)(*T*)_ are negative (**Figure 18**). Note that although β_G(fold)(*T*)_ is unity at *T*_*S*(α)_ and *T*_*S*(ω)_, the structures of the TSE and the NSE cannot be assumed to be identical as explained earlier.

**Figure 17.**
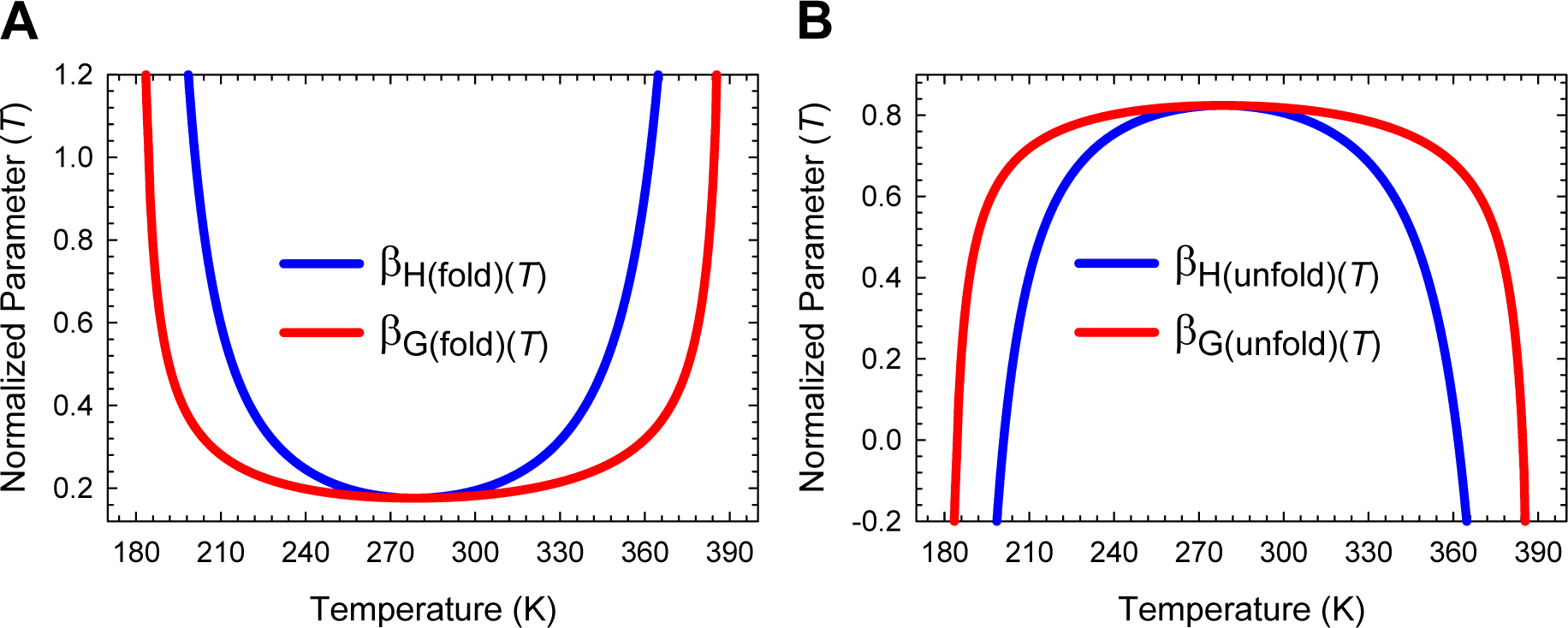
Comparison of the position of the TSE along the heat capacity and the entropic RCs. **(A)** The position of the TSE with respect to the DSE along the heat capacity and the entropic RCs are identical at *T_S_*, and non-identical for *T* ≠ *T_S_*. **(B)** The position of the TSE with respect to the NSE along the heat capacity and the entropic RCs are identical at *T_S_*, and non-identical for *T* ≠ *T_S_*. See **Figure 17−figure supplement 1** for unscaled plots of β_G(fold)(*T*)_ and β_G(unfold)(*T*)_.

**Figure 18.**
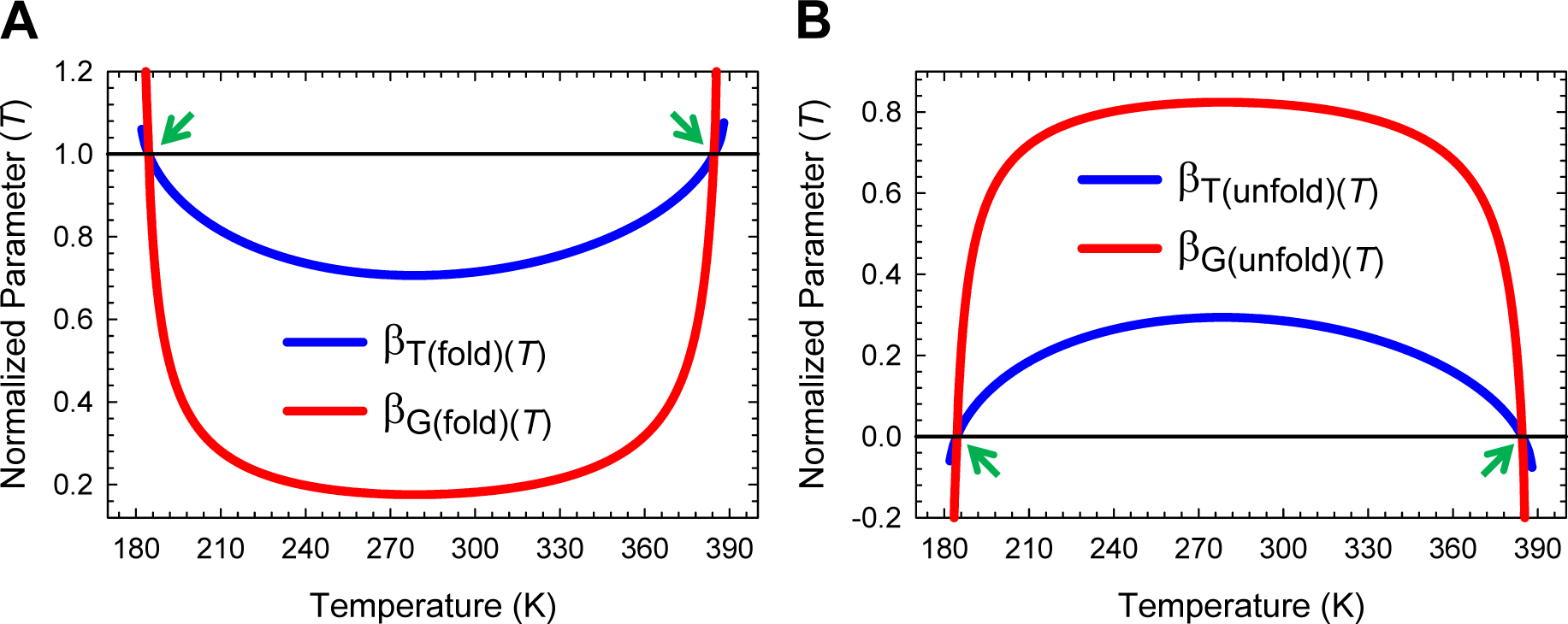
Comparison of the position of the TSE along the heat capacity and the entropic RCs. **(A)** The position of the TSE with respect to the DSE along the SASA and the entropic RCs are identical at *T*_*S*(α)_ and *T*_*S*(ω)_, and dissimilar for *T* ≠ *T*_*S*(α)_ and *T*_*S*(ω)_. **(B)** The position of the TSE with respect to the NSE along the SASA and the entropic RCs are identical at *T*_*S*(α)_ and *T*_*S*(ω)_, and dissimilar for *T* ≠ *T*_*S*(α)_ and *T*_*S*(ω)_. The green pointers indicate *T*_*S*(α)_ and *T*_*S*(ω)_.

Although it is beyond the scope of this manuscript to perform a large-scale survey of literature for corroborating evidence, the notion that these equations must hold for any two-state folder (as long as they conform to the postulates laid out in Paper-I) is readily apparent from the experimental data of Kelly, Gruebele and colleagues.^25,82-84^ However, the reader will note that what Gruebele and coworkers refer to as Φ_T_(*T*, *P*) (see Eq. (8) in Crane et al., 2000 and Jäger et al., 2001, Eq. (5) in Ervin and Gruebele, 2002, and Eq. (3) in Nguyen et al., 2003) is equivalent to β_G(*T*)_ in this article. We will reserve the letter Φ for Φ-value analysis which we will address later.^79^ Inspection of Fig. 7a in Crane et al., 2000 demonstrates that β_G(fold)(*T*)_ increases with temperature for *T* > *T_S_* for both the wild type hYAP WW domain and its mutant W39F (∽0.4 at 38 °C and ∽0.8 at 78 °C). This pattern is once again repeated for the wild type and several mutants of Pin WW domain (Fig. 8 in Jäger et al., 2001) and more importantly for ΔNΔC Y11R W30F, a variant of FBP28 WW (inset in Fig. 4B in Nguyen et al., 2003). Nevertheless, all is not in agreement since the shapes of their β_G(fold)(*T*)_ curves are distinctly different from what is expected from the formalism discussed in this article. This discrepancy most probably has to do with their use of Taylor expansion with three adjustable parameters to calculate the temperature-dependence of equilibrium stability and the Gibbs activation energies. While it is stated that the use of this non-classical model and the associated adjustable parameters in preference to the physically realistic Schellman formalism (which requires the model-independent calorimetrically determined value of Δ*C*_*p*DN_)^6^ makes little or no difference to the temperature-dependence of equilibrium stability over an extended temperature range, this may not be true for the activation energy. Once again in good agreement with prediction that β_G(unfold)(*T*)_ must decrease with temperature for *T* > *T_S_*, Tokmakoff and coworkers find that β_G(unfold)(*T*)_ for ubiquitin decreases with temperature (0.77 at 53 °C and 0.67 at 67 °C).^85^ Note that although raw data of the said groups and their conclusion that the position of the TSE shifts closer to the NSE as the temperature is raised for *T* > *T_S_* is in agreement with the predictions of the equations derived here, their Hammond-postulate-based inference of the structure of the TSE is flawed from the perspective of the parabolic approximation.

Now, at the midpoint of thermal (*T_m_*) or cold denaturation (*T_c_*), Δ*G*_D-N(*T*)_ = 0; therefore, Eqs. (1) and (2) become

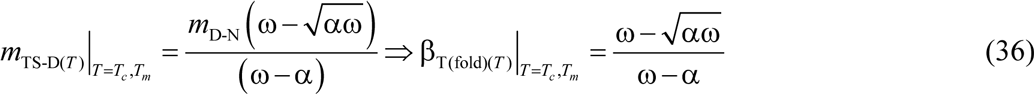

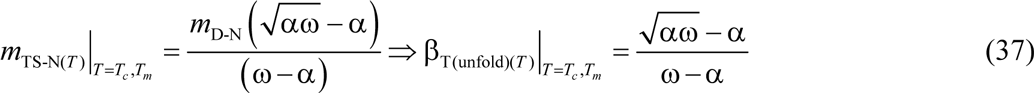

Substituting Eqs. (36) and (37), and 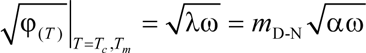 in (33) and (35), respectively, and simplifying gives

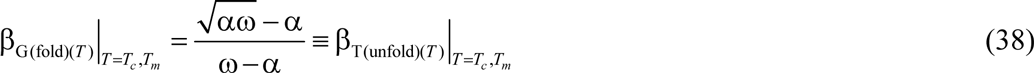

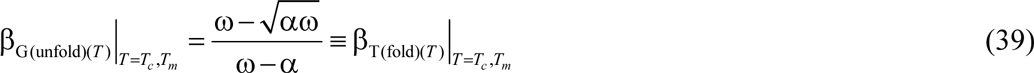

Simply put, at the midpoint of cold or heat denaturation, the position of the TSE relative to the DSE along the entropic RC is identical to the position of the TSE relative to the NSE along the SASA-RC (**Figure 19A**). Similarly, the position of the TSE relative to the NSE along the entropic RC is identical to the position of the TSE relative to the DSE along the SASA-RC (**Figure 19B**). Dividing Eq. (38) by (39) gives

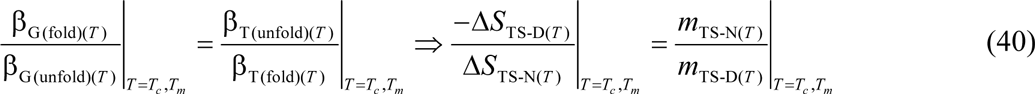

This seemingly obvious relationship has far deeper physical meaning. Simplifying further and recasting gives

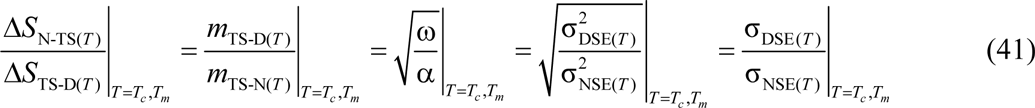

**Figure 19.**
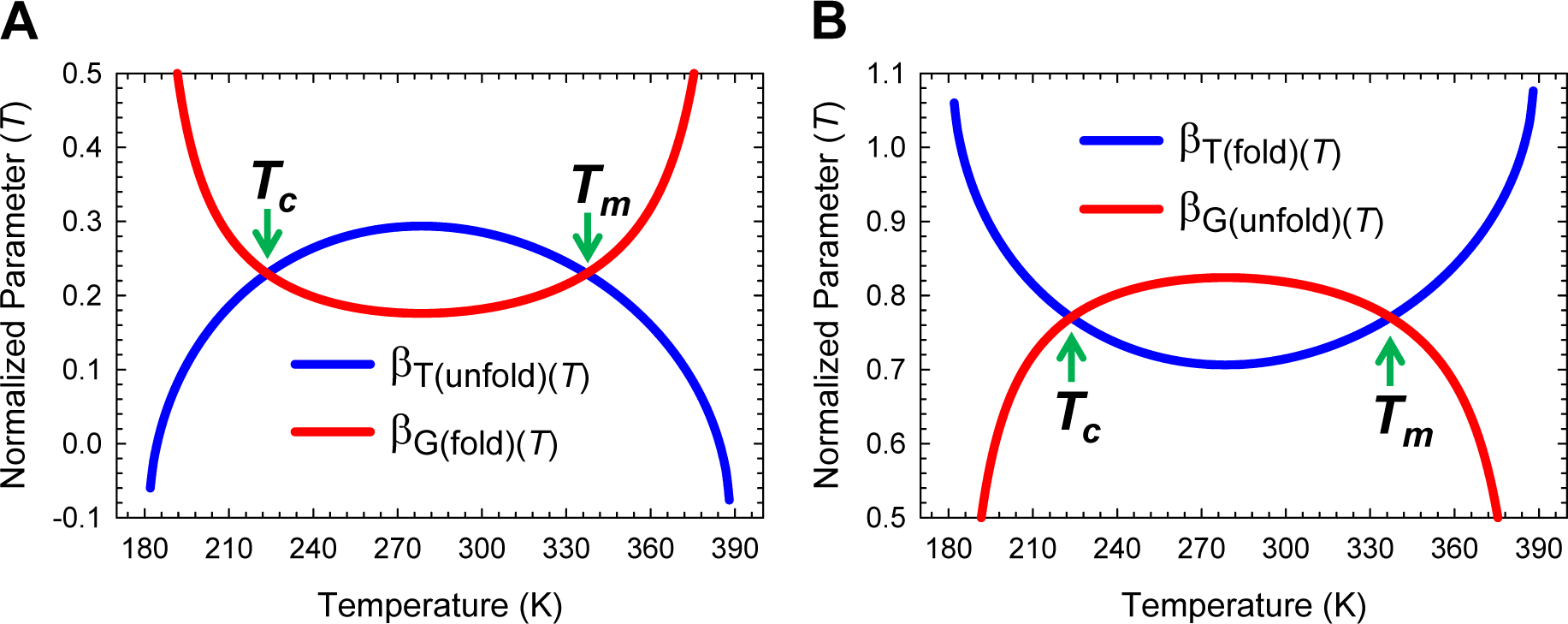
The intersection of β_G(*T*)_ and β_T(*T*)_ functions. **(A)** At the midpoint of cold or heat denaturation, the position of the TSE relative to the DSE along the entropic RC is identical to the position of the TSE relative to the NSE along the SASA-RC. **(B)** The position of the TSE relative to the NSE along the entropic RC is identical to the position of the TSE relative to the DSE along the SASA-RC at the midpoint of cold or heat denaturation.

Thus, at the temperatures *T_c_* and *T_m_* where the concentration of the DSE and the NSE are identical, the ratio of the slopes of the folding and unfolding arms of the chevron determined at the said temperatures are a measure of the ratio of the change in entropies for the partial folding reactions [*TS*] ⇋ *N* and *D* ⇋[*TS*], or the square root of the ratio of the Gaussian variances of the DSE 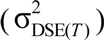 and the NSE 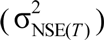 along the SASA-RC, or equivalently, the ratio of the standard deviations of the DSE σ_DES(*T*)_ and the NSE σ_NES(*T*)_ Gaussians **(Figure 19−figure supplement 1**; see Paper-I for the relationship between force constants, Gaussian variances and equilibrium stability). A corollary is that irrespective of the primary sequence, or the topology of the native state, or the residual structure in the DSE, if for a spontaneously folding two-state system at constant pressure and solvent conditions it is found that at a certain temperature the ratio of the distances by which the denatured and the native conformers must travel from the mean of their ensemble to reach the TSE along the SASA RC is identical to the ratio of the standard deviations of the Gaussian distribution of the SASA of the conformers in the DSE and the NSE, then at this temperature the Gibbs energy of unfolding or folding must be zero.

As an aside, the reader will note that β_G(fold)(*T*)_ and β_G(unfold)(*T*)_ are equivalent to the Brønsted exponents alpha and beta, respectively, in physical organic chemistry; and their classical interpretation is that they are a measure of the structural similarity of the transition state to either the reactants or the products.^81^ If the introduction of a systematic perturbation (often a change in structure *via* addition or removal of a substituent, pH, solvent etc.) generates a reaction-series, and if for this reaction-series it is found that alpha is close to zero (or beta close to unity), then it implies that the energetics of the transition state is perturbed to the same extent as that of the reactant, and hence inferred that the structure of the transition state is very similar to that of the reactant. Conversely, if alpha is close to unity (or beta is almost zero), it implies that the energetics of the transition state is perturbed to the same extent as the product, and hence inferred that the transition state is structurally similar to the product. Although the Brønsted exponents in many cases can be invariant with the degree of perturbation (i.e., a constant slope leading to linear free energy relationships),^70,86^ this is not necessarily true, especially if the degree of perturbation is substantial (Fig. 3 in Cohen and Marcus, 1968; Fig. 1 in Kresge, 1975).^y14,72,81^ Further, this seemingly straightforward and logical Hammond-postulate-based conversion of Brønsted exponents to similarity or dissimilarity of the structure of the transition states to either of the ground states nevertheless fails for those systems with Brønsted exponents greater than unity and less than zero (see page 1897 in Kresge, 1974).^24,81,87-91^

To summarise, a comparison of the position of the TSE along the solvent (β_T(*T*)_), heat capacity (β_H(*T*)_), and entropic (β_G(*T*)_) RCs leads to three important general conclusions (**Figure 20**): (*i*) as long as ΔSASA_D-N_ is large, and by extension Δ*C*_*p*D-N_ is large and positive, the position of the TSE relative to the ground states along the various RCs is neither constant nor a simple linear function of temperature when investigated over a large temperature range; (*ii*) for a given temperature, the position of the TSE along the RC depends on the choice of the RC; and (*iii*) although the algebraic sum of β_T(fold)(*T*)_ and β_T(unfold)(*T*)_, β_H(fold)(*T*)_ and β_H(unfold)(*T*)_, and β_G(fold)(*T*)_ and β_G(unfold)(*T*)_ must be unity for a two-state system for any particular temperature, individually they can be positive, negative, or zero. Consequently, the notion that the atomic structure of the transiently populated reaction-states in protein folding can be inferred from their position along the said RCs is flawed.^78^

**Figure 20.**
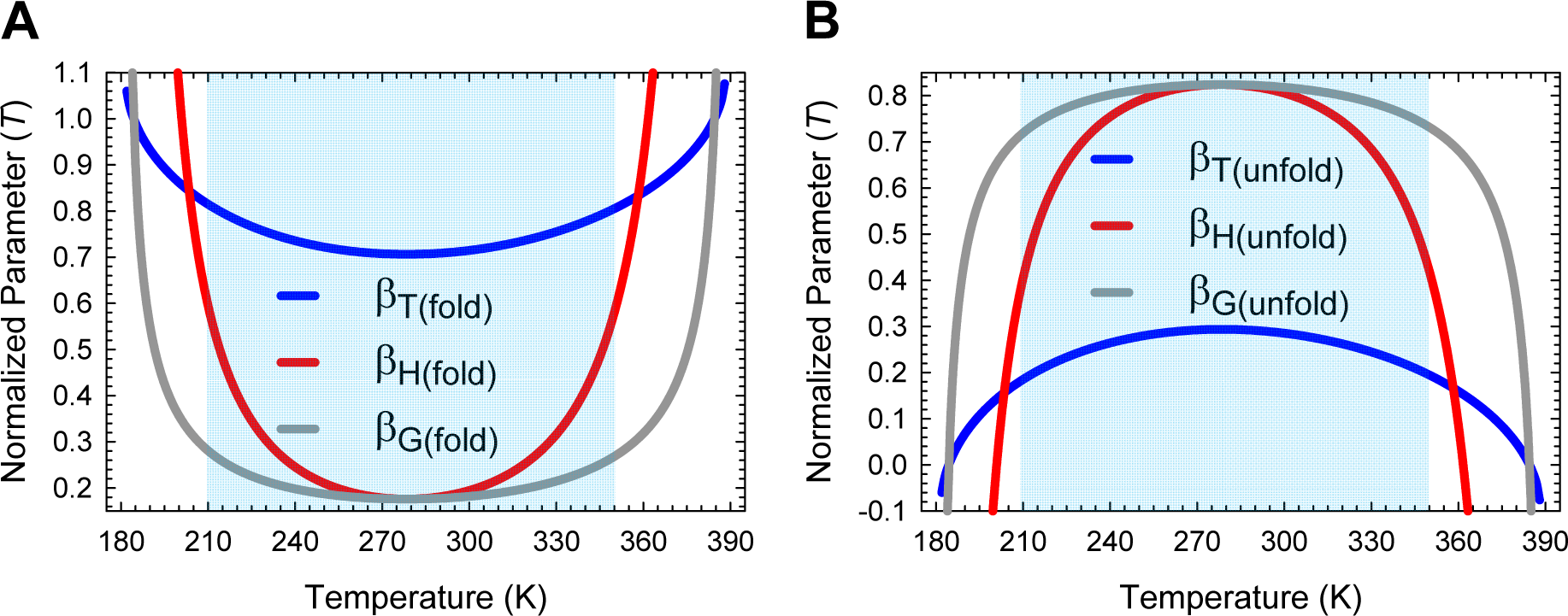
Comparison of the position of the TSE along the various reaction coordinates. The position of the TSE relative to the ground states depends on the choice of the RC and changes in a complex manner with temperature. **(A)** For 210 K∽ < *T* < ∽350 K (shaded region), the position of the TSE relative to the DSE is the most advanced along the solvent RC as compared to the heat capacity and entropic RCs; and for *T* ≠ *T_S_*, is the most advanced along the heat capacity RC as compared to the entropic RC **(B)** In contrast, for 210 K∽ < *T* < ∽350 K and *T* ≠ *T_S_*, the position of the TSE relative to the NSE is the most advanced along the entropic RC as compared to the heat capacity RC, and is the least advanced along the SASA-RC.

### Temperature-dependence of Φ-values

Φ-value analysis is a variation of the Brønsted procedure introduced by Fersht and coworkers which when properly implemented claims to provide a near-atomic-level description of the transiently populated reaction-states in protein folding.^79,80^ In this procedure, the primary sequence of the target protein is modified using protein engineering, and the effect of these perturbations are quantified through a parameter Φ (0 ≤ Φ ≤ 1) which by definition is the ratio of mutation-induced change in the Gibbs activation energy of folding/unfolding to the corresponding change in equilibrium stability. According to the canonical formulation, when Φ_F(*T*)_ = 0 (Φ-value for folding), it implies that the energetics of the TSE is perturbed to the same extent as that of the DSE upon mutation, and hence *inferred* that the said reaction-states are structurally identical with respect to the site of mutation. In contrast, when Φ_F(*T*)_ = 1, it implies that the energetics of the TSE is perturbed to the same extent as that of the NSE, and hence *inferred* that the structure at the site of mutation is identical in both the TSE and the NSE. Partial Φ-values are difficult to interpret and are thought to be due to partially developed interactions in the TSE, or multiple routes to the TSE. Thus, while Φ *per se* is the slope a two-point Brønsted plot, the conversion of this value to relative-structure is based on the Hammond postulate and the canonical range: The Hammond postulate provides the licence to infer structure from energetics, and the canonical scale enables one to infer how similar or dissimilar the TSE is to either the DSE or the NSE. Assuming that the prefactor is identical for the wild type and the mutant proteins, we may write for the partial folding (*D* ⇌ [*TS*]) and unfolding (*N* ⇌ [*TS*]) reactions

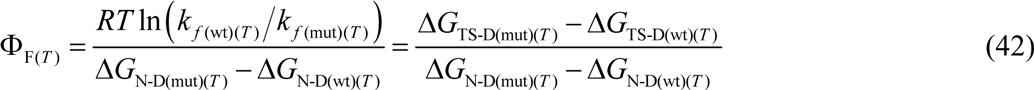

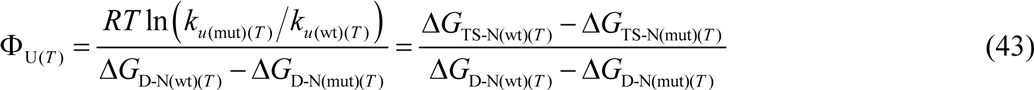

where the subscripts “wt” and “mut” denote the reference or the wild type, and the structurally perturbed protein, respectively, and Φ_U(*T*)_ is the Φ-value for unfolding. Inspection of Eqs. (42) and (43) shows that for a two-state system, Φ_F(*T*)_ + Φ_U(*T*)_ = 1. Now, although the primary sequence is intact in thermal denaturation experiments, we can readily calculate the temperature-dependence of Φ values for folding and unfolding using the protein at one unique temperature as the internal reference or the wild type, and protein at all the rest of the temperatures as the mutants. Thus, if the protein at *T_S_* is defined as the internal reference or the wild type, Eqs. (42) and (43) become

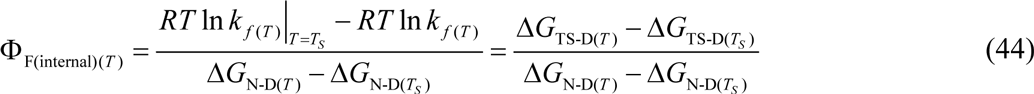

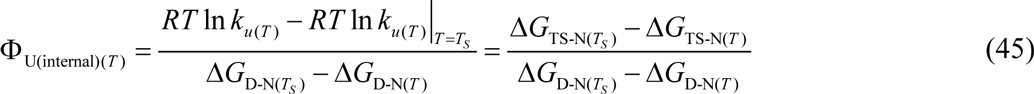

Similarly, if the protein at *T_m_* is defined as the internal reference or the wild type, Eqs. (42) and (43) become

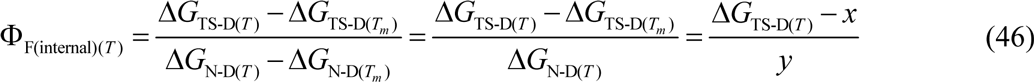

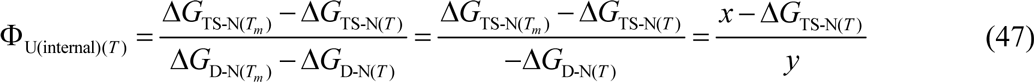

Where *x* = Δ*G*_TS-D(*T_m_*)_ = Δ*G*_TS-N(*T_m_*)_ and *y* = Δ*G*_N-D(*T*)_ ≡ −Δ*G*_D-N(*T*)_ (the denominator reduces to a single quantity since Δ*G*_D-N(*T_m_*)_ ≡ −Δ*G*_N-D(*T_m_*)_ = 0). The parameters Φ_F(internal)(*T*)_ and Φ_U(internal)(*T*)_ (which are obviously undefined for the reference temperatures) when interpreted according to the canonical Φ-value framework (i.e., the notion that 0 ≤ Φ ≤ 1) are a measure of the *global* similarity or dissimilarity of the structure of the TSE to either the DSE or the NSE. Thus, if Φ_F(internal)(*T*)_ = 0, it implies that the energetics of the TSE is perturbed to the same extent as that of the DSE upon a perturbation in temperature, and hence inferred that the global structure of the TSE is identical to that of the DSE. Conversely, if Φ_F(internal)(*T*)_ = 1, it implies that the energetics of the TSE is perturbed to the same extent as the NSE upon a perturbation in temperature, and hence inferred that the global structure of the TSE is identical to that of the NSE.

Inspection of **Figures 21** and **Figure 21−figure supplements 1**, **2**, **3** and **4** immediately demonstrates that: (*i*) irrespective of which temperature is defined as the internal reference (i.e., the wild type), Φ_F(internal)(*T*)_ must be a minimum and Φ_U(internal)(*T*)_ must be a maximum at *T_S_* (see **Appendix)**; (*ii*) the magnitude of Φ_F(internal)(*T*)_ is always the least, and the magnitude of Φ_U(internal)(*T*)_ is always the greatest when the protein at *T_S_* is defined as the reference or the wild type protein, and any deviation in the definition of the reference temperature from *T_S_* must lead to a uniform increase in Φ_F(internal)(*T*)_ and a uniform decrease in Φ_U(internal)(*T*)_ for all temperatures; (*iii*) although the algebraic sum of Φ_F(internal)(*T*)_ and Φ_U(internal)(*T*)_ is unity for all temperatures, the notion that they must independently be restricted to 0 ≤ Φ ≤ 1 is flawed; and (*iv*) although both Leffler β_G(*T*)_ and Fersht Φ values are derived from changes in Gibbs activation energies for folding and unfolding relative to changes in equilibrium stability upon a perturbation in temperature, their response is not the same since the equations that govern their behaviour are not the same. While the magnitude of the Leffler β_G(*T*)_ is independent of the reference owing to it being the ratio of the derivatives of the change in Gibbs energies with respect to temperature, the magnitude of Φ(internal)(*T*) is dependent on the definition of the reference state. For example, if the protein at *T_S_* is defined as the wild type, then β_G(fold)(*T*)_ ≈ Φ_F(internal)(*T*)_ and β_G(unfold)(*T*)_ ≈ Φ_U(internal)(*T*)_ around the temperature of maximum stability; but as the temperature deviates from *T_S_*, β_G(fold)(*T*)_ increases far more steeply than Φ_F(internal)(*T*)_, and β_G(unfold)(*T*)_ decreases far more steeply than Φ_U(internal)(*T*)_ such that for *T* ≠ *T_S_* we have β_G(fold)(*T*)_ > Φ_F(internal)(*T*)_ and β_G(unfold)(*T*)_ < Φ_U(internal)(*T*)_ (**Figure 21−figure supplement 3**). In contrast, if the protein at *T_m_* is defined as the wild type, then we have: (*i*) β_G(fold)(*T*)_ < Φ_F(internal)(*T*)_ for *T_c_* < *T* < *T_m_* and β_G(fold)(*T*)_ > Φ_F(internal)(*T*)_ for *T* < *T_c_* and *T* > *T_m_*; and (*ii*) β_G(unfold)(*T*)_ > Φ_U(internal)(*T*)_ for *T_c_* < *T* < *T_m_* and β_G(unfold)(*T*)_ < Φ_U(internal)(*T*)_ for *T* < *T_c_* and *T* > *T_m_*(**Figure 21−figure supplement 4**). The point we are trying to make is that a comparison of the position of the TSE along Leffler β_G(*T*)_ and Φ_(internal)(*T*)_ RCs is not straightforward since both β_G(*T*)_ and Φ_(internal)(*T*)_ are temperature-dependent, and importantly respond differently to temperature-perturbation; and even if we restrict the comparison to one particular temperature, the answer we get is still subjective since the magnitude of Φ_(internal)(*T*)_ is dependent on how we define the wild type.^92^

Although the mathematical formalism for why the extrema of Φ_F(internal)(*T*)_ (which is a minimum) and Φ_U(internal)(*T*)_ (which is a maximum) must always occur precisely at *T_S_* has been shown in the appendix, it is instructive to examine the same graphically. Inspection of **Figure 21−figure supplements 5, 6** and **7** demonstrates that this is a consequence of Δ*G*_TS-D(*T*)_ and Δ*G*_N-D(*T*)_ being a minimum, and Δ*G*_TS-N(*T*)_ and Δ*G*_D-N(*T*)_ being a maximum at *T*_*S*_. Subtracting the reference Gibbs energies from the numerator and the denominator (Eq. (44)) has the effect of lowering the Δ*G*_TS-D(*T*)_ curve and raising the Δ*G*_N-D(*T*)_, such that the value of the said curves are zero at the reference temperature, but the shapes of the curves are not altered in any way (**Figure 21−figure supplement 5**). On the other hand, for Δ*G*_TS-N(*T*)_ and Δ*G*_D-N(*T*)_ curves (Eq. (45)), apart from the value of the curves becoming zero at the reference, it causes them to flip vertically (**Figure 21−figure supplement 6**). Consequently, if we divide the transformed Gibbs activation energies by the transformed equilibrium Gibbs energies, we end up with Φ_F(internal)(*T*)_ and Φ_U(internal)(*T*)_ which are a minimum and a maximum, respectively, at *T_S_* (**Figure 21−figure supplement 7**).

Now that the process that leads to the temperature-dependence of Φ has been addressed, the question is “Can we infer the structure of the TSE as being similar to either the DSE or the NSE from these data?” The answer is “no” for several reasons. First, as argued earlier, the Hammond postulate cannot be valid for protein folding; and because the structural interpretation of Φ values is based on the Hammond postulate, it too must be deemed fallacious. Second, even if we accept the premise that Hammond postulate is applicable to protein folding, the inference that the global structure of the TSE as being denatured-like for Φ_F(internal)(*T*)_ = 0, and native-like for Φ_F(internal)(*T*)_ = 1 is flawed since Φ values need not necessarily be restricted to 0 ≤ Φ ≤ 1 (**Figure 21−figure supplement 2**). Third, even if we summarily exclude those wild types that lead to anomalous Φ values as being unsuitable for Φ analysis, we still have a problem since even within the restricted set of wild types that yield 0 ≤ Φ ≤ 1, their magnitude depends on the definition of the wild type; consequently, for the same temperature, the degree of structure in the TSE relative to that in the DSE appears to increase as the definition of the wild type deviates from *T_S_* (**Figure 21−figure supplement 1**). If we try to circumvent this interpretational problem by arguing that the “inference of the structure of the TSE” is always relative to the residual structure in the DSE, and that changing the definition of what constitutes the wild type will invariably affect Φ values, then we can’t really say much about the structure of the TSE without first solving the structure of the DSE. Fourth, even if through a judicious combination of various structural and biophysical methods (residual dipolar couplings, paramagnetic relaxation enhancement, small angle X-ray scattering, single molecule spectroscopy etc.), and computer simulation, we are able to determine the residual structure in the DSE,^93-96^ the structural interpretation of Φ values leads to physically unrealistic scenarios. For example, inspection of **Figure 21A** shows that around room temperature (298 K) Φ_F(internal)(*T*)_ ≈ 0.18. A canonical interpretation of this number implies that the global structure of the TSE is very similar to that of the DSE. However, inspection of **Figure 2−figure supplement 1A** shows that the denatured conformer has buried ∽70% of the total SASA to reach the TSE (i.e., advanced by about 70% along the SASA-RC). Similarly, inspection of **Figure 5A** shows that Δ*G*_TS-D(*T*)_ = 2.6 kcal.mol^-1^ at 298 K (note that this is not a small number that can be ignored since Δ*G*_D-N(*T*)_ = 2.1 kcal.mol^-1^ at 298 K). Further, we have shown earlier in the section on the “Inapplicability of the Hammond postulate to protein folding,” that even when two reaction-states have identical SASA, Gibbs energies, enthalpies, and entropies, there need not necessarily have identical structure. Thus, the question is: How can we conclude with any measure of certainty that the global structure of the TSE is very similar to that of the DSE at 298 K when they have such a large difference in SASA, and a substantial difference in Gibbs energy? To illustrate why it is difficult to rationalize the theoretical basis of Φ analysis, it is instructive to directly examine the ratio of the Gibbs activation energies and the difference in Gibbs energy between the ground states (**Figure 21−figure supplement 8**). It is immediately apparent that the ratios are a complex function of temperature; and although we can readily provide an explanation for the particular features of these complex dependences, it is difficult to see how subtracting reference energies from the numerator and denominator of the ratios Δ*G*_TS-D(*T*)_/Δ*G*_N-D(*T*)_ and Δ*G*_TS-N(*T*)_/Δ*G*_D-N(*T*)_ allows us to divine the structure of the TSE to a near-atomic resolution. This is once again readily apparent from the complex non-linear relationship between equilibrium stability and the rate constants (**Figure 21−figure supplement 9**).

**Figure 21.**
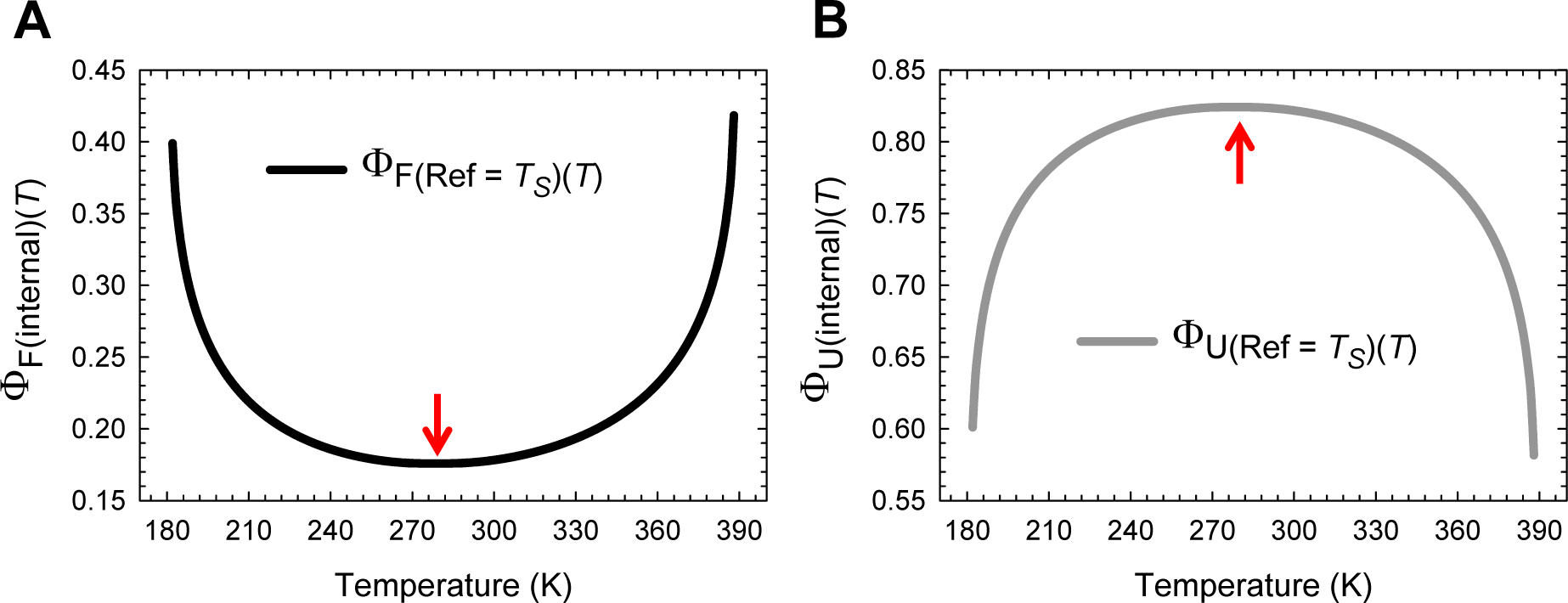
Temperature-dependence of Φ values when the protein at *T_S_* is defined as the wild type. **(A)** Temperature-dependence of Φ_F(internal)(*T*)_. **(B)** Temperature-dependence of Φ_U(internal)(*T*)_. The red pointers indicate the extrema of the functions. The discontinuities in the curves which must occur at *T_S_* have been removed by mathematically manipulating Eqs. (42) and (45) (manipulated equations not shown). Nevertheless, Φ_F(internal)(*T*)_ and Φ_U(internal)(*T*)_ are undefined at *T_S_* (i.e., the curves have holes at *T_S_* which is not obvious). Note that the mathematical stipulation that Φ_F(internal)(*T*)_ + Φ_U(internal)(*T*)_ = 1 for a two-state system is satisfied for all temperatures.

To further illuminate the difficulty in rationalizing the Φ-value procedure, it is instructive to apply Eqs. (42) and (45) to treat enthalpies. Thus, for the partial folding (*D* ⇌ [*TS*]) and unfolding (*N* ⇌ [*TS*]) reactions we have

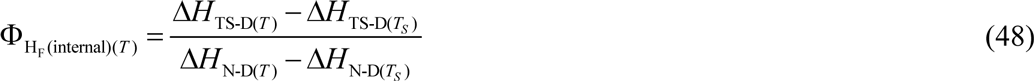

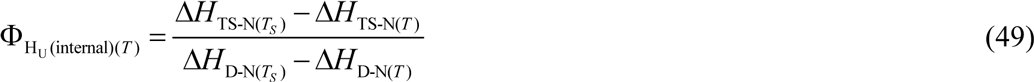

Where the parameters Φ_H_F_(internal)(*T*)_ and Φ_H_U_(internal)(*T*)_ are the “*enthalpic analogues*” of Φ_F(internal)(*T*)_ and Φ_U(internal)(*T*)_, respectively (the subscript “H” indicates we are using enthalpy instead of Gibbs energy), when the protein at the temperature *T_S_* is defined as the wild type. Now, if we apply an analogous version of the canonical interpretation given by Fersht and coworkers, it implies that when Φ_H_F_(internal)(*T*)_ = 0, the enthalpy of the TSE is perturbed to the same extent as that of the DSE upon a perturbation in temperature; and when Φ_H_F_(internal)(*T*)_ = 1, it implies that the enthalpy of the TSE is perturbed to the same extent as that of the NSE. It is easy to see that just as Φ_F(internal)(*T*)_ and Φ_U(internal)(*T*)_ are the *Fersht-analogues* of the Leffler β_G(fold)(*T*)_ and β_G(unfold)(*T*)_, respectively (see entropic RC), the parameters Φ_H_F_(internal)(*T*)_ and Φ_H_U_(internal)(*T*)_ are similarly the Fersht-analogues of the Leffler β_H(fold)(*T*)_ and β_H(unfold)(*T*)_, respectively (see heat capacity RC).

Inspection of **Figure 22** and its supplements immediately demonstrates that the same anomalies that prevent a straightforward structural interpretation of Φ_F(internal)(*T*)_ and Φ_U(internal)(*T*)_ are also emerge if we try to assign structure to their enthalpic analogues, Φ_H_F_(internal)(*T*)_ and Φ_H_U_(internal)(*T*)_. First, although the algebraic sum of Φ_H_F_(internal)(*T*)_ and Φ_H_U_(internal)(*T*)_ is unity for all temperatures, they need not independently be restricted to a canonical range of 0 ≤ Φ ≤ 1 (**Figure 22**). Second, the magnitude of Φ_HF(internal)(*T*)_ and Φ_H_U_(internal)(*T*)_ are dependent on the definition of the wild type (**Figure 22−figure supplement 1**). Third, changing the definition of the wild type has a dramatic effect on the relationship between the Leffler β_H(*T*)_ and its analogue, the Fersht Φ_H(internal)(*T*)_. Consequently, the question of whether Leffler β_H(*T*)_ underestimates or overestimates structure is dependent on how we analyse the system (**Figure 22−figure supplements 2** and **3**). Fourth, just as the temperature-dependent position of the TSE relative to the ground states depends on the choice of the RC (**Figure 20**), we see that Φ_(internal)(*T*)_ and its enthalpic analogue, Φ_H(internal)(*T*)_, change at different rates upon a perturbation in temperature (**Figure 22−figure supplement 4**). The difficulty in rationalizing how subtracting reference values from the numerator and the denominator of Eqs. (42) and (49) can yield residue-level information is once again apparent from the complex dependence of the ratios ∂ ln *k*_*f*(*T*)_/∂ ln *K*_N-D(*T*)_ = Δ*H*_TS-D(*T*)_/Δ*H*_N-D(*T*)_ and ∂ ln *k*_*u*(*T*)_/∂ ln *K*_D-N(*T*)_ = Δ*H*_TS-N(*T*)_/Δ*H*_D-N(*T*)_ on temperature (**Figure 22−figure supplement 5**).

**Figure 22.**
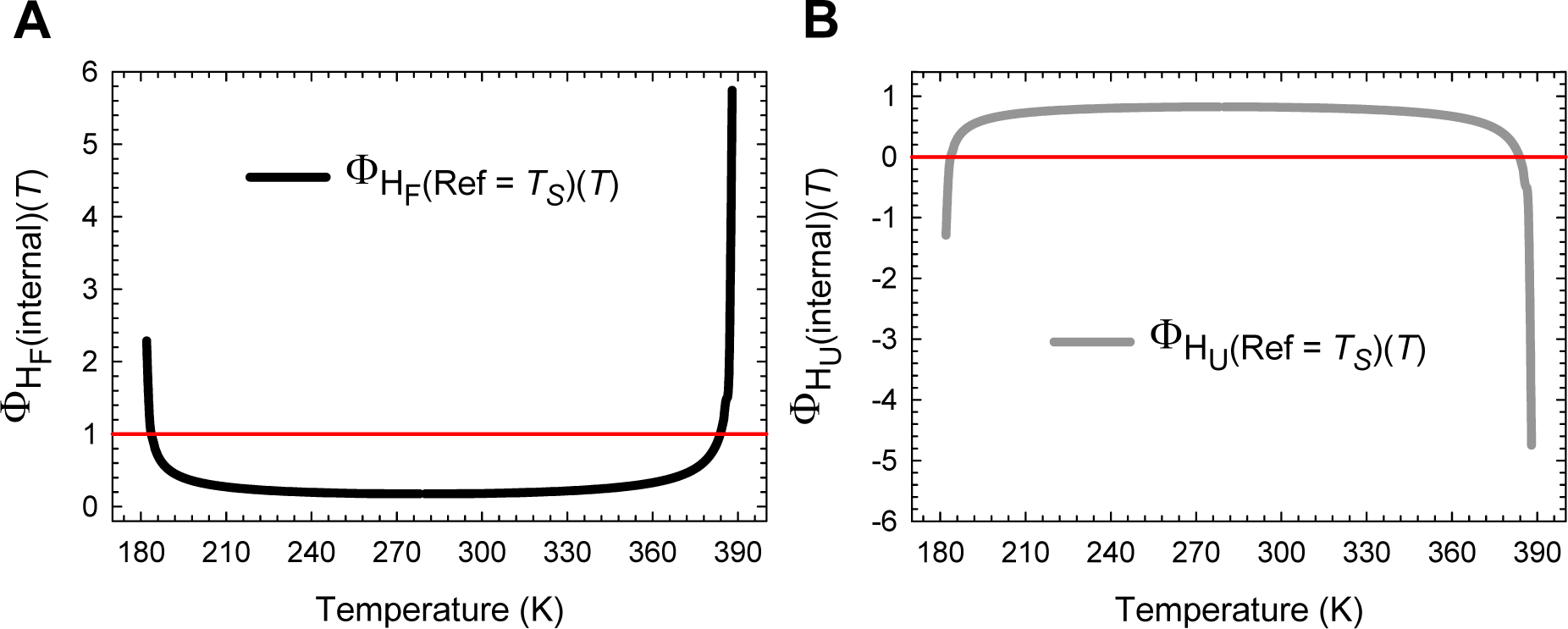
Temperature-dependence of Φ_H(internal)(*T*)_ when the protein at *T_S_* is defined as the wild type. **(A)** Temperature-dependence of Φ_H_F_(internal)(*T*)_. **(B)** Temperature-dependence of Φ_H_U_(internal)(*T*)_. Note that both these curves are undefined at *T_S_*. Although the algebraic sum of Φ_H_F_(internal)(*T*)_ and Φ_H_U_(internal)(*T*)_ is unity for all temperatures, they need not necessarily be are not restricted to a canonical range of 0 ≤ Φ ≤ 1. The parameters Φ_H_F_(internal)(*T*)_ and Φ_H_U_(internal)(*T*)_ are the “*enthalpic analogues*” of Φ_F_(internal)(*T*) and Φ_U_(internal)(*T*), respectively (the subscript “H” indicates we are using enthalpy instead of Gibbs energy). Consequently, this figure is the enthalpic equivalent of **Figure 21**.

### Comparison of theoretical and experimental Φ-values obtained from structural perturbation across 31 two-state systems

Given that the framework of Φ-value analysis was primarily developed to be used in conjunction with structural rather than temperature perturbation, and despite its anomalies has been used extensively for more than twenty years to divine the structures of the TSEs of not just globular but also membrane proteins, it is imperative to demonstrate that the notion that the structure of the TSE cannot be inferred from Φ-values is also valid for structural perturbation.^97-101^ Although a detailed reappraisal is beyond the scope of this article and will be presented elsewhere, because we have questioned the validity of Φ analysis, one is compelled to provide some justification in this article.

Consider the wild type of a hypothetical two-state folder whose equilibrium stability and the mean length of the RC at constant temperature, pressure and solvent conditions are given by Δ*G*_D-N(*T*)_ = 6 kcal.mol^-1^ and *m*_D-N_ = 2 kcal.mol^-1^.M^-1^, respectively. Although not necessarily true and addressed elsewhere, to limit the number of hypothetical scenarios to a manageable number, we will assume that the force constants of the DSE and the NSE-parabolas of the wild type and all its mutants are given by α = 1 M^2^.mol.kcal^-1^ and ω = 30 M^2^.mol.kcal^-1^. The effect of single point mutations on the wild type may be classified into a total of five unique scenarios (**Figure 23A**).

**Case I (Quadrant *x*2):** The introduced mutation causes a concomitant decrease in both the stability and the mean length of the RC (i.e., Δ*G*_D-N(*T*)(wt)_ > Δ*G*_D-N(*T*)(mut)_ and *m*_D-N(wt)_ > *m*_D-N(mut)_). This is equivalent to the introduced mutation causing the separation between the vertices of the DSE and the NSE-parabolas along the abscissa and ordinate to decrease (**Figure 23−figure supplement 1A**).

**Case II (Quadrant *y*1):** The introduced mutation causes a decrease in stability but concomitantly causes an increase in the mean length of the RC (i.e., Δ*G*_D-N(*T*)(wt)_ > Δ*G*_D-N(*T*)(mut)_ and *m*_D-N(wt)_ < *m*_D-N(mut)_). This is equivalent to the mutation causing a decrease in the separation between the vertices of the DSE and the NSE-parabolas along the ordinate, but an increase along the abscissa (**Figure 23−figure supplement 1B**).

**Case III (Quadrant *x*1):** The introduced mutation leads to an increase in stability but concomitantly causes a decrease in the mean length of the RC (i.e., Δ*G*_D-N(*T*)(wt)_ < Δ*G*_D-N(*T*)(mut)_ and *m*_D-N(wt)_ > *m*_D-N(mut)_). This is equivalent to the mutation causing an increase in the separation between the vertices of the DSE and the NSE-parabolas along the ordinate, but a decrease along the abscissa (**Figure 23−figure supplement 1C**).

**Case IV (Quadrant *y*2):** The introduced mutation leads to a concomitant increase in both the stability and the mean length of the RC (i.e., Δ*G*_D-N(*T*)(wt)_ < Δ*G*_D-N(*T*)(mut)_ and *m*_D-N(wt)_ < *m*_D-N(mut)_). This is equivalent to the mutation causing an increase in the separation between the vertices of the DSE and the NSE-parabolas along the ordinate and the abscissa (**Figure 23−figure supplement 1D**).

**Case V:** The introduced mutation leads to a change in stability but has no effect on the mean length of the RC (*m*_D-N(wt)_ = *m*_D-N(mut)_). This is equivalent to the mutation causing an increase or a decrease in the separation between the vertices of the DSE and the NSE-parabolas along the ordinate, but the separation along the abscissa is invariant (**Figure 23−figure supplement 2**).

**Figure 23.**
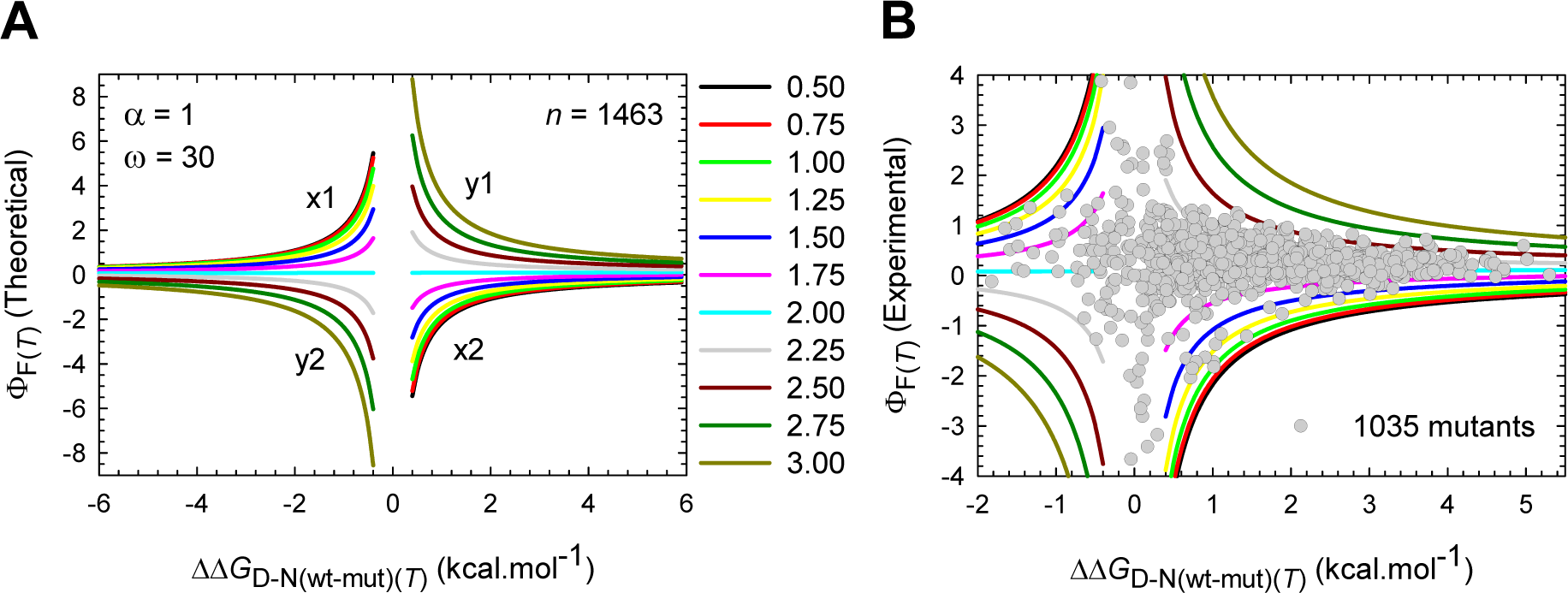
Comparison of the theoretical and experimental Φ_F(*T*)_ values (structural perturbation). **(A)** Theoretical limits of Φ_F(*T*)_ values according to parabolic approximation where all the 1463 theoretical mutants have the following common parameters: α = 1 M^2^.mol.kcal^-1^ (DSE-parabola); ω = 30 M^2^.mol.kcal^-1^ (NSE-parabola). The wild type was arbitrarily chosen to be the one with parameters Δ*G*_D-N(*T*)_= 6 kcal.mol^-1^ and *m*_D-N_ = 2 kcal.mol^-1^.M^-1^. The legend indicates the variation in *m*_D-N_ values in kcal.mol^-1^.M^-1^. The quadrants labelled *x*1 and *x*2 are for mutants whose *m*_D-N_ < *m*_D-N(wt)_ (i.e., a contraction of the RC) and the quadrants labelled *y*1 and *y*2 are for those mutants whose *m*_D-N_ > *m*_D-N(wt)_ (i.e., an expansion of the RC). Close inspection shows that for those mutants whose stabilities have changed but not their *m*_D-N_ values, the Φ_F(*T*)_ values are positive but very close to zero (shown in cyan). Theoretical Φ_F_ values corresponding to ΔΔ*G*_D-N(wt-mut)(*T*)_ = 0.0 ± 0.4 kcal.mol^-1^ have been excluded for clarity. This corresponds to about 6.7% error on the wild type Δ*G*_D-N(wt)(*T*)_. **(B)** An overlay of theoretical Φ_F(*T*)_ and experimental Φ_F(*T*)_ values in water for 1035 mutants from 31 two-state systems. Data used to calculate the experimental Φ_F(*T*)_ values were taken from published literature (detailed information is given elsewhere). The vertical asymptotes are a consequence of ΔΔ*G*_D-N(wt-mut)(*T*)_ approaching zero.

In summary, what we done is taken a pair of intersecting parabolas of differing curvature (ω > α), and systematically varied the separation between their vertices along the abscissa (*m*_D-N_) and ordinate (Δ*G*_D-N(*T*)_) without changing the curvature of the parabolas. Once this is done, we can calculate *a priori* the position of the *curve-crossings* relative to the vertex of the DSE-parabola along the abscissa (i.e., *m*_TS-D(*T*)_; Eq. (1)) and ordinate (i.e., Δ*G*_TS-D(*T*)_; Eq. (3)). Once the Δ*G*_TS-D(*T*)_ values for all combinations of Δ*G*_D-N(*T*)_ and *m*_D-N_ are obtained (each combination is equivalent to a point mutation), Φ_F(*T*)_ values can be readily calculated using Eq. (50) by arbitrarily choosing one particular combination of Δ*G*_D-N(*T*)_ (= 6 kcal.mol^-1^) and *m*_D-N_ (= 2 kcal.mol^-1^.M^-1^) as the wild type.

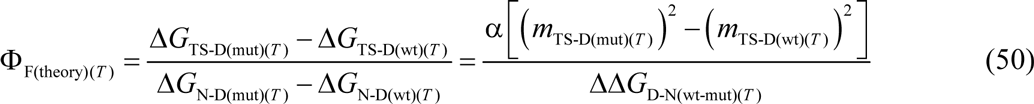

**Figure 23A** which has been generated by plotting the theoretical Φ_F(*T*)_ values as a function of ΔΔ*G*_D-N(wt-mut)(*T*)_ leads to two important conclusions: (*i*) Φ_F(*T*)_ values are not restricted to 0 ≤ Φ ≤ 1, and that the perceived unusualness of anomalous or non-classical Φ values is a consequence of flawed canonical limits; and (*ii*) the magnitude of Φ_F(*T*)_ values decrease as the difference in stability between the wild type and the mutant proteins increase, and at once debunks the idea that one must use an arbitrary ΔΔ*G*_D-N(wt-mut)(*T*)_ cut-off (± 0.6 kcal.mol^-1^ according to the Fersht lab, and ± 1.7 kcal.mol^-1^ according to Sanchez and Kiefhaber) for Φ_F(*T*)_ values to be interpretable.^98,102^ While it is true that Φ values would be error prone when |ΔΔ*G*_D-N(wt-mut)(*T*)_| is less than the error with which one can determine Δ*G*_D-N(*T*)_ of both the wild type and the mutant proteins (typically about ± 5-10% of Δ*G*_D-N(*T*)_),^103^ the increase in the magnitude of Φ_F(*T*)_ values when ΔΔ*G*_D-N(wt-mut)(*T*)_ approaches zero (the vertical asymptotes) is a mathematical certainty and not because of error as is commonly argued. Nevertheless, because these conclusions are based on the results of a model that is purely hypothetical, they would naturally be meaningless without experimental validation. Thus, as a test of the hypothesis, experimental Φ_F(*T*)_ values in water were calculated according to Eq. (51) using published kinetic data of a total of 1064 proteins (1035 mutants + 29 wild types) from 31 two-state systems (details of the systems analysed will be provided elsewhere).

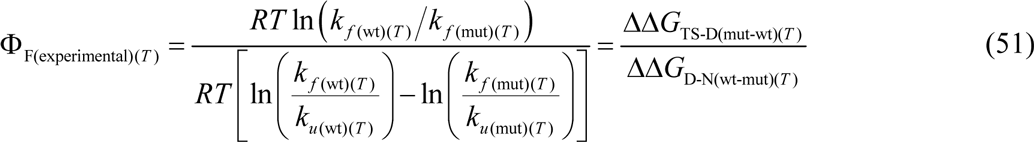

The remarkable agreement between theoretical prediction and experimental Φ_F(*T*)_ values is immediately apparent from an overlay of the said datasets (**Figure 23B**), and serves as arguably one of the most rigorous tests of the hypothesis for the following reasons: (1) The space enclosed by the curves in **Figure 23A** is complex and restricted. Therefore, if the experimental Φ_F(*T*)_ values fall within this restricted theoretical space it would be highly unlikely for it to be purely due to some dramatic coincidence. (2) The sample size of experimental dataset is sufficiently large (1035 mutations), and the two-state systems investigated include α, β, and α/β proteins (note that α and β refer to secondary structure in this context and not to the force constant of the DSE or the Tanford beta value, respectively), with size ranging from 37 to 107 residues. (3) The published kinetic data used to calculate experimental Φ_F(*T*)_ values were acquired by various labs under varying solvent conditions (buffers, co-solvents and pH; denaturant is either guanidine hydrochloride or urea) and temperature (as low as 278 K to as high as 301.16 K), over a period of about two decades using a variety of experimental methods, including infrared laser-induced and electrical discharge temperature-jump relaxation measurements, stopped flow and manual mixing experiments, and lineshape analysis of exchange-broadened NMR resonances. These results, including those on the temperature-dependence of Φ_F(*T*)_ values lead to an important conclusion: Because the canonical scale itself has no basis, Φ-value-based interpretation of the structure of the transiently populated protein reaction-states is dubious.

## Concluding Remarks

Although the temperature-dependent behaviour of FBP28 WW was analysed in great detail using the theory developed in the Papers I and II, and novel conclusions have been drawn, this is by no means sufficient since we have barely addressed the physical chemistry underlying the effect of temperature on the Gibbs energies, the enthalpies, the entropies, and the heat capacities of activation for folding and unfolding. These aspects will be dealt with in the accompanying articles. Further, there is a good reason why we have given little importance to the actual values of the reference temperatures and instead focussed on what they actually mean and how they relate to each other. Although the remarks in Table 1 are valid for all reference temperatures, except for the values of the equilibrium reference temperatures (*T_c_*, *T_H_*, *T_S_*, and *T_m_*), the values for the rest of them can change depending on the values of the force constants. However, what will not change is the inter-relationship between them. The nature of this limitation will be addressed when the mechanism of action of denaturants is investigated.

## Methods

The temperature-dependence of Δ*G*_D-N(*T*)_ of FBP28 WW wild type (**Figure 1**) was simulated according to Eq. (A1) using *T_m_* = 337.2 K, Δ*H*_D-N(*T_m_*)_ = 26.9 kcal.mol^-1^ and Δ*C*_*p*D-N_ = 417 cal.mol^-1^.K^-1^ (Table 1 in Petrovich et al., 2006).^4^ The values of *k*^0^ = 2180965 s^-1^, α = 7.594 M^2^.mol.kcal^-1^, ω = 85.595 M^2^.mol.kcal^-1^, and *m*_D-N_ = 0.82 kcal.mol^-1^.M^-1^ were extracted from the chevron of FBP28 WW (acquired at 283.16 K in 20 mM 3-[morpholino] propanesulfonic acid, ionic strength adjusted to150 mM with Na_2_SO_4_, pH 6.5) by fitting it to a modified chevron-equation using non-linear regression as described in Paper-I. The data required to simulate the chevron (*k*_*f*(H_2_O)(*T*)_, *k*_*u*(H_2_O)(*T*)_, *m*_TS-D(*T*)_ and *m*_TS-N(*T*)_) were taken from Table 4 in Petrovich et al., 2006.^4^ Once the parameters Δ*H*_D-N(*T_m_*)_, *T_m_*, Δ*C*_*p*D-N_, *m*_D-N_, the force constants α and ω, and *k*^0^ are known, the left-hand side of all the equations in this article may be readily calculated for any temperature. Note that the spring constants, *k*^0^, *m*_D-N_, and Δ*C*_*p*D-N_ are temperature-invariant.

## Appendix

### The temperature-dependence of Δ*G*_D-N(*T*)_, Δ*H*_D-N(*T*)_, and Δ*S*_D-N(*T*)_ functions

The temperature-dependence of the change in Gibbs energy, enthalpy and entropy of two-state systems upon unfolding at equilibrium are given by^6^

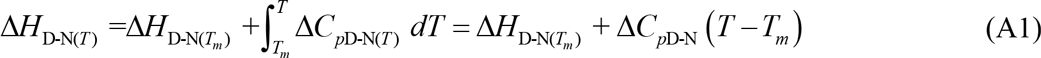

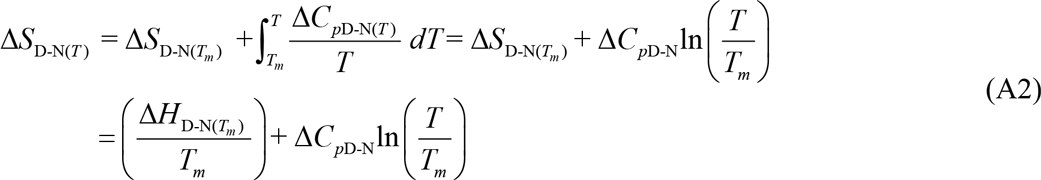

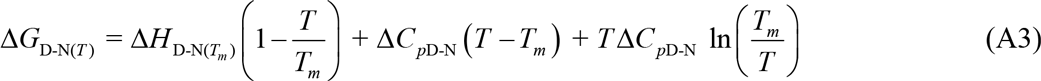

Where Δ*H*_D-N(*T*)_, Δ*H*_D-N(*T_m_*)_ and Δ*S*_D-N(*T*)_, Δ*S*_D-N(*T_m_*)_ denote the equilibrium enthalpies and entropies of unfolding, respectively, at any given temperature, and at the midpoint of thermal denaturation (*T_m_*), respectively, for a given two-state folder under defined solvent conditions. The temperature-invariant and the temperature-dependent difference in heat capacity between the DSE and NSE are denoted by Δ*C*_*p*D-N_ and Δ*C*_*p*D-N(*T*)_, respectively.

### The first derivatives of *m*_TS-D(*T*)_, *m*_TS-N(*T*)_, β_**T(fold)(*T*)**_ and β_**T(unfold)(*T*)**_ with respect to temperature

The first derivative of *m*_TS-D(*T*)_ is given by

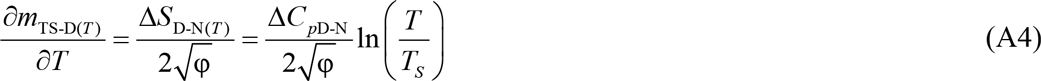

Because β_T(fold)(*T*)_ = *m*_TS-D(*T*)_/*m*_D-N_, we also have

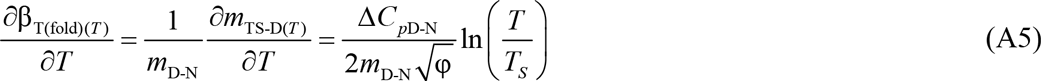

Since ∂*m*_TS-D(*T*)_/∂*T* and ∂β_T(fold)(*T*)_/∂*T* are physically undefined for φ < 0, their algebraic sign at any given temperature is determined by the ln(*T*/*T*_*S*_) term. This leads to three scenarios: (*i*) for *T* < *T_S_* we have ∂*m*_TS-D(*T*)_/∂*T* > 0 and ∂β_T(fold)(*T*)_/∂*T* > 0; and (*iii*) for *T* = *T*_*S*_ we have ∂*m*_TS-D(*T*)_/∂*T* = 0 and ∂β_T(fold)(*T*)_/∂*T* = 0.

Because *m*_TS-N(*T*)_ = (*m*_D-N_ − *m*_TS-D(*T*)_) for a two-state system, and β_T(unfold)(*T*)_ = *m*_TS-N(*T*)_/*m*_D-N_, we have

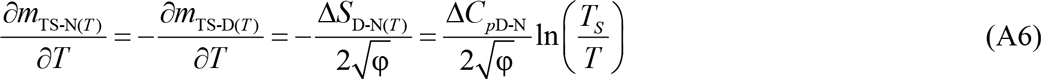

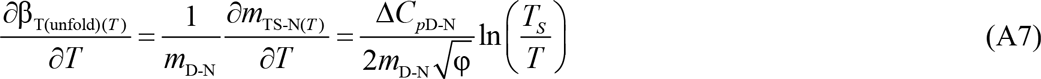

Eqs. (A6) and (A7) once again lead to three scenarios: (*i*) for *T* < *T_S_* we have ∂*m*_TS-N(*T*)_/∂*T* > 0 and ∂β_T(unfold)(*T*)_/∂*T* > 0; (*ii*) for *T* > *T*_*S*_ we have ∂*m*_TS-N(*T*)_/∂*T* < 0 and ∂β_T(unfold)(*T*)_/∂*T* > 0; and (*iii*) for *T* = *T_S_* we have ∂*m*_TS-N(*T*)_/∂*T* = 0 and ∂β_T(unfold)(*T*)_/∂*T* = 0.

### The second derivatives of *m*_TS-D(*T*)_ and *m*_TS-N(*T*)_ with respect to temperature

Differentiating Eq. (A4) with respect to temperature gives

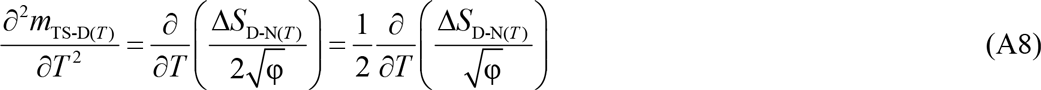

Simplifying Eq. (A8) yields

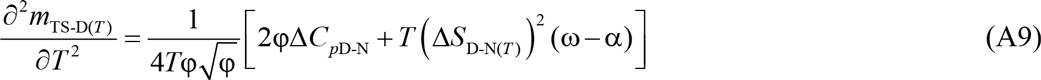

Similarly, we may show that

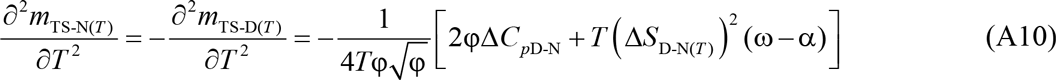

### Expression for the temperature-dependence of the observed rate constant

The observed rate constant *k*_obs(*T*)_ for a two-state system is the sum of *k*_*f*(*T*)_ and *k*_*u*(*T*)_.^104^ Therefore, we can write

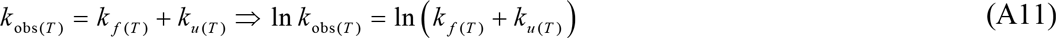

Substituting Eqs. (5) and (6) in (A11) gives

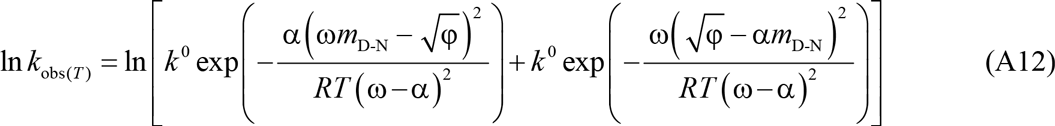

### Expressions to demonstrate why the extrema of Φ_F(internal)(*T*)_ and Φ_U(internal)(*T*)_ must occur at *T_S_*

Differentiating Eq. (44) with respect to temperature gives

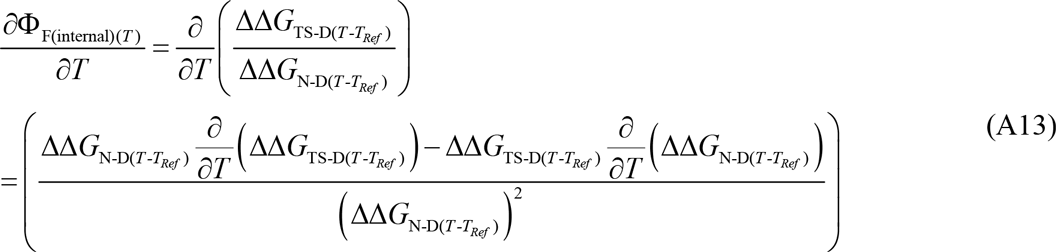

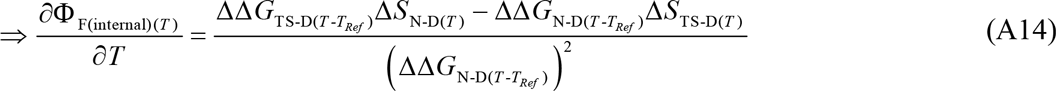

where the protein at the temperature *T_Ref_* is by definition the wild type protein. Because Δ*S*_N-D(*T*)_ and Δ*S*_TS-D(*T*)_ are both zero at *T_S_*, irrespective of *T_Ref_*, the derivative of Φ_F(internal)(*T*)_ will be zero at *T_S_*. Similarly, we can show by differentiating Eq. (45) that

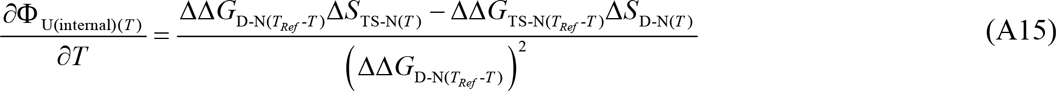

Once again, since Δ*S*_D-N(*T*)_ and Δ*S*_TS-N(*T*)_ are both zero at *T_S_*, irrespective of *T_Ref_*, the derivative of Φ_U(internal)(*T*)_ will be zero at *T_S_*.

---

**Figure 1-figure supplement 1.**
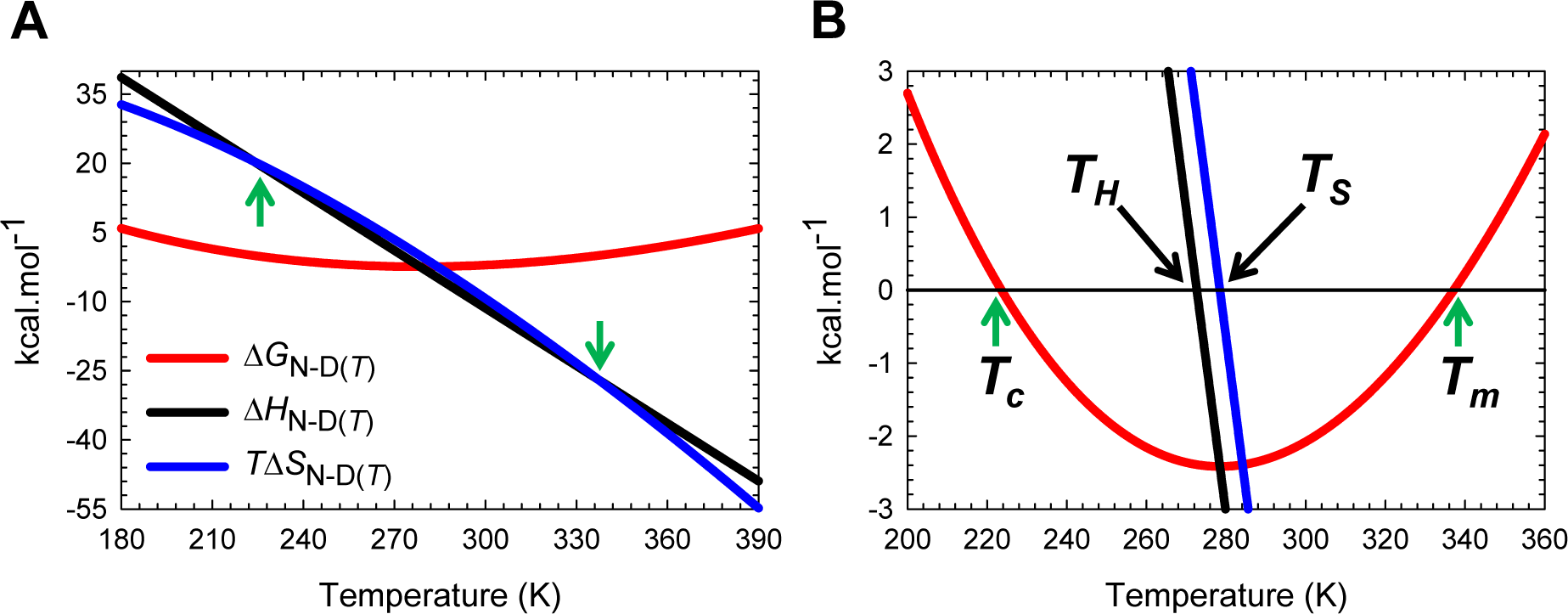
Stability curve for the folding reaction *N* ⇌ *D*. **(A)** Temperature-dependence of Δ*G*_N-D(*T*)_, Δ*H*_N-D(*T*)_, and *T*Δ*S*_N-D(*T*)_. The green pointers identify *T_c_* and *T_m_*. The slopes of the red and black curves are given by ∂Δ*G*_N-D(*T*)_/∂*T*=-Δ*S*_N-D(*T*)_ and ∂Δ*H*_N-D(*T*)_/∂*T*=Δ*C*_*p*N-D_, respectively. **(B)** An appropriately scaled version of plot on the left. The reference temperatures are as described in the parent figure.

**Figure 2−figure supplement 1.**
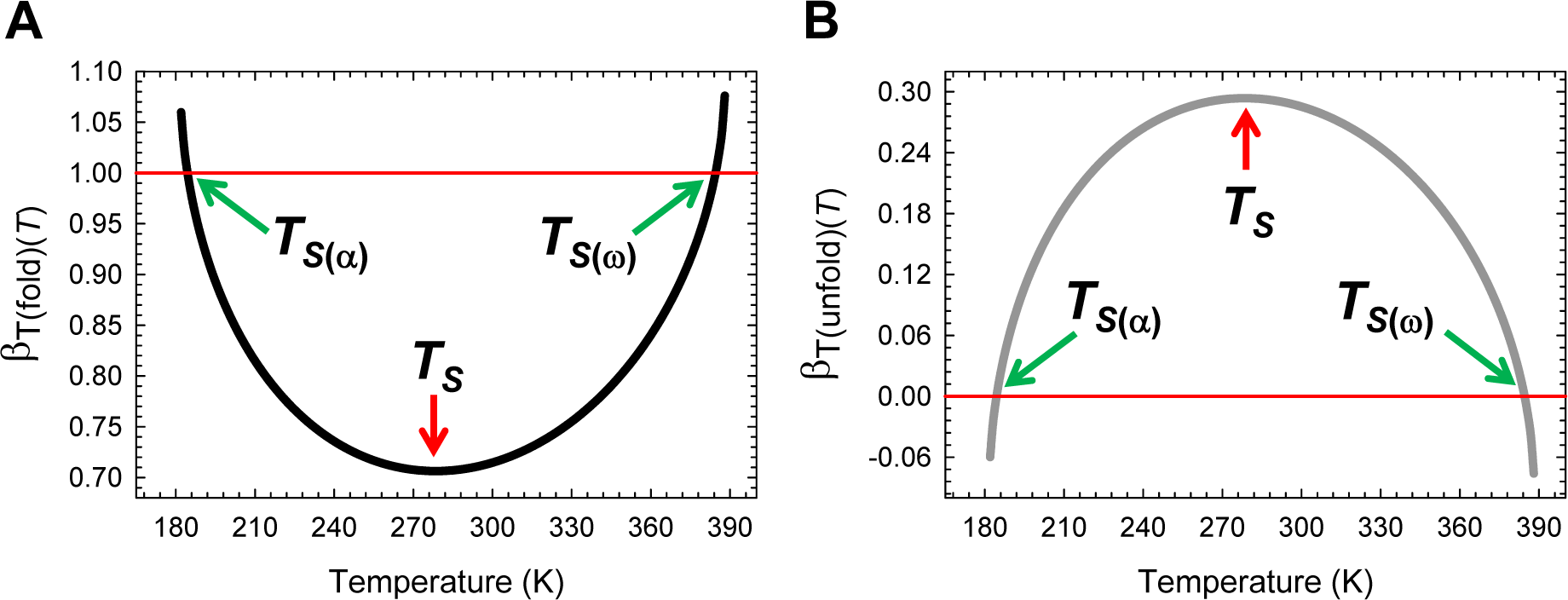
Temperature-dependence of β_T(fold)(*T*)_ and β_T(unfold)(*T*)_. **(A)** β_T(fold)(*T*)_ is a minimum at *T_S_*, unity at *T*_*S*(*α*)_ and *T*_*S*(*ω*)_, and greater than unity for *T_α_* ≤ *T* < *T*_*S*(*α*)_ and *T*_*S*(*ω*)_ < *T* ≤ *T_ω_*. The slope of this curve is given by 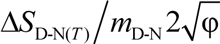 **(B)** β_T(unfold)(*T*)_ is a maximum at *T_S_*, zero at *T*_*S*(*α*)_ and *T*_*S*(*ω*)_, and negative for *T_α_* ≤ *T* < *T*_*S*(*α*)_ and *T*_*S*(*ω*)_ < *T* ≤ *T_ω_*. The slope of this curve is given by 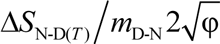. From the perspective of Tanford’s framework, the SASA of the TSE is the least native-like at *T_S_* but becomes progressively more native-like as the temperature deviates from the *T_S_*, and is identical to the SASA of the NSE at *T*_*S*(*α*)_ and *T*_*S*(*ω*)_; and for *T_α_* ≤ *T* < *T*_*S*(*α*)_ and *T*_*S*(*ω*)_ < *T* ≤ *T_ω_*, the TSE is more compact than the NSE.

**Figure 4−figure supplement 1.**
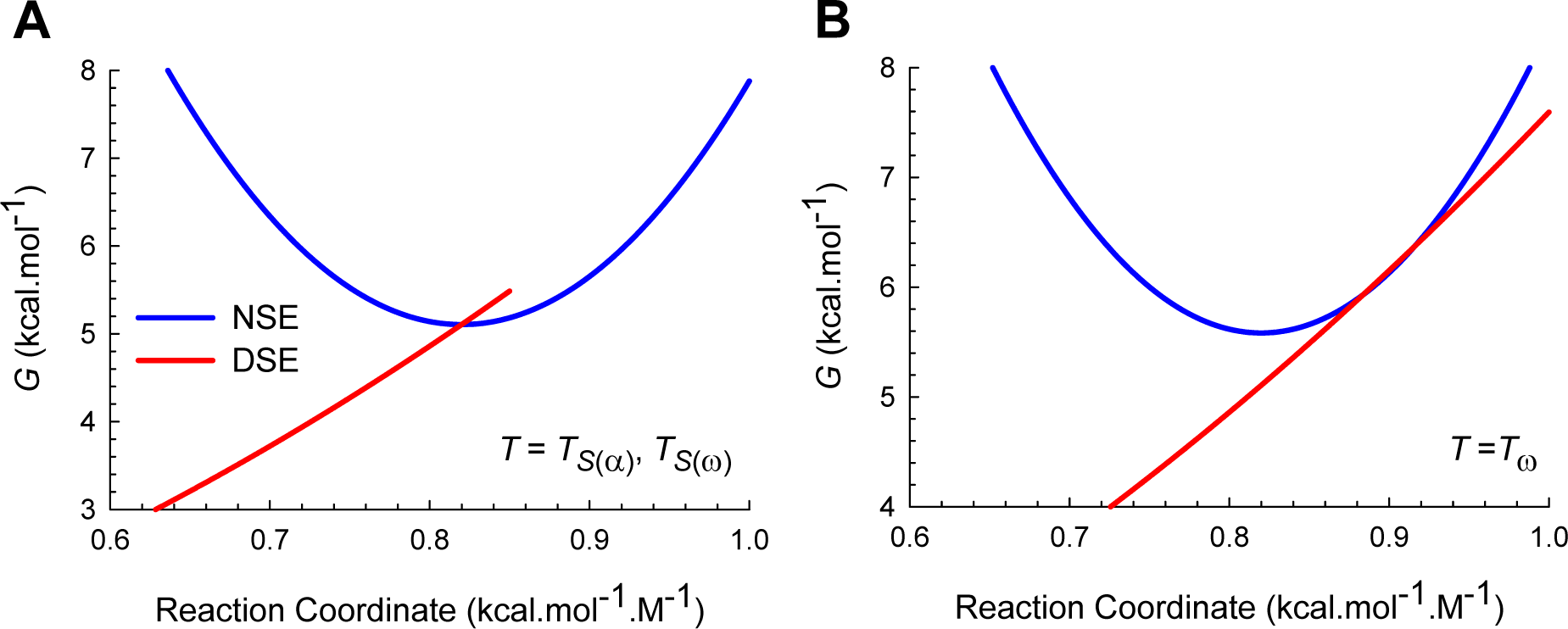
An appropriately scaled view of Marcus *curve-crossings* at *T*_*S*(ω)_ and *T*_ω_. **(A)** *Curve-crossing* at *T*_*S*(*α*)_ and *T*_*S*(*ω*)_ where *m*_TS-D(*T*)_ = *m*_D-N_, *m*_TS-N(*T*)_ = 0, Δ*G*_TS-N(*T*)_ = 0, Δ*G*_TS-D(*T*)_ = α(*m*_D-N_)^2^ = *λ*, Δ*G*_D-N(*T*)_ = − λ, and *k*_*u*(*T*)_ = *k*^0^. **(B)** *Curve-crossing* at *T_ω_* where *m*_TS-D(*T*)_ > *m*_D-N_.

**Figure 5−figure supplement 1.**
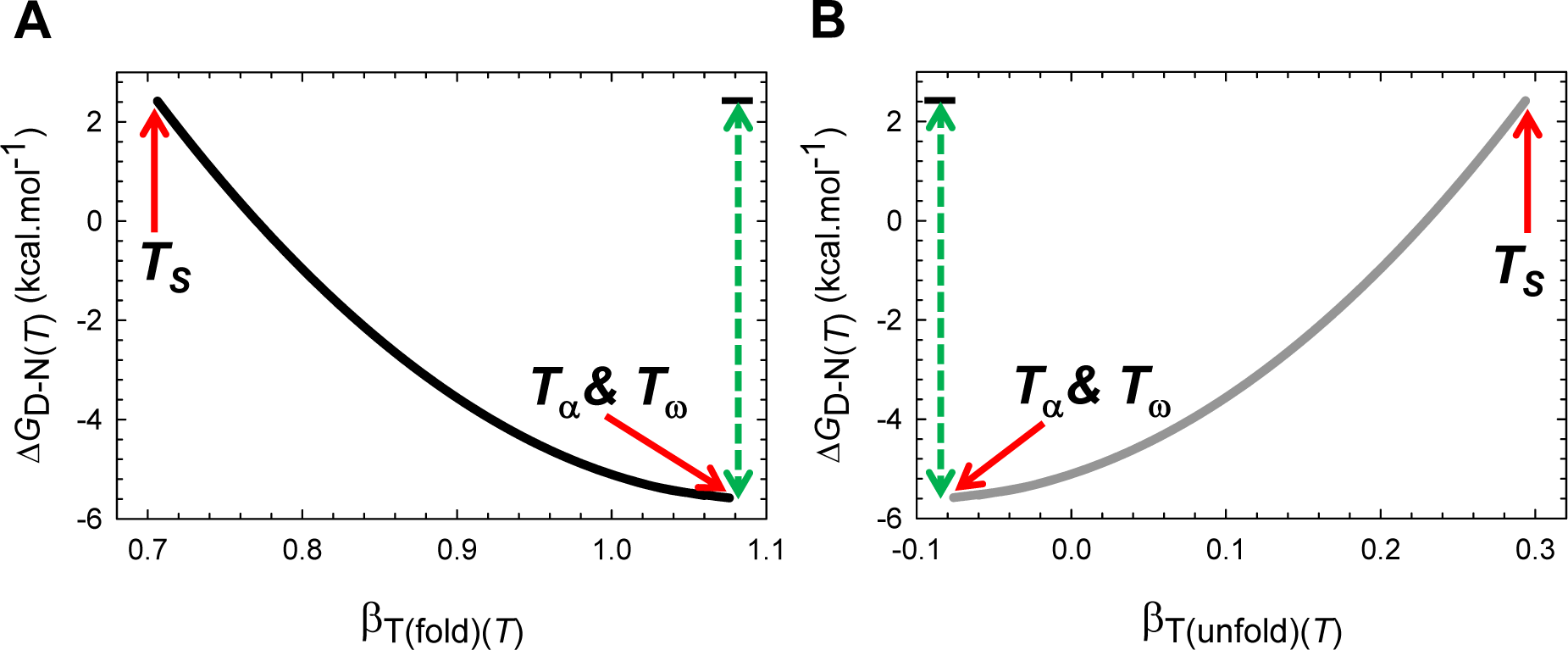
The principle of least displacement. **(A)** The stability of a two-state system at constant pressure and solvent conditions is the greatest when the denatured conformers are displaced the least from the mean of their ensemble along the SASA-RC to reach the TSE. The length of the green dotted line is identical to Δ*G*_D-N(*T_s_*)_ + [ λω/(ω-α)], where Δ*G*_D-N(*T_S_*)_ is the stability at *T_S_*. The slope of this curve equals 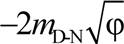. **(B)** Δ*G*_D-N(*T*)_ will be the greatest when the native conformers expose the greatest amount of SASA to reach the TSE. The slope of this curve equals 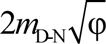.

**Figure 5−figure supplement 2.**
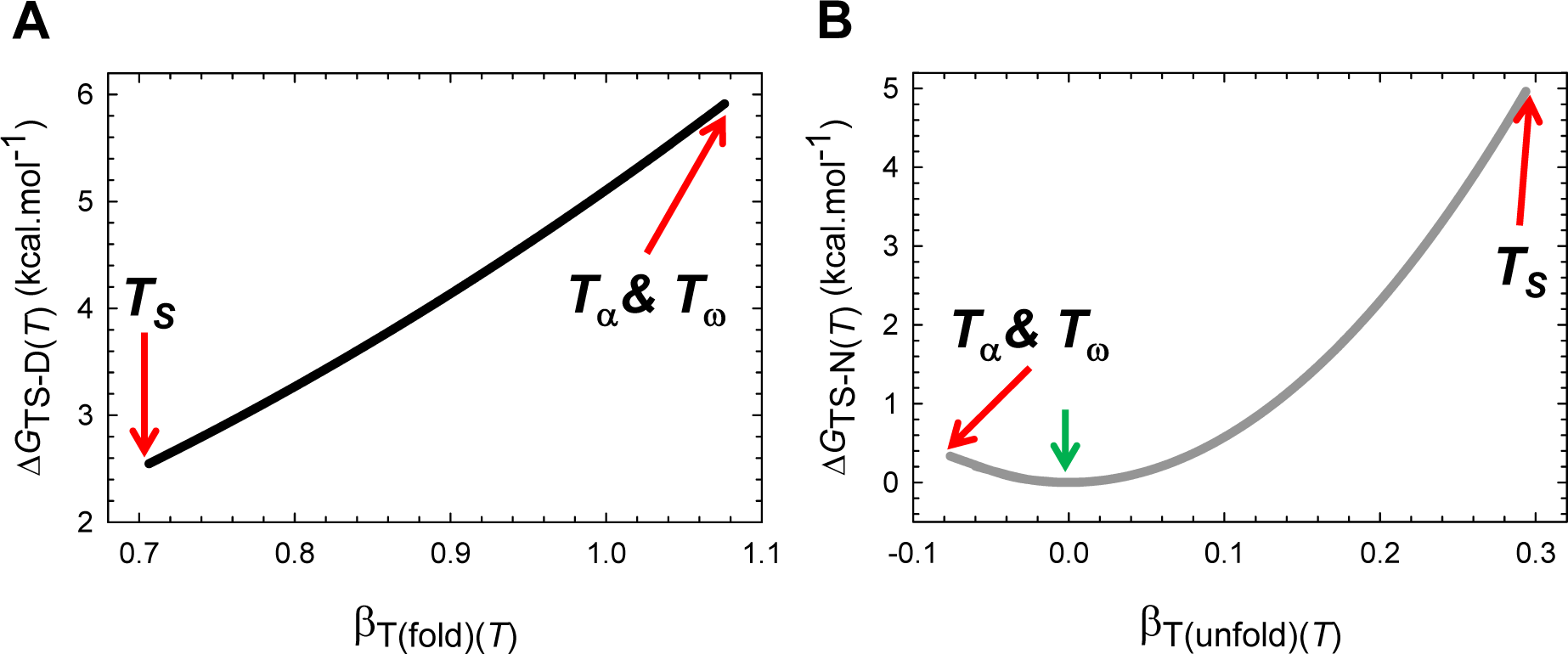
Gibbs activation energies as a function of the position of the TSE along the RC. **(A)** Δ*G*_TS-D(*T*)_ is the least when the denatured conformers bury the least amount of SASA to reach the TSE. The slope of this curve equals 2*λ*β_T(fold)(*T*)_. **(B)** Δ*G*_TS-N(*T*)_ is the greatest when the native conformers expose the greatest amount of SASA to reach the TSE. The green pointer indicates *T*_*S*(*α*)_ and *T*_*S*(*ω*)_ where *m*_TS-D(*T*)_ = *m*_D-N_, *m*_TS-N(*T*)_ = β_T(unfold)(*T*)_ = 0, Δ*G*_TS-N(*T*)_ = 0, Δ*G*_TS-D(*T*)_ = α(*m*_D-N_)^2^ = λ, and Δ*G*_D-N(*T*)_ = − λ. The slope of this curve equals 2ω*m*_TS-N(*T*)_*m*_D-N_. Because *m*_TS-N(*T*)_ < 0 for *T_α_* ≤ *T* < *T*_*S*(*α*)_ and *T*_*S*(*ω*)_ < *T* ≤ *T_ω_*, the slope is negative for the part that is to the left of the green pointer.

**Figure 6−figure supplement 1.**
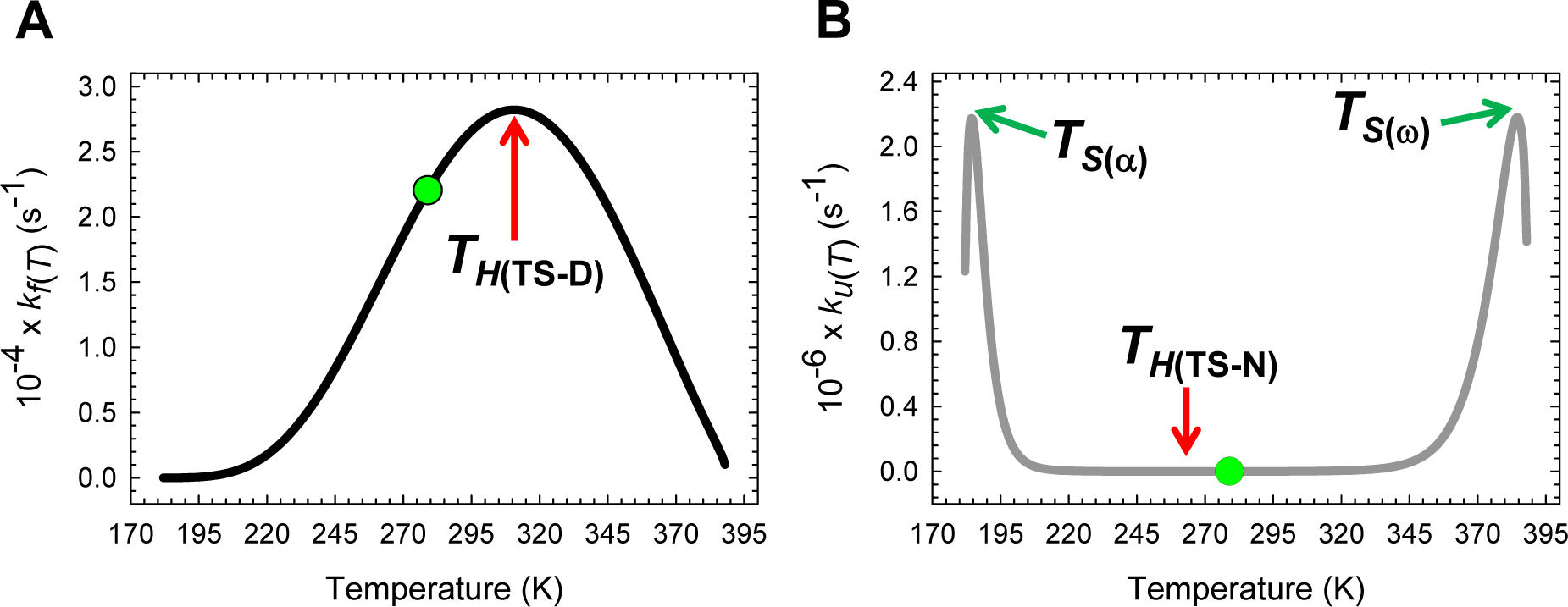
Temperature-dependence of *k*_*f*(*T*)_ and *k*_*u*(*T*)_ on a linear scale. **(A)** *k*_*f*(*T*)_ is a maximum and Δ*H*_TS-D(*T*)_ = 0 at *T*_*H*(TS-D)_. The slope of this curve is given by *k*_*f*(*T*)_∆*H*_TS-D(*T*)_/*RT*^2^. **(B)** Unlike *k*_*f*(*T*)_ which has only one extremum, *k*_*u*(*T*)_ is a minimum at *T*_*H*(TS-N)_ and a maximum at *T*_*S*(α)_ and *T*_*S*(ω)_. Although the minimum of *k*_*u*(*T*)_ is not apparent on a linear scale, the *barrierless* and *inverted-regimes* for unfolding are readily apparent. The slope of this curve is given by *k*_*u*(*T*)_∆*H*_TS-N(*T*)_/*RT*^2^. The features of these curves arise primarily from the temperature-dependence of the equilibrium constants for the partial folding (*D* ⇌ [*TS*]) and unfolding (*N* ⇌ [*TS*]) reactions as shown later. The green dots represent *T_S_*.

**Figure 6−figure supplement 2.**
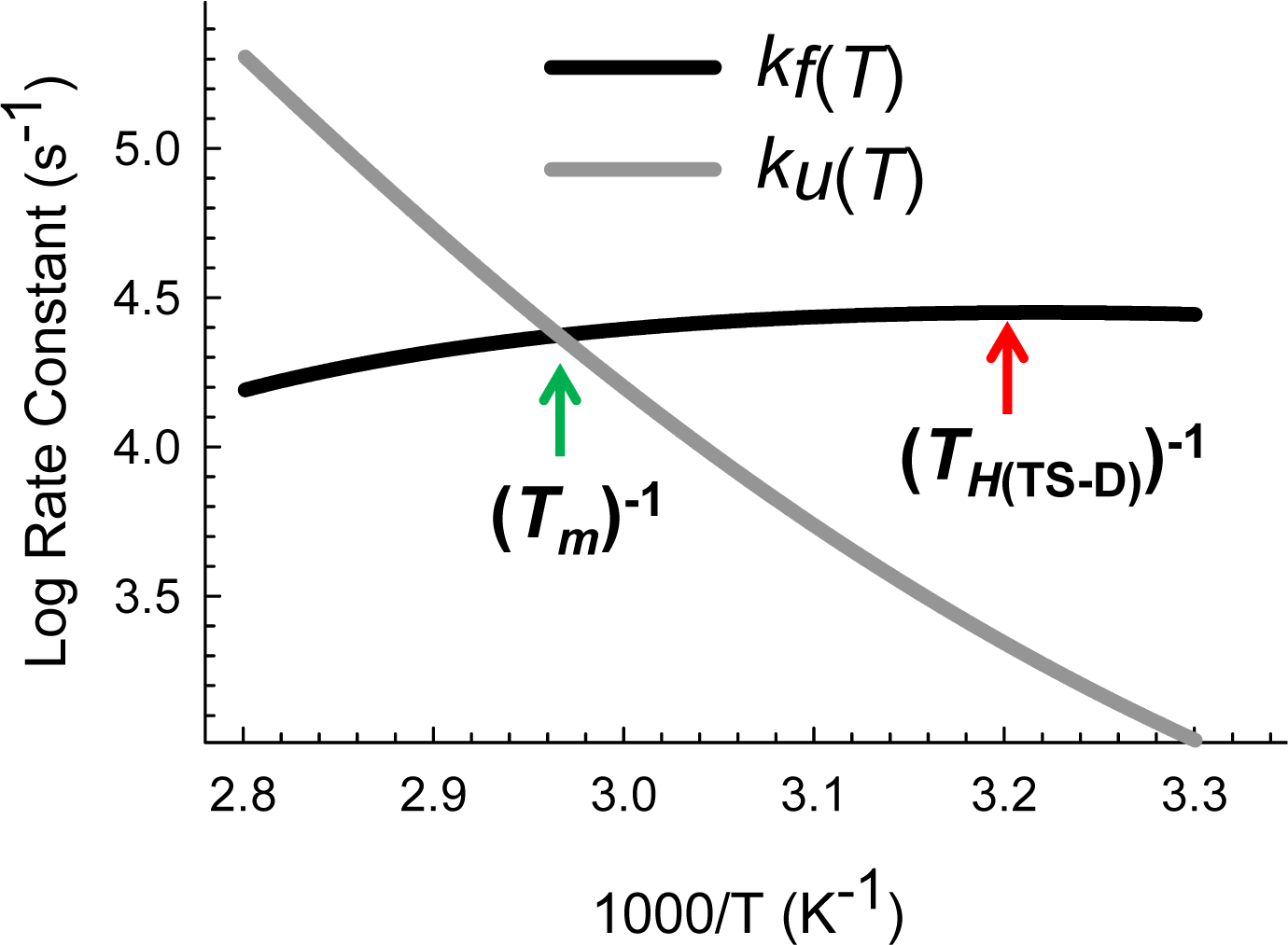
Arrhenius plot for the temperature-dependence of the rate constants with the ordinate on a Log scale (base10). A combined and appropriately rescaled version of Figure 6 to enable a ready comparison of the rate constants for FBP28 WW wild type (calculated using parabolic approximation) and the experimental rate constants for ΔNΔC Y11R W30F, a variant of FBP28 WW (reported by Nguyen et al., 2003, Fig. 4A). Note that the intersection of *k*_*f*(*T*)_ and *k*_*u*(*T*)_ is shifted to the left along the abscissa for the wild type FBP28 WW since its *T_m_* is ∽ 10 K greater than that of ΔNΔC Y11R W30F (see Table 1 in Nguyen et al., 2003).^25^

**Figure 7−figure supplement 1.**
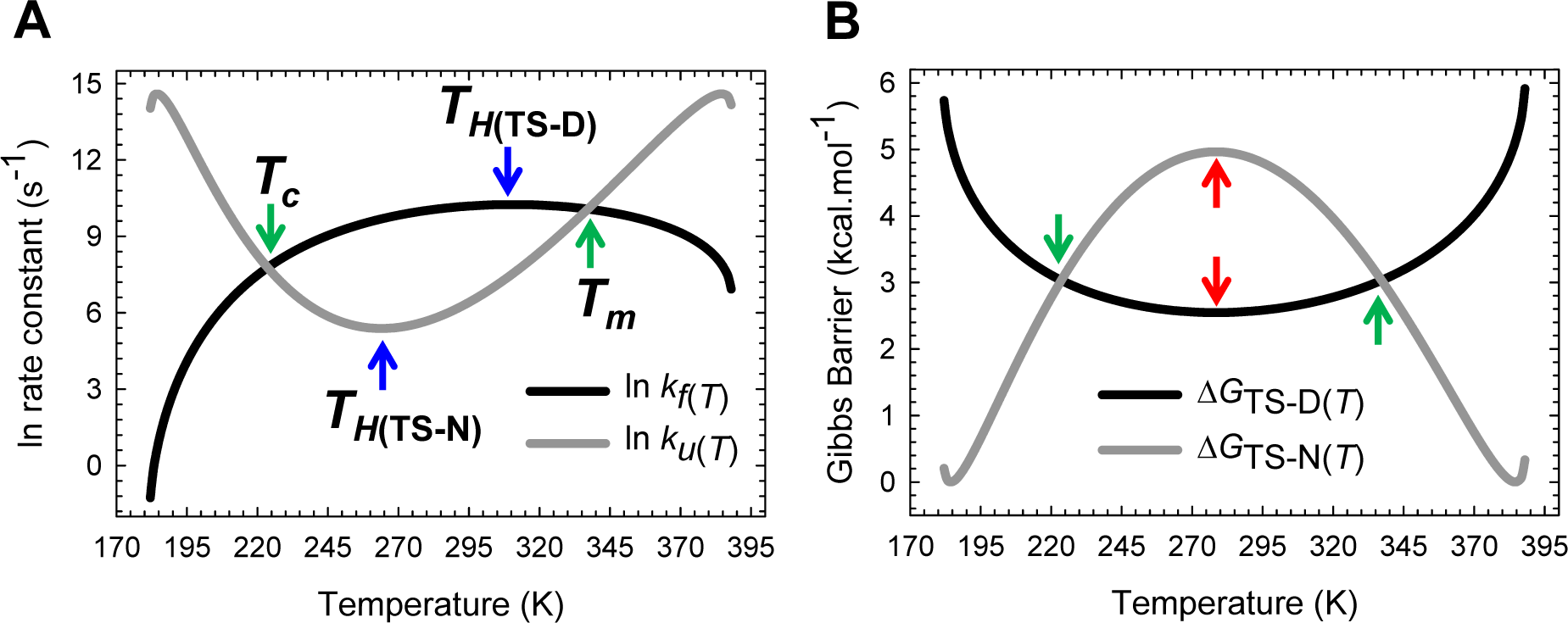
The principle of microscopic reversibility. **(A)** *k*_*f*(*T*)_ is a maximum at *T*_*H*(TS-D)_ and *k*_*u*(*T*)_ is a minimum at *T*_*H*(TS-N)_. The slopes of the black and grey curves are given by ∆*H*_TS-D(*T*)_/*RT*^2^ and ∆*H*_TS-N(*T*)_/*RT*^2^, respectively. **(B)** Δ*G*TS-D(*T*) and Δ*G*TS-N(*T*) are a minimum and a maximum, respectively, at *T_S_* (red pointers) leading to Δ*G*_D-N(*T*)_ being a maximum at *T_S_* (**Figure 1**). Equilibrium stability is thus a consequence or the equilibrium manifestation of the underlying kinetic behaviour. The rate constants are identical at *T_c_* and *T_m_*, leading to ∆*G*_D-N(*T*)_=*RT* ln (*k*_*f*(*T*)_/*k*_*u*(*T*)_)=∆*G*_TS-N(*T*)_-∆*G*_TS-D(*T*)_=0.

**Figure 9−figure supplement 1.**
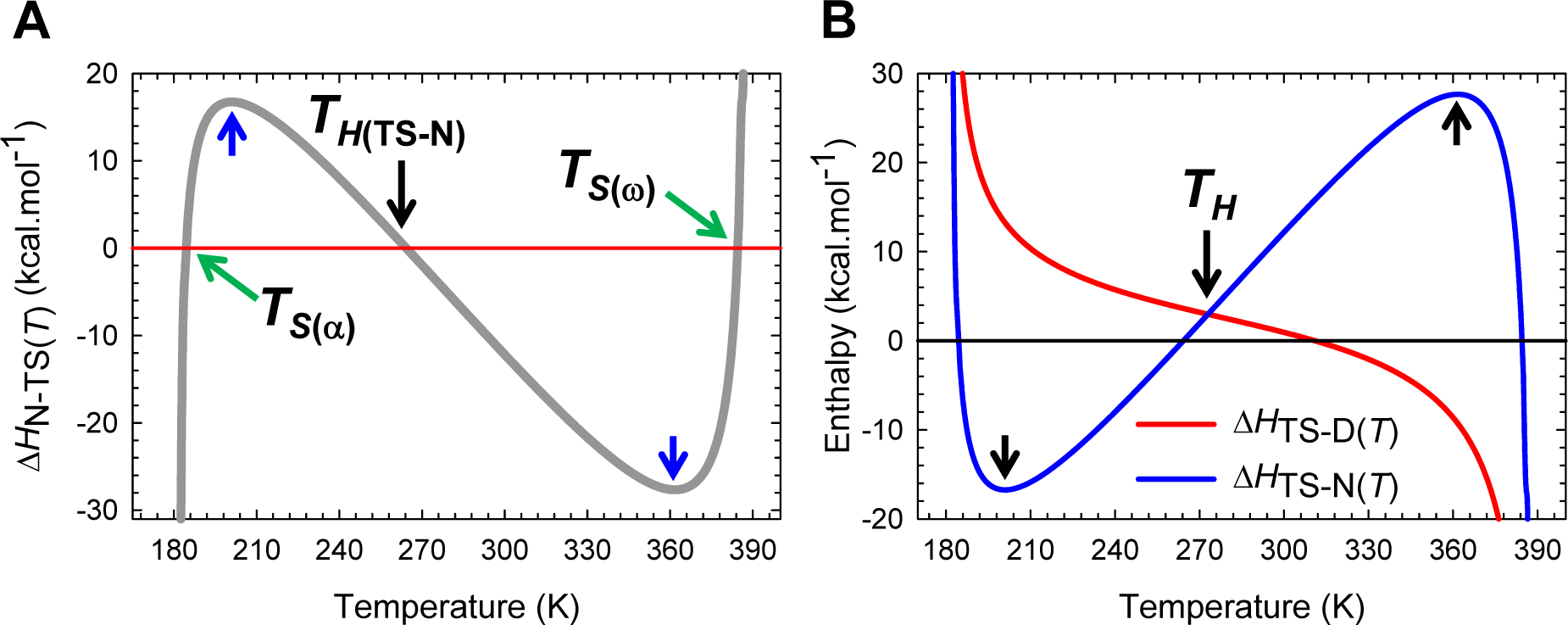
The variation in Δ*H*_N-TS(*T*)_ with temperature and the intersection of Δ*H*_TS-D(*T*)_ and Δ*H*_TS-N(*T*)_ functions. **(A)** An appropriately scaled view of the change in enthalpy for the partial folding reaction [*TS*] ⇌ *N*. The flux of the conformers from the TSE to the NSE is enthalpically: (*i*) favourable for *T*_*α*_ ≤ *T* < *T*_*S*(*α*)_ and *T*_*H*(TS-N)_ < *T* < *T*_*S*(*ω*)_ (Δ*H*_N-TS(*T*)_ < 0); (*ii*) unfavourable for *T*_*S*(*α*)_ < *T* < *T*_*H*(TS-N)_ and *T*_*S*(*ω*)_ < *T* ≤ *T_ω_* (Δ*H*_N-TS(*T*)_ > 0); and (*iii*) neither favourable nor unfavourable at *T*_*S*(*α*)_, *T*_*H*(TS-N)_, and *T*_*S*(*ω*)_. The blue pointers indicate the temperatures where Δ*C*_*p*N-TS(*T*)_ (or −Δ*C*_*p*TS-N(*T*)_) is zero. **(B)** The intersection of the Δ*H*_TS-D(*T*)_ and Δ*H*_TS-N(*T*)_ functions occurs precisely at *T_H_*. The requirement that both Δ*H*_TS-D(*T*)_ and Δ*H*_TS-N(*T*)_ be positive at the point of intersection is a consequence of the theoretical relationship: *T*_*H*(TS-N)_ < *T_H_* < *T_S_* < *T*_*H*(TS-D)_ and must be satisfied by all two-state systems (see Paper II). Note that the net flux of the conformers from the DSE to the NSE at *T_H_* is driven purely by entropy (Δ*G*_D-N(*T*)_ = –*T* Δ*S*_D-N(*T*)_).

**Figure 9−figure supplement 2.**
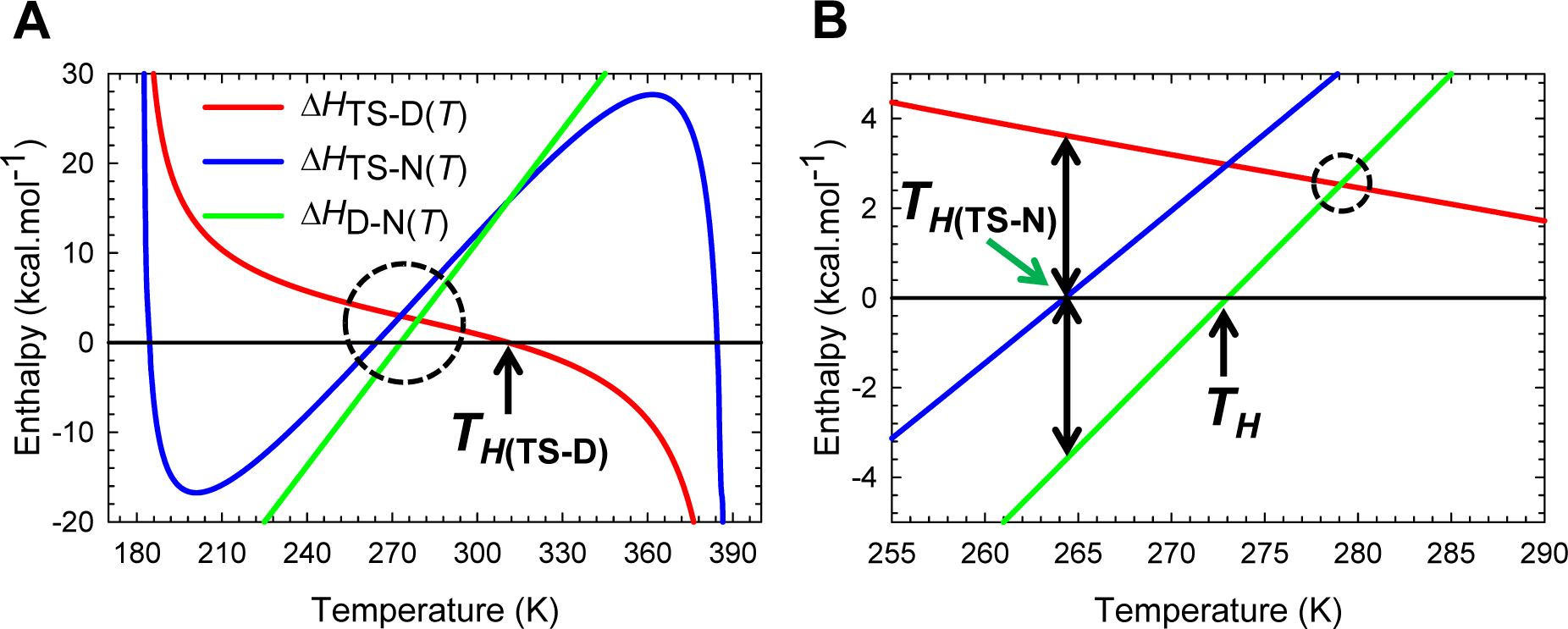
Comparison of equilibrium and activation enthalpies. **(A)** Δ*H*_D-N(*T*)_ for the reaction *N* ⇌ *D* is zero at the temperature where Δ*H*_TS-D(*T*)_ and Δ*H*_TS-N(*T*)_ functions intersect (the intersection of green curve and zero reference line must align vertically with the point where the blue and the red curves intersect). The intersection of Δ*H*_D-N(*T*)_ and Δ*H*_TS-N(*T*)_ functions (green and blue curves) occurs precisely when *T* = *T*_*H*(TS-D)_. This is expected since Δ*H*_TS-D(*T*)_ = 0 at *T*_*H*(TS-D)_. The similarity in the slopes of the Δ*H*_D-N(*T*)_ and Δ*H*_TS-N(*T*)_ functions between ∽ 240 K and ∽ 320 K implies that most of Δ*C*_*p*D-N_ stems from the first-half of the unfolding reaction *N* ⇌ [*TS*]. **(B)** An appropriately scaled view of the encircled area in the figure on the left. When *T* = *T*_*H*(TS-N)_, Δ*H*_TS-D(*T*)_ is identical to |Δ*H*_D-N(*T*)_| or Δ*H*_N-D(*T*)_. Further, at the temperature where Δ*H*_TS-D(*T*)_ and Δ*H*_D-N(*T*)_ functions intersect (i.e., the intersection of the red and the green curves), the absolute enthalpy of the DSE (*H*_D(*T*)_) is exactly half the algebraic sum of the absolute enthalpies of the TSE (*H*_TS(*T*)_) and the NSE (*H*_N(*T*)_), i.e., *H*_D(*T*)_ = (*H*_TS(*T*)_+*H*_N(*T*)_)/2. The various auxiliary relationships that may obtained from the intersection of various state functions are addressed in subsequent publications.

**Figure 10−figure supplement 1.**
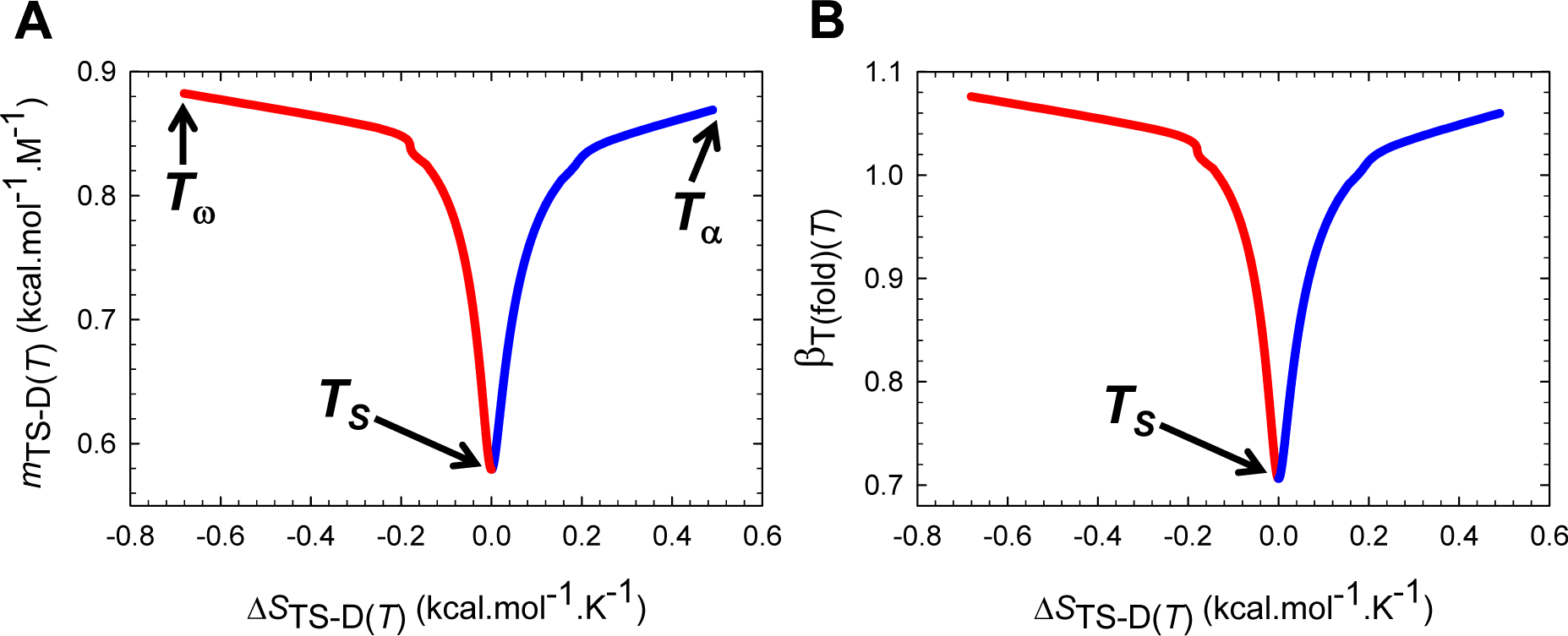
Activation entropy for folding *vs curve-crossing*. **(A)** Δ*S*_TS-D(*T*)_ is zero when the denatured conformers are displaced the least from the mean of their ensemble to reach the TSE along the SASA-RC. The slope of this curve is given by 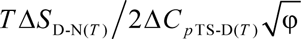 **(B)** Δ*S*_TS-D(*T*)_ is zero when the SASA of the TSE is the least native-like. The slope of this curve is given by 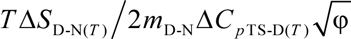. The blue and the red sections of the curves represent the temperature regimes *T_α_* ≤ *T* ≤ *T_S_* and *T_S_* ≤ *T* ≤ *T_ω_*, respectively.

**Figure 10−figure supplement 2.**
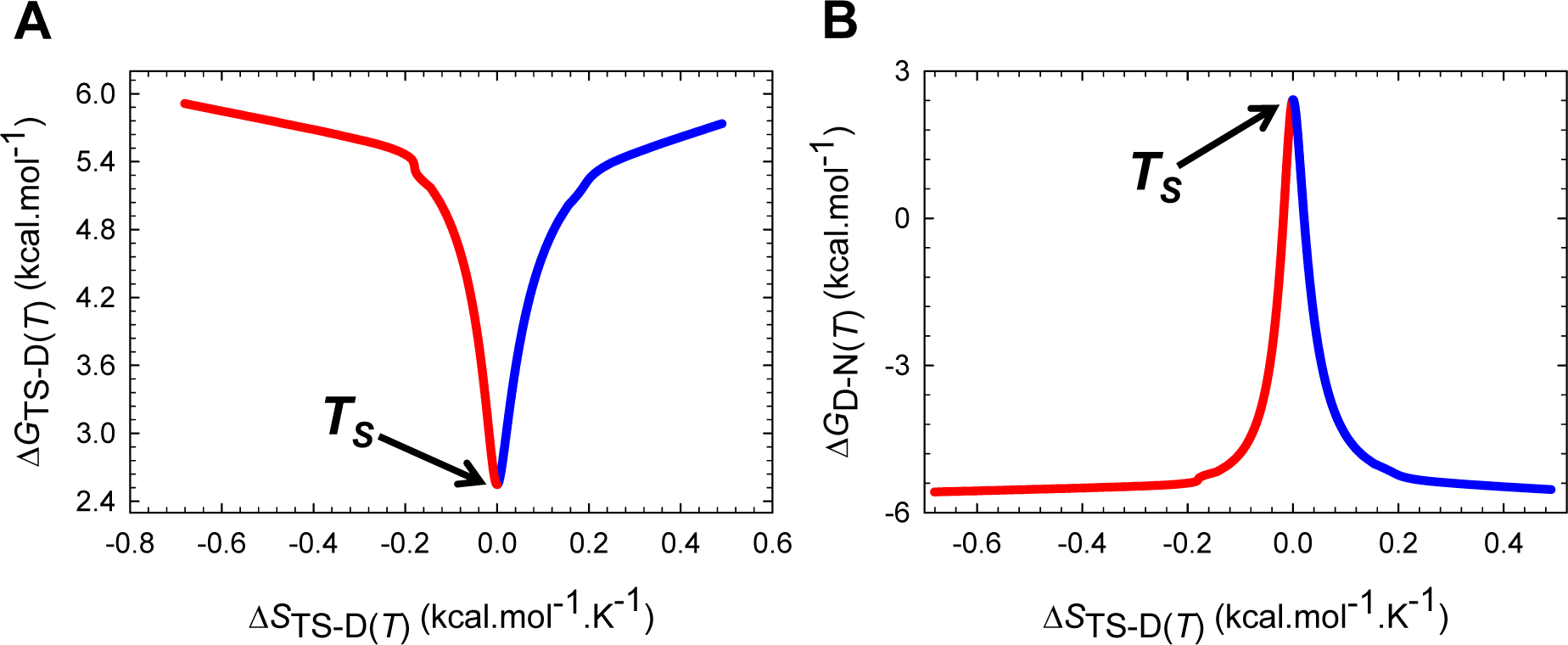
Activation entropy *vs* Δ*G*_TS-D(*T*)_ and Δ*G*_D-N(*T*)_. **(A)** Δ*G*_TS-D(*T*)_ is always the least when it is purely enthalpic. The slope of this curve equals-TΔ*S*_TS-D(*T*)_/Δ*C*_*p*TS-D(*T*)_. **(B)** The stability is always the greatest when the activation entropy for folding is the zero. The slope of this curve equals-TΔ*S*_TS-D(*T*)_/Δ*C*_*p*TS-D(*T*)_. The blue and the red sections of the curves represent the temperature regimes *T_α_* ≤ *T* ≤ *T_S_* and *T_S_* ≤ *T* ≤ *T_ω_*, respectively.

**Figure 11−figure supplement 1.**
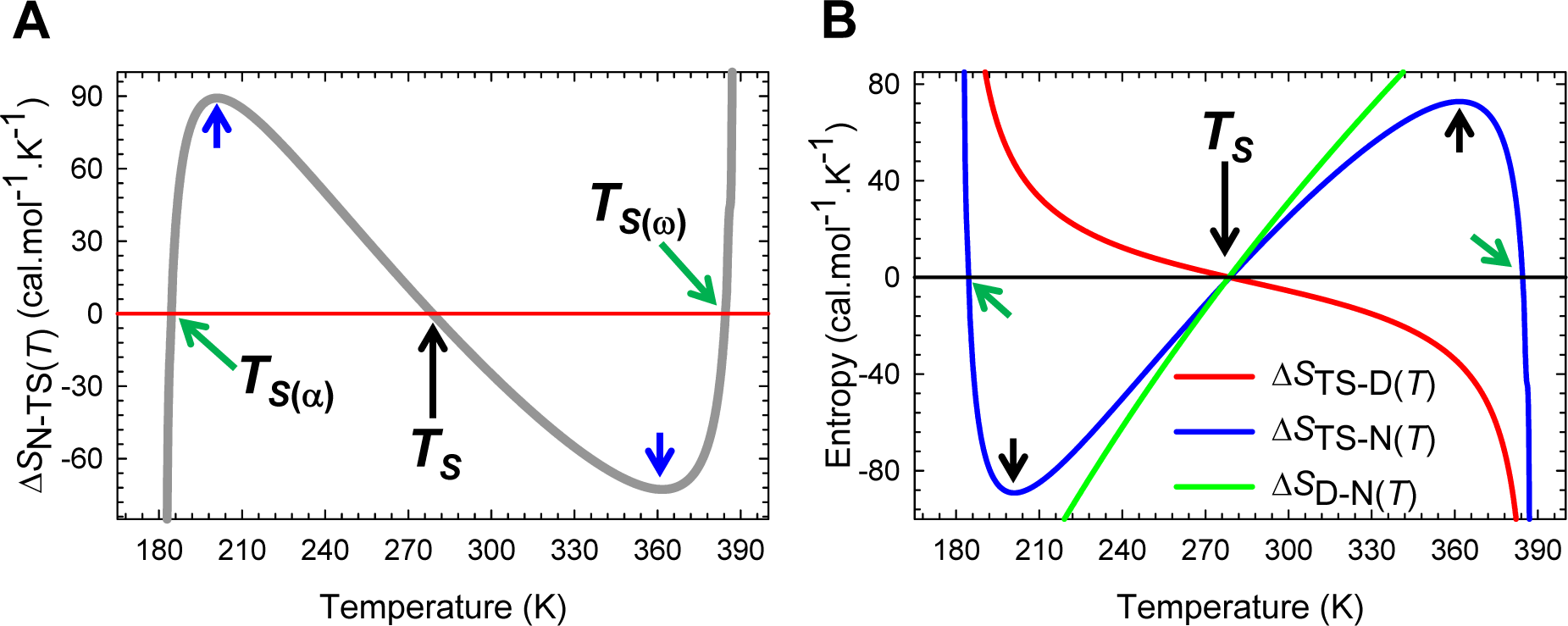
The variation in Δ*S*_N-TS(*T*)_ with temperature and comparison of equilibrium and activation entropies. **(A)** An appropriately scaled view of the change in entropy for the partial folding reaction [*TS*] ⇌ *N*. The slope of this curve equals Δ*C*_*p*N-TS(*T*)_/*T* (or −Δ*C*_*p*N-TS(*T*)_/*T* and is zero at *T*_*C_p_*TS-N(*α*)_ and *T*_*C_p_*TS-N(*ω*)_. The flux of the conformers from the TSE to the NSE is entropically: (*i*) unfavourable for *T_α_* ≤ *T* < *T*_*S*(*α*)_ and *T_S_* < *T* < *T*_*S*(*ω*)_ (Δ*S*_N-TS(*T*)_ < 0); (*ii*) favourable for *T*_*S*(*α*)_ < *T* < *T_S_* and *T*_*S*(*ω*)_ < *T* ≤ *T_ω_* (Δ*S*_N-TS(*T*)_ > 0); and (*iii*) neutral at *T*_*S*(*α*)_, *T_S_*, and *T*_*S*(*ω*)_. **(B)** An overlay of Δ*S*_D-N(*T*)_, Δ*S*_TS-D(*T*)_ and Δ*S*_TS-N(*T*)_ functions. Unlike the Δ*H*_TS-D(*T*)_ and Δ*H*_TS-N(*T*)_ functions which must be positive at the point of intersection (**Figure 9−figure supplement 1B**), theory dictates that both Δ*S*_TS-D(*T*)_ and Δ*S*_TS-N(*T*)_ functions must independently be equal to zero at *T_S_*, leading to the unique scenario: *S*_D(*T*)_ = *S*_TS(*T*)_ = *S*_N(*T*)_. The similarity in the slopes of the Δ*S*_D_N(*T*) and Δ*S*_TS-N(*T*)_ functions between ∽ 240 K and ∽ 320 K implies that most of Δ*C*_*p*D-N_ stems from the first-half of the unfolding reaction *N* ⇌ [*TS*]. Consequently at *T_S_*, Δ*G*_D-N(*T*)_ = Δ*H*_D_N(*T*), i.e., the net flux of the conformers from the DSE to the NSE is driven purely by enthalpy.

**Figure 11−figure supplement 2.**
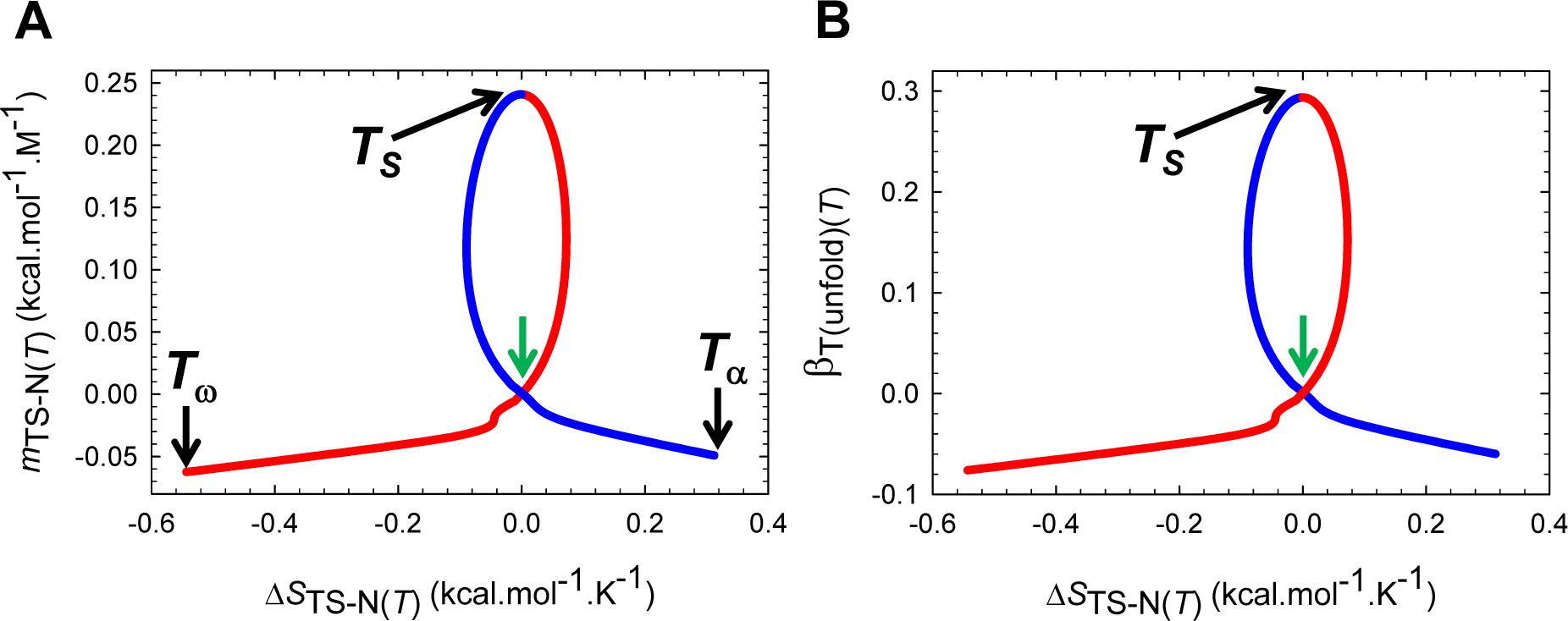
Activation entropy for unfolding *vs curve-crossing*. **(A)** Unlike the Δ*S*_TS-D(*T*)_ function which is zero only once, Δ*S*_TS-N(*T*)_ is zero once when the native conformers are displaced the greatest to reach the TSE (*T_S_*), and twice when this displacement is zero (green pointer; *T*_*S*(*α*)_ and *T*_*S*(*ω*)_). The slope of this curve is given by 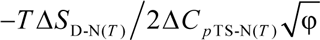. **(B)** Δ*S*_TS-N(*T*)_ is zero once when the difference in SASA between the TSE and the NSE is the greatest (*T_S_*), and twice when the SASA of the TSE is identical to that of the NSE (green pointer; *T*_*S*(*α*)_ and *T*_*S*(*ω*)_). The slope of this curve is given by 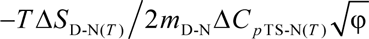. The blue and the red sections of the curves represent the temperature regimes *T_α_* ≤ *T* ≤ *T_S_* and *T_S_* ≤ *T* ≤ *T_ω_*, respectively.

**Figure 11−figure supplement 3.**
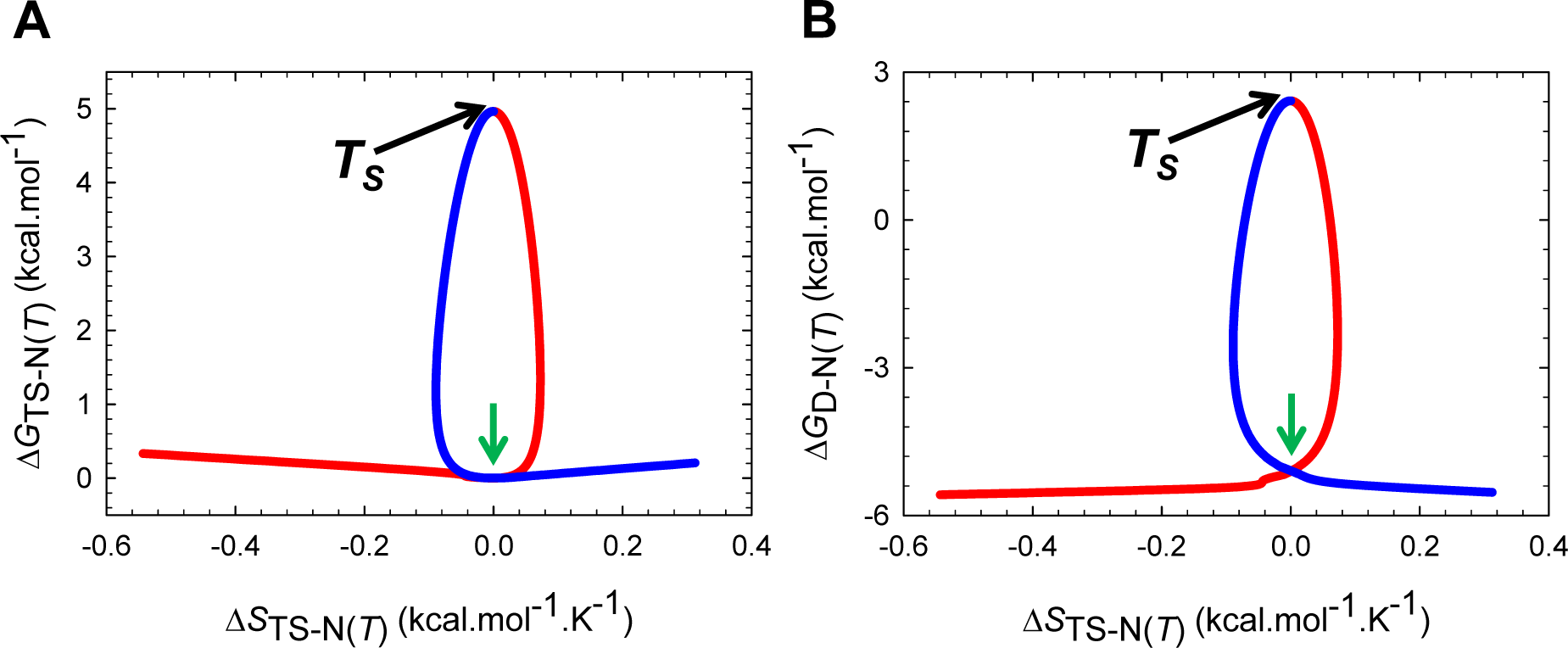
Activation entropy *vs* Δ*G*_TS-N(*T*)_ and Δ*G*_D-N(*T*)_ **(A)** Δ*G*_TS-N(*T*)_ is the greatest and also the least (zero) when it is purely enthalpic, with the former occurring at *T_S_*, and the latter occurring at *T*_*S*(*α*)_ and *T*_*S*(*ω*)_ (green pointer). The slope of this curve equals −*T*Δ*S*_TS-N(*T*)_/Δ*C*_*p*TS-N(*T*)_. **(B)** The stability is always the greatest at *T_S_* where the Gibbs barrier to unfolding is purely enthalpic; and at *T*_*S*(*α*)_ and *T*_*S*(*ω*)_ (green pointer), Δ*G*_D-N(*T*)_ = − *λ*. The slope of this curve equals −*T*Δ*S*_D-N(*T*)_/Δ*C*_*p*TS-N(*T*)_. The blue and the red sections of the curves represent the temperature regimes *T_α_* ≤ *T* ≤ *T_S_* and *T_S_* ≤ *T* ≤ *T_ω_*, respectively.

**Figure 11−figure supplement 4.**
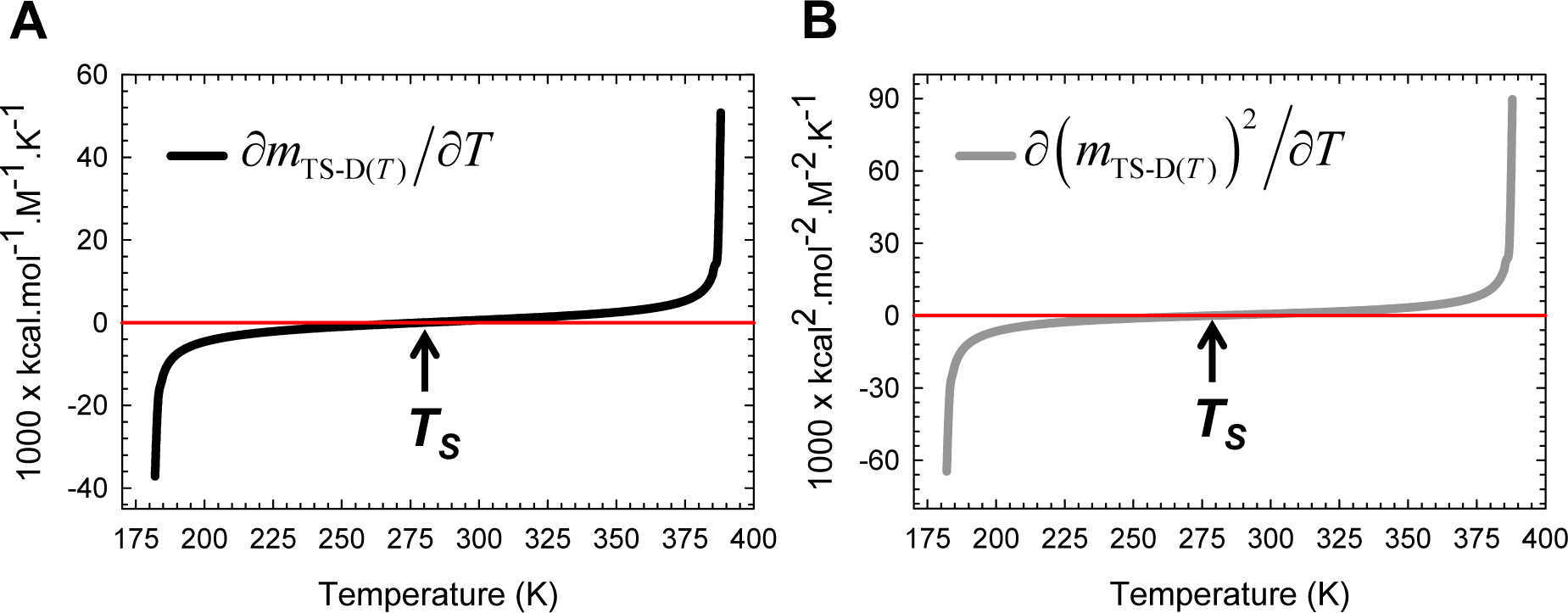
The first derivatives of *m*_TS-D(*T*)_ and the square of *m*_TS-D(*T*)_ with respect to temperature. **(A)** *∂m*_TS-D(*T*)_ is negative for *T_α_* ≤ *T* < *T_S_*, positive for *T_S_* < *T* ≤ *T_ω_*, and zero at *T_S_* and is dictated by Eq. (A4). **(B)** Because *∂*(*m*_TS-D(*T*)_)^2^/*∂T*=2*m*_TS-D(*T*)_(*∂m*_TS-D(*T*)_/*∂T*) and *m*_TS-D(*T*)_ > 0 throughout the temperature regime, the variation of its algebraic sign is identical to that of *∂m*_TS-D(*T*)_/*∂T*. The relationship between *∂m*_TS-D(*T*)_/*∂T* and Δ*S*_TS-D(*T*)_ is given by Eq. (8).

**Figure 11−figure supplement 5.**
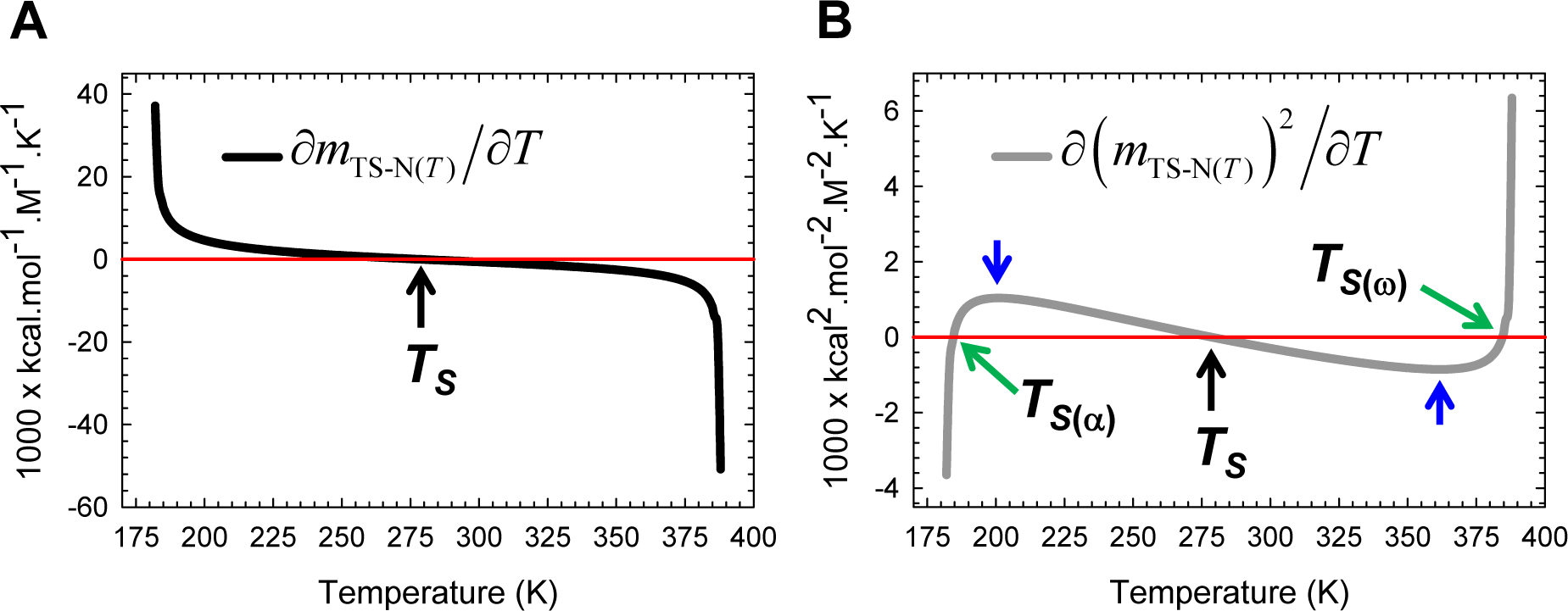
The first derivatives of *m*_TS-N(*T*)_ and the square of *m*_TS-N(*T*)_ with respect to temperature. **(A)** ∂*m*_TS-N(*T*)_/∂*T* is positive for *T*_α_≤*T* < *T_S_*, negative for *T_S_* < *T* ≤ *T*_ω_, and zero at *T_S_* and is governed by Eq. (A6). **(B)** Because ∂(*m*_TS-N(*T*)_)^2^/∂*T* = 2*m*_TS-N(*T*)_(∂*m*_TS-N(*T*)_/∂*T*) and *m*_TS-N(*T*)_ can be negative, zero or positive depending on the temperature, the variation of its algebraic sign with temperature is far more complex: (*i*) ∂(*m*_TS-N(*T*)_^2^/∂*T* is negative for *T*_α_≤ *T* < *T_S(α)_* and *T_S_* < *T* < *T_S(ω)_*; (*ii*) positive for *T_S(α)_* < *T* < *T_S_* and *T_S(ω)_* < *T* ≤ *T*_ω_; and (*iii*) zero at *T*_*S*(*α*)_, *T_S_*, and *T*_*S*(ω)_. The relationship between *∂m*_TS-N(*T*)_/∂*T* and Δ*S*_TS-N(*T*)_ is given by Eq. (9). The blue pointers indicate the temperatures at which the second derivative of the square of *m*_TS-N(*T*)_ is zero and are identical to the temperatures at which Δ*C*_*p*TS-N(*T*)_ is zero.

**Figure 12−figure supplement 1.**
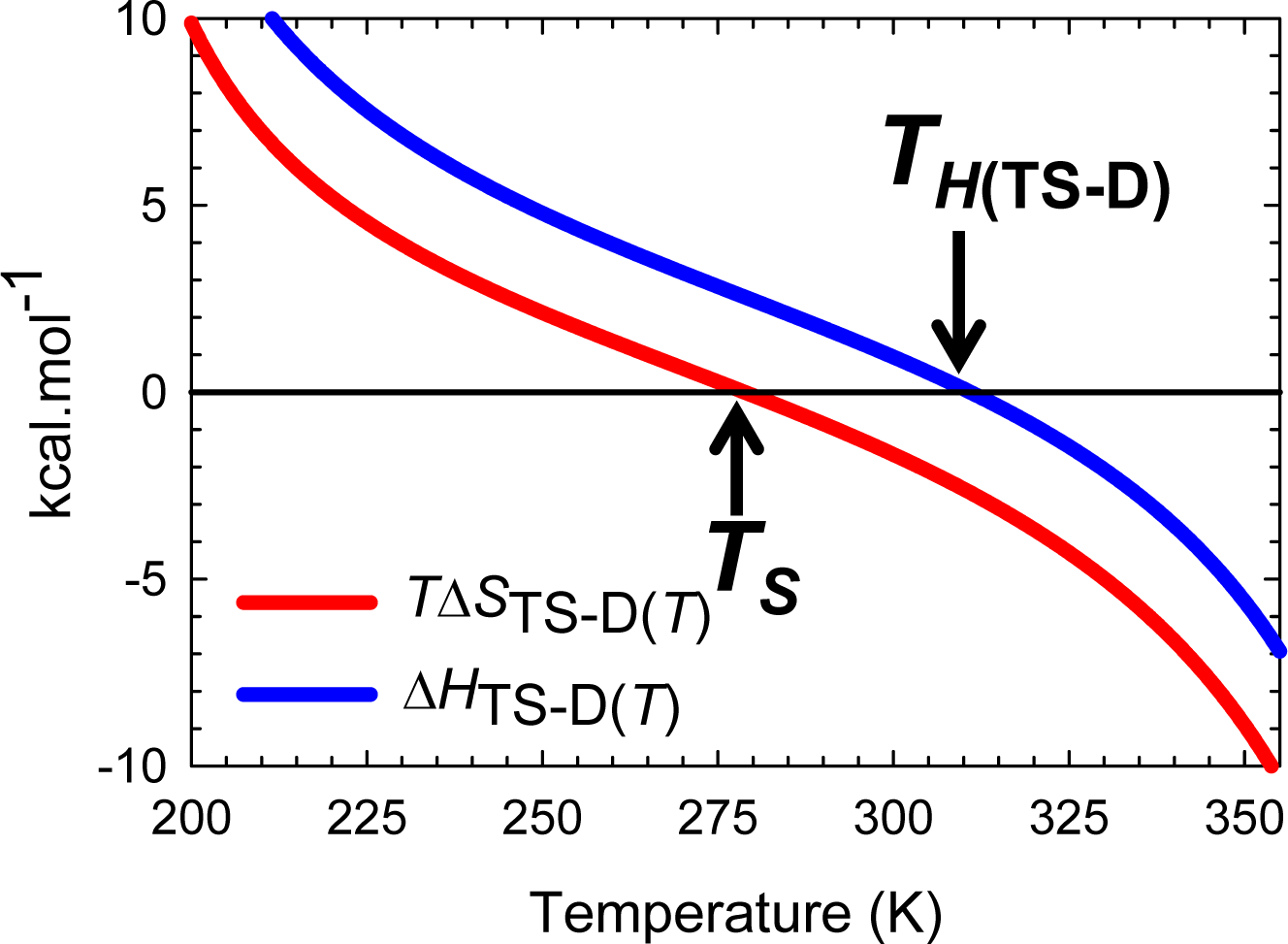
Deconvolution of the Gibbs activation energy for the reaction *D* ⇌ [*TS*]. This is an appropriately scaled view of **Figure 12B**. For *T*_α_ ≤ *T* < *T_S_*, *T*Δ*S*_TS-D(*T*)_ > 0 but is more than offset by unfavourable Δ*H*_TS-D(*T*)_, leading to incomplete compensation and a positive Δ*G*_TS-D(*T*)_ − (*ΔH*_TS-D(*T*)_ *T*Δ*S*_TS-D(*T*)_. When *T* = *T_S_*, Δ*G*_TS-D(*T*)_ is a minimum and purely enthalpic (Δ*G*_TS-D(*T*)_ = Δ*H*_TS-D(*T*)_ > 0). For *T_S_* < *T* < *T*_*H*(TS-D)_, the activation is enthalpically and entropically disfavoured (Δ*H*_TS-D(*T*)_ > 0 and *T*Δ*S*_TS-D(*T*)_< 0) leading to a positive Δ*G*_TS-D(*T*)_. In contrast, for *T*_*H*(TS-D)_ < *T* ≤ *T*_ω_, Δ*H*_TS-D(*T*) < 0_ but is more than offset by the unfavourable entropy (*T*Δ*S*_TS-D(*T*)_ < 0), leading once again to a positive Δ*G*_TS-D(*T*)_. When *T* = *T*_*H*(TS-D)_, Δ*G*_TS-D(*T*)_ is purely entropic (Δ*G*_TS-D(*T*)_ = −*TΔS*_TS-D(*T*)_ > 0) and *k*_*f*(*T*)_ is a maximum.

**Figure 13-figure supplement 1.**
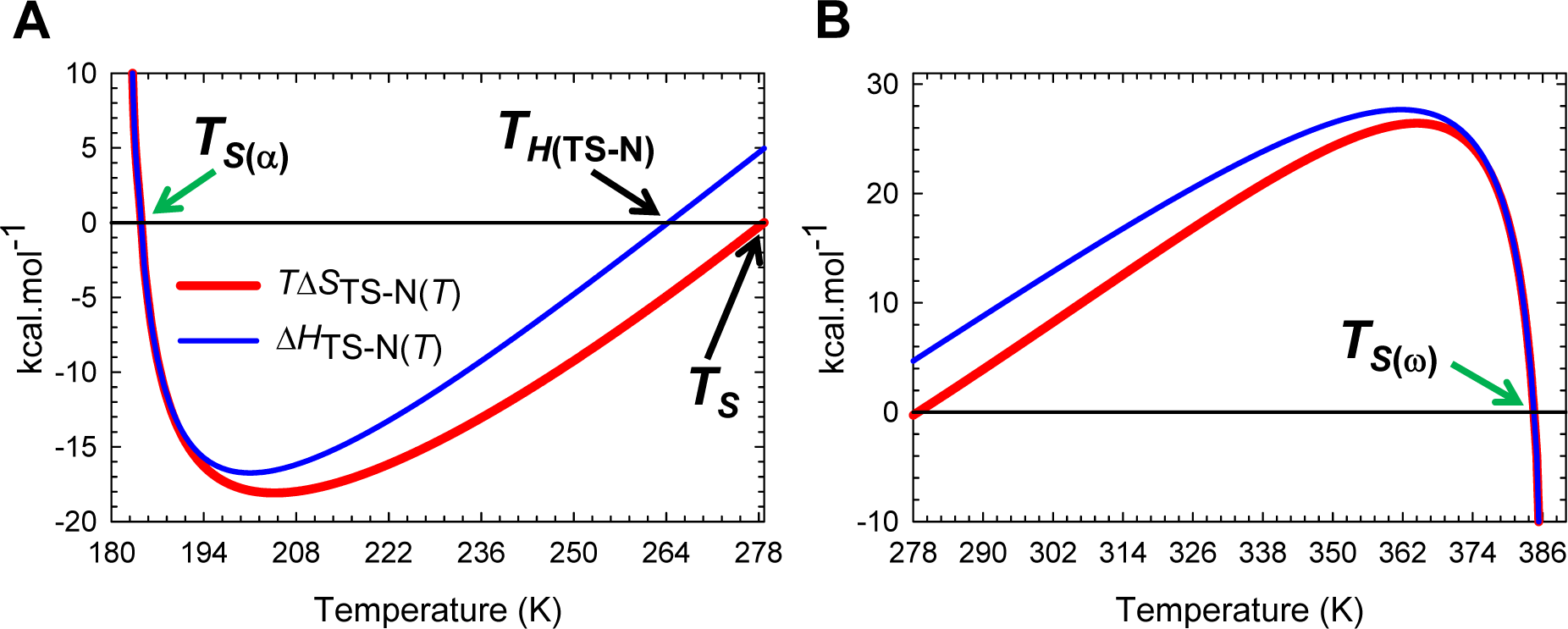
Deconvolution of the Gibbs activation energy for unfolding. These are appropriately scaled split views of **Figure 13B. (A)** For *T*_α_ ≤ *T* < *T*_*S*(*α*)_, *N* ⇌ [*TS*] entropically favoured (*T*Δ*S*_TS-N(*T*)_ > 0) but is more than offset by endothermic enthalpy (Δ*H*_TS-N(*T*)_ > 0) leading to Δ*H*_TS-N(*T*)_ − *T*Δ*S*_TS-N(*T*)_ > 0. When *T* = *T*_*S*(*α*)_, Δ*S*_TS-N(*T*)_ = Δ*H*_TS-N(*T*)_ = 0 ⇒ Δ*G*_TS-N(*T*)_ = 0, and *k*_*u*(*T*)_ = *k*^0^. For *T*_*S*(α)_ < *T* < *T*_*H*(TS-N)_, *N* ⇌ [*TS*] is enthalpically favourable (Δ*H*_TS-N(*T*)_ < 0) but is more than offset by the unfavourable negentropy (*T*Δ*S*_TS-N(*T*)_ < 0) leading to Δ*G*_TS-N(*T*)_ > 0. When *T* = *T_H_*_(TS-N)_, Δ*H*_TS-N(*T*)_ = 0 for the second time, Δ*G*_TS-N(*T*)_ is purely due to the negentropy (Δ*G*_TS-N(*T*)_ = −*T*Δ*S*_TS-N(*T*)_ 0), and *k*_*u*(*T*)_ is a minimum. For *T*_*H*(TS-N)_ < *T* < *T_S_*, *N* ⇌ [*TS*] is entropically and enthalpically unfavourable (Δ*H*_TS-N(*T*)_ > 0 and *T*Δ*S*_TS-N(*T*)_ < 0) leading to Δ*G*_TS-N(*T*)_ > 0. When *T* = *T_S_*, Δ*S*_TS-N(*T*)_ = 0 for the second time, and Δ*G*_TS-N(*T*)_ is a minimum and purely enthalpic (Δ*G*_TS-N(*T*)_ = Δ*H*_TS-N(*T*)_ 0). **(B)** For *T_S_* < *T* < *T*_*S*(ω)_, *N* ⇌ [*TS*] is entropically favourable (Δ*H*_TS-N(*T*)_ > 0) but is more than offset by the endothermic enthalpy (Δ*H*_TS-N(*T*)_ > 0) leading to a positive Δ*G*_TS-N(*T*)_. When *T* = *T*_*S*(ω)_, Δ*S*_TS-N(*T*)_ = Δ*H*_TS-N(*T*)_ = 0 for the third and the final time, Δ*G*_TS-N(*T*)_ = 0 for the second and final time, and *k*_*u*(*T*)_ = *k*^0^. For *T*_*S*(ω)_ < *T* ≤ *T*_ω_, *N* ⇌ [*TS* is enthalpically favourable (Δ*H*_TS-N(*T*)_ < 0) but is more than offset by the unfavourable negentropy (*T*Δ*S*_TS-N(*T*)_ < 0), leading to Δ*G*_TS-N(*T*)_ > 0 and *k*_*u*(*T*)_ < *k*^0^.

**Figure 13-figure supplement 2.**
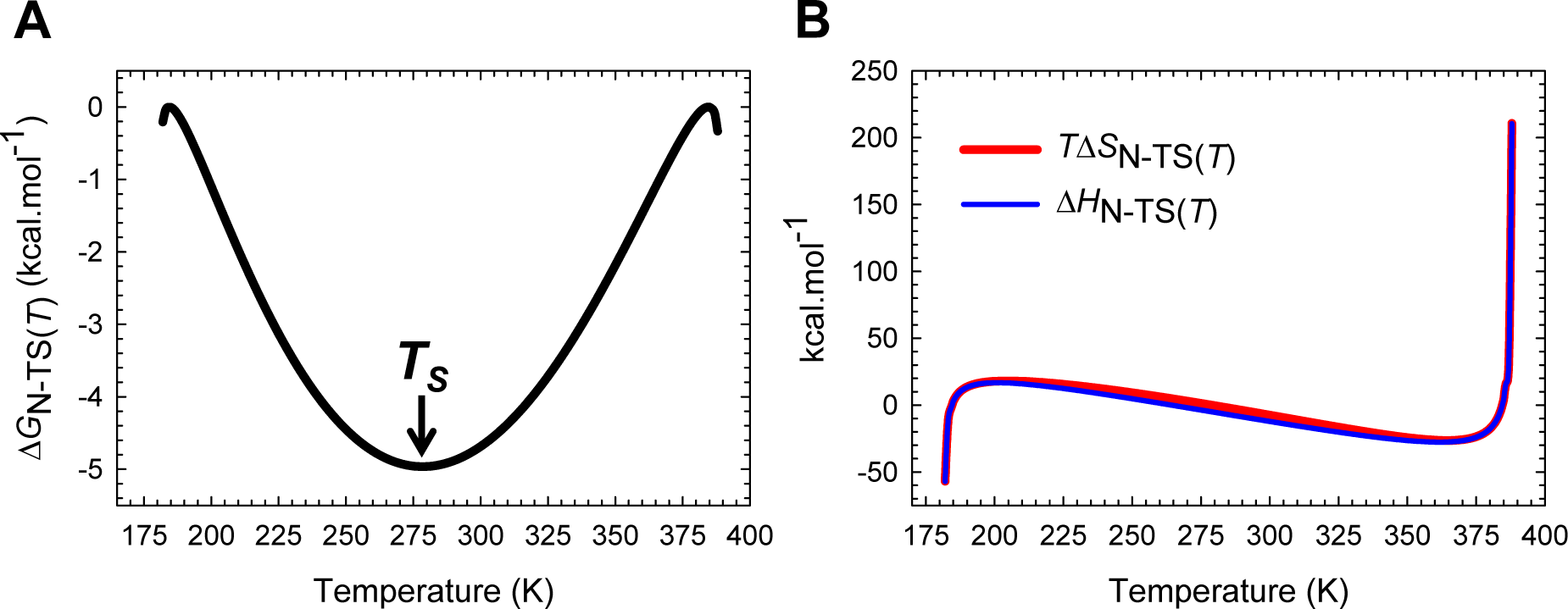
Entropy-enthalpy compensation for the partial folding reaction [*TS*] ⇌ *N*. Despite large changes in Δ*H*_N-TS(*T*)_, Δ*G*_N-TS(*T*)_ which is a minimum at *T_S_*, varies only by ∽5 kcal.mol^-1^ due to compensating changes in Δ*S*_N-TS(*T*)_. See the appropriately scaled figure supplement for description.

**Figure 13−figure supplement 3.**
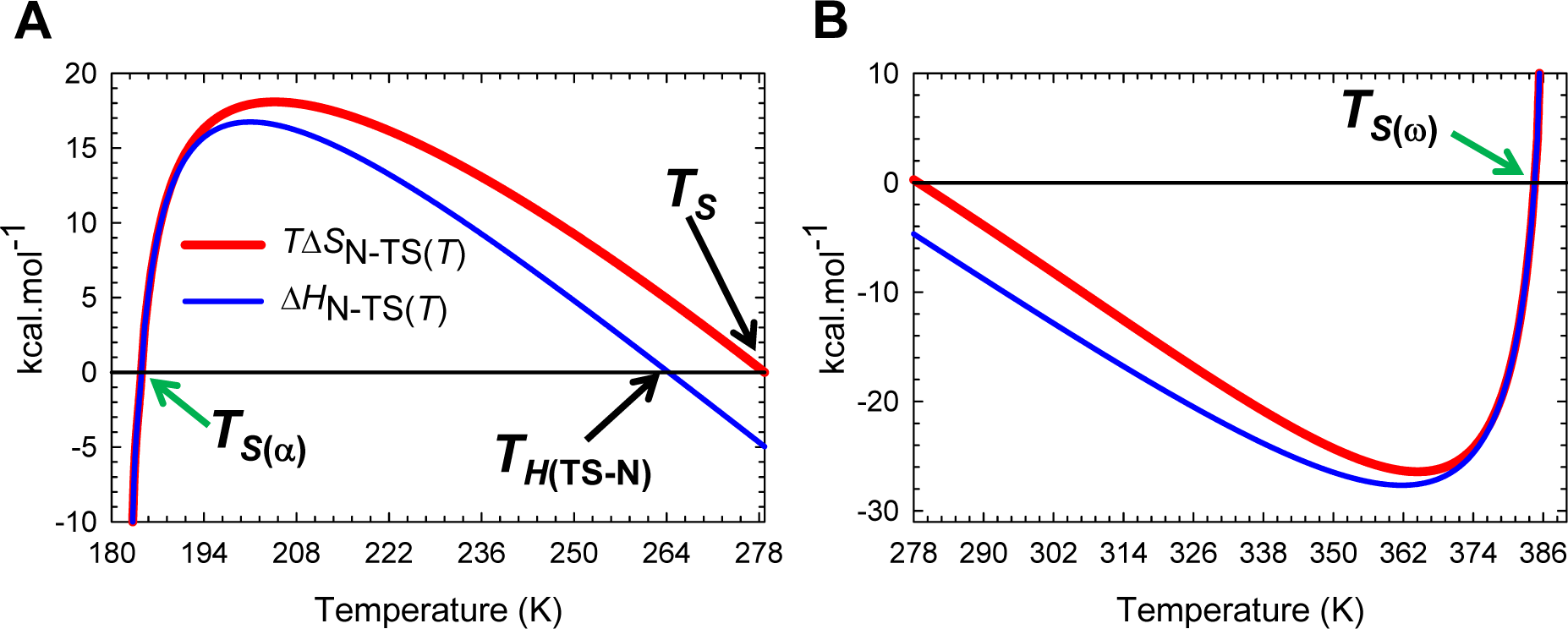
Deconvolution of the change in Gibbs energy for the partial folding reaction [*TS*] ⇌ *N*. These are appropriately scaled split views of **Figure 13−figure supplement 2B**. **(A)** For *T*_α_ ≤ *T* < *T*_*S*(α)_, [*TS*] ⇌ *N* is entropically disfavoured (*T*Δ*S*_N-TS(*T*)_ < 0) but is more than compensated by the exothermic enthalpy (Δ*H*_N-TS(*T*)_ < 0), leading to Δ*G*_N-TS(*T*)_ < 0. When *T* = *T*_*S*(α)_, Δ*S*_N-TS(*T*)_ = Δ*H*_N-TS(*T*)_ = Δ*G*_N-TS(*T*)_ = 0, and the net flux of the conformers from the TSE to the NSE is zero. For *T*_*S*(α)_ < *T* < *T*_*H*(TS-N)_, [*TS*] ⇌ *N* is enthalpically unfavourable (Δ*H*_N-TS(*T*)_ > 0) but is more than compensated by entropy (*T*Δ*S*_N-TS(*T*)_ > 0) leading to Δ*G*_N-TS(*T*)_ < 0. When *T* = *T*_*H*(TS-N)_, the net flux from the TSE to the NSE is driven purely by the favourable change in entropy (Δ*G*_N-TS(*T*)_ = –*T*Δ*S*_N-TS(*T*)_ < 0). For *T*_*H*(TS-N)_ < *T* < *T*_*S*_, the net flux of the conformers from the TSE to the NSE is entropically and enthalpically favourable (Δ*H*_N-TS(*T*)_ < 0 and *T*Δ*S*_N-TS(*T*)_ > 0) leading to Δ*G*_N-TS(*T*)_ < 0. When *T* = *T_S_*, the net flux is driven purely by the exothermic change in enthalpy (Δ*G*_N-TS(*T*)_ = Δ*H*_N-TS(*T*)_ < 0). **(B)** For *T_S_* < *T* < *T*_*S*(ω)_,[*TS*] ⇌ *N* is entropically unfavourable (*T*Δ*S*_N-TS(*T*)_ < 0) but is more than compensated by the exothermic enthalpy (Δ*H*_N-TS(*T*)_ < 0) leading to Δ*G*_N-TS(*T*)_ < 0. When *T* = *T*_*S*(ω)_, Δ*S*_N-TS(*T*)_ = Δ*H*_N-TS(*T*)_ = Δ*G*_N-TS(*T*)_ = 0, and the net flux of the conformers from the TSE to the NSE is zero. For *T*_*S*(ω)_< *T* ≤ *T*_ω_, [*TS*] ⇌ *N* is enthalpically unfavourable (Δ*H*_N-TS(*T*)_ > 0) but is more than compensated by the favourable change in entropy (*T*Δ*S*_N-TS(*T*)_ > 0), leading to Δ*G*_N-TS(*T*)_ < 0.

**Figure 14−figure supplement 1.**
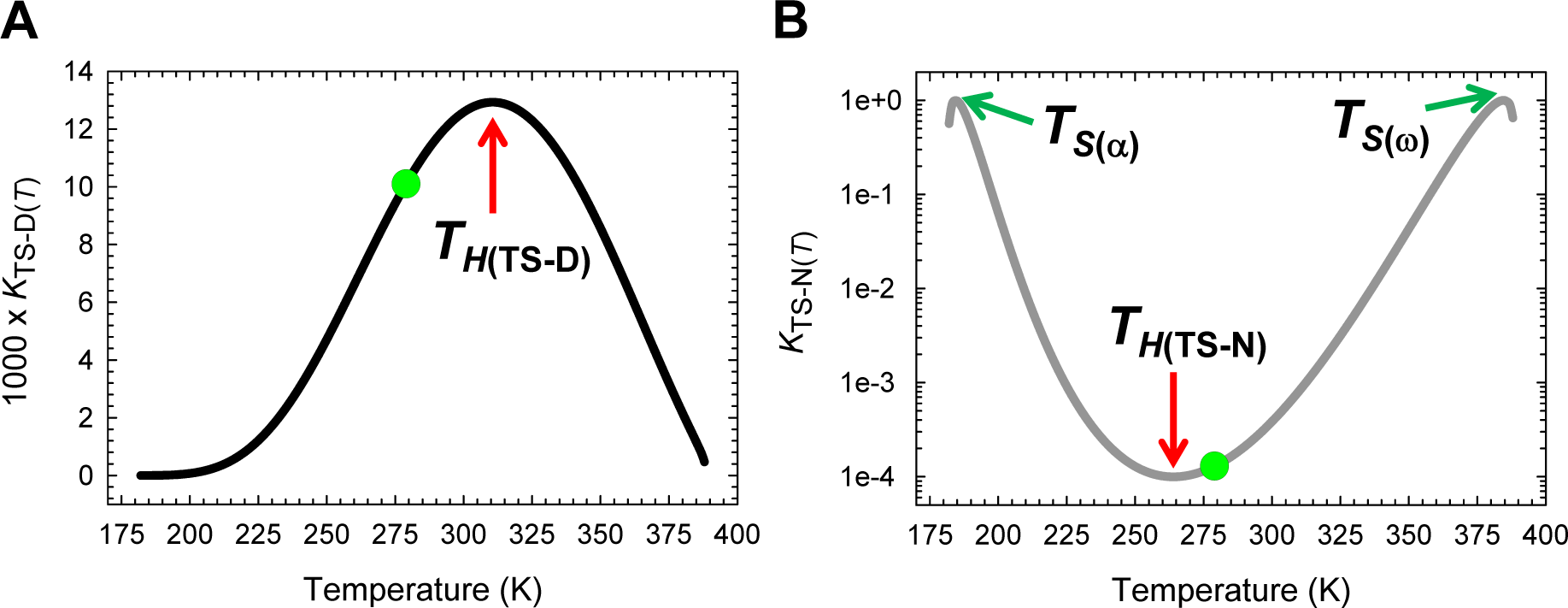
Temperature-dependence of *K*_TS-D(*T*)_ and *K*_TS-N(*T*)._ **(A)** Temperature-dependence of *K*_TS-D(*T*)_ = [*TS*]/[*D*] for the partial folding reaction *D* ⇌ [*TS*]. *K*_TS-D(*T*)_ is a maximum not when Δ*G*_TS-D(*T*)_ is a minimum (green circle) but when the Massieu-Planck activation potential for folding, Δ*G*_TS-D(*T*)_/*T*, is a minimum, and occurs precisely when *T* = *T*_*H*(TS-D)_. The slope of this curve is given by *K*_TS-D(*T*)_ Δ*H*_TS-D(*T*)_/*RT*^2^. **(B)** Temperature-dependence of *K*_TS-N(*T*)_ = [*TS*]/[*N*] for the partial unfolding reaction *N* ⇌ [*TS*]. *K*_TS-N(*T*)_ is a minimum not when Δ*G*_TS-N(*T*)_ is a maximum (green circle) but when the Massieu-Planck activation potential for unfolding, Δ*G*_TS-N(*T*)_/*T*, is a maximum, and occurs precisely when *T* = *T*_*H*(TS-N)_. The slope of this curve is given by *K*_TS-N(*T*)_ Δ*H*_TS-N(*T*)_/*RT*^2^. Note that *K*_TS-N(*T*)_ is unity at *T*_*S*(α)_ and *T*_*S*(ω)_. It is not possible to capture the minimum of *K*_TS-N(*T*)_ on a linear scale; hence the ordinate is shown on a log scale (base 10). The green circles represent the temperature *T_S_* at which Δ*G*_D-N(*T*)_ and Δ*G*_TS-N(*T*)_ are both a maximum, Δ*G*_TS-D(*T*)_ is a minimum, and the absolute entropies of the DSE, the TSE and the NSE are identical.

**Figure 14−figure supplement 2.**
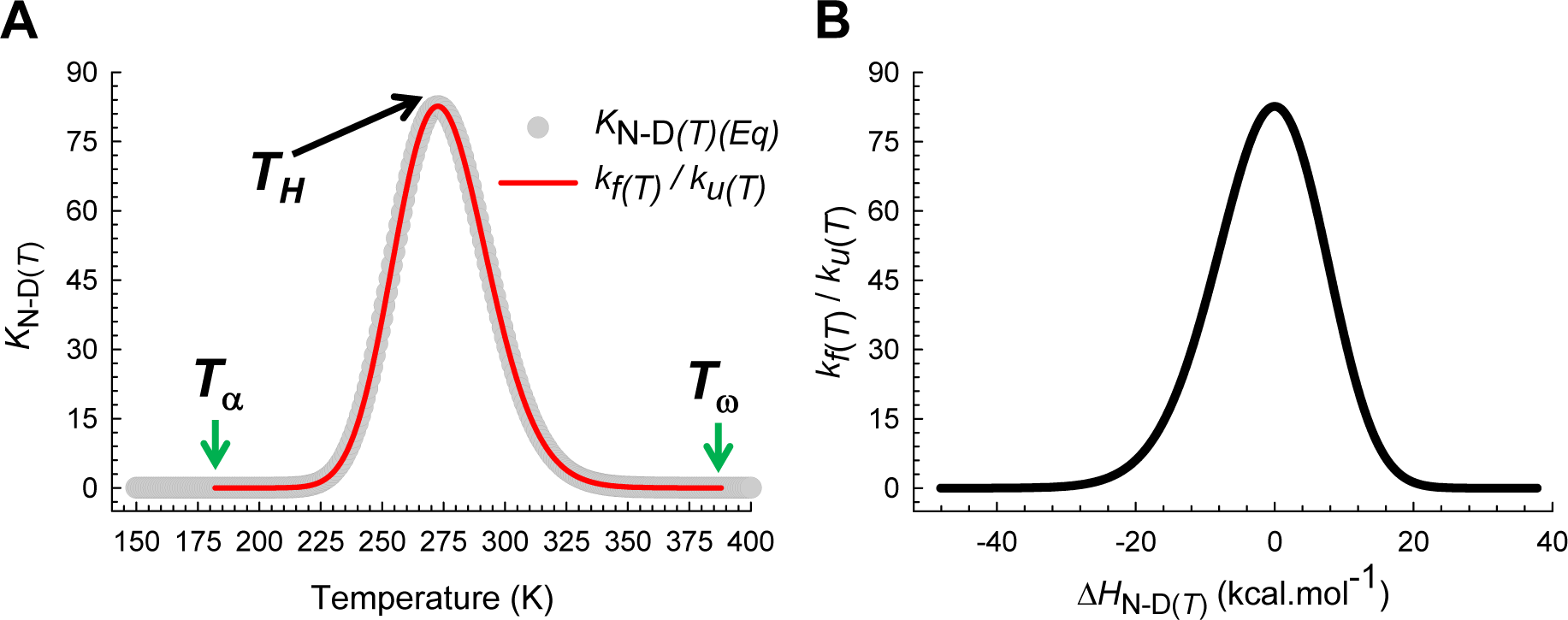
Temperature-dependence of the equilibrium constant for the reaction *D* ⇌ *N*. **(A)** An overlay of the ratio of the rate constants for folding and unfolding and the equilibrium constant derived from the Gibbs energy of folding at equilibrium. The curve fits to Boltzmann distribution and is a maximum at *T_H_*. The slope of this curve is given by *K*_N-D(*T*)_ Δ*H*_N-D(*T*)_/*RT*^2^. Although the value of Δ*H*_D-N(*T*)_ can be calculated for any temperature above absolute zero using Eq. (A1), it has physical meaning only for *T*_α_ ≤ *T* ≤ *T*_ω_. This applies to Δ*S*_D-N(*T*)_ and Δ*G*_D-N(*T*)_ as well (Eqs. (A2) and (A3)). **(B)** The solubility of the NSE as compared to the DSE is the greatest when the net flux of the conformers from the DSE to the NSE is driven purely by the difference in entropy between these two reaction-states. The slope of this curve is given by *K*_N-D(*T*)_ Δ*H*_N-D(*T*)_/Δ*C*_*p*N-D_*RT*^2^

**Figure 14−figure supplement 3.**
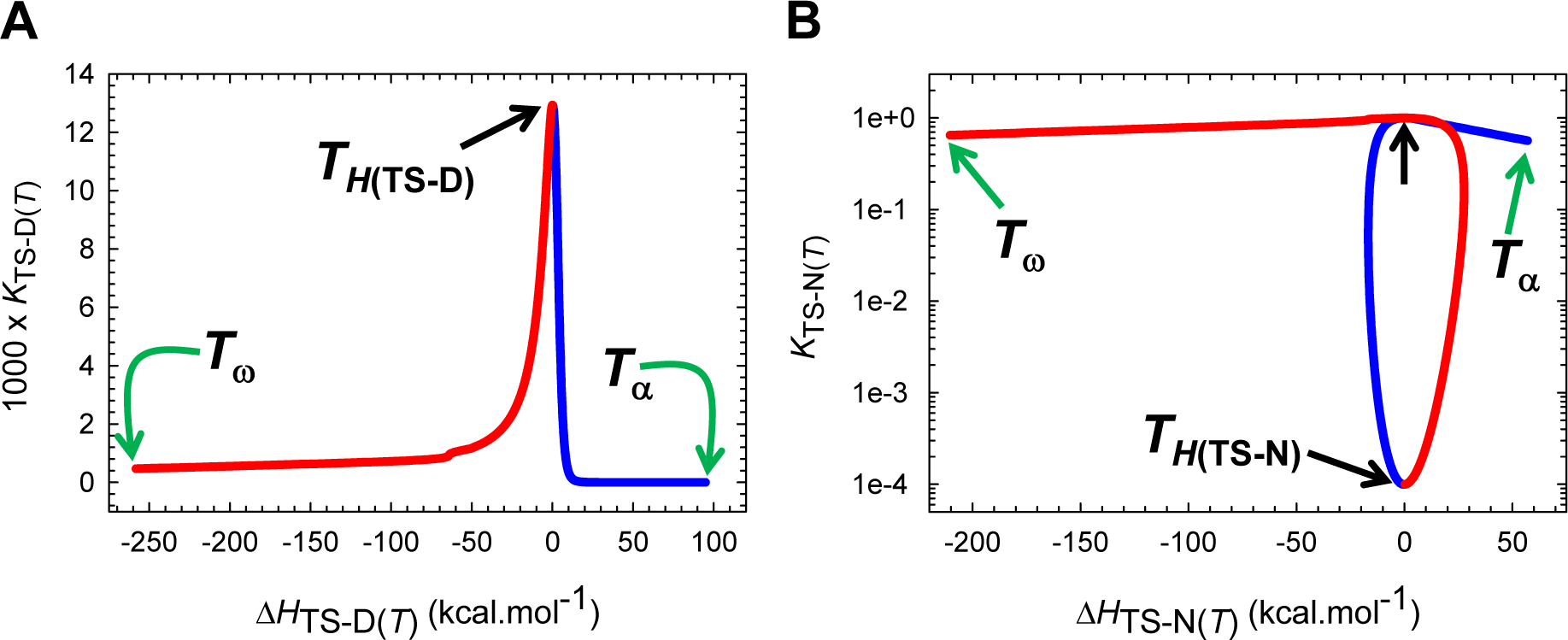
The solubility of the TSE relative to the DSE and the NSE across a broad temperature regime. **(A)** The solubility of the TSE as compared to the DSE is the greatest when Δ*H*_TS-D(*T*)_ = 0, or equivalently, when the Gibbs barrier to folding is purely entropic. The slope of this curve is given by *K*_TS-D(*T*)_ Δ*H*_TS-D(*T*)_/Δ*C*_*p*TS-D(*T*)_*RT*^2^. The blue and red sections of the curve represent the temperature regimes *T*_α_ ≤ *T* ≤ *T*_*H*(TS-D)_ and *T*_*H*(TS-D)_ ≤ *T* ≤ *T*_ω_, respectively. **(B)** The solubility of the TSE as compared to the NSE is the least when Δ*H*_TS-N(*T*)_= 0 and when the Gibbs barrier to unfolding is purely entropic. The slope of this curve is given by *K*_TS-N(*T*)_ Δ*H*_TS-N(*T*)_/Δ*C*_*p*TS-N(*T*)_*RT*^2^. The point where the solubility of the TSE is identical to that of the NSE is indicated by the unlabelled black pointer, and described earlier, occurs precisely at *T*_*S*(α)_ and *T*_*S*(ω)_. The blue and red sections of the curve represent the temperature regimes *T*_α_ ≤ *T* ≤ *T*_*H*(TS-N)_ and *T*_*H*(TS-N)_ ≤ *T* ≤ *T*_ω_, respectively. Note that the ordinate is on a log scale (base 10).

**Figure 14−figure supplement 4.**
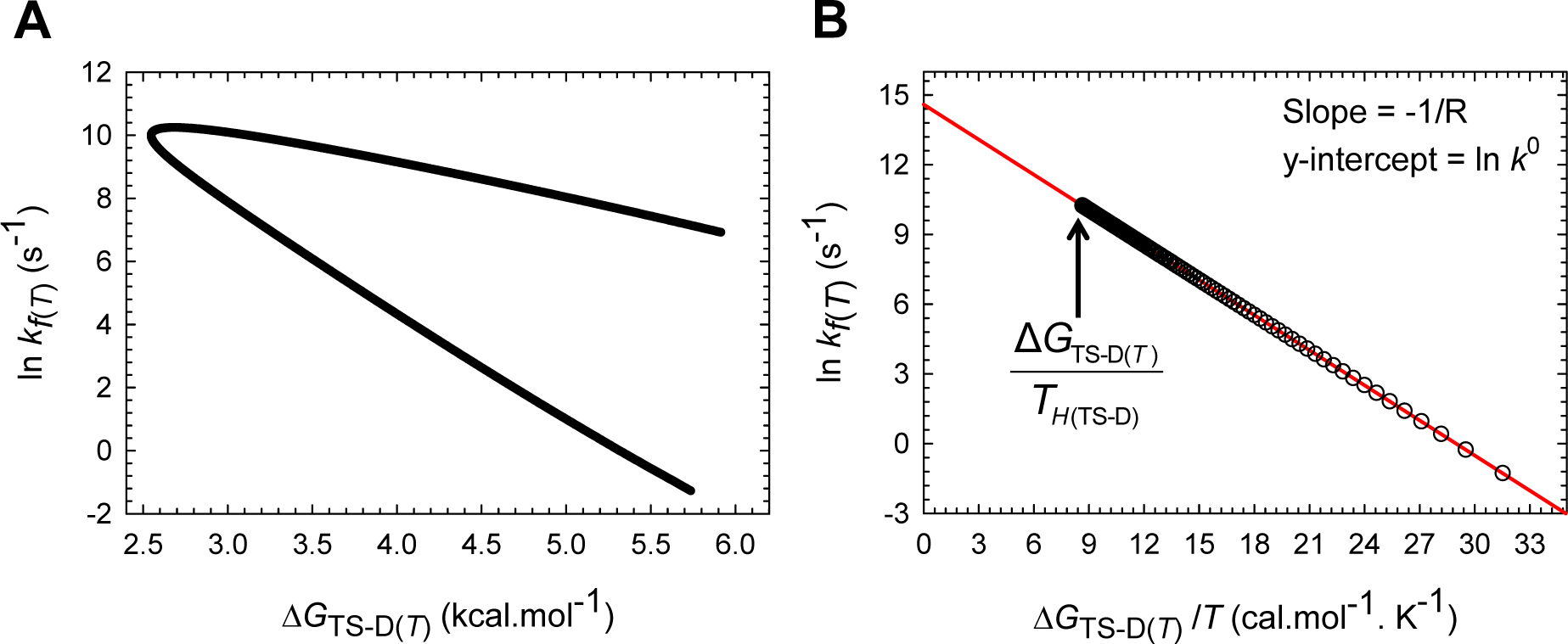
The natural logarithm of *k*_*f*(*T*)_ is linearly dependent on the Massieu-Planck activation potential for folding. **(A)** The natural logarithm of *k*_*f*(*T*)_ has a complex dependence on the Gibbs barrier to folding when explored over a large temperature range. The slope of this curve is given by −Δ*H*_TS-D(*T*)_/Δ*S*_TS-D(*T*)_*RT*^2^. **(B)** The natural logarithm of *k*_*f*(*T*)_ decreases linearly with an increase in the Massieu-Planck activation potential for folding, with the magnitude of the negative slope being given by the reciprocal of the gas constant. The *y*-intercept at zero Massieu-Planck potential yields the value of the prefactor. Naturally, *k*_*f*(*T*)_ is a maximum when the magnitude of the Massieu-Planck function for folding is a minimum, and this occurs precisely at *T*_*H*(TS-D)._

**Figure 14−figure supplement 5.**
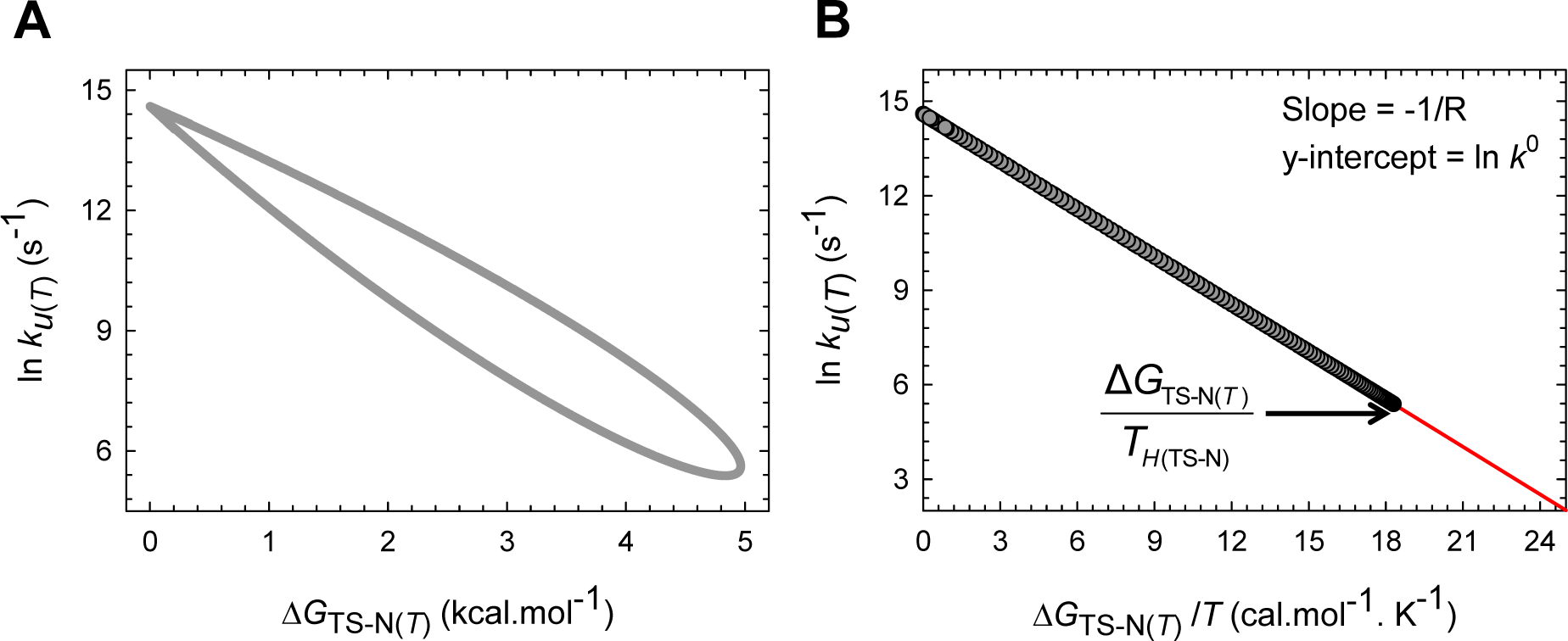
The natural logarithm of *k*_*u*(*T*)_ is linearly dependent on the Massieu-Planck activation potential for unfolding. **(A)** The natural logarithm of *k*_*u*(*T*)_ has a complex dependence on the Gibbs barrier to unfolding when explored over a large temperature range. The slope of this curve is given by −Δ*H*_TS-N(*T*)_/Δ*S*_TS-N(*T*)_*RT*^2^. **(B)** The natural logarithm of *k*_*u*(*T*)_ decreases linearly with an increase in the Massieu-Planck activation potential for unfolding, with the magnitude of the negative slope being given by the reciprocal of the gas constant. The *y*-intercept at zero Massieu-Planck potential yields the value of the prefactor. Naturally, *k*_*u*(*T*)_ is a minimum when the magnitude of the Massieu-Planck function for unfolding is a maximum, and this occurs precisely at *T*_*H*(TS-N)_. The reason why the data points for the unfolding rate constants extend all the way to the intercept is because the Gibbs barrier to unfolding becomes zero at *T*_*S*(α)_ and *T*_*S*(ω)_.

**Figure 15−figure supplement 1.**
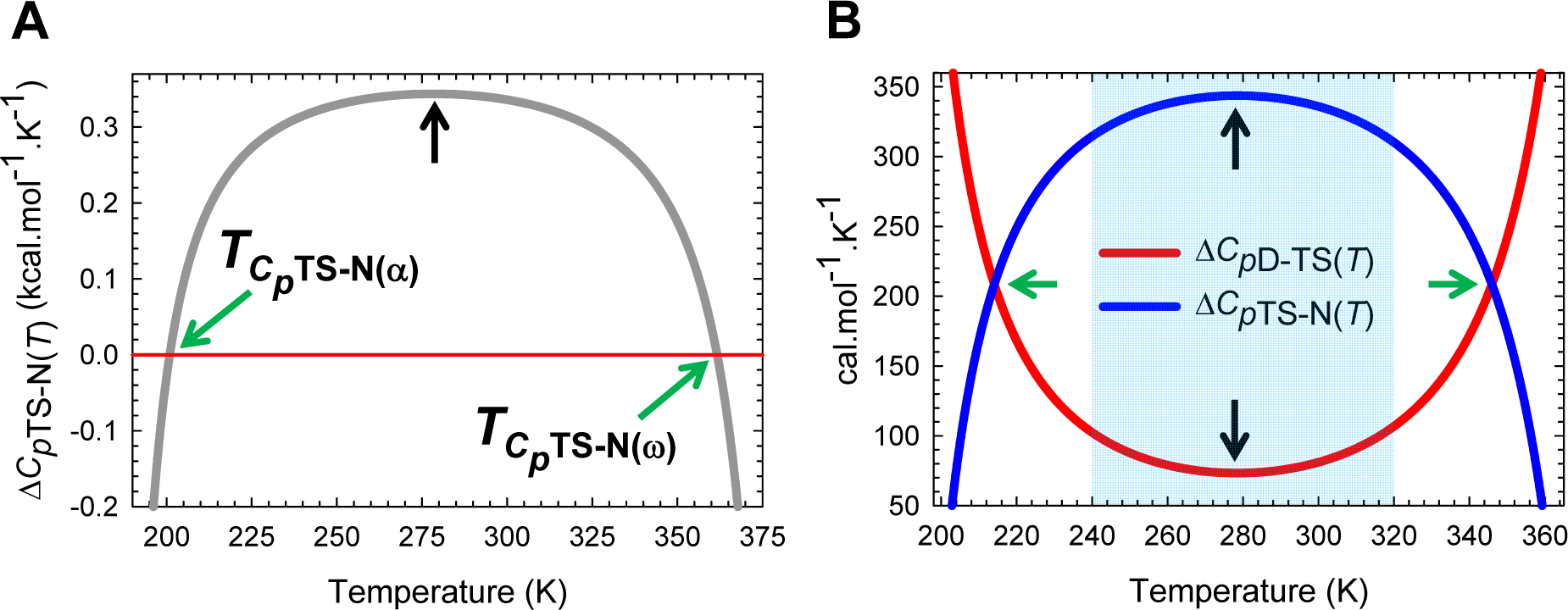
Appropriately scaled Δ*C*_*p*D-TS(*T*)_ and Δ*C*_*p*TS-N(*T*)_ functions to showcase their features. **(A)** Δ*C*_*p*TS-N(*T*)_ which is a maximum and positive at *T_S_*, decreases with any deviation in temperature from *T_S_*, is zero at *T*_*C_p_*TS-N(α)_ and *T*_*C_p_*TS-N(ω)_, and negative for *T*_α_ ≤ *T* < *T*_*C_p_*TS-N(α)_ and *T*_*C_p_*TS-N(ω)_ < *T* ≤ *T*_ω_. **(B)** At the temperatures where the Δ*C*_*p*D-TS(*T*)_ and Δ*C*_*p*TS-N(*T*)_ functions intersect (214.1K and 345.9 K), the absolute heat capacity of the TSE is exactly half the sum of the absolute heat capacities of the DSE and the NSE. The black pointers indicate that the extrema of Δ*C*_*p*D-TS(*T*)_ and Δ*C*_*p*TS-N(*T*)_ functions, while the green pointers indicate their intersection. Inspection shows that Δ*C*_*p*TS-N(*T*)_ > Δ*C*_*p*D-TS(*T*)_ for 240 K < *T* < 320 K (shaded region), and is approximately five fold greater than Δ*C*_*p*TS-N(*T*)_ at *T_S_* (343.7/73.3 = ∽ 4.7) despite ∽30% and ∽70% of the total change in SASA for the unfolding reaction *N* ⇌ *D*, occurring in the partial unfolding reactions *N* ⇌ [*TS*] and [*TS*] ⇌ *D*, respectively (**Figure 2−figure supplement 1**).

**Figure 15-figure supplement 2.**
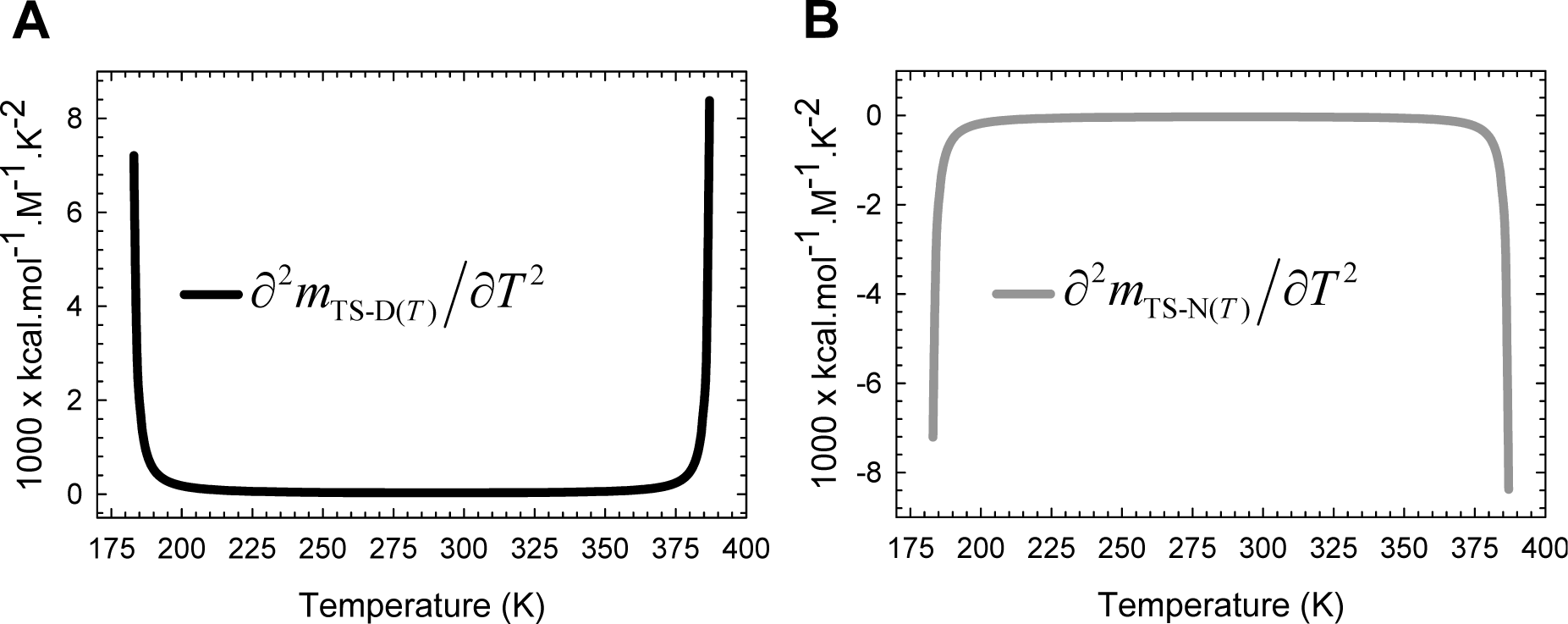
The second derivatives of *m*_TS-D(*T*)_ and *m*_TS-N(*T*)_ with respect to temperature. **(A)** The second derivative of *m*_TS-D(*T*)_ according to Eq. (A9). **(B)** The second derivative of *m*_TS-N(*T*)_ according to Eq. (A10). The sole intent of these figures is to demonstrate that the gross features of the temperature-dependence of the heat capacity functions arise primarily from the second derivatives of the temperature-dependent shift in the position of the TSE relative to the vertices of the DSE or the NSE Gibbs parabolas along the RC. See **Figure 15−figure supplement 3** for the location of the extrema of these two functions.

**Figure 15-figure supplement 3.**
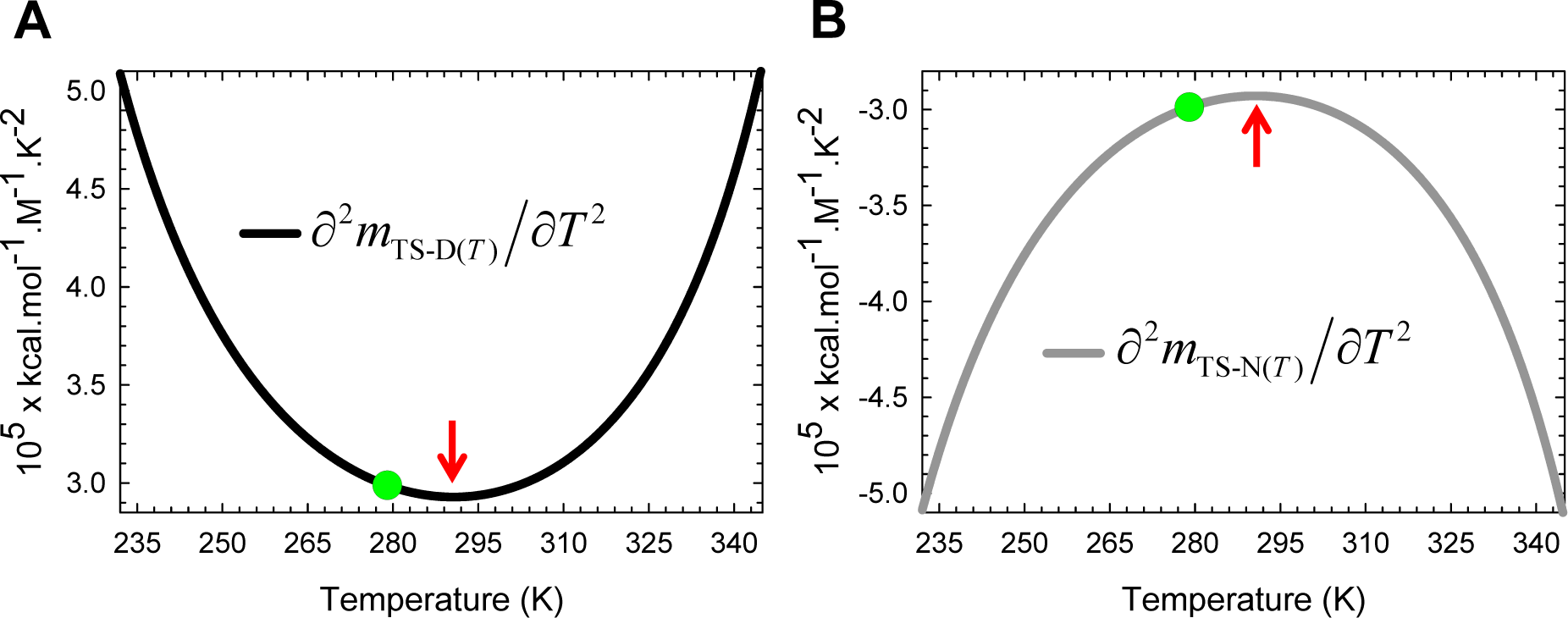
The extrema of the second derivatives of *m*_TS-D(*T*)_ and *m*_TS-N(*T*)_ with respect to temperature are not at *T_S_*. **(A)** The sole intent of these appropriately scaled figures is to demonstrate that although the gross features of the temperature-dependence of the heat capacity functions arise predominantly from the second derivatives of the temperature-dependent shift in the position of the TSE along the RC, the minimum of ∂^2^*m*_TS-D(*T*)_/∂*T*^2^ and the maximum of ∂^2^*m*_TS-N(*T*)_/∂*T*^2^ do not occur at *T_S_* (green circles), and is apparent from comparison of Eqs. (12), (13), (A9) and (A10).

**Figure 16-figure supplement 1.**
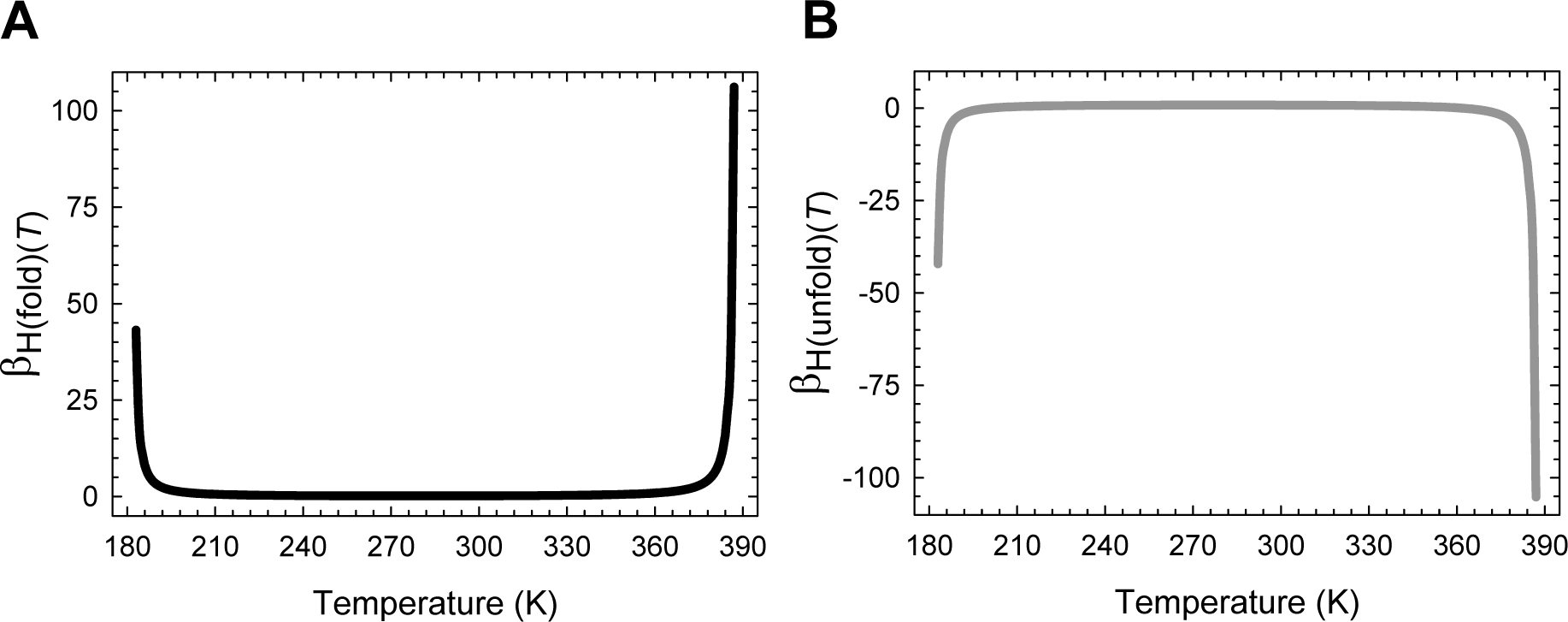
Temperature-dependence of β_H(fold)(*T*)_ and β_H(unfold)(*T*)_. **(A)** Variation in β_H(fold)(*T*)_ with temperature according to Eq. (26). **(B)** Variation in β_H(unfold)(*T*)_ with temperature according to Eq. (27). The location of the extrema is not apparent in these figures. Note that although the algebraic sum of β_H(fold)(*T*)_ and β_H(unfold)(*T*)_ must always be unity for a two-state system, they need not be individually restricted to a canonical range of 0 to 1.

**Figure 17-figure supplement 1.**
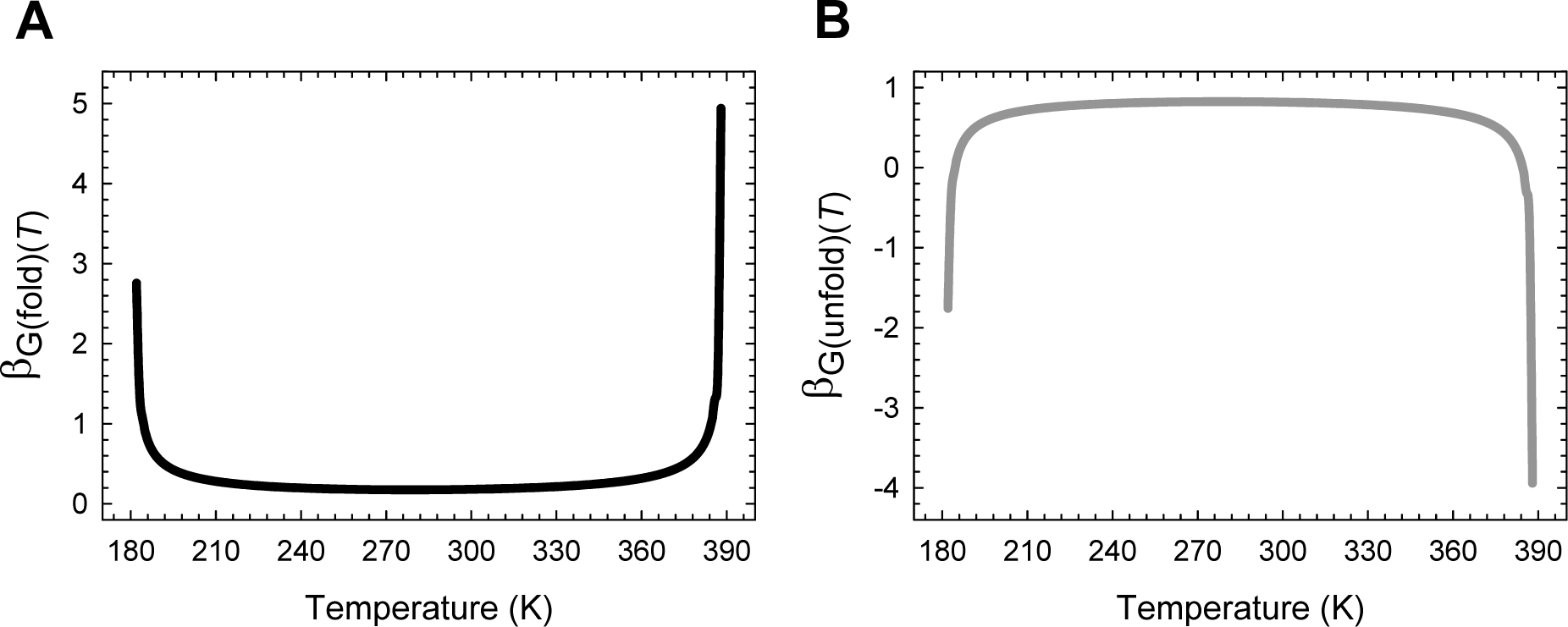
Temperature-dependence of β_G(fold)(*T*)_ and β_G(unfold)(*T*)_. **(A)** Variation in β_G(fold)(*T*)_ with temperature according to Eq. (33). **(B)** Variation in β_G(unfold)(*T*)_ with temperature according to Eq. (35). The location of the extrema is not apparent in these figures. Note that although the algebraic sum of β_G(fold)(*T*)_ and β_G(unfold)(*T*)_ must always be unity for a two-state system, they need not be individually restricted to a canonical range of 0 to 1.

**Figure 19-figure supplement 1.**
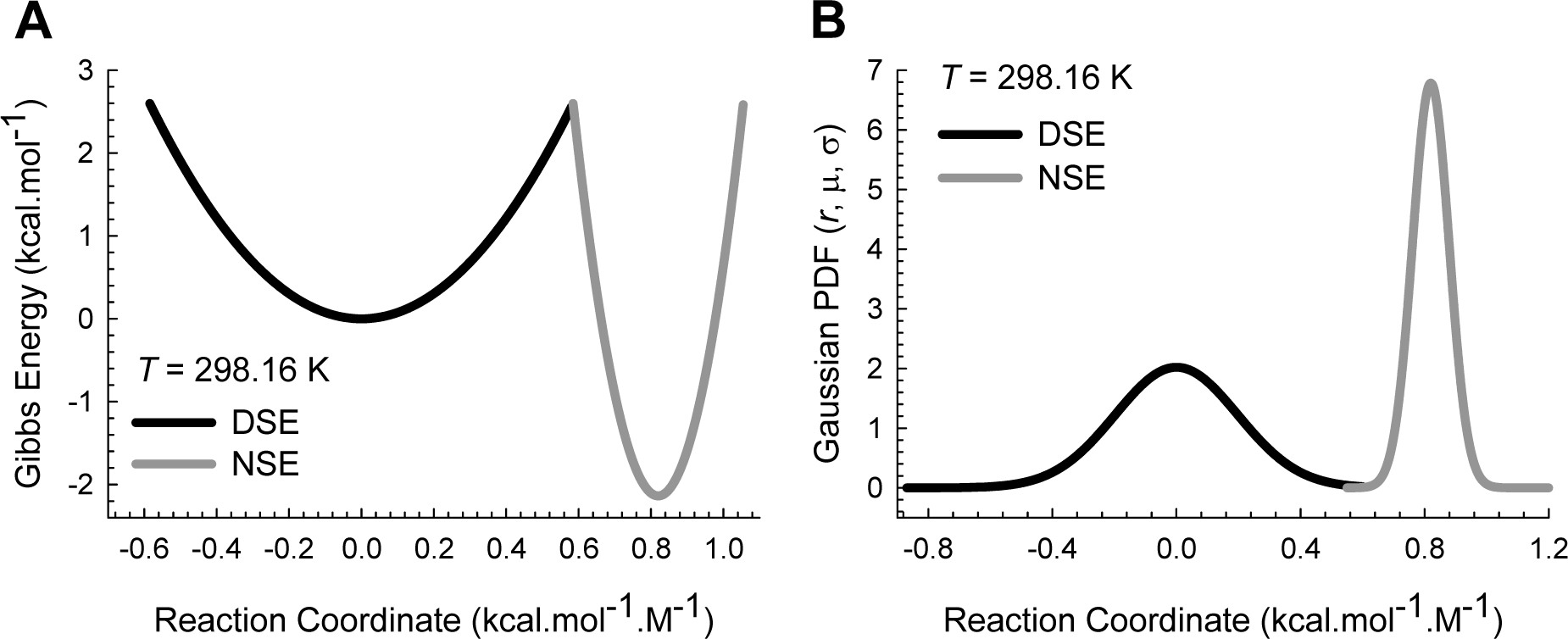
The correspondence between Gibbs parabolas and Gaussian PDFs. **(A)** Parabolic Gibbs energy curves with α = 7.594 M^2^.mol.kcal^-1^ and ω = 85.595 M^2^.mol.kcal^-1^, *m*_D-N_ = 0.82 kcal.mol^-1^.M^-1^ and Δ*G*_D-N(*T*)_ = 2.138 kcal.mol^-1^. The separation between *curve-crossing* and the vertices of the DSE and the NSE-parabolas along the abscissa are 0.5848 kcal.mol^-1^.M^-1^ and 0.2352 kcal.mol^-1^.M^-1^, respectively. The absolute values of Δ*G*_TS-D(*T*)_ and Δ*G*_TS-N(*T*)_ are 2.597 kcal.mol^-1^ and 4.735 kcal.mol^-1^, respectively. The parabolas have been generated as described in the legend for **Figure 3**. **(B)** Gaussian PDFs for the DSE and NSE generated using 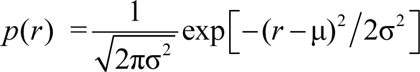, where *r* is any point on the abscissa, μ = 0 kcal.mol^-1^.M^-1^ and σ^2^ = *RT*/2α for the DSE-Gaussian, and μ = 0.82 kcal.mol^-1^.M^-1^ and σ^2^ = *RT*/2ω for the NSE-Gaussian. The units for the Gaussian variances are in kcal^2^.mol^-2^.M^-2^. The relationship between equilibrium stability and the areas enclosed by the DSE and the NSE Gaussians has been addressed in Paper-I.

**Figure 21-figure supplement 1.**
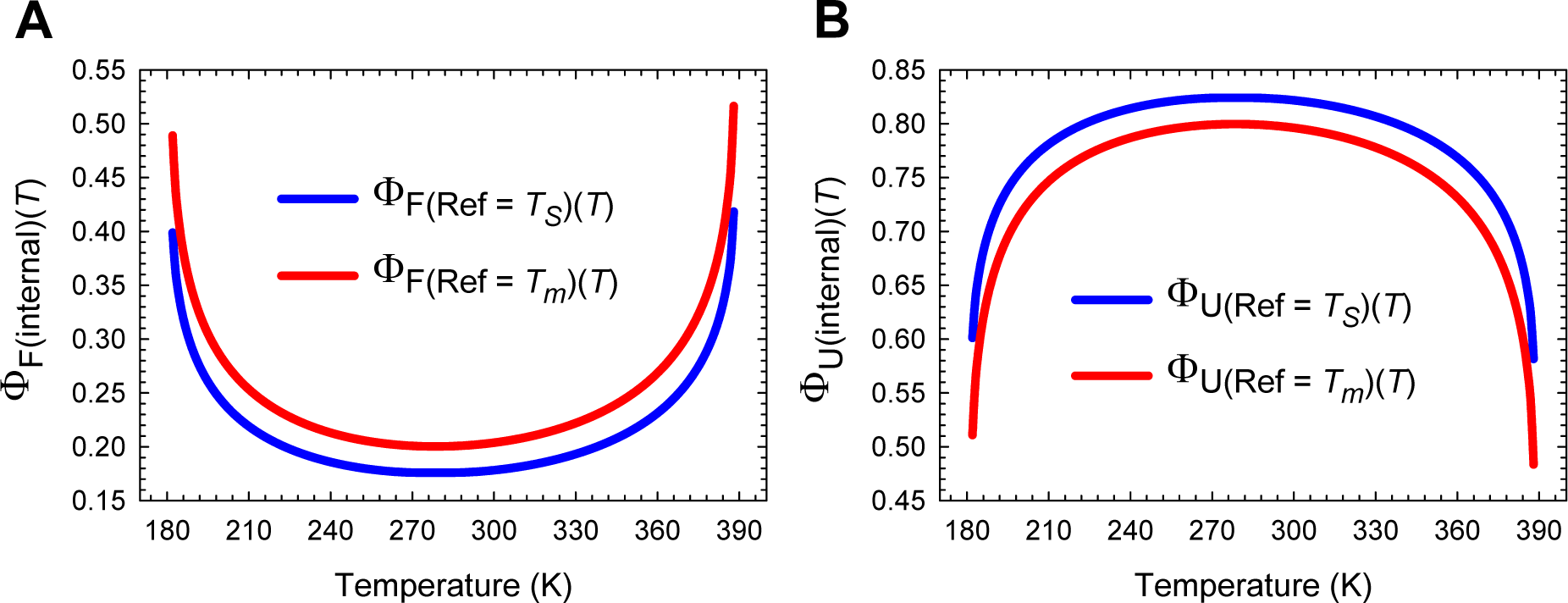
The magnitude of Φ_(internal)(*T*)_ is dependent on the definition of the wild type. **(A)** Φ_F(internal)(*T*)_ calculated using the protein at *T_S_* as the wild type must always be lower than Φ_F(internal)(*T*)_ calculated using protein at *T* ≠ *T_S_* as the wild type. **(B)** Φ_U(internal)(*T*)_ calculated using the protein at *T_S_* as the wild type must always be greater than Φ_U(internal)(*T*)_ calculated using protein at *T* ≠ *T_S_* as the wild type. For the blue curves we have Δ*G*_(wt)_ ≡ Δ*G*_(*T_S_*)_, and for the red curves, we have Δ*G*_(wt)_ ≡ Δ*G*_(*T_m_*)_. This notation applies to both equilibrium and activation energies. The blue curves are undefined (0/0) at *T_S_*, and the red curves are undefined at *T_c_* and *T_m_*. Note that the mathematical stipulation that Φ_F(internal)(*T*)_ + Φ_U(internal)(*T*)_ = 1 for a two-state system is satisfied for both the blue and the red curves for all temperatures.

**Figure 21-figure supplement 2.**
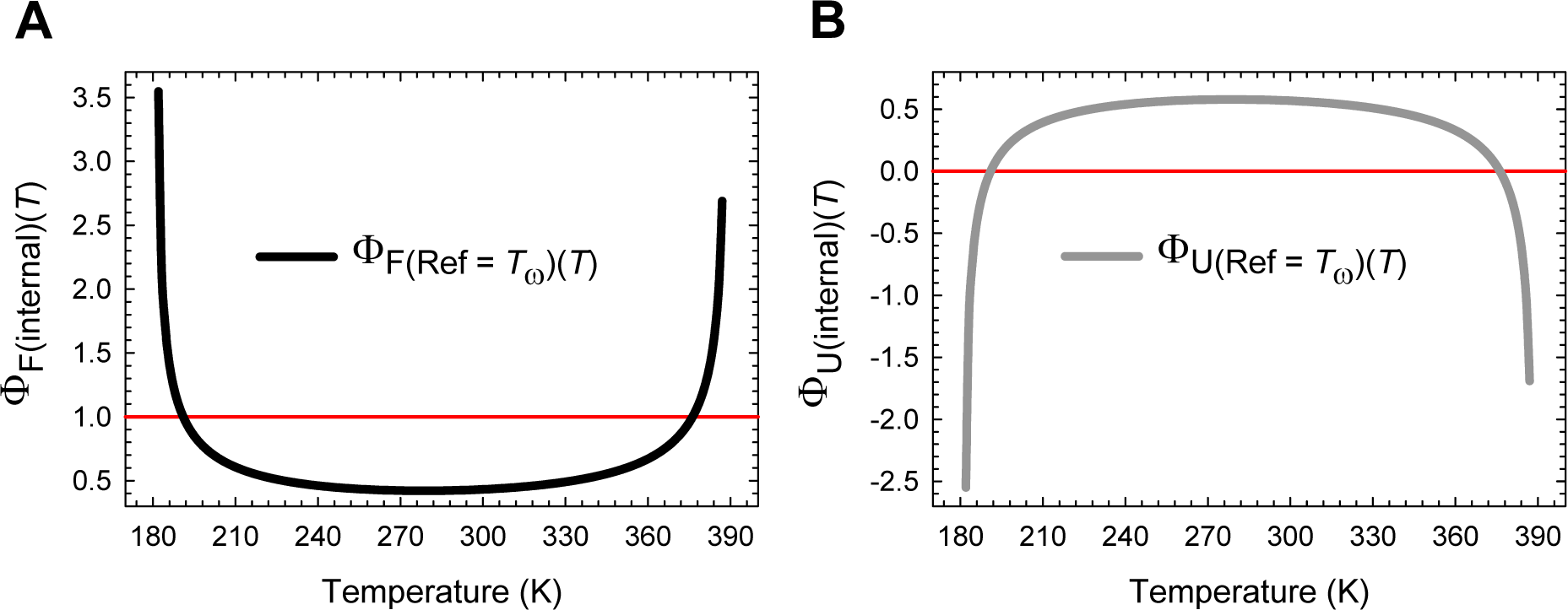
Φ values can be greater than unity, negative or zero depending on the definition of the reference state. **(A)** Φ_F(internal)(*T*)_ calculated by defining the protein at *T*_ω_ as the wild type. **(B)** Φ_U(internal)(*T*)_ calculated by defining the protein at *T*_ω_ as the wild type. Although the mathematical stipulation that Φ_F(internal)(*T*)_ + Φ_U(internal)(*T*)_ = 1 for a two-state system is satisfied for all temperatures, Φ values for folding and unfolding are not restricted to the canonical range of 0 ≤ Φ ≤ 1 when the protein at *T*_ω_ is defined as the reference or the wild type. Note that the curves are undefined at *T*_ω_.

**Figure 21-figure supplement 3.**
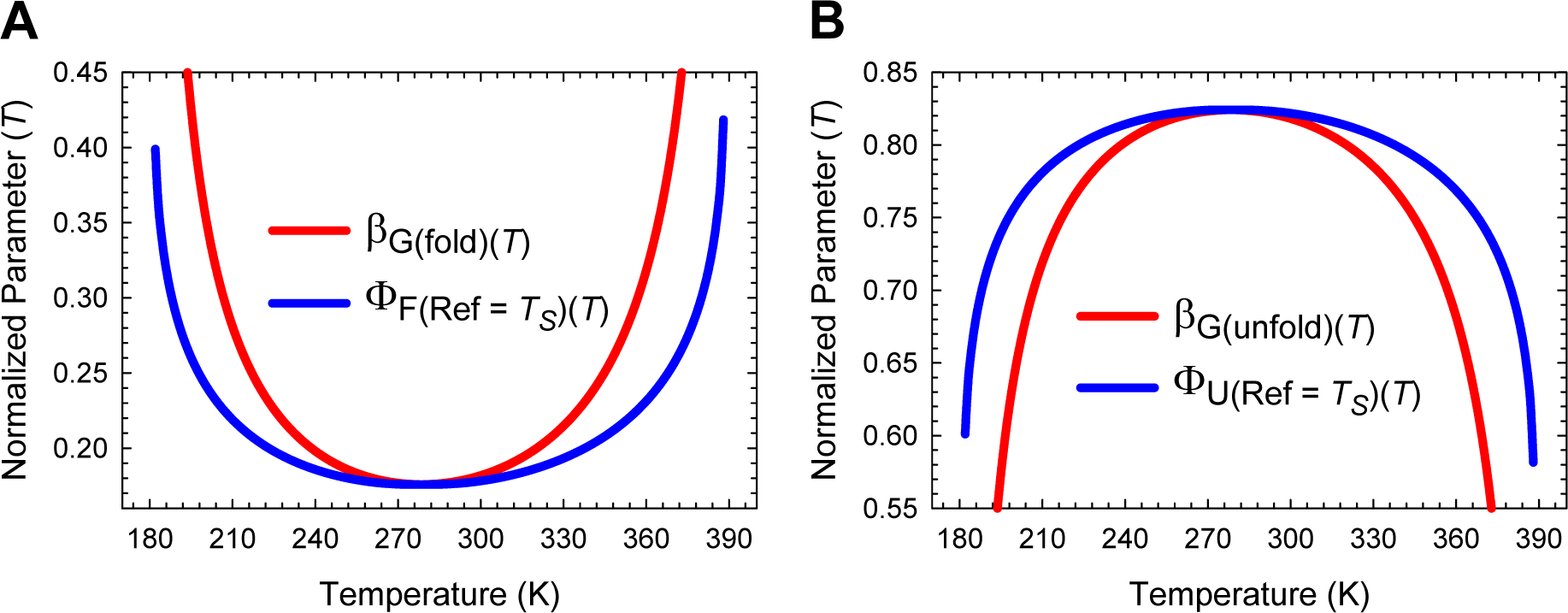
Comparison of Leffler β_G(*T*)_ and Fersht Φ_(internal)(*T*)_ when the protein at *T_S_* is defined as the wild type. **(A)** β_G(fold)(*T*)_ is almost identical to Φ_F(internal)(*T*)_ around the temperature of maximum stability, but as the temperature deviates from *T_S_*, β_G(fold)(*T*)_ increases far more steeply than Φ_F(internal)(*T*)_, such that for *T* ≠*T_S_* we have β_G(fold)(*T*)_ > Φ_F(internal)(*T*)_. **(B)** β_G(unfold)(*T*)_ is almost identical to Φ_U(internal)(*T*)_ around the temperature of maximum stability, but as the temperature deviates from *T_S_*, β_G(unfold)(*T*)_ decreases far more steeply than Φ_U(internal)(*T*)_, such that for *T* ≠*T_S_* we have β_G(unfold)(*T*)_ < Φ_U(internal)(*T*)_.

**Figure 21-figure supplement 4.**
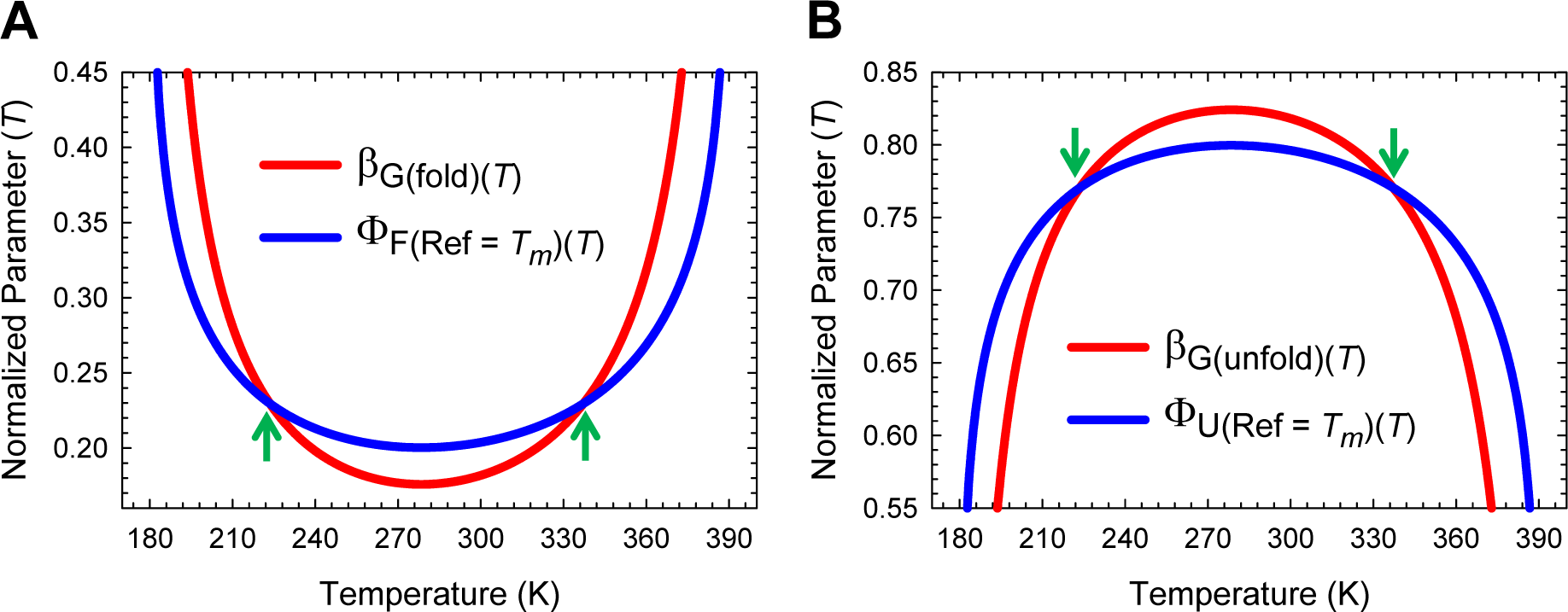
Comparison of the Leffler β_G(*T*)_ and Fersht Φ_(internal)(*T*)_ when the protein at *T_m_* is defined as the wild type. **(A)** β_G(fold)(*T*)_ < Φ_F(internal)(*T*)_ for *T_c_* < *T* < *T_m_* and β_G(fold)(*T*)_ > Φ_F(internal)(*T*)_ for *T* < *T_c_* and *T* > *T_m_*. **(B)** β_G(unfold)(*T*)_ > Φ_U(internal)(*T*)_ for *T_c_* < *T* < *T_m_* and β_G(unfold)(*T*)_ < Φ_U(internal)(*T*)_ for *T* < *T_c_* and *T* > *T_m_*. Note that Φ_F(internal)(*T*)_ and Φ_U(internal)(*T*)_ are undefined for *T_c_* and *T_m_* (green pointers).

**Figure 21-figure supplement 5.**
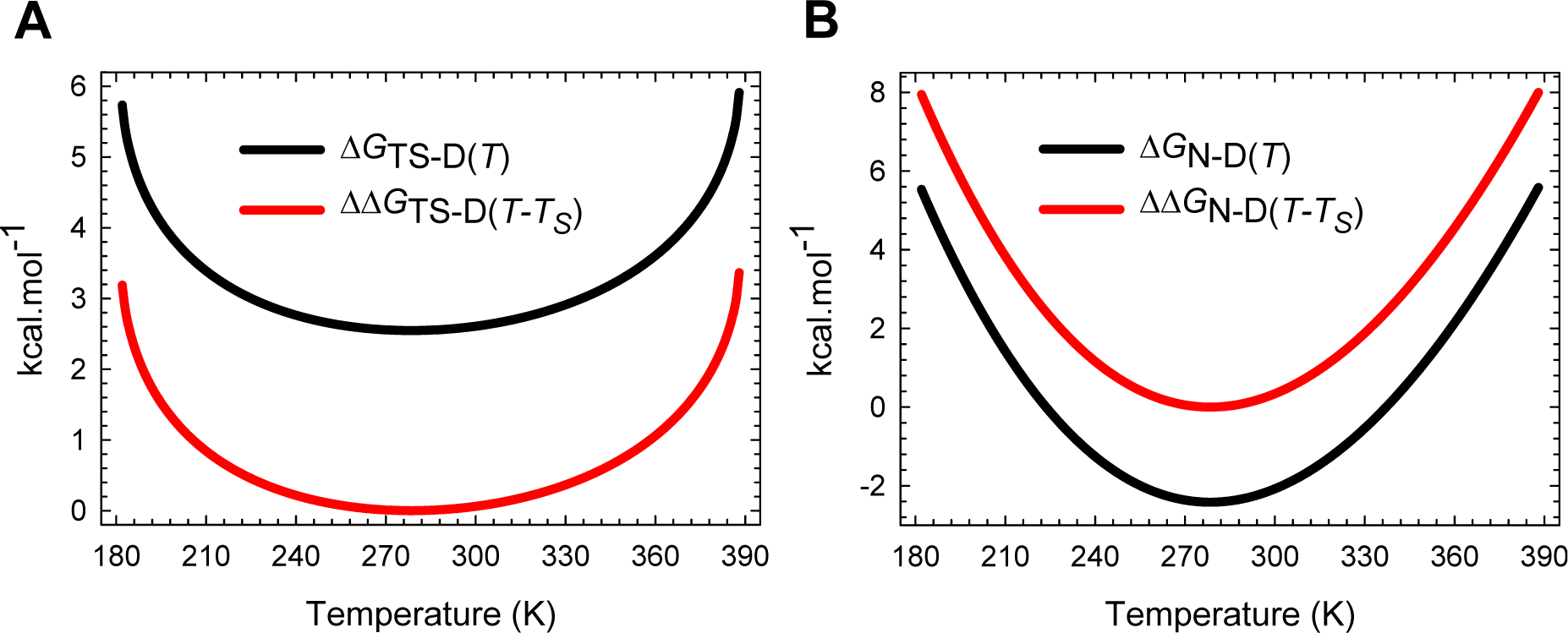
Transformation of Δ*G*TS-D(*T*) and Δ*G*_N-D(*T*)_to generateΦ_F(internal)(*T*)_ when the protein at *T_S_* is defined as the wild type. **(A)** The transformation Δ*G*_TS-D(*T*)_ − Δ*G*_TS-D(*T_S_*)_ (the numerator in Eq. (44)) lowers the Δ*G*_TS-D(*T*)_ function such that the value of ΔΔ*G*_TS-D(*T*-*T_S_*)_ is zero at the reference temperature. **(B)** The transformation Δ*G*_N-D(*T*)_ – Δ*G*_N-D(*T_S_*)_ (the denominator in Eq. (44)) raises the Δ*G*_N-D(*T*)_ function such that the value of ΔΔ*G*_N-D(*T*-*T_S_*)_ is zero at the reference temperature. The unmodified and the transformed curves are shown in black and red, respectively.

**Figure 21-figure supplement 6.**
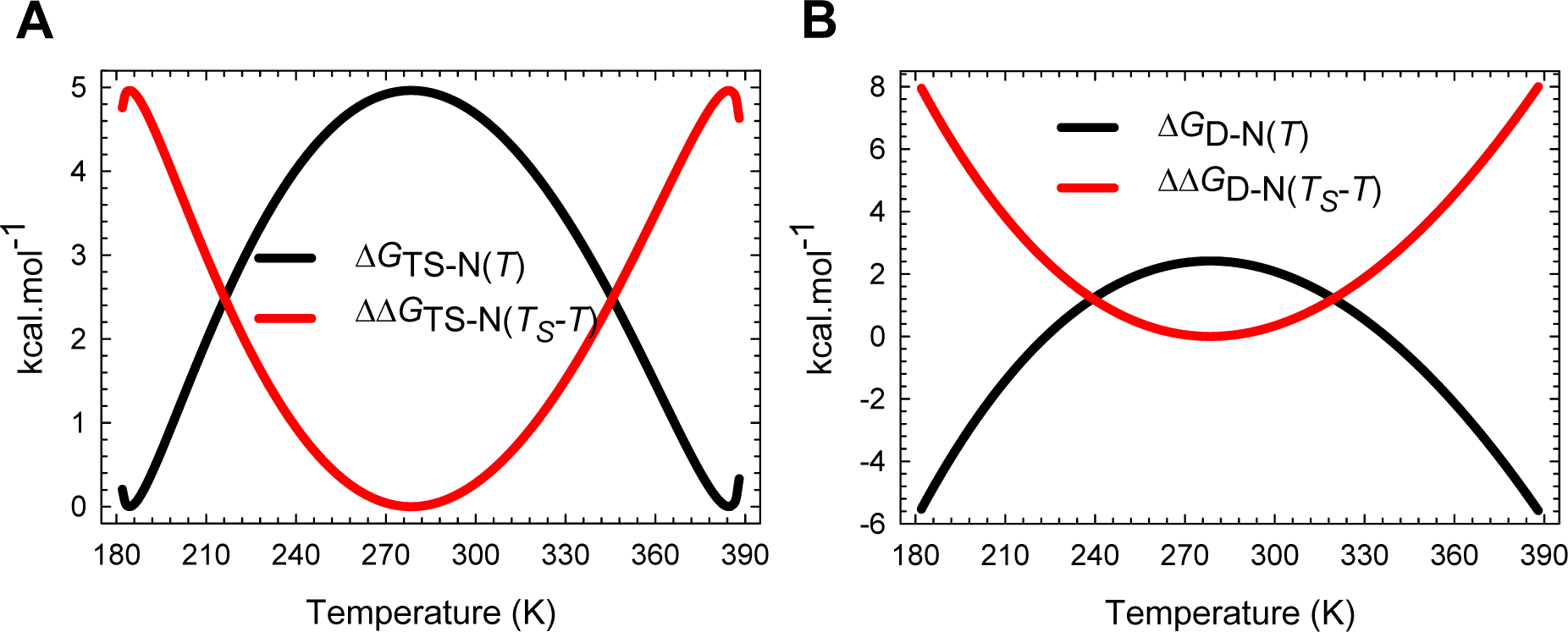
Transformation of Δ*G*_TS-N(*T*)_ and Δ*G*_D-N(*T*)_ to generate Φ_U(internal)(*T*)_ when the protein at *T_S_* is defined as the wild type. **(A)** The transformation Δ*G*_TS-N(*T_S_*)_ − Δ*G*_TS-N(*T*)_ (the numerator in Eq. (45)) flips the Δ*G*_TS-N(*T*)_ function vertically and concomitantly shifts it along the ordinate such that the value of ΔΔ*G*TS-N(*TS*-*T*) at the reference temperature is zero. **(B)** The transformation Δ*G*_D-N(*TS*)_ – Δ*G*_D_N(*T*) (the denominator in Eq. (45)) flips the Δ*G*_D-N(*T*)_ function vertically and concomitantly shifts it along the ordinate such that the value of ΔΔ*G*_D-N(*TS*-*T*)_ at the reference temperature is zero. The unmodified and the transformed curves are shown in black and red, respectively.

**Figure 21-figure supplement 7.**
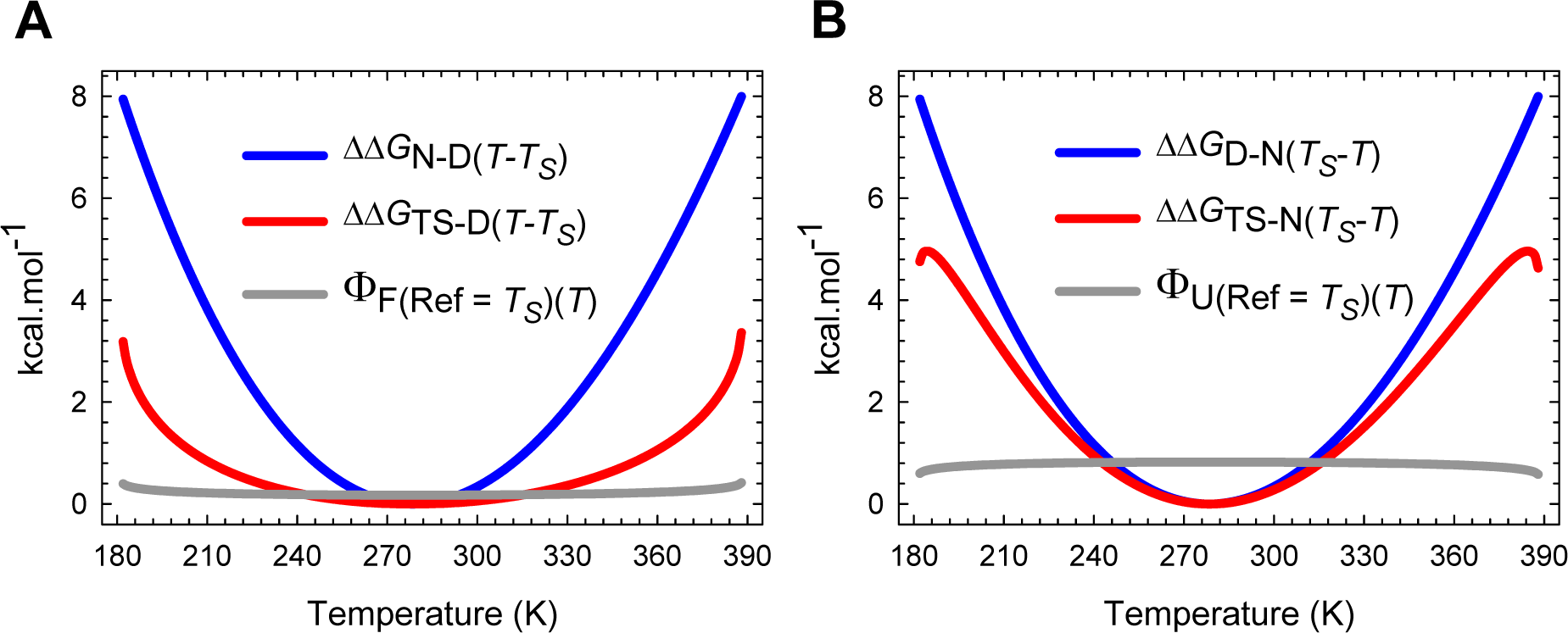
An overlay of transformed curves and Φ_(internal)(*T*)_ when the protein at *T_S_* is defined as the wild type. **(A)** Dividing ΔΔ*G*_TS-D(*T*-*T_S_*)_ by ΔΔ*G*_N-D(*T*-*T_S_*)_ generates Φ_F(internal)(*T*)_ with its minimum at *T_S_*. **(B)** Dividing ΔΔ*G*_TS-N(*T_S_*-*T*)_ by ΔΔ*G*_D-N(*T_S_*-*T*)_ generates Φ_U(internal)(*T*)_ with its maximum at *T_S_*. Note that the dimensions of the ordinate apply only to the red and the blue curves since Φ_(internal)(*T*)_ is dimensionless.

**Figure 21-figure supplement 8.**
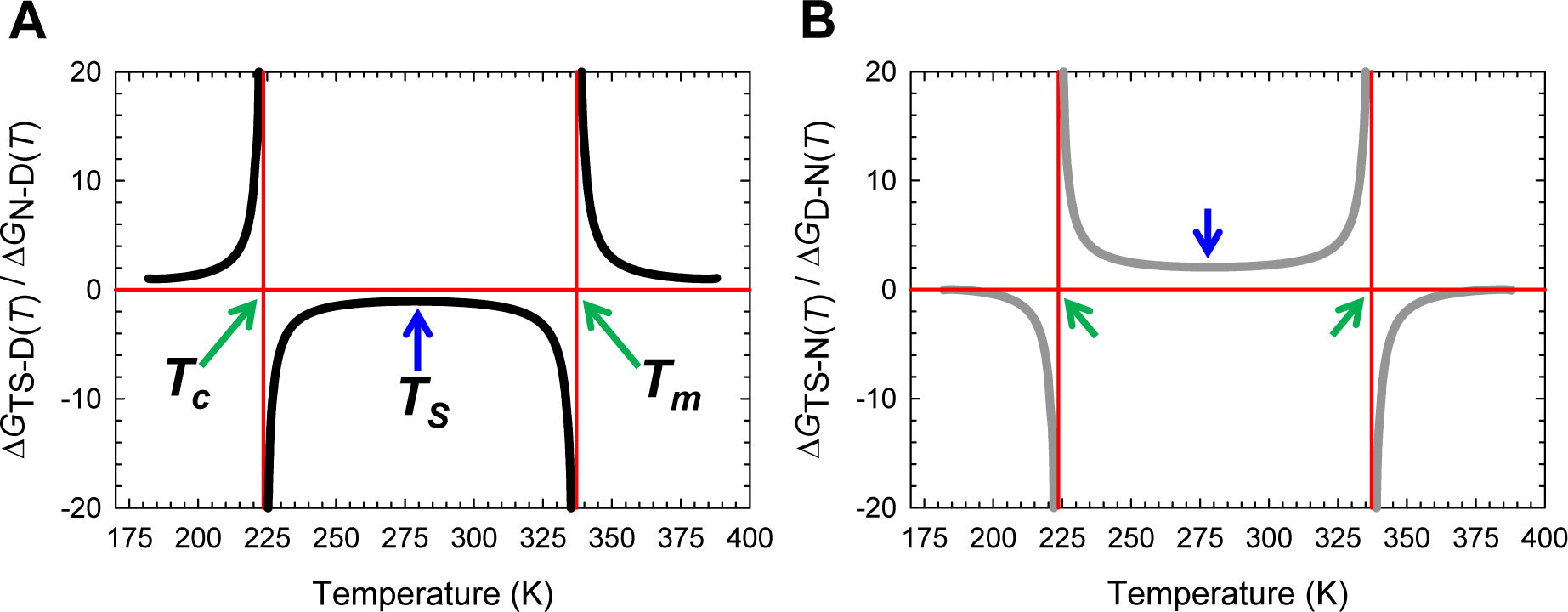
Temperature-dependence of the ratio of the Gibbs activation energies to stability. **(A)** The ratio Δ*G*_TS-D(*T*)_/Δ*G*_N-D(*T*)_ is negative for *T_c_* < *T* < *T_m_* and positive for *T* < *T_c_* and *T* < *T_m_*. **(B)** The ratio Δ*G*_TS-N(*T*)_/Δ*G*_D-N(*T*)_ is positive for *T_c_* < *T* < *T_m_* and negative for *T* < *T_c_* and *T* > *T_m_*. The vertical asymptotes are a consequence of Δ*G*_D-N(*T*)_ = −Δ*G*_N-D(*T*)_ approaching zero as the temperature approaches *T_c_* and *T_m_*. Note that the ordinate is dimensionless, and that (Δ*G*_TS-D(*T*)_/Δ*G*_N-D(*T*)_) + (Δ*G*_TS-N(*T*)_/Δ*G*_D-N(*T*)_) = 1 for a two-state system.

**Figure 21-figure supplement 9.**
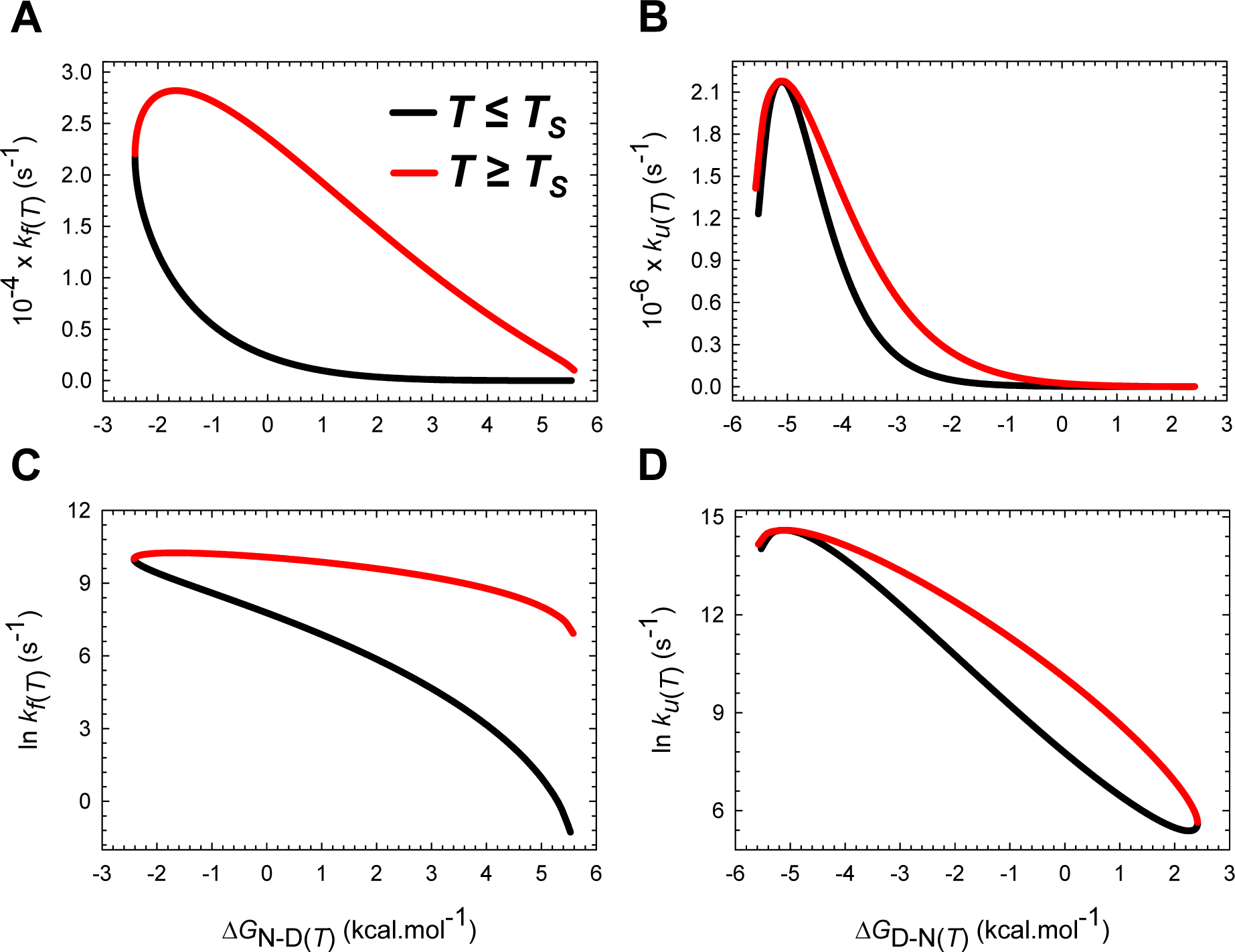
The complex non-linear relationship between the rate constants and the difference in Gibbs energies between the ground states. **(A)** *k*_*f*(*T*)_ *vs* the Gibbs energy of folding at equilibrium. The slope of this plot equals −*k_f(*T*)_* ΔH_TS-D(*T*)_/ΔS_TS-D(*T*)_/ΔS_N-D(*T*)_RT^2^. **(B)** *k*_*u*(*T*)_ *vs* the Gibbs energy of unfolding at equilibrium. The *Marcus-inverted-regimes* which occur at very low and high temperatures are towards the extreme left. The slope of this plot is given by −*k_u(*T*)_* ΔH_TS-D(*T*)_/ΔS_TS-D(*T*)_/ΔS_N-D(*T*)_RT^2^. **(C)** Natural logarithm of *k*_*f*(*T*)_ *vs* the Gibbs energy of folding at equilibrium (slope = -ΔH_TS-D(*T*)_/ΔS_N-D(*T*)_RT^2^). **(D)** Natural logarithm of *k*_*u*(*T*)_ *vs* the Gibbs energy of unfolding at equilibrium (slope = -ΔH_TS-D(*T*)_/ΔS_N-D(*T*)_RT^2^). The abscissae for plots A and C, and plots B and D are identical. The colour-code is identical for all the four plots.

**Figure 22-figure supplement 1.**
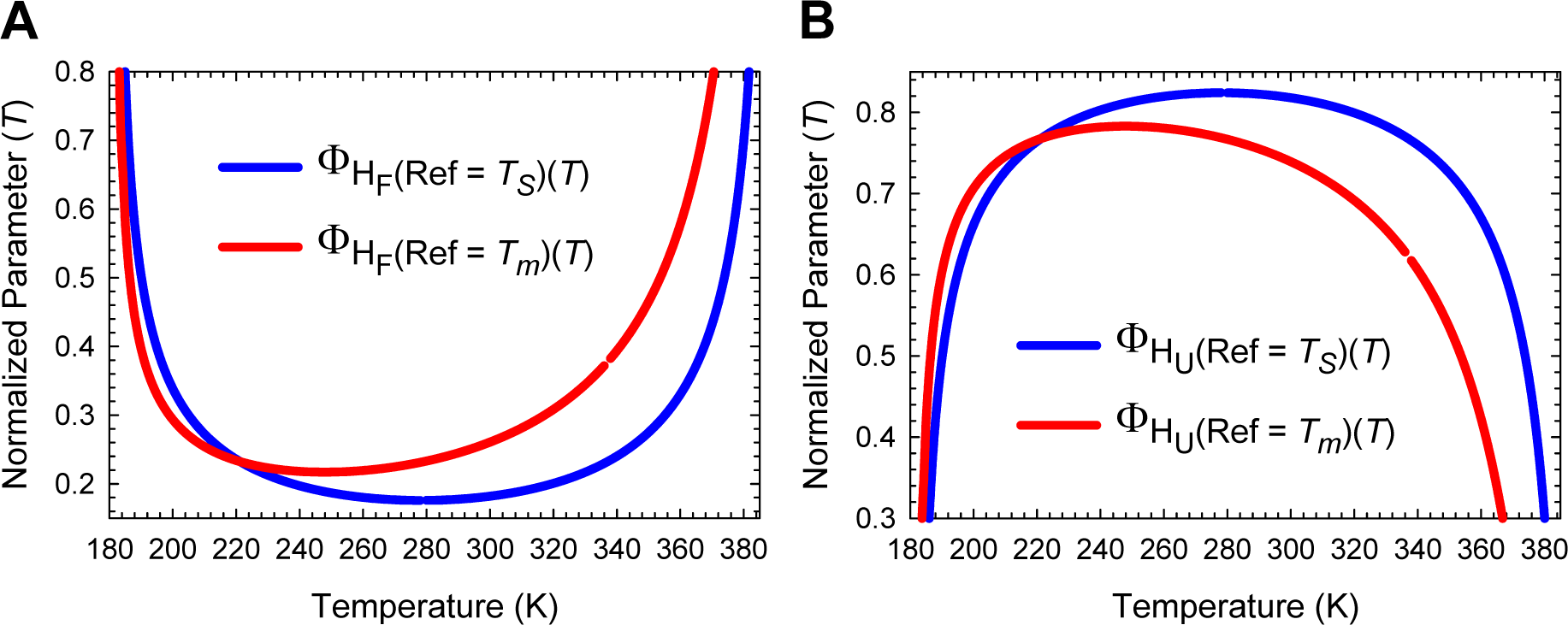
The magnitude of Φ_H(internal)(*T*)_ is dependent on the definition of the wild type. **(A)** A comparison of Φ_H_F_(internal)(*T*)_ calculated using proteins at *T_S_* and *T_m_* as the wild types. **(B)** A comparison of Φ_H_U_(internal)(*T*)_ calculated using proteins at *T_S_* and *T_m_* as the wild types. For the blue curves we have Δ*H*_(wt)_ ≡ Δ*H*_(*TS*)_, and for the red curves, we have Δ*H*_(wt)_ ≡ Δ*H*_(*T_m_*)_. This notation applies to both equilibrium and activation enthalpies. The blue curves are undefined (0/0) at *T_S_*, and the red curves are undefined at *T_c_* and *T_m_*. This figure is the enthalpic equivalent of **Figure 21−figure supplement 1**.

**Figure 22-figure supplement 2.**
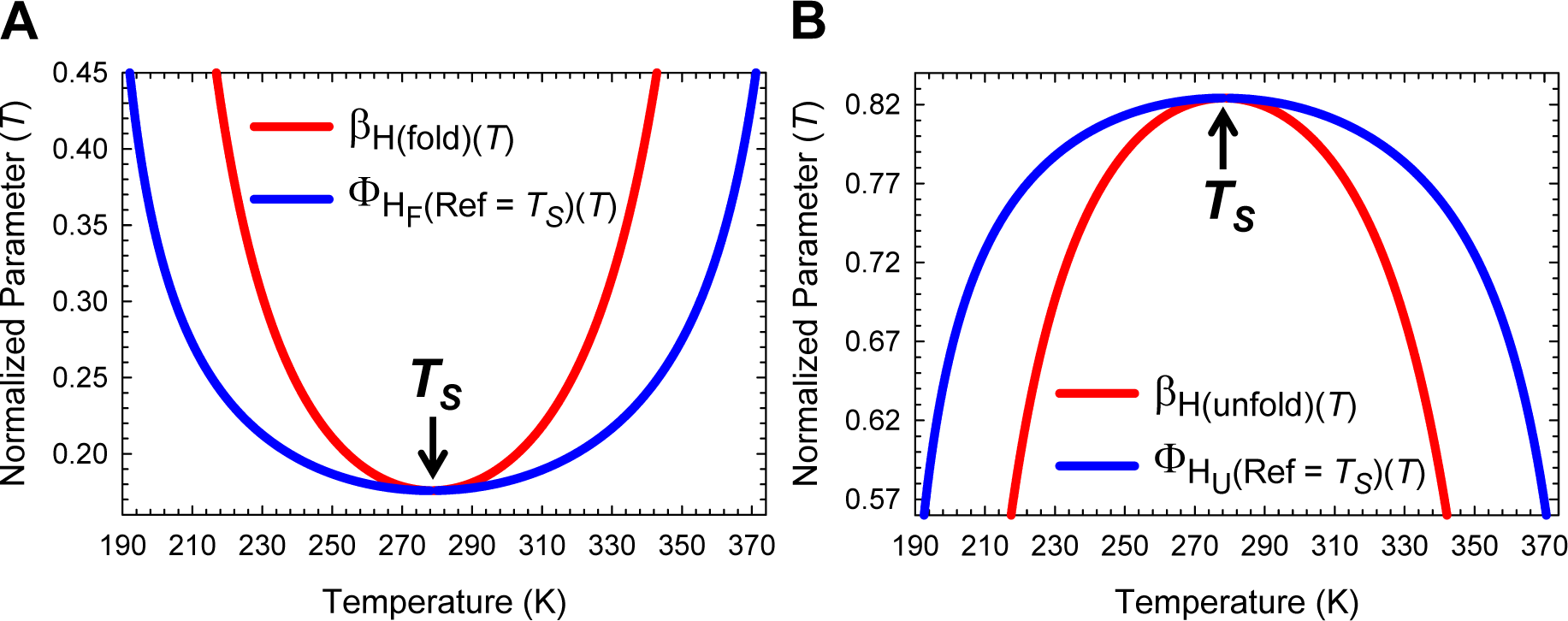
Comparison of the Leffler β_H(*T*)_ and Φ_H(internal)(*T*)_ when the protein at *T_S_* is defined as the wild type. **(A)** β_H(fold)(*T*)_ is almost identical to Φ_H_F_(internal)(*T*)_ around the temperature of maximum stability, but as the temperature deviates from *T_S_*, β_H(fold)(*T*)_ increases far more steeply than Φ_H_F_(internal)(*T*)_, such that for *T* ≠*T_S_* we have β_H(fold)(*T*)_ > Φ_H_F_(internal)(*T*)_. **(B)** β_H(unfold)(*T*)_ is almost identical to Φ_H_U_(internal)(*T*)_ around the temperature of maximum stability, but as the temperature deviates from *T_S_*, β_H(unfold)(*T*)_ decreases far more steeply than Φ_H_U_(internal)(*T*)_, such that for *T* ≠*T_S_* we have β_H(unfold)(*T*)_ < Φ_H_U_(internal)(*T*)_. Note that the parameters Φ_H_F_(internal)(*T*)_ and Φ_H_U_(internal)(*T*)_ are the Fersht-analogues of the Leffler β_H(fold)(*T*)_ and β_H(unfold)(*T*)_, respectively (see heat capacity RC). Consequently, this figure is analogous to a comparison of Leffler β_G(fold)(*T*)_ and Fersht Φ_F(internal)(*T*)_, and Leffler β_G(unfold)(*T*)_ and Fersht Φ_U(internal)(*T*)_ (see **Figure 21−figure supplement 3**).

**Figure 22-figure supplement 3.**
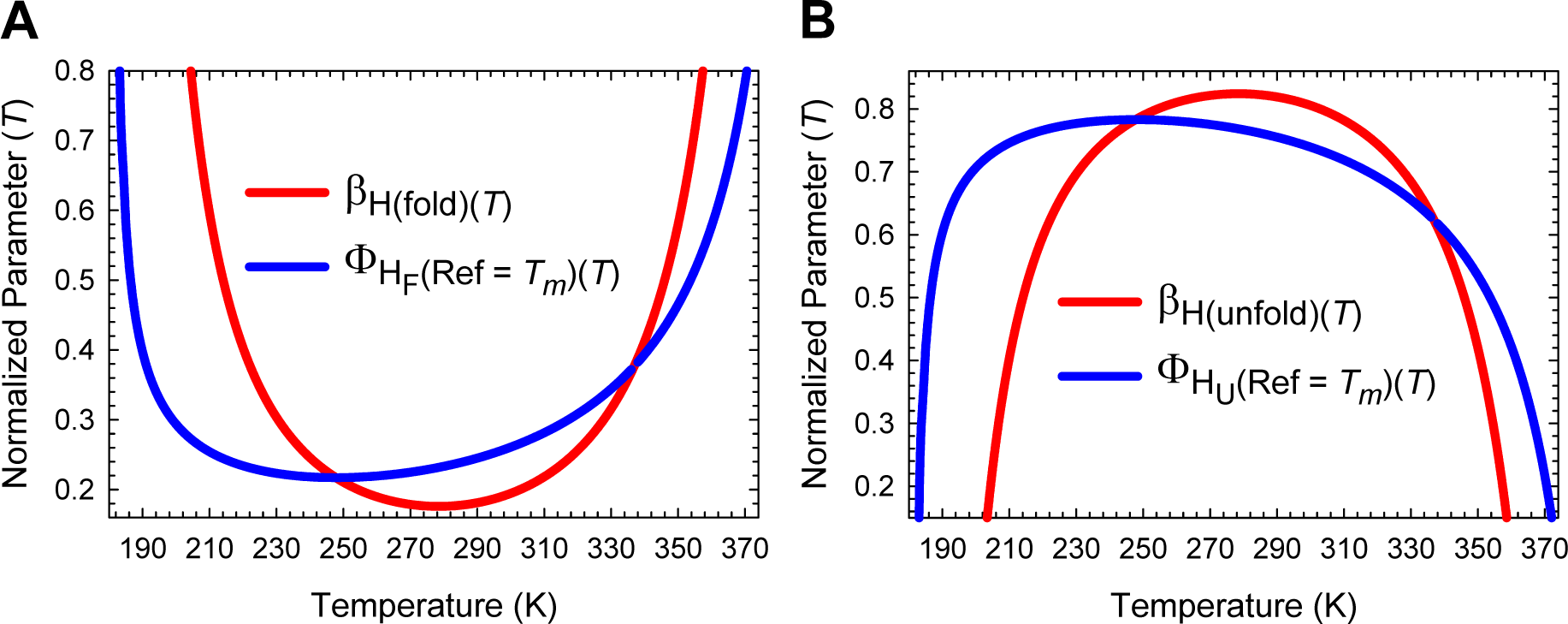
Comparison of the Leffler β_H(*T*)_ and Φ_H(internal)(*T*)_ when the protein at *T_m_* is defined as the wild type. Changing the definition of the wild type from *T_S_* (see previous figure) to *T_m_* has a dramatic effect on the relationship between the Leffler-β_H(*T*)_ and the Fersht-Φ_H(internal)(*T*)_. **(A)** β_H(fold)(*T*)_ < Φ_H_F_(internal)(*T*)_ for ∽248 K < *T* < *T_m_*, β_H(fold)(*T*)_ > Φ_H_F_(internal)(*T*)_ for *T* < ∽248 K and *T* > *T_m_*, and identical when *T* = ∽248 K. **(B)** β_H(unfold)(*T*)_ > Φ_H_U_(internal)(*T*)_ for ∽248 K < *T* < *T_m_*, β_H(unfold)(*T*)_ < Φ_H_U_(internal)(*T*)_ for *T* < ∽248 K and *T* > *T_m_*, and identical when *T* = ∽248 K. Note that Φ_H_F_(internal)(*T*)_ and Φ_H_U_(internal)(*T*)_ are undefined at *T_m_* and the discontinuity in the functions is apparent upon close inspection. This figure is the enthalpic equivalent of **Figure 21−figure supplement 4**.

**Figure 22-figure supplement 4.**
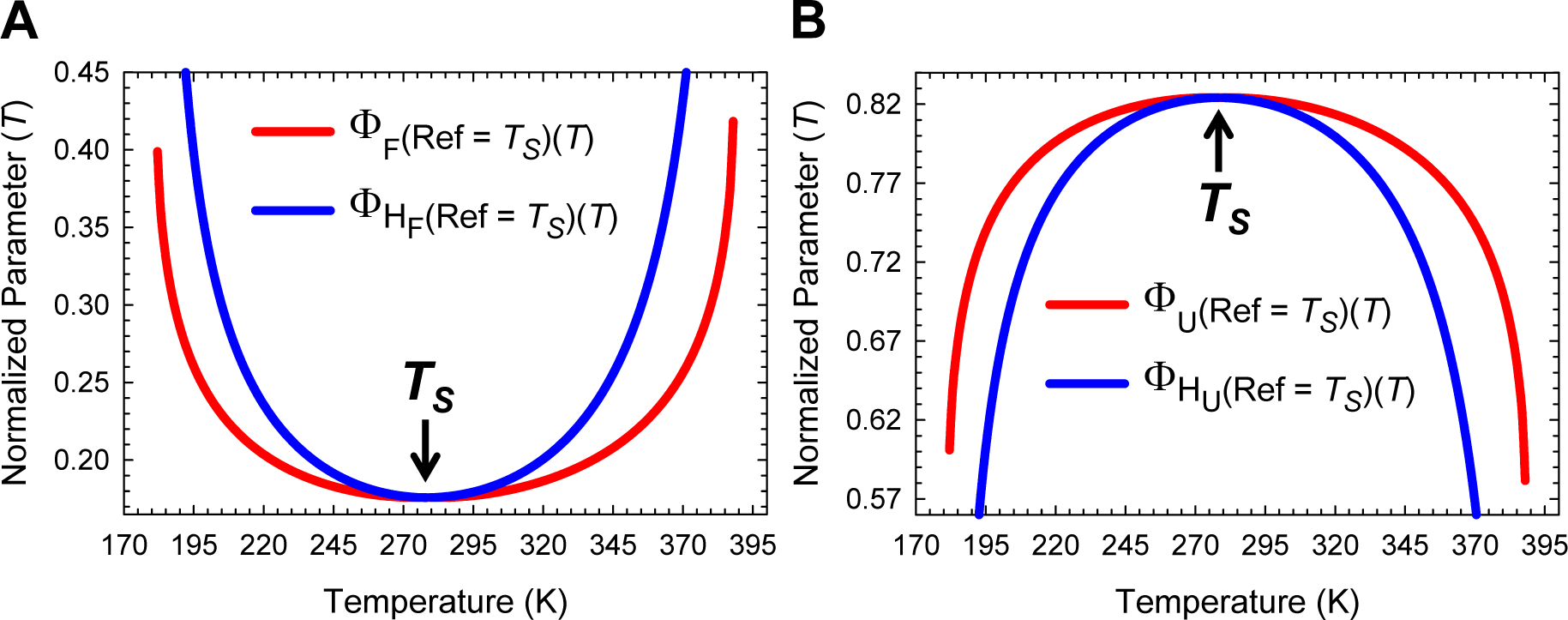
Comparison of Φ_(internal)(*T*)_ and Φ_H(internal)(*T*)_ when the protein at *T_S_* is defined as the wild type. **(A)** The normalized Gibbs parameter Φ_F(internal)(*T*)_ is almost identical to the normalized enthalpic parameter Φ_H_F_(internal)(*T*)_ around the temperature of maximum stability, but as the temperature deviates from *T_S_*, Φ_H_F_(internal)(*T*)_ increases far more steeply than Φ_F(internal)(*T*)_, such that for *T* ≠*T_S_* we have Φ_H_F_(internal)(*T*)_ > Φ_F(internal)(*T*)_. **(B)** The normalized Gibbs parameter Φ_U(internal)(*T*)_ is almost identical to the normalized enthalpic parameter Φ_H_U_(internal)(*T*)_ around the temperature of maximum stability, but as the temperature deviates from *T_S_*, Φ_H_U_(internal)(*T*)_ decreases far more steeply than Φ_U(internal)(*T*)_, such that for *T* ≠*T_S_* we have Φ_H_F_(internal)(*T*)_ < Φ_U(internal)(*T*)_. Since Φ_(internal)(*T*)_ and Φ_H(internal)(*T*)_ are the Fersht-analogues of Leffler β_G(*T*)_ and β_H(*T*)_, respectively, this figure is analogous to comparing the temperature-dependent position of the TSE along the entropic and heat capacity RCs (i.e., a comparison of the temperature-dependence of β_G(*T*)_ and β_H(*T*)_; see **Figure 17**).

**Figure 22-figure supplement 5.**
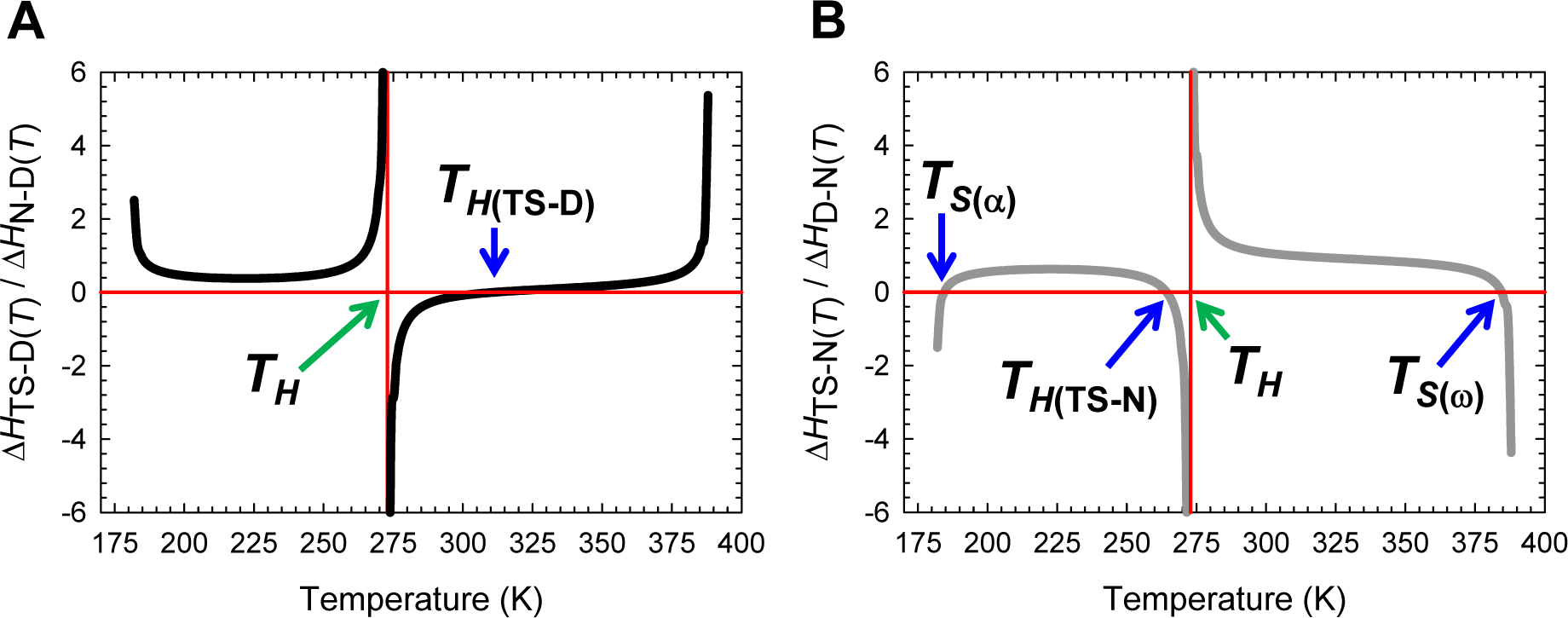
Temperature-dependence of the ratio of the activation enthalpies to equilibrium enthalpies. **(A)** The ratio Δ*H*_TS-D(*T*)_/Δ*H*_N-D(*T*)_ is positive for *T*_α_ ≤ *T* < *T_H_* and *T*_*H*(TS-D)_ < *T* ≤ *T*_ω_, negative for *T_H_* < *T* < *T*_*H*(TS-D)_ and zero at *T*_*H*(TS-D)_. **(B)** The ratio Δ*H*_TS-N(*T*)_/Δ*H*_D-N(*T*)_ is positive for *T*_*S*(α)_ < *T* < *T*_*H*(TS-N)_ and *T_H_* < *T* < *T*_*S*(ω)_, negative for *T*_α_ ≤ *T* < *T*_*S*(α)_, *T*_*H*(TS-N)_ < *T* < *T_H_*, and *T*_*S*(ω)_ < *T* ≤ *T*_ω_, and zero at *T*_*S*(α)_, *T*_*H*(TS-N)_, and *T*_*S*(α)_. The vertical asymptotes are a consequence of Δ*H*_D-N(*T*)_ = −Δ*H*_N-D(*T*)_ approaching zero as *T*→*T_H_*. Note that the ordinate is dimensionless, and that (Δ*H*_*TS*-D(*T*)_/Δ*H*_N-D(*T*)_)+(Δ*H*_*TS*-N(*T*)_/Δ*H*_D-N(*T*)_)=1 for a two-state system.

**Figure 23-figure supplement 1.**
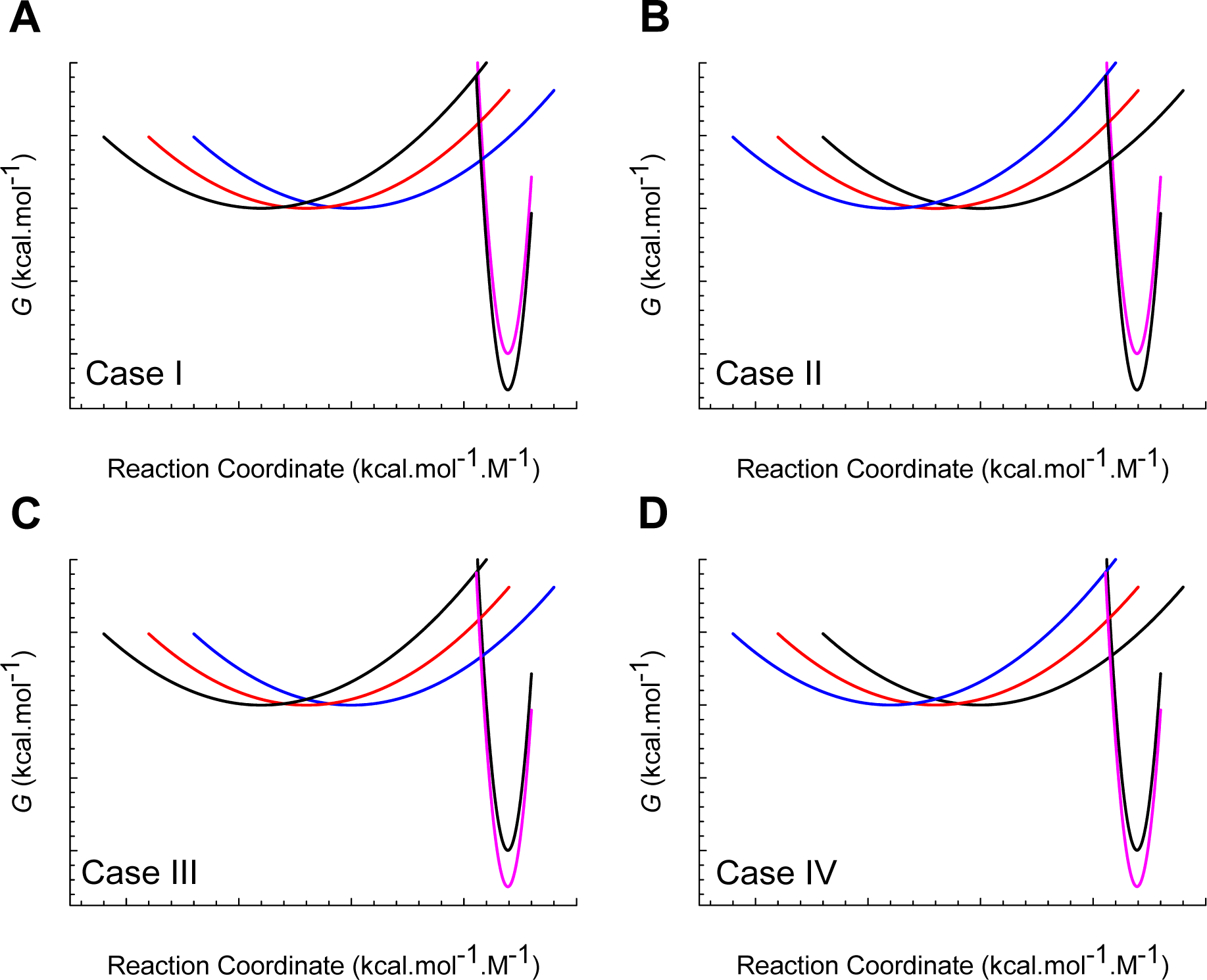
Parabolic Gibbs energy curves to illustrate the effect of concomitant changes in Δ*G*_D-N(*T*)_ and *m*_D-N_ on the position of the TSE along the abscissa and ordinate. The parabolas corresponding to the DSE and NSE of the wild type are shown in black in all the four plots. The mutant DSE-parabolas are shown in blue and red while the mutant NSEparabolas are shown in magenta. **(A)** The introduced mutation causes a concomitant decrease in Δ*G*_D-N(*T*)_ and the mean-length of the RC. **(B)** The introduced mutation causes a decrease in Δ*G*_D-N(*T*)_ but an increase in the mean-length of the RC. **(C)** The introduced mutation stabilizes the mutant but causes a concomitant decrease in the mean-length of the RC. **(D)** The introduced mutation stabilizes the protein but concomitantly causes an increase in the mean-length of the RC. The curvatures of all the DSE-parabolas (α = 1 M^2^.mol.kcal^-1^) and all the NSE-parabolas (ɷ = 30 M^2^.mol.kcal^-1^) are identical.

**Figure 23-figure supplement 2.**
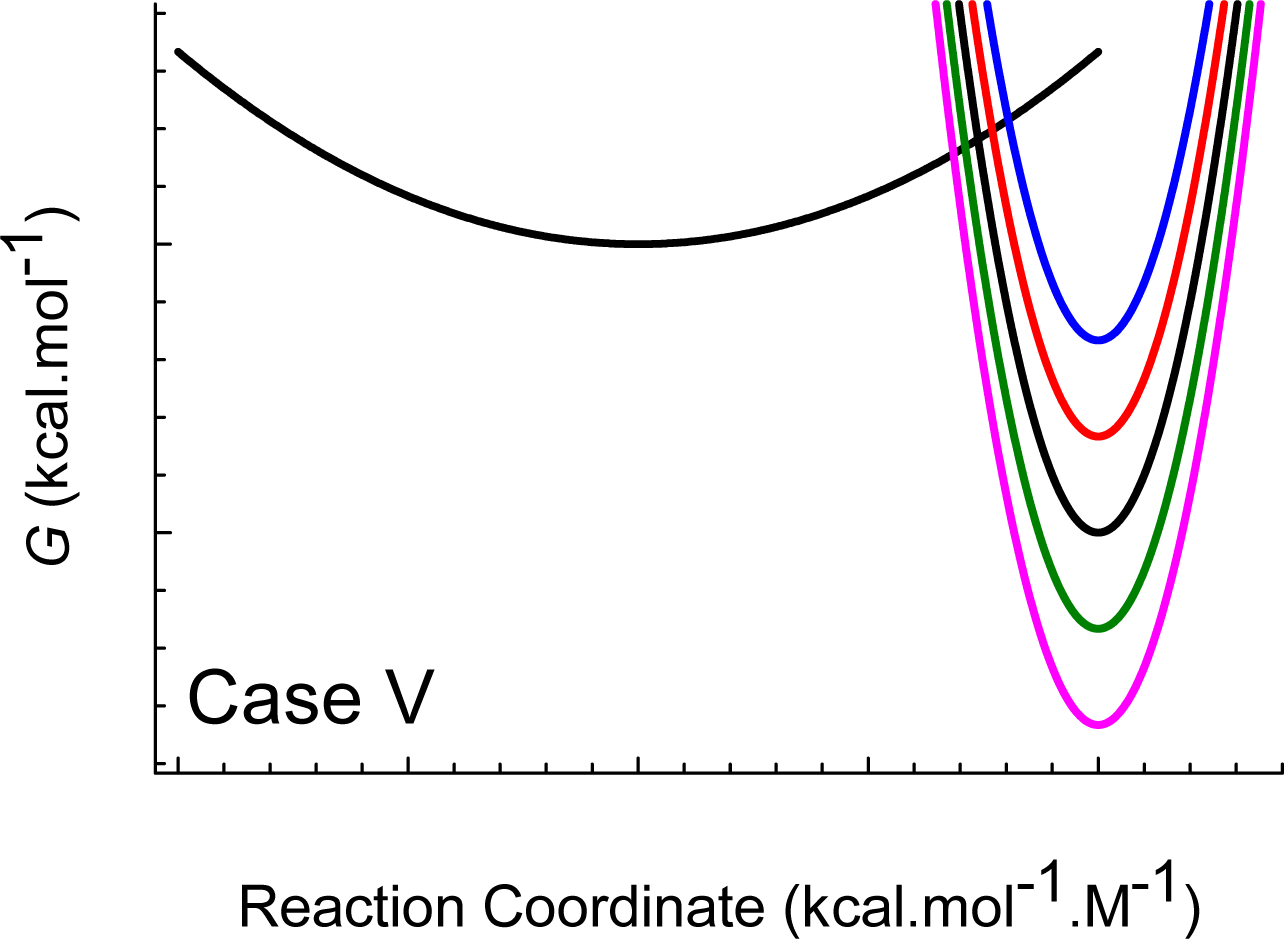
Parabolic Gibbs energy curves to illustrate the effect of a change in Δ*G*_D-N(*T*)_ on the position of the TSE along the abscissa and ordinate. The wild type DSE and NSE-parabolas are shown in black, the destabilized mutants are shown in red and blue, and the stabilized mutants are shown in green and magenta. As the protein is increasingly destabilized the *curve-crossing* along the RC shifts closer to the vertex of the NSE-parabola, and can be due to a stabilized DSE or a destabilized NSE, or both. Conversely, as the protein is increasingly stabilized, the *curve-crossing* along the RC shifts away from the vertex of NSE-parabola and this can be due to a destabilized DSE or a stabilized NSE, or both. The force constant for the DSE-parabola is α = 1 M^2^.mol.kcal^-1^. The curvatures of all the NSE-parabolas are identical (ɷ = 30 M^2^.mol.kcal^-1^).

## REFERENCES

1 Marcus, R. A. Chemical and Electrochemical Electron-Transfer Theory. Annu. Rev. Phys. Chem. 15, 155, 10.1146/annurev.pc.15.100164.001103 (1964).

2 Sade, R. S. Analysis of Two-State Folding Using Parabolic Approximation I: Hypothesis. bioRxiv, 10.1101/036491 (2016).

3 Sade, R. S. Analysis of Two-State Folding Using Parabolic Approximation II: Temperature-Dependence. bioRxiv, 10.1101/037341 (2016).

4 Petrovich, M., Jonsson, A. L., Ferguson, N., Daggett, V. & Fersht, A. R. Φ-Analysis at the Experimental Limits: Mechanism of Beta-Hairpin Formation. J. Mol. Biol. 360, 865–881, 10.1016/j.jmb.2006.05.050 (2006).

5 Tanford, C. Protein Denaturation. C. Theoretical Models for the Mechanism of Denaturation. Adv. Protein Chem. 24, 1–95, 10.1016/S0065-3233(08)60241-7 (1970).

6 Becktel, W. J. & Schellman, J. A. Protein Stability Curves. Biopolymers 26, 1859–1877, 10.1002/bip.360261104 (1987).

7 Privalov, P. L. Thermodynamic Problems of Protein Structure. Annu. Rev. Biophys. Biophys. Chem. 18, 47–69, 10.1146/annurev.bb.18.060189.000403 (1989).

8 Otzen, D. E. & Oliveberg, M. Correspondence between anomalous *m*- and Δ*C_p_-values* in protein folding. Protein Sci. 13, 3253–3263, 10.1110/ps.04991004 (2004).

9 Dimitriadis, G. et al. Microsecond folding dynamics of the F13W G29A mutant of the B domain of staphylococcal protein A by laser-induced temperature jump. Proc. Natl. Acad. Sci. U S A 101, 3809–3814, 10.1073/pnas.0306433101 (2004).

10 Taskent, H., Cho, J. H. & Raleigh, D. P. Temperature-Dependent Hammond Behaviour in a Protein-Folding Reaction: Analysis of Transition-State Movement and Ground-State Effects. J. Mol. Biol. 378, 699–706, 10.1016/j.jmb.2008.02.024 (2008).

11 Cellmer, T., Henry, E. R., Hofrichter, J. & Eaton, W. A. Measuring internal friction of an ultrafast-folding protein. Proc. Natl. Acad. Sci. U S A 105, 18320–18325, 10.1073/pnas.0806154105 (2008).

12 Sun, L., Noel, J. K., Sulkowska, J. I., Levine, H. & Onuchic, J. N. Connecting Thermal and Mechanical Protein (Un)folding Landscapes. Biophys. J. 107, 2941–2952, 10.1016/j.bpj.2014.10.021 (2014).

13 Marcus, R. A. Theoretical Relations among Rate Constants, Barriers, and Brønsted Slopes of Chemical Reactions. J. Phys. Chem. 72, 891–899, 10.1021/j100849a019 (1968).

14 Cohen, A. O. & Marcus, R. A. On the Slope of Free Energy Plots in Chemical Kinetics. J. Phys. Chem. 72, 4249–4256, 10.1021/j100858a052 (1968).

15 Koeppl, G. W. & Kresge, A. J. Marcus Rate Theory and the Relationship between Brønsted Exponents and Energy of Reaction. Chem. Commun., 371–373, 10.1039/C39730000371 (1973).

16 Rasmussen, B. F., Stock, A. M., Ringe, D. & Petsko, G. A. Crystalline ribonuclease A loses function below the dynamical transition at 220 K. Nature 357, 423–424, 10.1038/357423a0 (1992).

17 Kauzmann, W. The Nature of the Glassy State and the Behavior of Liquids at Low Temperatures. Chem. Rev. 43, 219–256, 10.1021/cr60135a002 (1948).

18 Adam, G. & Gibbs, J. H. On the Temperature Dependence of Cooperative Relaxation Properties in Glass Forming Liquids. J. Chem. Phys. 43, 139–146, 10.1063/1.1696442 (1965).

19 Goldstein, M. Viscous Liquids and the Glass Transition: A Potential Energy Barrier Picture. J. Chem. Phys. 51, 3728–3739, 10.1063/1.1672587 (1969).

20 Debenedetti, P. G. & Stillinger, F. H. Supercooled liquids and the glass transition. Nature 410, 259–267, 10.1038/35065704 (2001).

21 Dill, K. A. & Bromberg, S. Molecular Driving Forces – Statistical Thermodynamics in Chemistry and Biology. (Garland Science, 2003).

22 Dyre, J. Mysteries of the Glass Transition. Physics Today 61, 15, 10.1063/1.2835137 (2008).

23 Bauer, T., Lunkenheimer, P. & Loidl, A. Cooperativity and the Freezing of Molecular Motion at the Glass Transition. Phys. Rev. Lett. 111, 225702, 10.1103/PhysRevLett.111.225702 (2013).

24 Kresge, A. J. The Brønsted Relation - Recent Developments. Chem. Soc. Rev. 2, 475–503, 10.1039/CS9730200475 (1973).

25 Nguyen, H., Jäger, M., Moretto, A., Gruebele, M. & Kelly, J. W. Tuning the free-energy landscape of a WW domain by temperature, mutation, and truncation. Proc. Natl. Acad. Sci. U S A 100, 3948–3953, 10.1073/pnas.0538054100 (2003).

26 Ferguson, N. et al. Rapid amyloid fiber formation from the fast-folding WW domain FBP28. Proc. Natl. Acad. Sci. U S A 100, 9814–9819, 10.1073/pnas.1333907100 (2003).

27 Thomsen, J. S. Logical Relations among the Principles of Statistical Mechanics and Thermodynamics. Phys. Rev. 91, 1263–1266, 10.1103/PhysRev.91.1263 (1953).

28 Bryngelson, J. D., Onuchic, J. N., Socci, N. D. & Wolynes, P. G. Funnels, Pathways and the Energy Landscape of Protein Folding: A Synthesis. Proteins: Struct. Funct. Genet. 21, 167, 10.1002/prot.340210302 (1995).

29 Oliveberg, M., Tan, Y. J. & Fersht, A. R. Negative activation enthalpies in the kinetics of protein folding. Proc. Natl. Acad. Sci. U S A 92, 8926–8929, 10.1073/pnas.92.19.8926 (1995).

30 Scalley, M. L. & Baker, D. Protein folding kinetics exhibit an Arrhenius temperature dependence when corrected for the temperature dependence of protein stability. Proc. Natl. Acad. Sci. U S A 94, 10636–10640, 10.1073/pnas.94.20.10636 (1997).

31 Chan, H. S. & Dill, K. A. Protein Folding in the Landscape Perspective: Chevron Plots and Non-Arrhenius Kinetics. Proteins: Struct. Funct. Genet. 30, 2–33, 10.1002/(SICI)1097-0134(19980101)30:1<2::AID-PROT2>3.0.CO;2-R (1998).

32 Ghosh, K., Ozkan, S. B. & Dill, K. A. The Ultimate Speed Limit to Protein Folding Is Conformational Searching. J. Am. Chem. Soc. 129, 11920–11927, 10.1021/ja066785b (2007).

33 Myers, J. K., Pace, C. N. & Scholtz, J. M. Denaturant *m* values and heat capacity changes: Relation to changes in accessible surface areas of protein unfolding. Protein Sci. 4, 2138–2148, 10.1002/pro.5560041020 (1995).

34 Robertson, A. D. & Murphy, K. P. Protein Structure and the Energetics of Protein Stability. Chem. Rev. 97, 1251–1268, 10.1021/cr960383c (1997).

35 Schellman, J. A. Temperature, Stability, and the Hydrophobic Interaction. Biophys. J. 73, 2960–2964, 10.1016/S0006-3495(97)78324-3 (1997).

36 Baldwin, R. L. Dynamic hydration shell restores Kauzmann’s 1959 explanation of how the hydrophobic factor drives protein folding. Proc. Natl. Acad. Sci. U S A 111, 13052–13056, 10.1073/pnas.1414556111 (2014).

37 Privalov, P. L. & Gill, S. J. Stability of Protein Structure and Hydrophobic Interaction. Adv. Protein Chem. 39, 191–234, 10.1016/S0065-3233(08)60377-0 (1988).

38 Gómez, J., Hilser, V. J., Xie, D. & Freire, E. The Heat Capacity of Proteins. Proteins: Struct. Funct. Bioinf. 22, 404–412, 10.1002/prot.340220410 (1995).

39 Shortle, D. The Expanded Denatured State: An Ensemble of Conformations Trapped in a Locally Encoded Topological Space. Adv. Protein Chem. 62, 1–23, 10.1016/S0065-3233(02)62003-0 (2002).

40 Bowler, B. E. Residual structure in unfolded proteins. Curr. Opin. Struct. Biol. 22, 4–13, 10.1016/j.sbi.2011.09.002 (2012).

41 Richards, F. M. Areas, Volumes, Packing, and Protein Structure. Annu. Rev. Biophys. Bioeng. 6, 151–176, 10.1146/annurev.bb.06.060177.001055 (1977).

42 Bernadó, P., Blackledge, M. & Sancho, J. Sequence-Specific Solvent Accessibilities of Protein Residues in Unfolded Protein Ensembles. Biophys. J. 91, 4536–4543, 10.1529/biophysj.106.087528 (2006).

43 Gong, H. & Rose, G. D. Assessing the solvent-dependent surface area of unfolded proteins using an ensemble model. Proc. Natl. Acad. Sci. U S A 105, 3321–3326, 10.1073/pnas.0712240105 (2008).

44 Behe, M. J., Lattman, E. E. & Rose, G. D. The protein-folding problem: The native fold determines packing, but does packing determine the native fold? Proc. Natl. Acad. Sci. U S A 88, 4195–4199, 10.1073/pnas.88.10.4195 (1991).

45 Chothia, C. Principles that Determine the Structure of Proteins. Annu. Rev. Biochem. 53, 537–572, 10.1146/annurev.bi.53.070184.002541 (1984).

46 Miller, S., Janin, J., Lesk, A. M. & Chothia, C. Interior and Surface of Monomeric Proteins. J. Mol. Biol. 196, 641–656, 10.1016/0022-2836(87)90038-6 (1987).

47 Tan, Y. J., Oliveberg, M. & Fersht, A. R. Titration Properties and Thermodynamics of the Transition State for Folding: Comparison of Two-state and Multi-state Folding Pathways. J. Mol. Biol. 264, 377–389, 10.1006/jmbi.1996.0647 (1996).

48 Leffler, J. Parameters for the Description of Transition States. Science 117, 340–341, 10.1126/science.117.3039.340 (1953).

49 Sanchez, I. E. & Kiefhaber, T. Non-linear rate-equilibrium free energy relationships and Hammond behavior in protein folding. Biophys. Chem. 100, 397–407, 10.1016/S0301-4622(02)00294-6 (2003).

50 Bilsel, O. & Matthews, C. R. Barriers in Protein Folding Reactions. Adv. Protein Chem. 53, 153–207, 10.1016/S0065-3233(00)53004-6 (2000).

51 Gloss, L. M. & Matthews, C. R. The Barriers in the Bimolecular and Unimolecular Folding Reactions of the Dimeric Core Domain of Escherichia Coli Trp Repressor are Dominated by Enthalpic Contributions. Biochemistry 37, 16000–16010, 10.1021/bi981694f (1998).

52 Lee, A. L. & Wand, A. J. Microscopic origins of entropy, heat capacity and the glass transition in proteins. Nature 411, 501–504, 10.1038/35078119 (2001).

53 Chandler, D. Interfaces and the driving force of hydrophobic assembly. Nature 437, 640–647, 10.1038/nature04162 (2005).

54 Ben-Naim, A. Myths and verities in protein folding theories: From Frank and Evans iceberg-conjecture to explanation of the hydrophobic effect. J. Chem. Phys. 139, 165105, 10.1063/1.4827086 (2013).

55 Sturtevant, J. M. Heat capacity and entropy changes in processes involving proteins. Proc. Natl. Acad. Sci. U S A 74, 2236–2240, 10.1073/pnas.74.6.2236 (1977).

56 Lazaridis, T. & Karplus, M. Heat capacity and compactness of denatured proteins. Biophys. Chem. 78, 207–217, 10.1016/S0301-4622(99)00022-8 (1999).

57 Frauenfelder, H. et al. A unified model of protein dynamics. Proc. Natl. Acad. Sci. U S A 106, 5129–5134, 10.1073/pnas.0900336106 (2009).

58 Lewandowski, J. R., Halse, M. E., Blackledge, M. & Emsley, L. Direct observation of hierarchical protein dynamics. Science 348, 578–581, 10.1126/science.aaa6111 (2015).

59 Cooper, A. Heat capacity of hydrogen-bonded networks: an alternative view of protein folding thermodynamics. Biophys. Chem. 85, 25–39, 10.1016/S0301-4622(00)00136-8 (2000).

60 Cooper, A. Heat capacity effects in protein folding and ligand binding: a re-evaluation of the role of water in biomolecular thermodynamics. Biophys. Chem. 115, 89–97, 10.1016/j.bpc.2004.12.011 (2005).

61 Cooper, A. Protein Heat Capacity: An Anomaly that Maybe Never Was. J. Phys. Chem. Lett. 1, 3298–3304, 10.1021/jz1012142 (2010).

62 Karplus, M., Ichiye, T. & Pettitt, B. M. Configurational Entropy of Native Proteins. Biophys. J. 52, 1083–1085, 10.1016/S0006-3495(87)83303-9 (1987).

63 Finkelstein, A. V. & Ptitsyn, O. B. Protein Physics: A Course of Lectures. (Academic Press, 2002).

64 Dill, K. A. Dominant Forces in Protein Folding. Biochemistry 29, 7133–7155, 10.1021/bi00483a001 (1990).

65 Privalov, P. L. & Makhatadze, G. I. Contribution of Hydration and Non-covalent Interactions to the Heat Capacity Effect on Protein Unfolding. J. Mol. Biol. 224, 715–723, 10.1016/0022-2836(92)90555-X (1992).

66 Privalov, P. L. & Makhatadze, G. I. Contribution of Hydration to Protein Folding Thermodynamics: II. The Entropy and Gibbs Energy of Hydration. J. Mol. Biol. 232, 660–679, 10.1006/jmbi.1993.1417 (1993).

67 Makhatadze, G. I. & Privalov, P. L. Contribution of Hydration to Protein Folding Thermodynamics I. The Enthalpy of Hydration. J. Mol. Biol. 232, 639–659, 10.1006/jmbi.1993.1416 (1993).

68 Hammond, G. S. A Correlation of Reaction Rates. J. Am. Chem. Soc. 77, 334–338, 10.1021/ja01607a027 (1955).

69 Hammett, L. P. Linear free energy relationships in rate and equilibrium phenomena. Trans. Faraday Soc. 34, 156–165, 10.1039/TF9383400156 (1938).

70 Wells, P. R. Linear Free Energy Relationships. Chem. Rev. 63, 171–219, 10.1021/cr60222a005 (1963).

71 Farcasiu, D. The use and misuse of the Hammond Postulate. J. Chem. Edu. 52, 76, 10.1021/ed052p76 (1975).

72 Agmon, N. Quantitative Hammond postulate. J. Chem. Soc., Faraday Trans. 2 74, 388–404, 10.1039/f29787400388 (1978).

73 Albery, W. J. The Application of the Marcus Relation to Reactions in Solution. Annu. Rev. Phys. Chem. 31, 227–263, 10.1146/annurev.pc.31.100180.001303 (1980).

74 Formosinho, S. J. On the Validity of Linear Free Energy Relationships in Chemical Kinetics. Rev. Port. Quim. 27, 521 (1985).

75 Estell, D. A. Artifacts in the application of linear free energy analysis. Protein Eng. 1, 441–442, 10.1093/protein/1.6.441 (1987).

76 Fersht, A. R. Linear free energy relationships are valid! Protein Eng. 1, 442–445, 10.1093/protein/1.6.442 (1987).

77 Sanchez, I. E. & Kiefhaber, T. Hammond behavior versus ground state effects in protein folding: evidence for narrow free energy barriers and residual structure in unfolded states. J. Mol. Biol. 327, 867–884, 10.1016/S0022-2836(03)00171-2 (2003).

78 Straub, J. E. & Karplus, M. The interpretation of site-directed mutagenesis experiments by linear free energy relations. Protein Eng. 3, 673–675, 10.1093/protein/3.8.673 (1990).

79 Fersht, A. R., Matouschek, A. & Serrano, L. The Folding of an Enzyme I. Theory of Protein Engineering Analysis of Stability and Pathway of Protein Folding. J. Mol. Biol. 224, 771–782, 10.1016/0022-2836(92)90561-W (1992).

80 Fersht, A. R. & Daggett, V. Protein Folding and Unfolding at Atomic Resolution. Cell 108, 573–582, 10.1016/S0092-8674(02)00620-7 (2002).

81 Kresge, A. J. in Proton-Transfer Reactions (eds Edward Caldin & Victor Gold) Ch. 7: The Brønsted Relation: Significance of the Exponent, 179–199 (Springer US, 1975).

82 Crane, J. C., Koepf, E. K., Kelly, J. W. & Gruebele, M. Mapping the Transition State of the WW Domain Beta-Sheet. J. Mol. Biol. 298, 283–292, 10.1006/jmbi.2000.3665 (2000).

83 Jager, M., Nguyen, H., Crane, J. C., Kelly, J. W. & Gruebele, M. The Folding Mechanism of a Beta-Sheet: The WW Domain. J. Mol. Biol. 311, 373–393, 10.1006/jmbi.2001.4873 (2001).

84 Ervin, J. & Gruebele, M. Quantifying protein folding transition States with Φ_T_. J. Biol. Phys. 28, 115–128, 10.1023/A:1019930203777 (2002).

85 Chung, H. S. & Tokmakoff, A. Temperature-dependent downhill unfolding of ubiquitin. I. Nanosecond-to-millisecond resolved nonlinear infrared spectroscopy. Proteins: Struct. Funct. Bioinf. 72, 474–487, 10.1002/prot.22043 (2008).

86 Johnson, C. D. Linear Free Energy Relations and the Reactivity-Selectivity Principle. Chem. Rev. 75, 755–765, 10.1021/cr60298a004 (1975).

87 Bordwell, F. G., Boyle, W. J., Hautala, J. A. & Yee, K. C. Brønsted Coefficients Larger Than 1 and less than 0 for Proton Removal from Carbon Acids. J. Am. Chem. Soc. 91, 4002–4003, 10.1021/ja01042a082 (1969).

88 Marcus, R. A. Unusual Slopes of Free Energy Plots in Kinetics. J. Am. Chem. Soc. 91, 7224–7225, 10.1021/ja01054a003 (1969).

89 Kresge, A. J. The Nitroalkane Anomaly. Can. J. Chem. 52, 1897–1903, 10.1139/v74-270 (1974).

90 Shapiro, I. O., Zharova, N. G., Ranneva, Y. I., Terekhova, M. I. & Shatenshtein, A. I. Reasons for the “Alpha” Coefficient Anomaly in the Brønsted Relation for the Ionization of CH Acids. Theor. Expt. Chem. 22, 425–430, 10.1007/BF00523820 (1987).

91 Würthwein, E.-U., Lang, G., Schappele, L. H. & Mayr, H. Rate-Equilibrium Relationships in Hydride Transfer Reactions: The Role of Intrinsic Barriers. J. Am. Chem. Soc. 124, 4084–4092, 10.1021/ja0121540 (2002).

92 Fersht, A. R. Relationship of Leffler (Brønsted) alpha values and protein folding Phi values to position of transition-state structures on reaction coordinates. Proc. Natl. Acad. Sci. U S A 101, 14338–14342, 10.1073/pnas.0406091101 (2004).

93 Gillespie, J. R. & Shortle, D. Characterization of Long-range Structure in the Denatured State of Staphylococcal Nuclease. I. Paramagnetic Relaxation Enhancement by Nitroxide Spin Labels. J. Mol. Biol. 268, 158–169, 10.1006/jmbi.1997.0954 (1997).

94 Shortle, D. & Ackerman, M. S. Persistence of Native-Like Topology in a Denatured Protein in 8 M Urea. Science 293, 487–489, 10.1126/science.1060438 (2001).

95 Klein-Seetharaman, J. et al. Long-range Interactions Within a Nonnative Protein. Science 295, 1719–1722, 10.1126/science.1067680 (2002).

96 Bernadó, P. et al. A structural model for unfolded proteins from residual dipolar couplings and small-angle x-ray scattering. Proc. Natl. Acad. Sci. U S A 102, 17002–17007, 10.1073/pnas.0506202102 (2005).

97 Goldenberg, D. P. Finding the right fold. Nat. Struct. Biol. 6, 987–990, 10.1038/14866 (1999).

98 Sanchez, I. E. & Kiefhaber, T. Origin of Unusual Phi-Values in Protein Folding: Evidence Against Specific Nucleation Sites. J. Mol. Biol. 334, 1077–1085, 10.1016/j.jmb.2003.10.016 (2003).

99 Weikl, T. R. & Dill, K. A. Transition-States in Protein Folding Kinetics: The Structural Interpretation of Φ values. J. Mol. Biol. 365, 1578–1586, 10.1016/j.jmb.2006.10.082 (2007).

100 Naganathan, A. N. & Munoz, V. Insights into protein folding mechanisms from large scale analysis of mutational effects. Proc. Natl. Acad. Sci. U S A 107, 8611–8616, 10.1073/pnas.1000988107 (2010).

101 Booth, P. J. & Clarke, J. Membrane protein folding makes the transition. Proc. Natl. Acad. Sci. U S A 107, 3947–3948, 10.1073/pnas.0914478107 (2010).

102 Fersht, A. R. & Sato, S. Phi-value analysis and the nature of protein-folding transition states. Proc. Natl. Acad. Sci. U S A 101, 7976–7981, 10.1073/pnas.0402684101 (2004).

103 LiCata, V. J. & Liu, C.-C. Analysis of Free Energy Versus Temperature Curves in Protein Folding and Macromolecular Interactions. Methods Enzymol. 488, 219–238, 10.1016/B978-0-12-381268-1.00009-4 (2011).

104 Fersht, A. Structure and Mechanism in Protein Science: A Guide to Enzyme Catalysis and Protein Folding. (W.H. Freeman and Company, 1999).

